# Genetically Encoded, Multivalent Liquid Glycan Array (LiGA)

**DOI:** 10.1101/2020.03.24.997536

**Authors:** Mirat Sojitra, Susmita Sarkar, Jasmine Maghera, Emily Rodrigues, Eric Carpenter, Shaurya Seth, Daniel Ferrer Vinals, Nicholas Bennett, Revathi Reddy, Amira Khalil, Xiaochao Xue, Michael Bell, Ruixiang Blake Zheng, Ping Zhang, Corwin Nycholat, Chang-Chun Ling, Todd L. Lowary, James C. Paulson, Matthew S. Macauley, Ratmir Derda

## Abstract

The Central Dogma of Biology does not allow for the study of glycans using DNA sequencing. We report a “Liquid Glycan Array” (LiGA) platform comprising a library of DNA ‘barcoded’ M13 virions that display 30-1500 copies of glycans per phage. A LiGA is synthesized by acylation of phage pVIII protein with a dibenzocyclooctyne, followed by ligation of azido-modified glycans. Pulldown of the LiGA with lectins followed by deep sequencing of the barcodes in the bound phage decodes the optimal structure and density of the recognized glycans. The LiGA is target agnostic and can measure the glycan-binding profile of lectins such as CD22 on cells *in vitro* and immune cells in a live mouse. From a mixture of multivalent glycan probes, LiGAs identifies the glycoconjugates with optimal avidity necessary for binding to lectins on living cells *in vitro* and *in vivo*; measurements that cannot be performed with canonical glass slide-based glycan arrays.

**Dedication:** The paper is dedicated to Laura L. Kiessling on the occasion of her 60^th^ birthday.

## Introduction

The Central Dogma of Biology, DNA → RNA → Protein, facilitates the study of these biopolymers using a unified toolbox of next-generation DNA sequencing^1^. This ability has revolutionized and transformed all areas of biomedical and life science^2^. In contrast, investigating the biological roles of carbohydrates cannot rely on DNA sequencing directly. Akin to the DNA microarrays used in the 1990s, glycan arrays – made by printing carbohydrates on distinct locations on a glass surface^3^ – are a high throughput approach for identifying carbohydrates that bind to glycan binding proteins (GBPs)^4-8^. The information provided by glycan arrays, a glycan binding profile for a receptor, is a critical starting point for downstream fundamental applications such as improved design of inhibitors, vaccines and therapeutics^9-15^. DNA arrays have been largely replaced by *de novo* analysis of DNA by deep sequencing^16^. A technology that allows the one-to-one correspondence between a DNA sequence and a carbohydrate structure would enable application of powerful deep sequencing approaches to identify the binding profiles of GBPs in cells *in vitro* and *in vivo*.

Valency and spatial presentation of glycans are critical variables in glycan–GBP interactions^17-19^. Traditional ‘solid’ glycan arrays mimic the multivalent nature of these interactions^20, 21^, and the valency of glycans on the surface can be adjusted^22-24^. Indeed, the density of the multivalent display influences both the affinity and specificity of binding for lectins, carbohydrate processing enzymes, and antibodies^3, 24, 25^. For example, systematic variation of glycan density on microarrays enables differentiating subpopulations of serum antibodies that are not detectable using a single glycan density^24^. The solid-phase nature of canonical glycans arrays, however, fundamentally limits the ability to study the cross talk and dynamic competition between multiple glycans with a GBP. The format has also demonstrated limited utility for investigating cell-surface GBPs and is incompatible with *in vivo* studies. Ligation of glycans to DNA^26-32^ or peptide-nucleic-acid (PNA)^33^ is possible, but the monovalent display on DNA cannot mimic the multivalent presentation of glycans, a critical requirement because the very large majority of GBPs recognize their biologically relevant ligands with weak affinity. Tri- and tetravalent displays of glycans on RNA^34^ and recently DNA^35^, have been reported. However, investigating protein-glycan interactions in a cellular milieu and *in vivo* would place such platforms at risk of degradation by nucleases, which are ubiquitous.

One of the most successful platforms for investigating interactions between glycans and cell surface GBPs *ex vivo* and *in vivo* are glycan-decorated liposomal nanoparticles^36, 37^ and viral capsids^38^. However, neither technology permits encoding or tracking of different glycan structures. The multivalent presentation of different glycans was reported on micron-size beads encoded via the Luminex® strategy^39-41^ or encoded chemical tags^42^, but cell-binding assays and *in vivo* applications, which would necessitate injection of microbeads, have not been reported. This summary of these state-of the art technologies leads to the ideal platform for glycan arrays: it should display glycans on a multivalent, monodisperse carrier of sub-micron size equipped with an internal DNA barcode. This carrier must be stable and non-interfering with binding assays *in vitro*, in cell culture or *in vivo*. The internal location of the barcode would protect the DNA from nucleases and would also prevent undesired interactions between the coding element and other biological species. To satisfy these design elements, we describe here a liquid glycan array (LiGA) format built on M13 phage particles with silent DNA barcodes^43^ inside the phage genome. The LiGA technology combines the multivalent presentation of traditional ‘solid’ or bead-based arrays and liposome constructs with a soluble format and DNA encoding of both the composition and density of glycans. The components of a LiGA—a multivalent display of chemically-defined synthetic glycans—are fundamentally different from another reported glycan-functionalized phage: ‘glycophage display’^44-46^, which was obtained via the heterologous expression of the *Campylobacter jejuni* glycosylation machinery in *E. coli*. LiGAs retain most of the benefits of other M13 phage-displayed libraries, importantly genetic-encoding and robust protection of the DNA message inside the virion, which has allowed successful profiling of ligand–receptor interactions on the surface of cells, organs in live animals^47, 48^, and in humans^49, 50^.

## Results

### Construction of the LiGA building blocks: glycan–phage conjugates

Although many types of phage and viruses have been used for genetically-encoded display, we selected the M13 platform^51^ as a scalable, stable platform, compatible with a wide range of cloning methods, as well as chemical and enzymatic conjugation strategies. As we show here, while the M13 virion is not naturally glycosylated, desired glycans can be chemically conjugated to a subset of the 2700 copies of the major coat protein pVIII to produce a multivalent display of ∼200–1500 glycans on the 700 × 5 nm virion, creating a phage population with a single glycan composition. For DNA-encoding, the previously reported silent barcode^43^ cloned near the pIII cloning site of vector M13KE can be used (Supplementary Fig. S1a,b). This strategy allows the assembly of a glycan library of phage with distinct DNA encoded barcodes, each with a different glycan. Cloning of a degenerate nucleotide sequences with 10^10^ theoretically-possible combinations into the N-terminal region of protein pIII yielded a library of ‘silently’ barcoded vectors that encode chemically-identical phage particles. After serial propagation of the library to remove the unfit clones, we subcloned and amplified separate populations of phage particles that contained DNA barcode sequences separated by least three nucleotide substitutions (Supplementary Table S1). This design allowed us to correct any mistakes during DNA deep sequencing (Supplementary Fig. S1).

As the glycan source, we used synthetic glycans with an alkyl-azido linker, which are common intermediates in oligosaccharide synthesis; 63 glycans functionalized in this way were obtained as part of the public Consortium for Functional Glycomics (CFG) collection^52^, and an additional 12 azido-glycans were synthesized in this report (Supplementary Schemes S1–S5) or previous reports (see Supplementary Table S2 for list of glycans). We first tested a reported^38^ ligation of azido glycans by Cu-Activated Azido Alkyne Cycloaddition (CuAAC) to phage particles acylated by *N*-succinimidyl 4-pentynoate; however, CuAAC reduced the number of infective M13 particles indicating damage to either DNA or the capsid of phage (Supplementary Fig. S2). We tested Strain Promoted Azido Alkyne Cycloaddition^53^ (SPAAC) as an alternative (Fig. 1a) and, importantly, SPAAC ligation did not decrease the number of infectious particles (Supplementary Fig. S2c,d). Acylation of the N-terminus of the pVIII with dibenzocyclooctyne-*N*-hydroxysuccinimidyl ester (DBCO-NHS) followed by SPAAC ligation could be monitored by MALDI-TOF mass spectrometry (Fig. 1b). As the spacing of pVIII on the surface of M13 phage is well defined, the degree of conjugation would be expected to influence binding to antibodies or glycan binding proteins (Fig. 1c). The peak intensities in MALDI yielded a semi-quantitative estimate of the densities of the glycans on phage; such analysis was reliable down to ∼10^7^ phage particles (Supplementary Fig. S3). Control of the concentration of DBCO–NHS (Fig. 1d) and reaction conditions (Supplementary Fig. S4) enabled modification from 1% to 50% of pVIII proteins, yielding phage particles with 30–1400 copies of glycan per virion. For example, 0.5–1.5 mM of DBCO led to acylation of 15–50% of the pVIII population (Fig. 1d). Subsequent addition of 2 mM azido-glycan quantitatively consumed the alkyne and ligated the glycan to pVIII as evidenced by the pVIII–DBCO–glycan signal and concomitant disappearance of pVIII– DBCO signal (Fig. 1d). Quality control by MALDI confirmed the conjugation of diverse glycan structures (Fig. 1e) and detected rare problems in reactivity: for example, the use of glycosyl azides in the SPAAC yielded suboptimal coupling leaving unreactive pVIII-DBCO even after 24 h reaction time (Fig. 1b). This problem was resolved by introducing a linker between the glycan and the azido group (Fig. 1d,e). When the density of modification exceeded 50%, we observed that ∼5% of pVIII proteins contained two modifications per protein (Supplementary Fig. S5). To assess the regioselectivity of the acylation, we took advantage of the known spontaneous cleavage of the Asp7–Pro8 bond in pVIII by TFA in a sinapinic acid matrix^54^. We observed that the amide bond is formed predominantly at an N-terminal amine and not at the Lys10 of the pVIII sequence (Supplementary Fig. S6).

**Fig. 1:**
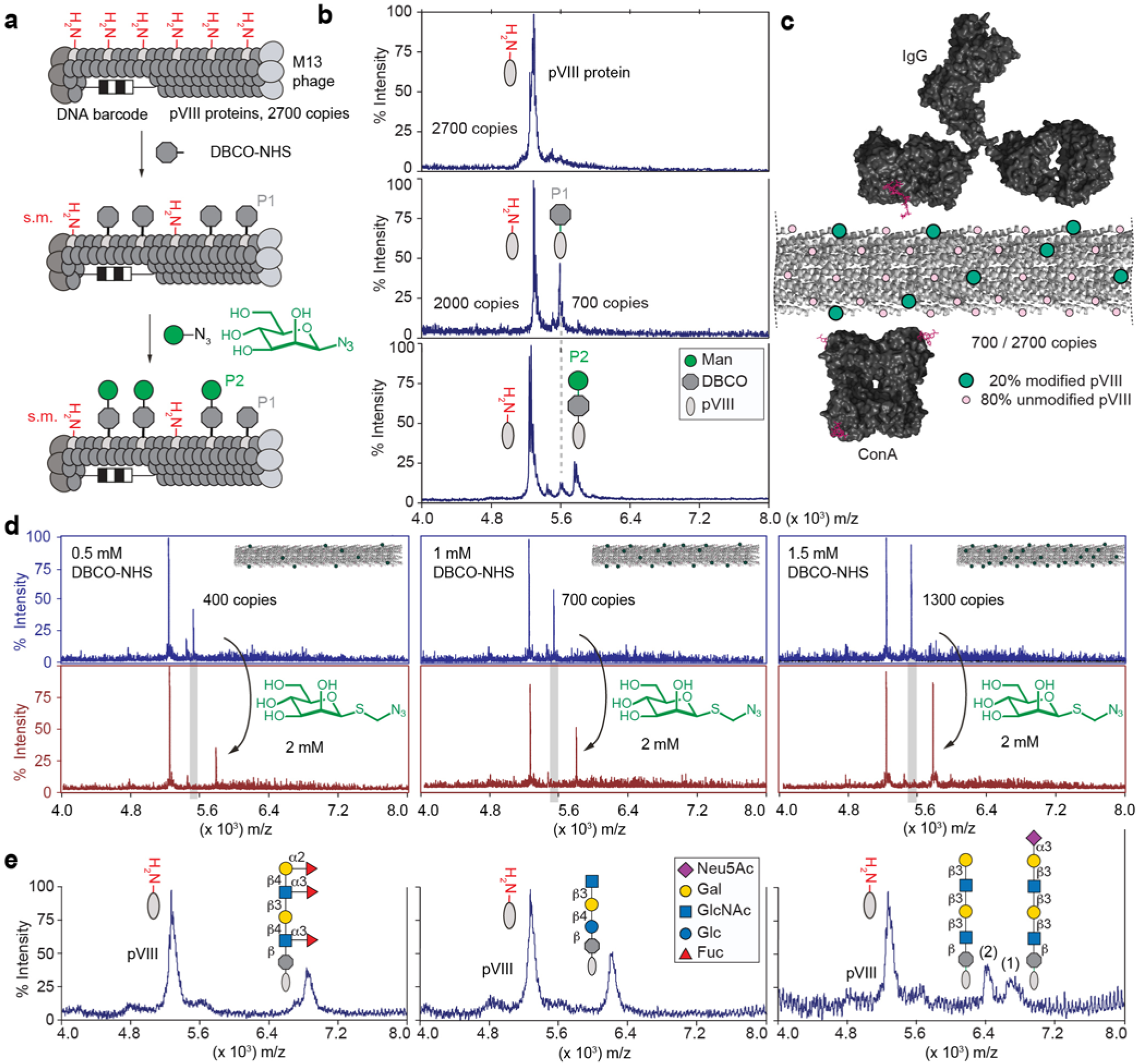
Synthesis and characterization of LiGA components. **a**, Representation of two-step chemical glycosylation of phage. **b**, MALDI mass spectrometry characterization of starting material (protein pVIII), alkyne-functionalized product (DBCO-pVIII, P1), and glycoconjugate product (P2). **c**, Spacing of glycans on M13 virion (PDB: 2MJZ and 2C0W) compared to spacing of binding sites of IgG antibody and lectin ConA. **d**, Control of the density of the modification by controlling the concentration of DBCO-NHS. **e**, Representative spectra of chemical modification of coat protein pVIII with glycans of different structural complexity. In MALDI conditions, sialoglycans exhibited a loss of sialic acid during ionization in acidic matrix; therefore MALDI of phage particles decorated with glycans containing terminal Neu5Ac contained two peaks (intact glycoconjugate and that with cleaved Neu5Ac).

### Functional validation of LiGAs

We produced ∼140 glycan-modified phage conjugates and characterized the density of glycans on phage by MALDI (summarized in Data/Maldi.pdf in the Supplementary Information). Mixing a subset of these differentially barcoded glycan-labeled phages produces a LiGA library that contains any desired combinations of glycans and glycans at different densities. Here, we employed several related LiGA libraries by mixing of 50–80 conjugates that display different glycan types (Supplementary Table S3). An ELISA confirmed the functional integrity of glycans on individual glycosylated phage (Supplementary Fig. S7); however, analysis by ELISA was incompatible with optimization of assays that contained mixtures of phage clones. We thus employed a LiGA composed of phage that transduce galactosidase, mNeonGreen^55^, or mCherry^56^ reporters within host cells (Fig. 2a-b). Composition of the mixture of these phage clones can be measured as plaque forming units (PFU). Plating of the input and the output of the assay on a bacterial agar overlay enabled measurement of blue, white, green-fluorescent and red-fluorescent plaques (Fig. 2c) and define recovery of each subpopulation of phage as PFU^output^/PFU^input^. This recovery tracked the binding of the associated glycans: such as binding the Manα-[green] to ConA but not Galβ1-4Glc1β-[red] and unmodified wild type [white] present in the same mixture (Fig. 2c). It further demonstrated that binding of α-Man-decorated clone to ConA either alone, or as part of the complex LiGA mixture, was statistically indistinguishable (Fig. 2c). This observation suggested that in an assay with super-stoichiometric GBP, the binding of the glycosylated particles is not influenced by the composition of the LiGA. The PFU assay validated that ConA or Galectin-3 (G3C) proteins immobilized on agarose beads can enrich clones decorated by Manα1-6(Manα1-3)Manα1-[red]-phage (Man_3_-[red]-phage) and Galβ1-3GlcNAcβ1-3Galβ1-4GlcNAcβ1-phage (LNT-NAc-[green]-phage) from a complex mixture of 75 glycosylated clones. Enrichment of LNT-NAc by G3C was specifically inhibited by soluble lactose (Fig. 2d). The PFU-assay is target agnostic and allows optimization of both protein and cell-binding assays. For example, it confirmed that Chinese hamster ovary (CHO) cells engineered to express murine CD22 (mCD22) preferentially bound phage with Neu5Gcα2-6LacNAc over those with Neu5Acα2-6LacNAc, reflecting the known specificity of mCD22 for sialosides with Neu5Gc. In contrast, CHO cells that expressed human CD22 (hCD22) enriched both glycans, consistent with its equal preference for α2-6 sialosides with NeuAc or NeuGc. As expected, the parental CHO cell line exhibited no enrichment of either sialoglycan (Fig. 2f).

**Fig. 2:**
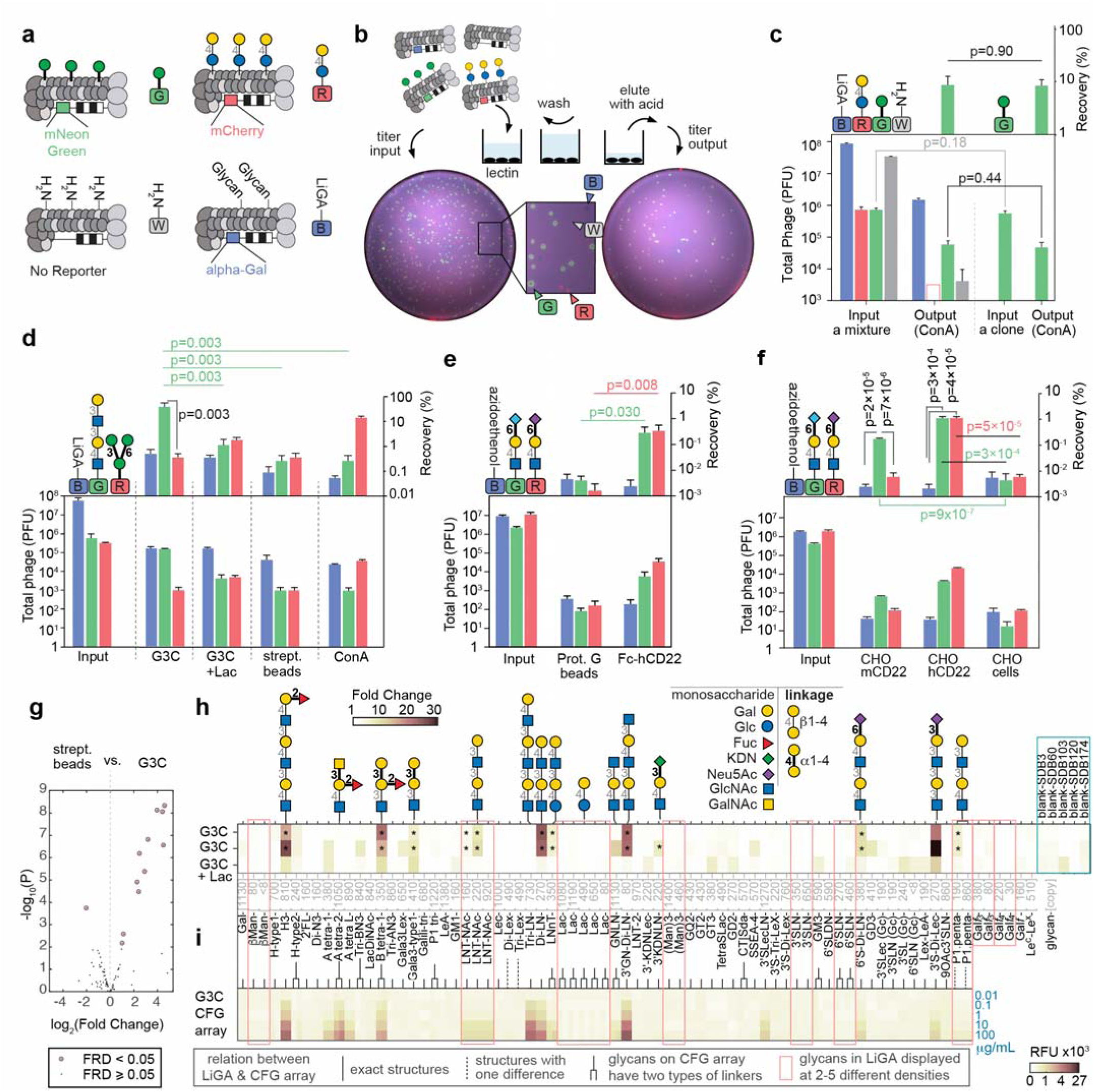
A colormetric PFU assay for rapid validation of LiGA. **a**, Clonal phage containing mNeonGreen reporter gene modified with mannose (α-Man), clonal phage containing mCherry gene with lactose (Lac), LiGA 3 × 3 array, constructed with LacZ reporter gene. **b**, Scheme of screenings against a immobilized GBPs and a fluorescent image of agarose plate used in plaque forming assay. **c**, Recovery of α-Man (mNeonGreen) in complex mixture and alone. n=3 input, n=4 output, Student’s t-test. **d**, The titer results suggested specific retention of G3C binders and ConA binders. n=4, Student’s t-test. **e**, hCD22 retains α2-6 sialoside, Neu5Gc (mNeonGreen) and Neu5Ac (mCherry) indiscriminately and has significantly higher recoveries compared to agarose beads. n=2 inputs, n=6 other, Student’s t-test. **f**, mCD22 expressing CHO cells have preference for recognizing α2-6 sialoside, Neu5Gc while hCD22 on cells do not have preference for Neu5Gc and Neu5Ac. n=5 CHO-hCD22, n=4 others, Student’s t-test. **g**, Bioconductor EdgeR differential enrichment (DE) analysis of deep-sequencing of LiGA binding to G3C and streptavidin beads (n=3) as described in (**d**). **h-i**, comparison of the (**h**) Fold Change (FC) values observed in DE analysis. n=4 for G3C+Lac, n=3 others with (**i**) data observed in glass-based glycan array (n=6); public microarray data, CFG request #2564; searchable as “primscreen_6003” to “primscreen_6011” at CFG website^52^. Data in (**c-f**) are presented as mean + s.d. In (**g**-**h**) the FC, FDR and p-values were calculated using negative binomial model, TMM-normalization and BH-correction for FDR (* designate FDR<0.05).

We used the PFU assay to optimize the input and wash stringency of LiGA binding assays, to check the quality of the biotin-labeled protein (Supplementary Fig. S8) or to detect whether a complex LiGA mixture contained target-binding clones (Supplementary Fig. S9). The PFU assay was a critical proxy for downstream PCR and sequencing. For example, robust PCR-based preparation of phage DNA for sequencing requires >10^3^ copies of phage DNA. The PFU assay identified the optimal conditions that minimized non-specific binding and yielded >10^4^ phage particles (Fig. 2d), which were then used as DNA templates for deep-sequencing and further analysis of glycan-protein interaction (Fig. 2g-h).

### LiGA characterizes the glycan recognition profile of glycan binding proteins (GBPs)

Next we analyzed the interactions of several GBPs with LiGAs (Fig. 2g,h). Here we used LiGA-75, which contains 75 glycoconjugates produced from 62 different glycans with nine of these glycans displayed at 2–5 different densities. Differential enrichment (DE) analysis was employed to examine phage DNA sequences associated with the control streptavidin agarose (SA) beads and phage DNA sequences associated with biotinylated GBP immobilized on SA-beads. Significantly enriched glycans were identified by Biocondutor EdgeR DE analysis^57, 58^ using negative binomial model Trimmed Mean of M-values (TMM) values normalization^59^, Benjamini–Hochberg (BH)^60^ correction to control the false discovery rate (FDR) at α□=□0.05. This analysis identified 11 glycans associating significantly with Galectin-3 (G3C)-SA-beads and not SA-beads alone (Fig. 2g). The enrichment pattern was consistent between independently conducted experiments and it was ablated in the presence of soluble lactose (Fig. 2h). The results for glycan recognition by G3C using LiGA-75 (Fig. 2h) is consistent with results obtained using a glass slide-based printed microarray deposited to the CFG database^52^ (Fig. 2i). To expand on these observations, we tested binding of a panel of biotinylated GBP and antibodies to LiGA-65 comprising 65 glycoconjugates produced from 56 different glycans with four of the glycans (βMan, Lac, 3’SLN and Gal*f*_4_) displayed at 2-4 different densities. Fig. 3a describes enrichment by FLAG monoclonal antibody (Flag) and anti-galactofuranose monoclonal antibody (Galf4) and nine lectins: Cholera Toxin B subunit (CTb), *Lens culinaris* agglutinin (LCA), *Pisum sativum* agglutinin (PSA), Soybean agglutinin (SBA), *Agrocybe cylindracea* lectin (ACG), *Narcissus pseudonarcissus* lectin (NPL), *Maackia amurensis* leucoagglutinin (MAL I), *Ulex europaeus* agglutinin (UEA), and *Sambucus nigra* lectin (SNA). Fig. 3b describes public CFG microarray data for eight of the aforementioned lectins.

**Fig. 3:**
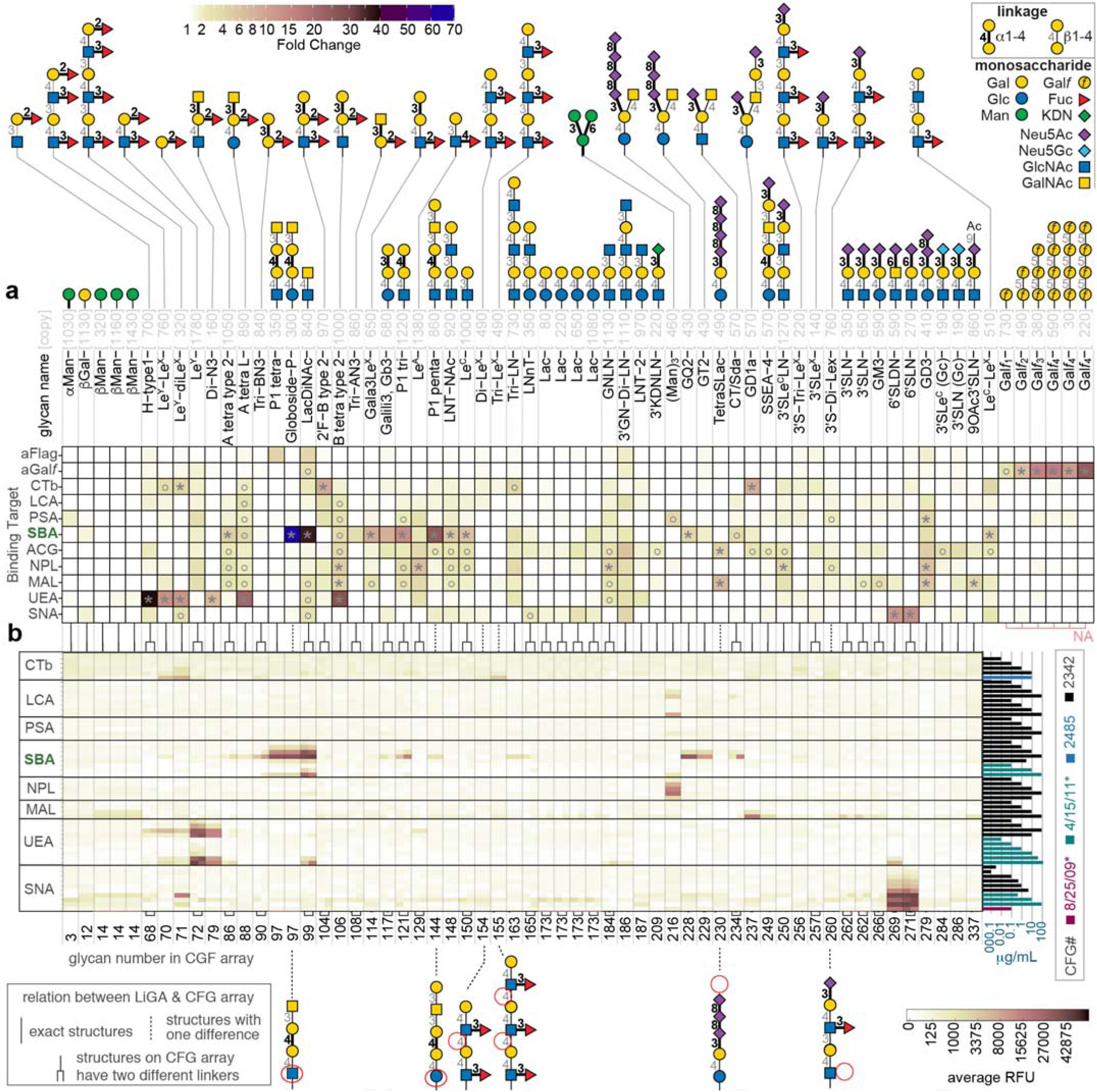
Summary of lectin binding to LiGA composed of 66 different glycophage constructs. **a**, Heat map describing the binding of 11 proteins to LiGA-65. Glycan notations, color-code of α/β linkages is shown in the legend, n=4. **b**, Binding specificity of the glycan binding proteins to glass-based glycan array. The data is an aligned subset of multiple publicly available CFG array data. CFG request numbers or experiment dates are provided for each dataset, n=6 (see details in Supplementary Fig. S13). The lines between (**a**) and (**b**) connect either identical or structural related glycans in LiGA and CFG. In (**a**) * FC>3.8, whereas circles represents 3.8>FC>2 with FDR≤0.05 in both cases. FC was calculated by Bioconductor EdgeR DE analysis using negative binomial model, TMM-normalization and BH-correction for FDR. Error bars represent s.d. propagated from the variance of the TMM-normalized sequencing data.

There was general agreement between glycan-GBP interactions measured by LiGAs and binding of lectins to printed glycan microarrays. Similarly to previous reports that evaluated the binding of lectins to glycans presented on different platforms^30, 61^, lectin binding to glycans in LiGAs and printed arrays highlighted some cross-platform differences. We noted several glycans that exhibited binding to GBP in LiGA experiments and in the published literature but not in printed arrays. For example (Neu5Acα2-8)_n_Neu5Acα2-3Galβ1-4Glcβ-Sp glycans (n=3: TetraSLac, n=1: GD3) do not bind to MAL-I in printed array (Fig. 3b); however, these glycans bind to this lectin as part of a LiGA (Fig. 3a) and when printed on glass as a multivalent BSA conjugate^61^. Binding of Galili-tri (Galα1-3Galβ1-4Glcβ-Sp) to SBA was not observed in printed array but this glycan bound to SBA as part of a LiGA and interaction of this glycan with SBA was confirmed by lectin frontier affinity chromatography^62^ (entry LfDB0166 at Lectin Frontier Database^63^). Interactions of cholera toxin B (CTB) with blood group B antigens on type 2 chains (2’F-B type 2) is not detected in a printed array but is detected in a LiGA. Such interactions have been confirmed by X-ray crystallography to occur at the non-primary binding site of CTB with low mM monovalent affinity^64^. We also note that some glycans observed to bind to lectins in printed arrays did not exhibit enrichment in pull down of a LiGA by lectins. Notable examples were: (i) phage decorated with Manα1-6[Manα1-3]Manα1-was not enriched by NPL and LCA; (ii) Phage decorated Le^Y^ was not recognized by UEA nor CTB; in the same LiGA mixture, both UEA and CTB enriched Le^Y^-(Le^X^)_1-2_ glycans but not the Le^X^-repeats alone; and (iii) Phage clones decorated by tri-LN and group A type-2 glycans were not recognized by G3C (Fig. 2h). Diminished recognition of glycan by lectins may result from suboptimal display of glycans on phage. Indeed, we noted that that when the same glycan was presented to G3C at several different densities on phage (e.g., LNT-NAc Fig. 2h), enrichment was the most optimal at one of those densities. Controlling the valency of the displayed glycans over several orders of magnitude is a critical technological benefit of LiGAs when compared to a monovalent DNA-glycan platform or low-valency phage-displayed glycan platforms. Building on this advantage, we further tested the effect of density of display of glycans on phage virion on recognition by some lectins.

To more systematically investigate the importance of density, a LiGA-9×6 was produced that contains nine glycans displayed at six different densities (Fig. 4a). Binding of LiGA-9×6 components to tetravalent ConA, Galectin-3 and mAb-Galf4 target uncovered bimodal density-dependent response in addition to the expected affinity-dependent response (Fig. 4b-d). Within the same glycan type, we observed a bimodal response: phage decorated with high copy of Manα1-6(Manα1-3)Manα1- (>300 per phage) exhibited weaker binding when compared to particles that contained intermediate (130–300) copies (Fig. 4b). We observed similar bimodal binding at different densities of Gal*f*-glycans to anti-Gal*f* IgG and the binding of LNT NAc to biotinylated Galectin-3 displayed on streptavidin beads (Fig. 4c-d). The LiGA technology decoupled affinity-based and avidity-based observations, highlighting both optimal recognition units and their density. Similar observations have been made in prior studies that employed arrays of glycan-BSA conjugates printed on a glass slide^24^. Critical to both approaches is the quality control of every synthesized glycoconjugate or LiGA components by MALDI to create arrays with predefined, consistent valency of glycans.

**Fig. 4:**
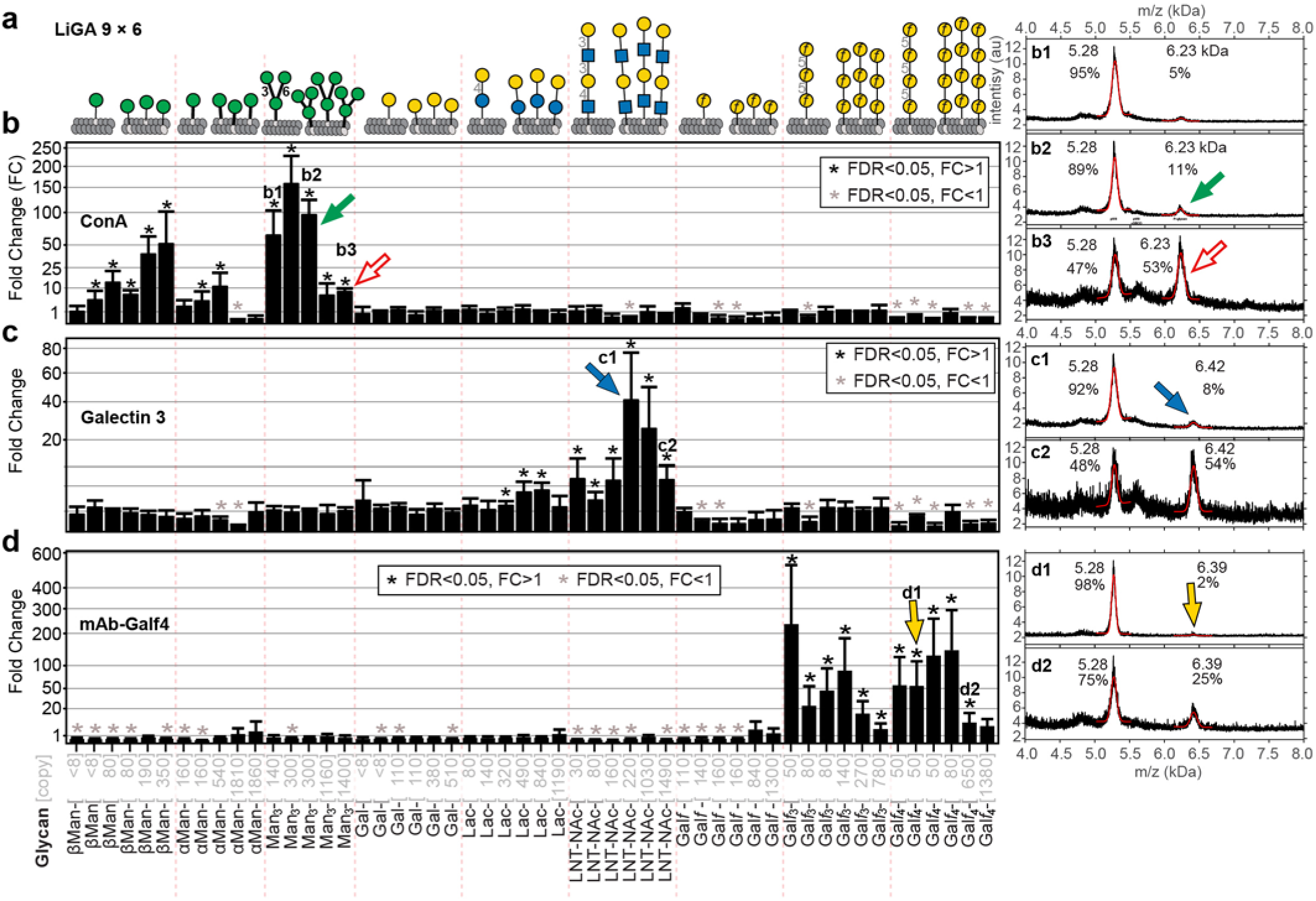
LiGA measures affinity and avidity-based responses in purified GBPs. **a**, LiGA 9×6 is composed of nine glycans displayed at up to six densities. **b**, Binding of LiGA 9×6 to ConA. n=3 test, n=4 control. **c**, Galectin-3, n=4. **d**, mAb-Galf_4_, n=3. Measurements by deep-sequencing confirms enrichment of specific glycans; for each glycan-GBP pair, the phage that display high density of the glycan showed significantly lower enrichment than those that display intermediate density. MALDI spectra of the phage glycoconjugates of low and high density. LiGA components **b1, b2, b3, c1, c2, d1** and **d2** described in (**b**-**d**). FC was calculated by Bioconductor EdgeR DE analysis using negative binomial model, TMM-normalization and BH-correction for FDR. Error bars represent s.d. propagated from the variance of the TMM-normalized sequencing data.

### LiGAs characterize the glycan recognition profile of cell-surface lectins

The previously mentioned PFU assay highlighted the possibility of interrogating specific recognition of glycans on the cell surface. Therefore, we probed two important immunomodulatory receptors, hCD22 and DC-SIGN, expressed on the cell surface with LiGA-71 produced from 66 different glycans with five of these glycans displayed at two different densities. To initially dissect hCD22 recognition, we immobilized hCD22-Fc on protein G-resin and tested with LiGA-71. Deep-sequencing confirmed known recognition of α2-6-linked sialic acid glycans but not the α2-3-linked regioisomers or asialo-glycans (Fig. 5a). We incubated the same LiGA-71 with CHO cells expressing hCD22 or control CHO cells, and sequenced the associated phage (Fig. 5b). Differential enrichment analysis uncovered indistinguishable recognition profile within LiGA-71 for hCD22 on the surface of CHO cells compared to recombinant soluble hCD22 immobilized on agarose beads.

**Fig. 5:**
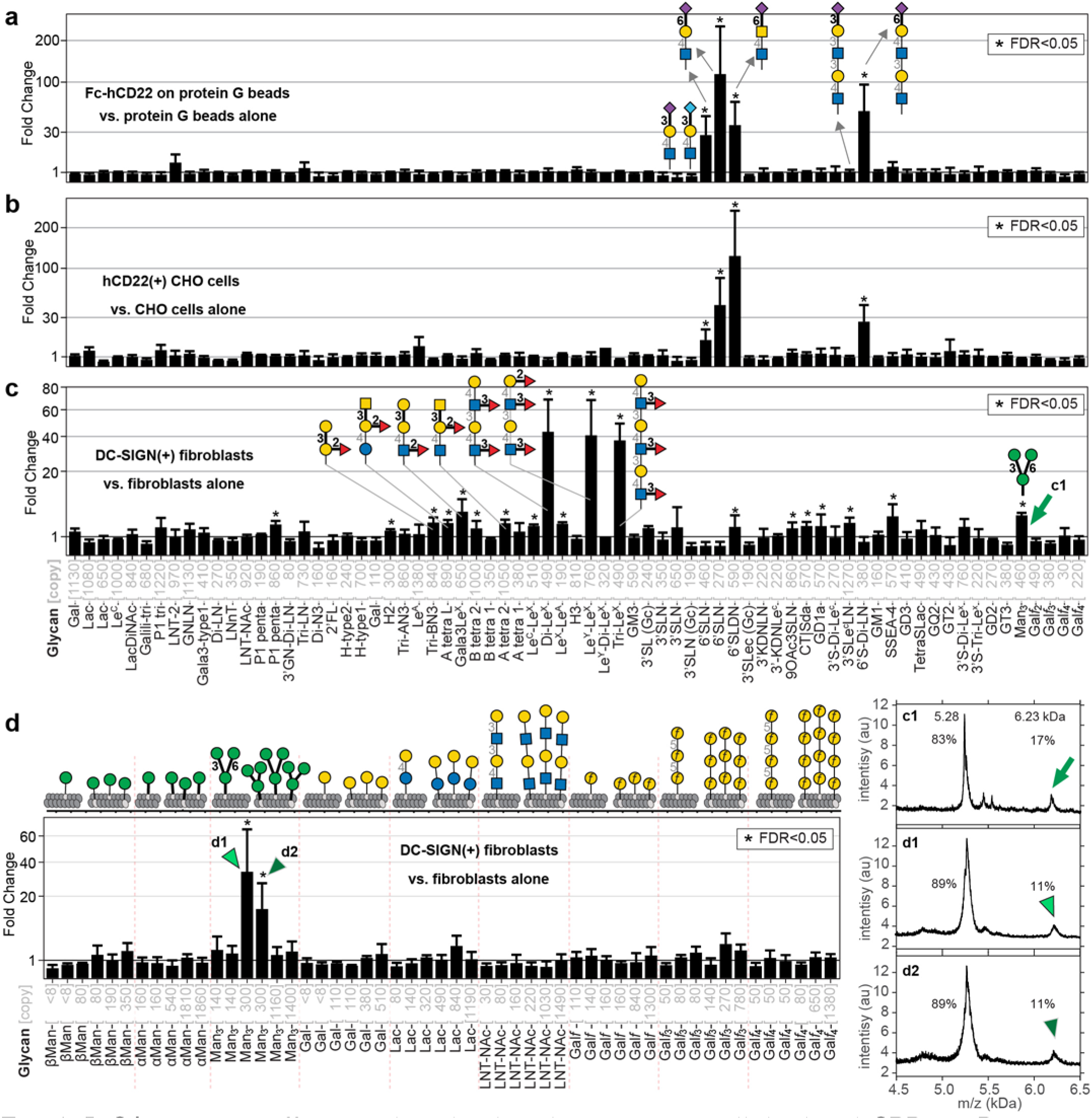
LiGAs measure affinity and avidity-based responses in cell-displayed GBPs. **a**, Binding of LiGA-71 to construct hCD22-Fc normalized by binding of the same LiGA to protein-G beads, n=3. **b**, Binding of LiGA-71 to hCD22(+) CHO cells normalized by binding of the same LiGA to CHO cells, n=4. **c**, Binding of LiGA-71 to DC-SIGN(+) fibroblasts normalized by binding of the same LiGA to fibroblasts, n=4. **d**, Binding of LiGA 9×6 to DC-SIGN(+) fibroblasts performed analogously to (**c**), n=3. Phage virions that display 300 copies (**d**) or 460 copies (**c**) of core trimannoside exhibit significant binding to DC-SIGN clusters on the cell surface, virions with <150 copies or >1200 copies exhibit insignificant enrichment. Calculations of FC, FDR and error bars are identical to those described in Fig. 2h, Fig. 3a, and 4b-d.

Next, we incubated LiGA-71 with rat fibroblasts expressing DC-SIGN or isogenic fibroblasts without DC-SIGN and compared the barcoded DNA associated with the DC-SIGN(+) and DC-SIGN(–) cell pellets. The resulting binding profile from DC-SIGN (+) cells revealed enrichment of αMan3 (Manα1-6(Manα1-3)Manα1-), glycans with Le^X^ motifs and other fucosylated (α1-3 or α1-2-linked) glycans (Fig. 5c). We observed a statistically significant but modest fold change (FC) enrichment of blood-group glycans, which was considerably lower than FC in interaction of these glycans with blood-group antibodies (Supplementary Fig. S10). Binding of LiGAs to DC-SIGN on cells was qualitatively similar to the glycan-binding profile of purified DC-SIGN measured by canonical printed glycan arrays deposited to CFG website^52^ (Supplementary Fig. S11), with only minor differences. For example, αMan3 (Manα1-6(Manα1-3)Manα1-(CH_2_)_6_-phage) was reproducibly enriched in LiGA-71 as well as in LiGA mixtures of other composition (Supplementary Fig. S11a), whereas the CFG glycan microarray only detected binding only to penta- and decamannose glycans. Recognition of alpha-linked Man3-motifs by DC-SIGN was postulated based on structural studies^65^ and specific binding of Manα1-6[Manα1-3]Manα1-(CH_2_)_6_-N_3_ to DC-SIGN was measured^66^. Thus, lack of detection of the Manα1-6(Manα1-3)Manα1-(CH_2_)_9_-amino glycan on the CFG glycan microarray may be the result of differences in the presentation of the glycan in the LiGA and printed array formats (Supplementary Fig. S11b-c). Deploying the density-scanning LiGA-9×6 array with DC-SIGN(+)/(-) fibroblasts not only confirmed the specific recognition of αMan3 but also uncovered pronounced bell-shaped dependence on density of αMan3 (Fig. 5d, Supplementary Fig. S12). Phage particles that displayed 300 copies exhibited binding significantly higher than those with <150 or >1000 copies of αMan3. This density scan further suggests that presentation of the αMan3 on the slide-based arrays may not have been optimal for the recognition by DC-SIGN. These observations highlight the ability of the LiGA platform to robustly and reproducibly measure both affinity-based and avidity-based effects on glycan-GBPs interactions and in a complex biological milieu on the surface of mammalian cells.

LiGAs, like other M13-display platforms, makes it possible to detect interactions between displayed ligands and receptors on the surface of organs and cells in live animals^47^. To demonstrate this capacity, we tracked the biodistribution of a LiGA to different organs in mice and its association with two subpopulations of B cells. We carried out a mixed bone marrow chimera to generate mice that contain a 50:50 mixture of B cells expressing transgenic hCD22^67^ or lacking hCD22 expression, both of which are on a mCD22^-/-^ background (Fig. 6a and SI for details). We used a red-green-blue-white (RGBW) LiGA, which was a 1:1 mixture of LiGA-70 linked to Lac-reporter phage (‘blue’) and non-glycosylated phage associate with reporter-free phage (‘white’) supplemented with ∼1% of two phage clones displaying Neu5Gcα2-6LacNAc and Neu5Acα2-6LacNAc linked to NeonGreen and mCherry transducing phage. The latter ‘red’ and ‘green’ clones previously demonstrated significant enrichment by cell surface-displayed hCD22 protein (Fig. 2f). We injected this RGBW-LiGA in the tail vein of mice (n=3), recovered the organs after one hour, sorted the B cell populations from the spleen and measured the composition of the library associated with each organ or B cell type by PFU assay (Fig. 6b). Gratifyingly, we observed significant enrichment of the phage displaying the NeuAcα2-6Gal sialosides only with B cells that expressed hCD22 sorted from the same spleen of the same mouse (Fig. 6c). No enrichment of the phage displaying the NeuAcα2-6Gal sialosides was found in the bulk organs, including the lungs, heart, liver, kidney. This experiment demonstrates the feasibility of studying glycan-GBP interactions on the surface of cells *in vivo* using the LiGA technology.

**Fig. 6:**
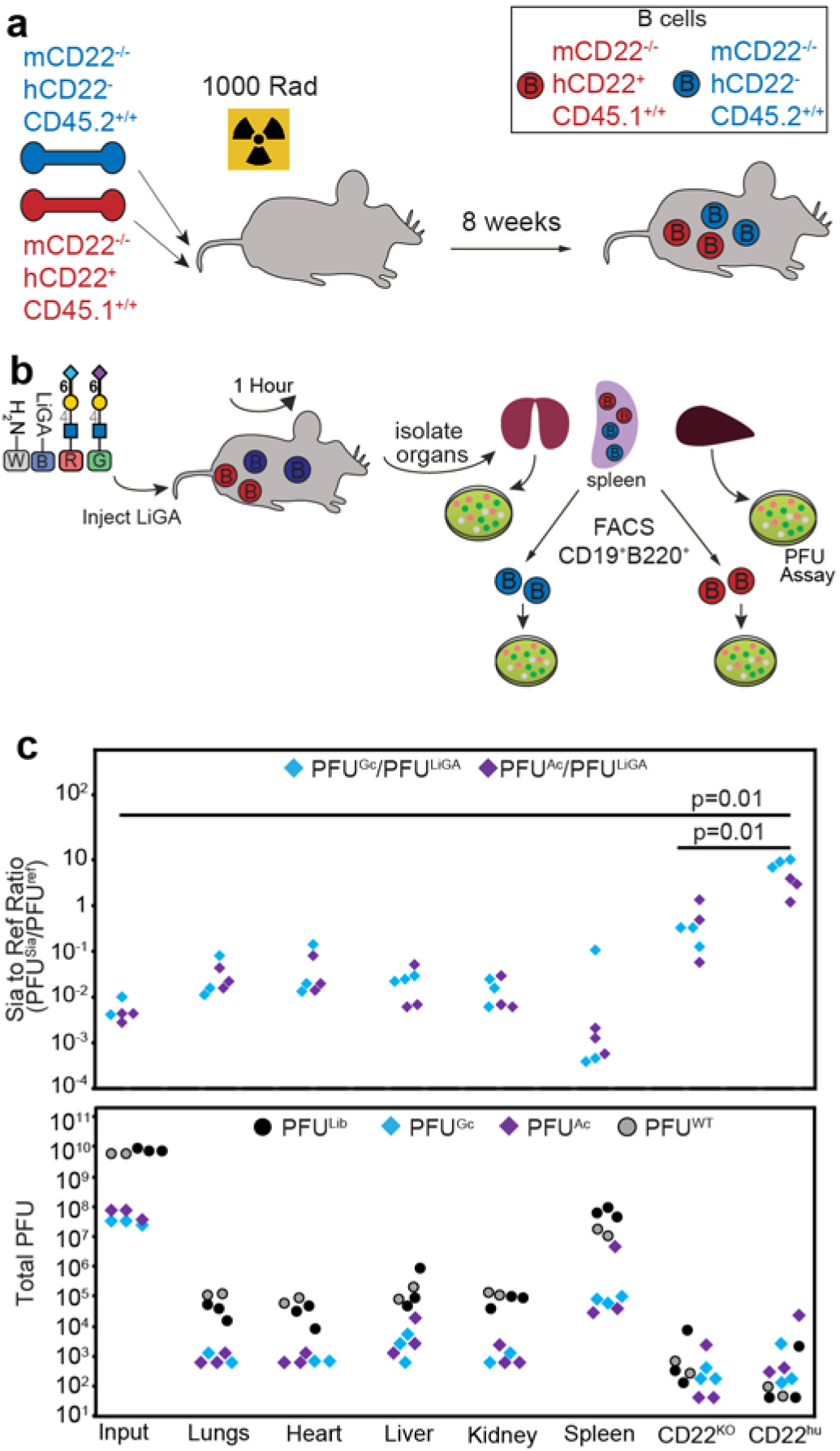
**a**, Lethally irradiated mice were reconstituted with a 50:50 mixture of bone marrow from the two indicated recipient bone marrow to generate chimeric mice that have a mixture of hCD22^+^ and hCD22^-^ B cells both lacking mCD22 and with different CD45 congenic markers for sorting purposes. **b**, The LiGA was injected into mice via the tail vein. One hour later, the mice were euthanized, and organs collected. The B cells were stained with the appropriate antibodies and sorted for differential expression of hCD22. Sorted cells and cells from organs were analyzed for phage by PFU assay. **c**, Analysis of types of phage from the PFU assay expressed as PFU enrichment (*top*) or total PFU (*bottom*). Tissues from n=3 mice, p-values calculated by Student’s t-test.

## Discussion

Multivalent interactions are ubiquitous in biological systems^68^. At a cell surface interface, multivalent receptor–ligand interactions are central to signal transduction, cell–cell recognition and the interpretation of external signals by cells^11^. The LiGA technology makes interrogation of these interactions possible using a mixture of monodisperse multivalent probes^19^: each probe with a DNA barcode and well-defined glycan density. Modern advances in DNA barcoding, deep-sequencing and bioorthogonal modification enabled high-throughput manufacturing of LiGA components via a parallel conjugation of the glycans to phage virion. Like the majority of arrays with glycans printed on glass, the linkage between the glycan and the carrier (phage) is not natural. However, chemical conjugation affording 30–1500 copies of glycan per phage particle was key to addressing two critical limitations of prior glycophage: density and glycan diversity. The pioneering reports of glycophage display by groups of DeLisa^44^ and Aebi^45^ employed M13 “3+3” phagemid particles displaying a D/E-X_1_-N-X_2_-S/T acceptor motif on pIII protein. Like all 3+3 display systems, it is a monovalent display of one copy of engineered pIII fusion and four copies of wild type pIII (on average). Importantly, both groups observed that only 1 in 100 phagemid particles carried a glycan biosynthesized by the bacterial host. The display of truncated glycopeptides was common. The latter problem was subsequently resolved by DeLisa and co-workers^46^ by employing truncated pIII instead of full-length pIII. However, even in the optimized construct, the authors could not quantify the density of glycans on the different glycophages due to the low level of phage glycosylation^46^. The biosynthesis of a DNA-encoded *bona fide* multivalent display of diverse glycan structures on phage remains an interesting bioengineering challenge. Recent advances in this area include biosynthesis of multivalent display of glycoproteins on mammalian cell surface^69^ and multivalent display of biosynthesized glycans on DNA-free virus-like particles^70^. Unlike these and other display systems that employ the biosynthesis of glycans, LiGA decouples DNA encoding and glycan display from biosynthesis. Chemical manufacturing of LiGA components allows the repurposing of many chemical approaches previously employed in the construction of traditional slide-based glycan arrays^20^.

Although SPAAC is a convenient linking strategy, other linking strategies may also work. For example, ligation of glycans with a carboxy linker to lysine side chains of bovine serum albumin is routinely employed in microarray manufacturing^24^ (reviewed in^20^). This approach is straightforward to adapt to LiGA manufacturing via acylation of N-termini of the pVIII protein instead of BSA. A growing palette of site-specific chemical conjugation strategies can inspire new and improved variants of LiGAs. The source of glycans in this paper is chemical, but it should also be possible to develop LiGAs using glycans isolated from natural sources. For example, ongoing efforts in our laboratory are focused on exploring the chemoenzymatic synthesis of LiGAs from biantennary N-linked glycan precursors.

We have shown that the multivalent LiGA platform effectively measured the glycan-binding preferences of purified GBPs and GBPs displayed on cells, including cells in organs in live animals. A typical preparation of one of the individual LiGA components requires only 50–100 μg of glycan and yields 10^12^ PFU of glycosylated phage particles. This amount is sufficient to perform on the order of 100,000 lectin-binding or cell-binding assays using 10^6^–10^7^ copies of each glycosylated clone as input. The deployment of the LiGA technology further benefits from accurate knowledge of copy number, composition, and valency of LiGA components. This critical information will facilitate the investigation and modeling of complex interactions between a multivalent display of glycans and multivalent display of receptors on the cell surface. The LiGA platform has the potential to uncover critical glycan binding information of between glycans and cell-displayed GBP that has remained elusive due to limitation of other glycan array platforms.

## Materials

Detailed **Biochemical Methods 1.1-1.17, Data Processing Methods 2.1-2.6**, and **Synthetic Methods 3.1-3.6** are available as part of the Supplementary Information document.

## Supplementary information

Supplemental Figures S1-S13, Supplemental Schemes S1-S4, Supplemental Tables S1-S3; detailed biochemical methods describing the synthesis of LiGA components, protein- and cell-binding assays, animal experiments; synthetic methods describing the synthesis of the glycans, and data processing methods describing the analysis of the DNA sequencing data, statistical methods.

### Source data

submitted as “data.zip” contain files describing (i) MALDI spectra of the LIGA components, (ii) *.xlsx tables describing the correspondences between DNA and glycan structures (“LiGA dictionaries”); (iii) *.xlsx tables describing raw sequencing data and statistical analysis used to generate Figures 2h, 3a, 4b-d, 5a-d, S9c, S10, S11a, S12; (iv) CFG glass array data used to generate Figures 3b, S11b, S13.

## Data Availability

All raw deep-sequencing data is publically available on http://ligacloud.ca/ with data-specific URL listed in Table S3. DNA sequences of the three LiGA phage constructs have been deposited to GeneBank (#MN865131, MN865132, MN872303). MatLab, Python and R scripts have been deposited to https://github.com/derdalab/liga

## Contributions

N.J.B. constructed and characterized all LiGA plasmids and libraries, M.S., S.Sa., J.M., D.F.V., R.R., performed isolation of phage clones, modifications of phage by glycans, MALDI analysis, protein-binding experiments, and *in vitro* assays. M.S., E.R., J.M., and R.D. performed cell-based assays, M.S.M and S.Sa. performed animal experiments, E.J.C., M.S. and R.D. performed statistical analysis., S.Se., performed custom analysis of public CFG data., T.L.L., P.Z., C.C.L., C.N. and J.C.P. contributed synthetic glycan reagents, A.K. and M.B. constructed synthetic antigens to raise and characterize anti-galactofuranose antibodies, which were produced by S.Sa. X.X. and R.B.Z. constructed antigens for incorporation into the LiGA. R.D., and M.S. wrote the manuscript, R.D., M.S.M., T.L.L., and J.C.P. edited the final manuscript and contributed intellectual and strategic input. All authors approved the final manuscript.

## Acknowledgments

We thank the staff at the University of Alberta mass spectrometry facility (Chemistry Department) for help with MALDI analysis and Sophie Dang at the molecular biology service unit for assistance with Illumina sequencing. Cell sorting was performed at the University of Alberta, Faculty of Medicine and Dentistry Flow Cytometry Facility with financial support from the Faculty of Medicine and Dentistry and Canada Foundation for Innovation (CFI) awards to contributing investigators. We thank Kurt Drickamer (Imperial College, London), Bruce Turnbull (University of Leeds), David Bundle, Christopher Cairo and Lori West (University of Alberta) for provision of critical reagents. The authors acknowledge funding from NSERC (RGPIN-2018-04365 to T.L.L., RGPIN-2018-03815 to M.S.M., and RGPIN-2016-402511 to R.D.) and NSERC Accelerator Supplement (to R.D.), GlycoNet (SD-1 to T.L.L., TP-22 to R.D.), Alberta Innovates Strategic Research Project to R.D., and NIH projects (AI118842 to M.S.M., and GM062116 and AI050143 to J.C.P.). Many compounds were prepared by the Consortium for Functional Glycomics supported by NIH GM061126. Infrastructure support was provided by CFI New Leader Opportunity (to R.D. and M.M.). J.M. acknowledges summer research fellowship from GlycoNet and Alberta Innovates Health Solutions.

## References

1. Shendure, J. & Ji, H. Next-generation DNA sequencing. Nat. Biotechnol. 26, 1135 (2008).

2. Lander, E.S. Array of hope. Nat. Genet. 21, 3–4 (1999).

3. Blixt, O. et al. Printed covalent glycan array for ligand profiling of diverse glycan binding proteins. Proc. Natl. Acad. Sci. U. S. A. 101, 17033–17038 (2004).

4. van Kooyk, Y. & Rabinovich, G.A. Protein-glycan interactions in the control of innate and adaptive immune responses. Nat. Immunol. 9, 593–601 (2008).

5. Varki, A. Glycan-based interactions involving vertebrate sialic-acid-recognizing proteins. Nature 446, 1023 (2007).

6. Stevens, J. et al. Structure and Receptor Specificity of the Hemagglutinin from an H5N1 Influenza Virus. Science 312, 404–410 (2006).

7. Stevens, J., Blixt, O., Paulson, J.C. & Wilson, I.A. Glycan microarray technologies: tools to survey host specificity of influenza viruses. Nat. Rev. Microbiol. 4, 857–864 (2006).

8. Raman, R., Raguram, S., Venkataraman, G., Paulson, J.C. & Sasisekharan, R. Glycomics: an integrated systems approach to structure-function relationships of glycans. Nat. Methods 2, 817 (2005).

9. Geissner, A. & Seeberger, P.H. Glycan Arrays: From Basic Biochemical Research to Bioanalytical and Biomedical Applications. Annu. Rev. Anal. Chem. (Palo Alto Calif.) 9, 223–247 (2016).

10. Smith, D.F., Cummings, R.D. & Song, X. History and future of shotgun glycomics. Biochem. Soc. Trans. 47, 1–11 (2019).

11. Bertozzi, C.R. & Kiessling, L.L. Chemical glycobiology. Science 291, 2357–2364 (2001).

12. Zhang, Y., Li, Q., Rodriguez, L.G. & Gildersleeve, J.C. An array-based method to identify multivalent inhibitors. J. Am. Chem. Soc. 132, 9653–9662 (2010).

13. Wang, C.C. et al. Glycan microarray of Globo H and related structures for quantitative analysis of breast cancer. Proc. Natl. Acad. Sci. U. S. A. 105, 11661–11666 (2008).

14. Xia, L., Schrump, D.S. & Gildersleeve, J.C. Whole-Cell Cancer Vaccines Induce Large Antibody Responses to Carbohydrates and Glycoproteins. Cell Chem. Biol. 23, 1515–1525 (2016).

15. Jacob, F. et al. Serum antiglycan antibody detection of nonmucinous ovarian cancers by using a printed glycan array. Int. J. Cancer 130, 138–146 (2012).

16. Fukui, S., Feizi, T., Galustian, C., Lawson, A.M. & Chai, W. Oligosaccharide microarrays for high-throughput detection and specificity assignments of carbohydrate-protein interactions. Nat. Biotechnol. 20, 1011–1017 (2002).

17. Demetriou, M., Granovsky, M., Quaggin, S. & Dennis, J.W. Negative regulation of T-cell activation and autoimmunity by Mgat5 N-glycosylation. Nature 409, 733–739 (2001).

18. Cecioni, S., Imberty, A. & Vidal, S. Glycomimetics versus multivalent glycoconjugates for the design of high affinity lectin ligands. Chem. Rev. 115, 525–561 (2015).

19. Kiessling, L.L., Gestwicki, J.E. & Strong, L.E. Synthetic multivalent ligands as probes of signal transduction. Angew. Chem. Int. Ed. 45, 2348–2368 (2006).

20. Park, S., Gildersleeve, J.C., Blixt, O. & Shin, I. Carbohydrate microarrays. Chem. Soc. Rev. 42, 4310–4326 (2013).

21. Rillahan, C.D. & Paulson, J.C. Glycan Microarrays for Decoding the Glycome. Annu. Rev. Biochem. 80, 797–823 (2011).

22. Godula, K., Rabuka, D., Nam, K.T. & Bertozzi, C.R. Synthesis and microcontact printing of dual end-functionalized mucin-like glycopolymers for microarray applications. Angew. Chem. Int. Ed. Engl. 48, 4973–4976 (2009).

23. Godula, K. & Bertozzi, C.R. Density variant glycan microarray for evaluating cross-linking of mucin-like glycoconjugates by lectins. J. Am. Chem. Soc. 134, 15732–15742 (2012).

24. Oyelaran, O., Li, Q., Farnsworth, D. & Gildersleeve, J.C. Microarrays with Varying Carbohydrate Density Reveal Distinct Subpopulations of Serum Antibodies. J. Proteome Res. 8, 3529–3538 (2009).

25. Dam, T.K. & Brewer, C.F. Lectins as pattern recognition molecules: The effects of epitope density in innate immunity. Glycobiology 20, 270–279 (2010).

26. Kwon, S.J. et al. Signal amplification by glyco-qPCR for ultrasensitive detection of carbohydrates: applications in glycobiology. Angew. Chem. Int. Ed. Engl. 51, 11800–11804 (2012).

27. Tom, J.K., Mancini, R.J. & Esser-Kahn, A.P. Covalent modification of cell surfaces with TLR agonists improves & directs immune stimulation. Chem. Commun. (Camb.) 49, 9618–9620 (2013).

28. Zhang, J. et al. Specific recognition of lectins by oligonucleotide glycoconjugates and sorting on a DNA microarray. Chem. Commun. (Camb.), 6795–6797 (2009).

29. Ciobanu, M. et al. Selection of a synthetic glycan oligomer from a library of DNA-templated fragments against DC-SIGN and inhibition of HIV gp120 binding to dendritic cells. Chem. Commun. (Camb.) 47, 9321–9323 (2011).

30. Yan, M.M. et al. Next-Generation Glycan Microarray Enabled by DNA-Coded Glycan Library and Next-Generation Sequencing Technology. Anal. Chem. 91, 9221–9228 (2019).

31. Chevolot, Y. et al. DNA-based carbohydrate biochips: A platform for surface glyco-engineering. Angew. Chem. Int. Ed. 46, 2398–2402 (2007).

32. Thomas, B. et al. Application of Biocatalysis to on-DNA Carbohydrate Library Synthesis. ChemBioChem 18, 858–863 (2017).

33. Novoa, A., Machida, T., Barluenga, S., Imberty, A. & Winssinger, N. PNA-encoded synthesis (PES) of a 10 000-member hetero-glycoconjugate library and microarray analysis of diverse lectins. ChemBioChem 15, 2058–2065 (2014).

34. Horiya, S., Bailey, J.K., Temme, J.S., Schippe, Y.V.G. & Krauss, I.J. Directed Evolution of Multivalent Glycopeptides Tightly Recognized by HIV Antibody 2G12. J. Am. Chem. Soc. 136, 5407–5415 (2014).

35. Kondengaden, S.M. et al. DNA Encoded Glycan Libraries as a next-generation tool for the study of glycan-protein interactions. bioRxiv, doi: https://doi.org/10.1101/2020.1103.1130.017012 (2020).

36. Macauley, M.S. et al. Antigenic liposomes displaying CD22 ligands induce antigen-specific B cell apoptosis. J. Clin. Invest. 123, 3074–3083 (2013).

37. Chen, W.C. et al. In vivo targeting of B-cell lymphoma with glycan ligands of CD22. Blood 115, 4778–4786 (2010).

38. Kaltgrad, E. et al. On-virus construction of polyvalent glycan ligands for cell-surface receptors. J. Am. Chem. Soc. 130, 4578–4579 (2008).

39. Pochechueva, T. et al. Comparison of printed glycan array, suspension array and ELISA in the detection of human anti-glycan antibodies. Glycoconj. J. 28, 507–517 (2011).

40. Pochechueva, T. et al. Multiplex suspension array for human anti-carbohydrate antibody profiling. Analyst 136, 560–569 (2011).

41. Purohit, S. et al. Multiplex glycan bead array for high throughput and high content analyses of glycan binding proteins. Nat. Commun. 9, 258 (2018).

42. Liang, R. et al. Parallel synthesis and screening of a solid phase carbohydrate library. Science 274, 1520–1522 (1996).

43. Tjhung, K.F. et al. Silent Encoding of Chemical Post-Translational Modifications in Phage-Displayed Libraries. J. Am. Chem. Soc. 138, 32–35 (2016).

44. Celik, E., Fisher, A.C., Guarino, C., Mansell, T.J. & DeLisa, M.P. A filamentous phage display system for N-linked glycoproteins. Protein Sci. 19, 2006–2013 (2010).

45. Durr, C., Nothaft, H., Lizak, C., Glockshuber, R. & Aebi, M. The Escherichia coli glycophage display system. Glycobiology 20, 1366–1372 (2010).

46. Celik, E. et al. Glycoarrays with engineered phages displaying structurally diverse oligosaccharides enable high-throughput detection of glycan-protein interactions. Biotechnol. J. 10, 199–209 (2015).

47. Pasqualini, R. & Ruoslahti, E. Organ targeting In vivo using phage display peptide libraries. Nature 380, 364–366 (1996).

48. Kolonin, M.G., Saha, P.K., Chan, L., Pasqualini, R. & Arap, W. Reversal of obesity by targeted ablation of adipose tissue. Nat. Med. 10, 625–632 (2004).

49. Krag, D.N. et al. Selection of Tumor-binding Ligands in Cancer Patients with Phage Display Libraries. Cancer Res. 66, 7724–7733 (2006).

50. Arap, W. et al. Steps toward mapping the human vasculature by phage display. Nat. Med. 8, 121–127 (2002).

51. Scott, J.K. & Smith, G.P. Searching for peptide ligands with an epitope library. Science 249, 386–390 (1990).

52. http://www.functionalglycomics.org.

53. Jewett, J.C. & Bertozzi, C.R. Cu-free click cycloaddition reactions in chemical biology. Chem. Soc. Rev. 39, 1272–1279 (2010).

54. Crimmins, D.L., Mische, S.M. & Denslow, N.D. Chemical Cleavage of Proteins in Solution. Current Protocols in Protein Science 41, 11.14.11-11.14.11 (2005).

55. Shaner, N.C. et al. A bright monomeric green fluorescent protein derived from Branchiostoma lanceolatum. Nat. Methods 10, 407–409 (2013).

56. Shaner, N.C. et al. Improved monomeric red, orange and yellow fluorescent proteins derived from Discosoma sp. red fluorescent protein. Nat. Biotechnol. 22, 1567–1572 (2004).

57. Robinson, M.D. & Smyth, G.K. Moderated statistical tests for assessing differences in tag abundance. Bioinformatics 23, 2881–2887 (2007).

58. Robinson, M.D. & Smyth, G.K. Small-sample estimation of negative binomial dispersion, with applications to SAGE data. Biostatistics 9, 321–332 (2008).

59. Robinson, M.D. & Oshlack, A. A scaling normalization method for differential expression analysis of RNA-seq data. Genome Biol. 11, R25 (2010).

60. Benjamini, Y. & Hochberg, Y. Controlling the False Discovery Rate - a Practical and Powerful Approach to Multiple Testing. J. R. Statist. Soc. B 57, 289–300 (1995).

61. Wang, L.L. et al. Cross-platform comparison of glycan microarray formats. Glycobiology 24, 507–517 (2014).

62. Tateno, H., Nakamura-Tsuruta, S. & Hirabayashi, J. Frontal affinity chromatography: sugar-protein interactions. Nat. Protoc. 2, 2529–2537 (2007).

63. Hirabayashi, J., Tateno, H., Shikanai, T., Aoki-Kinoshita, K.F. & Narimatsu, H. The Lectin Frontier Database (LfDB), and Data Generation Based on Frontal Affinity Chromatography. Molecules 20, 951–973 (2015).

64. Heggelund, J.E. et al. High-Resolution Crystal Structures Elucidate the Molecular Basis of Cholera Blood Group Dependence. PLoS Pathog. 12, e1005567 (2016).

65. Feinberg, H., Mitchell, D.A., Drickamer, K. & Weis, W.I. Structural basis for selective recognition of oligosaccharides by DC-SIGN and DC-SIGNR. Science 294, 2163–2166 (2001).

66. Ng, S. et al. Genetically-encoded fragment-based discovery of glycopeptide ligands for DC-SIGN. Biorg. Med. Chem. 26, 5368–5377 (2018).

67. Bednar, K.J. et al. Human CD22 Inhibits Murine B Cell Receptor Activation in a Human CD22 Transgenic Mouse Model. J. Immunol. 199, 3116–3128 (2017).

68. Mammen, M., Choi, S.K. & Whitesides, G.M. Polyvalent interactions in biological systems: Implications for design and use of multivalent ligands and inhibitors. Angew. Chem. Int. Ed. 37, 2755–2794 (1998).

69. Narimatsu, Y. et al. An Atlas of Human Glycosylation Pathways Enables Display of the Human Glycome by Gene Engineered Cells. Mol. Cell 75, 394–407 (2019).

70. Tytgat, H.L.P. et al. Cytoplasmic glycoengineering enables biosynthesis of nanoscale glycoprotein assemblies. Nat. Commun. 10, 5403 (2019).

