## Supporting Information for "Genetically Encoded, Multivalent Liquid Glycan Array (LiGA)"

### Abbreviations

| | |
|---|---|
| BSA | Bovine Serum Albumin |
| CHO | Chinese Hamster ovary [cells] |
| CD | cluster of differentiation |
| ConA | Concanavalin A |
| CuAAC | Copper-catalyzed azide alkyne cycloaddition |
| DBCO | dibenzylcyclooctyne |
| DNA | Deoxyribonucleotide |
| DMEM | Dulbecco's Modified – Eagle's Medium |
| dsDNA | Double stranded DNA |
| EDTA | ethylenediaminetetraacetic acid |
| ELISA | Enzyme linked immuno-assay |
| FACS | Fluorescence activated cell sorting |
| Gzip | file format used for file compression / decompression |
| FASTQ | test-based format for storing DNA sequence and its **Q**uality score |
| HEPES | Hydroxyethyl piperazineethanesulfonic acid |
| HRPO | Horseradish peroxidase |
| LB | Lysogeny Broth |
| MALDI-TOF | Matrix-assisted laser desorption/ionization-Time of flight |
| MWCO | Molecular weight cutoff |
| NHS | N-hydroxysuccinimide |
| qPCR | Quantitative polymerase chain reaction |
| PBS | Phosphate buffered saline |
| PCR | Polymerase chain reaction |
| PEG | Poly(ethylene glycol) |
| PFU | Plaque forming units |
| RBC | red blood cells |
| rt | room temperature |
| TRIS | Tris(hydroxymethyl)aminomethane |

### 1. Biochemical Methods

**1.1** *Materials and general information.*
Sanger sequencing and deep sequencing was performed at the Molecular Biology Service Unit (University of Alberta) using Illumina NextSeq500 and MiniSeq. All DNA primers with ordered from Integrated DNA Technologies. Biochemical reagents were purchased from Thermo Fisher Scientific unless noted otherwise. HEPES buffer contains 20 mM HEPES, 150 mM NaCl, 2 mM CaCl2, pH 7.4. PBS buffer contains 137 mM NaCl, 10 mM Na2HPO4, 2.7 mM KCl, pH 7.4. Solutions used for phage work were sterilized by filtration through 0.22 µm filters. The mannose-binding protein concanavalin-A (ConA) was purchased from Sigma-Aldrich (#C2010). The carbohydrate recognition domain of the human Gal3 (G3C), residues 107−250 (MW: 16327 Da) was a generous gift of Christopher Cairo (University of Alberta). Biotinylated Cholera Toxin pentamer B subunit was a generous gift of Bruce Turnbull (University of Leeds). Murine monoclonal anti-A IgM and anti-B IgM were a generous gift of Lori West (University of Alberta). Origin of the 76 anomeric azido-glycans used in LiGA: (i) 67 was prepared by the Consortium for Functional Glycomics at the Scripps Research Institute (La Jolla, CA, USA); (ii) preparation of 5 glycans is described here; (ii) synthesis of 4 glycans was described elsewhere (**Table S2** and references within). The azidomethyl-1-thio-β-D-mannopyranoside[1] and N-succinimidyl 4-pentynoate (NP) used in pVIII modification studies and were donated by David Bundle (University of Alberta). MS-MALDI-TOF spectra were recorded on AB Sciex Voyager Elite MALDI, mass spectrometer equipped with MALDI-TOF pulsed nitrogen laser (337 nm) (3 ns pulse up to 300 µJ/pulse) operating in Full Scan MS in positive ionization mode. NanodropTM (Thermo Fisher) was used to measure the absorbance of protein and DNA solutions.

**1.2** *Construction of silent double barcoded (SDB) M13 phage library.*
A library of silent double barcode-codons (SDB) in the phage genome proximal to the pIII cloning site was created using the Gibson Assembly cloning kit (NEB#E5510) purchased from New England Biolabs. The SDB regions were introduced into M13KE vector using NEBuilder HiFi DNA Assembly (NEB# E2621S). As starting point, we used double stranded DNA of M13KE vector containing the stuffer sequence CAGTTTACGTAGCTGCATCAGGGTGGAGGT corresponding to the peptide QFT*LHQ, with * representing a stop codon. The insert fragment was PCR amplified using the primers **P1** and **P2** and the vector fragment was PCR amplified using primers **P3** and **P4**:

```
Name Sequence (5'->3'):
P1 GAGATTTTCAACGTGAAAAAACTNCTNTTYGCNATHCCNCTNGTGGTACCTTTCTATTCTCA
P2 TTAAGACTCCTTATTACGCAGTA
P3 TTGCTAACATACTGCGTAATAAG
P4 TTTTTTCACGTTGAAAATCTC
P5 GTGGTACCTTTCTATTCTCACTCGAGYGTNGARAARAAYGAYCARAARACNTAYCAYGCNGGNGGNGGNT-
CGGCCGAAACTGTTGAAAG
P6 CGAGTGAGAATAGAAAGGTAC
```

PCR was performed using 50 ng phage dsDNA with 1 mM dNTPs, 0.5 µM primers, 0.5 µL Phusion High Fidelity DNA polymerase in 1x PCR buffer (NEB #B0518S) in a total volume of 50 µL. The temperature cycling protocol was performed as follows:

a) 98 °C 3 min,

b) 98 °C 30 s,
c) 60 °C 30 s,
d) 72 °C 4 min s,
e) repeat b) - d) for 35 cycles,
f) 72 °C 10 min,
g) 4 °C hold.

PCR amplified fragments were treated with restriction enzyme DPN 1 (NEB #R0176S) and then gel purified. NEBuilder Hifi DNA assembly was then carried out following the manufacturer protocols by mixing 100 ng of vector, 4 ng insert, 10 µL of NEBuilder Hifi DNA assembly master mix, and deionized $H_2O$ up to a total volume of 20 µL. The resulting ligated DNA was transformed into *E.coli* K12 ER2738 and propagated overnight at 37 °C. The overnight culture was then centrifuged to separate bacteriophage from host cells. Deep sequencing of the resulting SB1 QFT*LHQ cloning vector with 6,144 theoretical sequence combinations in the leader region is available at: https://48hd.cloud/file/20161105-68OOooIC-NB. We observed that many of the 6,144 theoretically possible sequences were eliminated due to grow disadvantage and top 200 sequences occupied ~90% of the available diversity (Figure S1).

The cloning of SVEKNDQKTYHAGGG peptide was conducted as follows. Primers **P5** and **P2** were annealed to and amplified by PCR following the 35-cycle protocol described above to produce a dsDNA insert. The vector SB1 QFT*LHQ was PCR amplified using primers **P4** and **P6**. PCR fragment were processed using NEBuilder Hifi DNA assembly kit using steps described above. The resulting ligated DNA was transformed into electrocompetent cells *E.coli* SS320 (Lucigen). The resulting overnight culture was centrifuged to remove host cells and incubated with 5% PEG-8000, 0.5 M NaCl for 8 h at 4 °C, followed by 15 min centrifugation at 13000 g to concentrate released phage. PEG precipitated phage were re-suspended in PBS-Glycerol 50% and stored at -20 °C. M13-SDB-SVEKY is a library of chemically identical phage with 6,144 theoretical redundant sequence combinations SB-1 region (~200 practically observed), and $2.1\times10^6$ theoretical redundant sequence combinations in SVEK region yielding between $10^8$ to $10^{10}$ possible sequence combinations. Sequence of the vector containing M13-SDB-SVEKY library is available on GeneBank (#MN865131). Deep sequencing of the resulting library is available at: https://48hd.cloud/file/20161215-67OOooOO-NB

**1.3** *SDB clone isolation and amplification.*
The M13-SDB-SVEKY library described in the previous section was used to isolate each monoclonal silently encoded phage. A 10 µL aliquot of phage was diluted and plated at a density of 100 plaques per plate. Single colonies were manually picked, and individually transferred into a clean 1.7 mL plastic tube containing 0.5 mL of PBS-Glycerol 50% and incubated at room temperature for 30 min. The tubes were then placed in 55 °C heating block for 10 min to inactivate any remaining bacterial cells. After the incubation, 20 µL sample of each colony suspension was amplified for 4.5 h in 5 mL of LB supplemented with a 0.5 mL of log phase *E. coli* K12 ER2738. After amplification the phage clones were collected from the culture supernatant by centrifugation at 4500 g for 10 min. Next, bacterial cell pellet and culture supernatant were processed separately. The supernatant was incubated with 5% PEG-8000, 0.5 M NaCl for 8 h at 4 °C, followed by 15 min centrifugation at 13000 g to precipitate the viral particles. The phages were re-suspended into 1 mL PBS-Glycerol 50%, titered, and stored at -20 °C until further use. The bacterial cell pellet was processed for phage-DNA extraction using GeneJET Plasmid Miniprep kit (Thermo Fisher, #K0502).

For SDB identification, a sample of 400 ng of the phage DNA was submitted for Sanger sequencing. We selected the clones that contained three or more base pair substitutions from one another (i.e., Hamming distance (H)≥3). H≥3 permits correction of any point mutations that may have arisen during the analysis by deep sequencing. A list of isolated clones is available in **Supplementary Table S1**. Solution of clonal phage (2 µL) was PCR amplified using barcoded sequencing primers (Fig. S1G-H**)** using a protocol similar to that described in *"PCR Protocol Section*" and analyzed by Illumina sequencing. The results of the Illumina sequencing of the isolated clones is available at http://ligacloud.ca/ by searching for specific SDB number. Example of searching for "SDB50":
http://ligacloud.ca/searchDB?search=SDB50

**1.4** *Construction of phage that transduce mCherry and mNeonGreen fluorescent proteins*.
We produced filamentous phage vector that contains the gene for fluorescent protein mCherry and mNeonGreen cloned in place of the lacZα fragment. Vectors containing mCherry and mNeonGreen genes were generous gifts from Robert Campbell (University of Alberta). Both constructs were built using NEBuilder Hifi DNA assembly ligation of fluorescent protein (FP) fragment and M13 fragment. Vector pBAD-mCherry was used as the source for the mCherry insert, whereas Vector pBAD-mNeonGreen was used as a source for the mNeonGreen insert. FP fragments were PCR amplified following the protocol described in section "*Construction of silent distal barcoded (SDB) M13 phage library*" using primers **P7** and **P8**, M13-SDB-SVEKY with sequence CTTCTATTTGCTATTCCTCTA was used to produce M13 fragment for the mCherry construct, and M13-SDB-SVEKY with sequence CTACTGTTCGCAATCCCGCTA was used to produce M13 fragment for the mNeonGreen construct and both were amplified using primers **P9** and **P10**.

| **Name:** | Sequence (5'->3') |
|---|---|
| **P7** | GCGGATAACAATTTCACACAGGAAACAGCTATGGTGAGCAAGGGCGAG |
| **P8** | TTAAATTTTTGTTAAATCAGCTCATTTTTTACTTGTACAGCTCGTCCA |
| **P9** | AAAATGAGCTGATTTAACAAAAATTTAA |
| **P10** | AGCTGTTTCCTGTGTGAAAT |

The sequences of the resulting vectors are available as
M13 mNeonGreen SDB SVEKY: GeneBank MN865132
M13 mCherry SDB SVEKY: GeneBank MN872303

**1.5** *Monitoring phage viability under CuAAC reaction*.
The CuAAC reaction followed the published protocol[2] with minor modifications. To prepare bathophenantroline/$Cu^+$ catalyst, $Cu_2SO_4 \cdot 5H_2O$ (10 mg) and bathophenantroline sulfonate (64.4 mg, GFS Chemicals Inc.) were added to a glass vial (4 mL) containing Tris buffer (1 mL, 0.2 M Tris, pH 8.0). Copper powder (50 mg) was added; the vial was closed with a rubber septa and purged with argon and rotated for 2 h. The appearance of a dark green color indicated the reduction of copper II to copper I. Solution containing the bathophenantroline/$Cu^+$ catalyst was used on the same day.
A solution of clone SDB3 (Table S1) (150 µL, 3.4 x $10^{13}$ PFU/mL, PBS pH 7.4) was concentrated using 0.5 mL Amicon filter (10 MWCO, Sigma, #UFC5010) and resuspended in Tris-borate 200 mM pH 8.0, to a final volume of 150 µL. The phage solution was titered by plaque forming assay (Fig. S2C-D). To the solutions of SDB3 (50 µL, in Tris-Borate, pH 8.0) in 1.7 mL Eppendorf™ tubes, NHS-4-pentynoate (50 mM in DMF) was added to afford a reaction

mixture with 1 mM final concentration of NHS-4-pentynoate. After incubation at rt for 1 h with gentle rocking, the reaction mixtures were purified on Zeba Spin Desalting Columns (40K MWCO, Thermo Fisher, #87766) to eliminate unreacted linker and 10 μL of filtrate was analyzed by plaque forming assay (Fig. S2C-D). To the filtered phage solution, azido-glycan Manb1-Ss1 was added to afford the 2 mM final concentration followed by ~1 mg of copper powder. The CuAAC reaction was then initiated by adding 3 μL of Bathophenantroline/$Cu^+$ catalyst. After incubation for 8 h at 4 °C under argon atmosphere, the mixture was purified on Zeba Spin column,[a] and the filtrate was analyzed by plaque forming assay (Fig. S2C-D).

**1.6** *Monitoring phage viability under SPAAC reaction.*
To a solution of phage clone SBD3 (50 μL, 3.4 x $10^{13}$ PFU/mL PBS pH 7.4) in 1.7 mL Eppendorf™ tubes, DBCO-NHS (2.5 μL, 20 mM in DMF) was added to afford a reaction mixture with 1 mM final concentration of DBCO-NHS. Three analogous reactions were tested in parallel. Each reaction mixture was incubated at rt for 1 h with gentle rocking. The reaction mixtures were purified on Zeba Spin column to remove the unreacted and hydrolysed DBCO-NHS and 10 μL of the filtrate was analyzed by plaque forming assay (Fig. S2C-D). To the filtrate, solution of azido-glycan Manβ1-Ss1 was added to afford the 2 mM final concentration. The solution was incubated for 8 h at 4 °C. The reaction mixture was purified on Zeba Spin column and analyzed by plaque forming assay (Fig. S2C-D).

**1.7** *Analysis of glycosylation of phage samples by MALDI-TOF MS.*
The sinapinic acid matrix[3] was formed by deposition of two layers. Layer1 was prepared as a 10mg/mL solution of sinapinic acid (Sigma, #D7927) in acetone-methanol (4:1). Layer2 was prepared as 10 mg/mL solution of sinapinic acid in acetonitrile:water (1:1) with 0.1% TFA. In a typical sample preparation, 2 μL of phage solution in PBS was combined with 4 μL of layer2, then a mixture of 1:1 layer1:layer2+phage was deposited in that order onto the MALDI inlet plate ensuring that layer 1 is completely dry before adding layer2+phage. The spots were washed with 10 μL of water with 0.1% TFA to remove salt ions from the PBS. To estimate the ratio of modified to unmodified pVIII, we fit and plotted the data using MatLab.

**1.8** *Quantification of phage-surface chemical modifications.*
This procedure describes example of modification of phage with different surface density of glycan using phage clone SDB3, DBCO-NHS and the azido-glycan Manb1-Ss1. A solution of SDB3 phage (60 μL, ~$10^{13}$ PFU/mL in PBS, pH 7.4) was distributed into three Eppendorf™ tubes and solution of DBCO was added to the tubes to yield 0.5 or 1.0 or 1.5 mM final concentration. After incubation for 1 h at rt with gentle rocking, the reaction mixtures were purified on Zeba Spin column. Each experiment was analyzed in technical duplicates. 10 μL of the filtrate was analyzed by MALDI-TOF (Fig. 1E). To the filtrate, solution of azido-glycan Manβ1-Ss1 (10 μL, 10 mM in DMF) was added to afford the 2 mM final concentration. The solution was incubated for 12 h at 4 °C. The reaction mixture was purified on Zeba Spin column and analyzed by MALDI-TOF.

**1.9** *Modification of phage clones with glycans to build components of LiGA.*
A solution of each SDB phage clone ($10^{12}$-$10^{13}$ PFU/mL in PBS) was combined with DCBO-NHS (20 mM in DMF) to afford a 1 mM concentration of DCBO-NHS in reaction mixture, which typically yields 25% of pVIII modification after 45 min incubation. Solutions of glycan-azide (10 mM in Nuclease Free $H_2O$) were added directly to the reaction mixture to afford a 2

mM concentration of glycan-azide and the solutions were further incubated overnight at 4 °C. Conjugated phage were stored at 4 °C in the solution that contained an excess of azido-glycan. LiGA mixture was prepared by combining these stock solutions followed by purification of the mixture by Zeba™ Spin Desalting column (7K MWCO, 0.5 mL, Thermo Fisher cat #89882).

*Alternative Procedure*: A solution of SDB phage clone ($10^{12}$-$10^{13}$ PFU/mL in PBS) was combined with DCBO-NHS (20 mM in DMF) to afford a 0.2-2.0 mM concentration of DCBO-NHS in reaction mixture, which typically yields 5-50% of pVIII modification after 45 min of incubation. After conjugation of DBCO-NHS, each clone was individually purified on a Zeba™ column following the manufacturer instructions. Solutions of azido-glycans (10 mM stock in Nuclease Free $H_2O$) were added to the filtrates to afford a 2 mM concentration of glycan-azide and the solutions were further incubated overnight at 4 °C. All chemical reactions were verified and quantified by MALDI-TOF as described on the previous sections. If reactions were incomplete and residual pVIII-DBCO peak was detected in MALDI, we added additional amount of azido glycan and extended the incubation. If reactions were completed, the conjugates were purified by Zeba column and stored at 4 °C or supplemented by glycerol and stored as 50% glycerol stock at -20 °C. LiGA mixture was prepared by combining these stock solutions.

**1.10** *Preparation of LiGA from glycosylated clones*.
In a typical protocol, LiGA was prepared by mixing $10^8$ PFU of desired glycan-phage conjugates into a single tube. The mixture was characterized by titering and N×$10^6$ PFU (N = glycan-phage conjugates) was used for a typical lectin or cell-binding experiment. Each unique LiGA mixture was assigned a two-letter identifier (e.g., “YZ”) and a “dictionary” (e.g., YZ.xlsx), a table that describes the correspondence between the DNA barcodes and the glycans in the LiGA mixture (**Table S3**). These dictionaries were subsequently used to translate from nucleotide sequences in the deep-sequencing files to the corresponding glycans (including density). Examples of dictionaries are available as part of the supporting information.

In characterizing the LiGA mixture by deep sequencing, we noted that the mixing of the solutions matched by the titer of phage stock did not afford uniform distribution of barcodes after sequencing. The composition of each naïve library or a naïve library binding to a control target, thus, was used as normalization factor in each experiment. Dictionaries for LiGA mixtures relevant to this manuscript are available in Data/LiGA Dictionaries/ folder. Naïve compositions of these mixtures and references to the deep-sequencing data on http://ligacloud.ca are available in **Table S3**.

| SDB_id | SB-1 region | SB-2 region |
|---|---|---|
| SDB1 | CTGCTGTTCGCAATACCACTC | AGTGTGGAGAAGAATGATCAGAAGACTTATCATGCGGGTGGAGGT |
| SDB2 | CTTCTATTCGCAATTCCGCTC | AGTGTGGAGAAGAATGATCAGAAGACTTATCATGCGGGTGGAGGT |
| SDB3 | CTGCTTTTCGCAATTCCGCTT | AGTGTGGAGAAGAATGATCAGAAGACTTATCATGCGGGTGGAGGT |
| SDB6 | CTGCTGTTTGCGATTCCACTG | AGTGTGGAGAAGAATGATCAGAAGACTTATCATGCGGGTGGAGGT |
| SDB9 | CTTCTTTTTGCAATTCCTCTA | AGTGTGGAGAAGAATGATCAGAAGACTTATCATGCGGGTGGAGGT |
| SDB10 | CTACTGTTTGCTATACCGCTG | AGTGTGGAGAAGAATGATCAGAAGACTTATCATGCGGGTGGAGGT |
| SDB12 | CTTCTGTTCGCGATACCTCTA | AGTGTGGAGAAGAATGATCAGAAGACTTATCATGCGGGTGGAGGT |
| SDB13 | CTACTTTTCGCAATTCCTCTG | AGTGTGGAGAAGAATGATCAGAAGACTTATCATGCGGGTGGAGGT |
| SDB15 | CTGCTGTTCGCCATACCCCTT | AGTGTGGAGAAGAATGATCAGAAGACTTATCATGCGGGTGGAGGT |
| SDB17 | CTACTCTTCGCGATTCCGCTT | AGTGTGGAGAAGAATGATCAGAAGACTTATCATGCGGGTGGAGGT |
| SDB18 | CTGCTGTTTGCTATCCCTCTG | AGTGTGGAGAAGAATGATCAGAAGACTTATCATGCGGGTGGAGGT |
| SDB20 | CTGCTCTTTGCCATCCCGCTT | AGTGTGGAGAAGAATGATCAGAAGACTTATCATGCGGGTGGAGGT |
| SDB21 | CTACTCTTTGCAATTCCCCTT | AGTGTGGAGAAGAATGATCAGAAGACTTATCATGCGGGTGGAGGT |
| SDB22 | CTACTGTTTGCTATCCCACTT | AGTGTGGAGAAGAATGATCAGAAGACTTATCATGCGGGTGGAGGT |
| SDB23 | CTGCTCTTTGCAATACCTCTT | AGTGTGGAGAAGAATGATCAGAAGACTTATCATGCGGGTGGAGGT |
| SDB24 | CTACTATTCGCGATCCCGCTC | AGTGTGGAGAAGAATGATCAGAAGACTTATCATGCGGGTGGAGGT |
| SDB26 | CTGCTATTCGCTATCCCACTC | AGTGTGGAGAAGAATGATCAGAAGACTTATCATGCGGGTGGAGGT |
| SDB29 | CTGCTATTTGCGATCCCGCTG | AGTGTGGAGAAGAATGATCAGAAGACTTATCATGCGGGTGGAGGT |
| SDB34 | CTTCTTTTTGCGATTCCGCTG | AGTGTGGAGAAGAATGATCAGAAGACTTATCATGCGGGTGGAGGT |
| SDB35 | CTGCTCTTCGCTATTCCACTT | AGTGTGGAGAAGAATGATCAGAAGACTTATCATGCGGGTGGAGGT |
| SDB40 | CTTCTGTTCGCTATTCCGCTT | AGCGTGGAAAAGAACGATCAAAAGACCTATCACGCCGGGGGAGGG |
| SDB42 | CTTCTGTTTGCAATACCCCTG | AGCGTGGAGAAGAATGATCAAAAGACGTACCATGCGGGGGGAGGG |
| SDB45 | CTGCTGTTTGCGATTCCTCTG | AGCGTGGAAAAAAATGACCAAAAAACCTACCATGCAGGGGGGGGA |
| SDB47 | TTATTATTCGCAATACCGCTA | AGTGTTGAGAAGAATGACCAGAAGACGTATCACGCCGGGGGAGGG |
| SDB48 | TTATTATTCGCAATTCCTTTA | AGCGTAGAGAAAAACGACCAGAAGACCTATCACGCGGGAGGAGGT |
| SDB49 | CTACTGTTCGCTATCCCGCTG | AGTGTGGAAAAGAATGATCAGAAGACTTACCACGCTGGTGGTGGG |
| SDB51 | CTGCTCTTTGCTATACCCCTA | AGCGTCGAGAAAAATGATCAAAAAACCTATCACGCCGGGGGGGGA |
| SDB52 | CTACTGTTTGCTATCCCGCTG | AGTGTAGAGAAGAATGACCAAAAGACATATCATGCGGGAGGAGGT |
| SDB56 | CTACTGTTTGCTATACCTCTT | AGCGTTGAAAAAAATGACCAGAAAACTTACCATGCGGGTGGAGGT |
| SDB57 | TTATTATTCGCAATTCCTTTA | AGCGTCGAGAAGAACGACCAAAAAACCTATCACGCAGGTGGCGGG |
| SDB58 | CTACTCTTCGCTATACCCCTC | AGCGTGGAGAAAAACGACCAAAAGACCTACCATGCAGGTGGTGGC |
| SDB59 | TTATTATTCGCAATTCCTTTA | AGTGTCGAAAAGAATGACCAAAAAACTTATCATGCAGGTGGGGGA |
| SDB60 | CTACTGTTCGCCATCCCTCTG | AGTGTAGAAAAGAACGACCAAAAGACTTACCATGCTGGTGGGGGG |
| SDB66 | TTATTATTCGCAATTCCTTTA | AGTGTCGAGAAGAACGATCAGAAGACCTACCACGCAGGGGGGGGT |
| SDB67 | CTTCTGTTCGCTATACCTCTC | AGCGTGGAGAAGAATGACCAGAAAACTTATCACGCAGGAGGTGGA |
| SDB68 | CTACTATTCGCGATCCCCCTC | AGCGTGGAGAAGAACGACCAGAAGACGTATCACGCAGGGGGGGGG |
| SDB69 | CTTCTGTTTGCGATTCCACTG | AGCGTGGAGAAGAACGACCAGAAAACATACCATGCTGGTGGTGGA |
| SDB70 | CTACTGTTCGCAATCCCGCTC | AGTGTTGAAAAAAACGATCAAAAAACGTATCATGCTGGTGGAGGT |
| SDB71 | CTGCTTTTTGCTATTCCTCTG | AGTGTTGAAAAAAACGATCAGAAGACTTATCACGCGGGGGGCGGG |
| SDB73 | TTATTATTCGCAATTCCTTTA | AGCGTTGAAAAAAACGATCAAAAGACATATCACGCTGGCGGAGGA |
| SDB74 | CTACTCTTTGCGATCCCTCTG | AGTGTCGAAAAAAATGACCAAAAGACCTATCACGCGGGCGGGGGA |
| SDB77 | CTGCTATTTGCCATCCCGCTG | AGCGTCGAGAAAAACGATCAAAAGACATACCATGCCGGCGGAGGA |
| SDB79 | CTACTGTTCGCAATACCTCTC | AGCGTGGAGAAAAACGACCAGAAAACGTATCATGCTGGTGGTGGA |
| SDB80 | CTTCTGTTCGCGATCCCCCTA | AGTGTAGAGAAGAATGATCAGAAAACGTATCATGCGGGAGGGGGT |
| SDB81 | CTACTCTTTGCTATTCCGCTT | AGCGTGGAGAAGAATGATCAGAAAACGTACCACGCCGGTGGTGGC |
| SDB82 | TTATTATTCGCAATTCCTTTA | AGCGTGGAGAAGAACGACCAAAAGACTTATCACGCGGGCGGGGGG |
| SDB83 | CTGCTCTTCGCGATCCCTCTG | AGTGTCGAAAAGAATGATCAAAAAACGTATCATGCGGGCGGTGGT |
| SDB87 | CTACTCTTCGCTATACCCCTC | AGCGTAGAAAAAAATGACCAAAAGACCTACCATGCGGGAGGTGGG |
| SDB88 | CTGCTGTTCGCGATTCCTCTA | AGTGTAGAGAAGAATGACCAGAAGACCTACCACGCGGGCGGGGGG |
| SDB99 | CTTCTCTTCGCGATACCGCTA | AGTGTTGAGAAAAATGATCAAAAAACGTATCATGCGGGTGGTGGG |
| SDB103 | TTATTATTCGCAATTCCTTTA | AGTGTTGAAAAAAACGACCAGAAGACTTATCACGCGGGTGGGGGG |
| SDB105 | CTGCTTTTTGCTATTCCTCTT | AGTGTCGAGAAGAATGATCAAAAAACTTACCACGCGGGTGGTGGA |
| SDB112 | CTACTGTTCGCGATCCCACTG | AGTGTAGAAAAAAATGACCAGAAGACGTATCACGCAGGCGGAGGC |
| SDB113 | CTACTGTTTGCCATTCCCCTG | AGTGTCGAAAAAAATGATCAGAAAACTTATCACGCAGGAGGGGGG |
| SDB115 | CTACTGTTCGCTATCCCGCTG | AGTGTCGAAAAAAATGACCAGAAAACGTATCATGCGGGGGGGGGT |
| SDB118 | CTACTCTTCGCGATACCCCTC | AGTGTCGAAAAGAATGATCAGAAGACTTATCATGCAGGTGGTGGT |
| SDB120 | CTGCTGTTTGCTATTCCACTA | AGTGTTGAAAAAAACGATCAGAAGACTTATCATGCTGGGGGAGGA |
| SDB126 | CTGCTCTTTGCAATTCCGCTG | AGTGTAGAAAAAAATGATCAGAAAACATACCATGCTGGTGGCGGA |
| SDB127 | CTGCTATTTGCCATCCCACTA | AGTGTTGAGAAGAACGATCAGAAAACGTATCATGCAGGTGGTGGA |
| SDB149 | CTACTCTTTGCAATCCCGCTT | AGCGTGGAGAAGAACGACCAGAAGACTTACCACGCGGGTGGTGGG |
| SDB150 | CTGCTCTTTGCTATCCCTCTA | AGTGTTGAAAAAAACGACCAGAAGACTTATCACGCAGGTGGTGGG |
| SDB152 | CTGCTTTTTGCTATACCTCTC | AGCGTAGAAAAGAATGACCAGAAGACGTACCATGCCGGTGGGGGG |

**Supplementary Table S1.** Examples of silent double barcodes (SDB) used to build LiGA Locations of SB1 and SB2 regions in M13 genome is illustrated in Fig. S1.

| | IUPAC | Common | Ref | MJ | MK | YG | YO | YT | YU | YX | YY | YZ |
|---|---|---|---|---|---|---|---|---|---|---|---|---|
| 1 | Galfβ1-5Galfβ-S8 | (Galf)2 | **SI** | | | | | | | | | |
| 2 | Galfβ1-5Galfβ1-5Galfβ-S8 | (Galf)3 | **SI** | | | | | | | | | |
| 3 | Galfβ1-5Galfβ1-5Galfβ1-5Galfβ-S8 | (Galf)4 | **SI** | | | | | | | | | |
| 4 | Manα1-6[Manα1-3]Mana-S6 | (Man)3 | **Ref**[4] | | | | | | | | | |
| 5 | Galα1-3[Fucα1-2]Galβ1-4[Fucα1-3]GlcNAcβ-Sp | 2'F-B type 2 | Te262 | | | | | | | | | |
| 6 | Fucα1-2Galβ1-4Glcβ-Sp | 2'FL | Tr120 | | | | | | | | | |
| 7 | KDNα2-3Galβ1-3GlcNAcβ-Sp | 3'-KDNLec | Tr48 | | | | | | | | | |
| 8 | GlcNAcβ1-3Galβ1-4GlcNAcβ1-3Galβ1-4GlcNAcβ-Sp | 3'GN-Di-LN | Te99 | | | | | | | | | |
| 9 | KDNα2-3Galβ1-4GlcNAcβ-Sp | 3'KDNLN | Tr47 | | | | | | | | | |
| 10 | Neu5Acα2-3Galβ1-3GlcNAcβ1-3Galβ1-3GlcNAcβ-Sp | 3'S-Di-Le$^{C}$ | Te321 | | | | | | | | | |
| 11 | Neu5Acα2-3(Galβ1-4[Fucα1-3]GlcNAcβ1-3)2β-Sp | 3'S-Di-Le$^{X}$ | Te140 | | | | | | | | | |
| 12 | Neu5Acα2-3(Galβ1-4[Fucα1-3]GlcNAcβ1-3)3β-Sp | 3'S-Tri-Le$^{X}$ | Te193 | | | | | | | | | |
| 13 | Neu5Gcα2-3Galβ1-4Glcβ-Sp | 3'SL (Gc) | Tr39 | | | | | | | | | |
| 14 | Neu5Gcα2-3Galβ1-3GlcNAcβ-Sp | 3'SLec (Gc) | Tr41 | | | | | | | | | |
| 15 | Neu5Acα2-3Galβ1-3GlcNAcβ1-3Galβ1-4GlcNAcβ-Sp | 3'SLecLN | Te288 | | | | | | | | | |
| 16 | Neu5Acα2-3Galβ1-4[Fucα1-3]GlcNAcβ-Sp | 3'SLex | Te64 | | | | | | | | | |
| 17 | Neu5Acα2-3Galβ1-4GlcNAcβ-Sp | 3'SLN | Tr33 | | | | | | | | | |
| 18 | Neu5Gcα2-3Galβ1-4GlcNAcβ-Sp | 3'SLN (Gc) | Tr40 | | | | | | | | | |
| 19 | Neu5Acα2-6(Galβ1-4GlcNAcβ1-3)2β-Sp | 6'S-Di-LN | Te176 | | | | | | | | | |
| 20 | Neu5Acα2-6GalNAcβ1-4GlcNAcβ-Sp | 6'SLDN | Tr269 | | | | | | | | | |
| 21 | Neu5Acα2-6Galβ1-4GlcNAcβ-Sp | 6'SLN | Tr36 | | | | | | | | | |
| 22 | Neu5Gcα2-6Galβ1-4GlcNAcβ-Sp | 6'SLN (Gc) | Tr43 | | | | | | | | | |
| 23 | Neu5,9Acα2-3Galβ1-4GlcNAcβ-Sp | 9OAc3'SLe$^{C}$ | Tr323 | | | | | | | | | |
| 24 | GalNAcα1-3[Fucα1-2]Galβ1-4Glcβ-Sp | A tetra L | Te224 | | | | | | | | | |
| 25 | GalNAcα1-3[Fucα1-2]Galβ1-3GlcNAcβ-Sp | A tetra type 1 | Te259 | | | | | | | | | |
| 26 | GalNAcα1-3[Fucα1-2]Galβ1-4GlcNAcβ-Sp | A tetra type 2 | Te222 | | | | | | | | | |
| 27 | Manα-S6 | aMan | **Ref**[5] | | | | | | | | | |
| 28 | Galα1-3[Fucα1-2]Galβ1-3GlcNAcβ-Sp | B tetra type 1 | Te258 | | | | | | | | | |
| 29 | Galα1-3[Fucα1-2]Galβ1-4GlcNAcβ-Sp | B tetra type 2 | Te223 | | | | | | | | | |
| 30 | Manβ-S0 | bMan | **Ref**[6] | | | | | | | | | |
| 31 | Neu5Acα2-3[GalNAcβ1-4]Galβ1-4GlcNAcβ-Sp | CT\|Sda | Te201 | | | | | | | | | |
| 32 | (Galβ1-4[Fucα1-3]GlcNAcβ1-3)2β-Sp | Di-Le$^{X}$ | Te101 | | | | | | | | | |
| 33 | (Galβ1-4GlcNAcβ1-3)2β-Sp | Di-LN | Te98 | | | | | | | | | |
| 34 | Fucα1-2Galβ-Sp | Di-N3 | Di-N3 | | | | | | | | | |
| 35 | Galβ-P4 | Gal | **SI** | | | | | | | | | |
| 36 | Galα1-3Galβ1-3GlcNAcβ-Sp | Gala3-type1 | Tr260 | | | | | | | | | |
| 37 | Galα1-3Galβ1-4[Fucα1-3]GlcNAcβ-Sp | Gala3Lex | Te221 | | | | | | | | | |
| 38 | Galfβ-S8 | Galf | **SI** | | | | | | | | | |
| 39 | Galα1-3Galβ1-4Glcβ-Sp | Galili-tri | Tr59 | | | | | | | | | |
| 40 | Neu5Acα2-3[Neu5Acα2-3Galβ1-3GalNAcβ1-4]Galβ1-4Glcβ-Sp | GD1a | Te303 | | | | | | | | | |
| 41 | Neu5Acα2-8Neu5Acα2-3[GalNAcβ1-4]Galβ1-4Glcβ-Sp | GD2 | Te78 | | | | | | | | | |
| 42 | Neu5Acα2-8Neu5Acα2-3Galβ1-4Glcβ-Sp | GD3 | Te79 | | | | | | | | | |
| 43 | GalNAcβ1-3Galα1-4Galβ1-4Glcβ-Sp | Globoside-P | Te272 | | | | | | | | | |
| 44 | Neu5Acα2-3[Galβ1-3GalNAcβ1-4]Galβ1-4Glcβ-Sp | GM1 | Te75 | | | | | | | | | |
| 45 | Neu5Acα2-3Galβ1-4Glcβ-Sp | GM3 | Tr32 | | | | | | | | | |
| 46 | GlcNAcβ1-3Galβ1-4GlcNAcβ-Sp | GNLN | Tr55 | | | | | | | | | |
| 47 | Neu5Acα2-8Neu5Acα2-8Neu5Acα2-8Neu5Acα2-3[GalNAcβ1-4]Galβ1-4Glcβ-Sp | GQ2 | Te306 | | | | | | | | | |
| 48 | Neu5Acα2-8Neu5Acα2-8Neu5Acα2-3[GalNAcβ1-4]Galβ1-4Glcβ-Sp | GT2 | Te119 | | | | | | | | | |
| 49 | Neu5Acα2-8Neu5Acα2-8Neu5Acα2-3Galβ1-4Glcβ-Sp | GT3 | Te97 | | | | | | | | | |
| 50 | Fucα1-2Galβ1-3GlcNAcβ-Sp | H-type1 | Tr116 | | | | | | | | | |
| 51 | Fucα1-2Galβ1-4GlcNAcβ-Sp | H-type2 | Tr117 | | | | | | | | | |
| 52 | Fucα1-2Galβ1-4GlcNAcβ1-3Galβ1-4GlcNAcβ-Sp | H2 | Te134 | | | | | | | | | |
| 53 | Fucα1-2Galβ1-4GlcNAcβ1-3 Galβ1-4GlcNAcβ1-3Galβ1-4GlcNAcβ-Sp | H3 | Te135 | | | | | | | | | |
| 54 | Galβ1-4Glcβ-Sp | Lac | D9 | | | | | | | | | |
| 55 | Galβ1-4Glcβ-P4 | Lac-peg4 | **Ref**[7] | | | | | | | | | |
| 56 | GalNAcβ1-4GlcNAcβ-Sp | LacDiNAc | D21 | | | | | | | | | |
| 57 | Galβ1-3[Fucα1-4]GlcNAcβ-Sp | Le$^{A}$ | Tr57 | | | | | | | | | |
| 58 | Galβ1-3GlcNAcβ-Sp | Le$^{C}$ | D8 | | | | | | | | | |
| 59 | Galβ1-3GlcNAcβ1-3Galβ1-4[Fucα1-3]GlcNAcβ-Sp | Le$^{C}$-Le$^{X}$ | Te286 | | | | | | | | | |
| 60 | Galβ1-4[Fucα1-3]GlcNAcβ1-3Galβ1-3[Fucα1-4]GlcNAcβ-Sp | Le$^{X}$-Le$^{A}$ | Te319 | | | | | | | | | |
| 61 | Fucα1-2Galβ1-4[Fucα1-3]GlcNAcβ-Sp | Le$^{Y}$ | Te118 | | | | | | | | | |
| 62 | Fucα1-2Galβ1-4[Fucα1-3]GlcNAcβ1-3Galβ1-4[Fucα1-3]GlcNAcβ-Sp | Le$^{Y}$-Le$^{X}$ | Te212 | | | | | | | | | |
| 63 | Galβ1-4GlcNAcβ1-3Galβ1-4Glcβ-Sp | LNnT | Te72 | | | | | | | | | |
| 64 | GlcNAcβ1-3Galβ1-4Glcβ-Sp | LNT-2 | Tr54 | | | | | | | | | |
| 65 | Galβ1-3GlcNAcβ1-3Galβ1-4GlcNAcβ-Sp | LNT-NAc | Te271 | | | | | | | | | |
| 66 | Galα1-4Galβ1-4GlcNAcβ1-3Galβ1-4Glcβ-Sp | P1 penta | Te327 | | | | | | | | | |
| 67 | GalNAcβ1-3Galα1-4Galβ1-4GlcNAcβ-Sp | P1 tetra | Te289 | | | | | | | | | |
| 68 | Galα1-4Galβ1-4GlcNAcβ-Sp | P1 tri | Tr62 | | | | | | | | | |
| 69 | Galβ1-3GalNAcβ1-3Galα1-4Galβ1-4GlcNAcβ-Sp | P1x penta | Te302 | | | | | | | | | |
| 70 | Neu5Acα2-3Galβ1-3GalNAcβ1-3Galα1-4Galβ1-4Glcβ-Sp | SSEA-4 | Te291 | | | | | | | | | |
| 71 | Neu5Acα2-8Neu5Acα2-8Neu5Acα2-8Neu5Acα2-3Galβ1-4Glcβ-Sp | TetraSLac | Te305 | | | | | | | | | |
| 72 | GalNAcα1-3[Fucα1-2]Galβ-Sp | Tri-AN3 | Tri-AN3 | | | | | | | | | |
| 73 | Galα1-3[Fucα1-2]Galβ-Sp | Tri-BN3 | Tri-BN3 | | | | | | | | | |
| 74 | Galβ1-4[Fucα1-3]GlcNAcβ1-3Galβ1-4[Fucα1-3]GlcNAcβ1-3Galβ1-4[Fucα1-3]GlcNAcβ-Sp | Tri-Le$^{X}$ | Te102 | | | | | | | | | |
| 75 | Galβ1-4GlcNAcβ1-3Galβ1-4GlcNAcβ1-3Galβ1-4GlcNAcβ-Sp | Tri-LN | Te100 | | | | | | | | | |

**Supplementary Table S2**. List of glycans, their source and usage in LiGA.

Linker abbreviations:

Sp -O-$(CH_2)_2$-$N_3$; S0 -$N_3$;

S6 -O-$(CH_2)_6$-$N_3$; S8 -O-$(CH_2)_8$-$N_3$;

P4: -(O-$CH_2$ -$CH_2$)$_4$-$N_3$ Ss1 -S-$CH_2$-$N_3$;

Synthesis of reagents denoted by "**SI**" is described in the supporting information.

Glycans with D/Te/Tr labels were prepared by Consortium of Functional Glycomics (CFG). Synthesis of glycans with "**Ref**" label is described in the indicated reference. Two letter identifiers (MJ, MK, YG, etc) describe specific LiGA mixture; their usage is further described in **Table S3**. Green rectangle indicates that the glycan was included in the specific LiGA. Synthesis of the CFG glycans Tr33[8], Tr36[8], Te303[9], Te305[9], Te306[9], Te271[10], are available in the corresponding publications. Preparation of Glycans with D/Te/Tr labels can be found on the CFG website using the corresponding S-core as the reference on http://www.functionalglycomics.org/. For example, synthesis of Te262 can be found using "S245" as a search term. The correspondence between D/Te/Tr labels, S-references and direct URL link is available below. This information is tabulated in the SyntheticGlycanTable.xlsx file available in the Data.zip.

| | | |
|---|---|---|
| Te262 | S245 | http://www.functionalglycomics.org/glycomics/CarbProductionServlet?pageType=view&view=view&operationType=view&sideMenu=no&carbProductionId=carbSynthe_493_D000_S245-1_07152004 |
| Tr120 | S133 | http://www.functionalglycomics.org/glycomics/CarbProductionServlet?pageType=view&view=view&operationType=view&sideMenu=no&carbProductionId=carbSynthe_0172_D000_S133-1_11292002 |
| Tr48 | S54 | http://www.functionalglycomics.org/glycomics/CarbProductionServlet?pageType=view&view=view&operationType=view&sideMenu=no&carbProductionId=carbSynthe_0180_D000_S54-1_03212002 |
| Te99 | S199 | http://www.functionalglycomics.org/glycomics/CarbProductionServlet?pageType=view&view=view&operationType=view&sideMenu=no&carbProductionId=carbSynthe_465_D000_S199-1_01132004 |
| Tr47 | S53 | http://www.functionalglycomics.org/glycomics/CarbProductionServlet?pageType=view&view=view&operationType=view&sideMenu=no&carbProductionId=carbSynthe_0181_D000_S53-1_03212002 |
| Te321 | S304 | http://www.functionalglycomics.org/glycomics/CarbProductionServlet?pageType=view&view=view&operationType=view&sideMenu=no&carbProductionId=carbSynthe_942_D000_S304-1_07262005 |
| Te140 | S204 | http://www.functionalglycomics.org/glycomics/CarbProductionServlet?pageType=view&view=view&operationType=view&sideMenu=no&carbProductionId=carbSynthe_0561_D000_S204-1_01052004 |
| Te193 | S174 | http://www.functionalglycomics.org/glycomics/CarbProductionServlet?pageType=view&view=view&operationType=view&sideMenu=no&carbProductionId=carbSynthe_495_D000_S174-1_05292003 |
| Tr39 | S65 | http://www.functionalglycomics.org/glycomics/CarbProductionServlet?pageType=view&view=view&operationType=view&sideMenu=no&carbProductionId=carbSynthe_0189_D000_S65-1_05012002 |
| Tr41 | S67 | http://www.functionalglycomics.org/glycomics/CarbProductionServlet?pageType=view&view=view&operationType=view&sideMenu=no&carbProductionId=carbSynthe_0188_D000_S67-1_12102003 |
| Te64 | S143 | http://www.functionalglycomics.org/glycomics/CarbProductionServlet?pageType=view&view=view&operationType=view&sideMenu=no&carbProductionId=carbSynthe_487_D000_S143-1_05192003 |
| Tr40 | S66 | http://www.functionalglycomics.org/glycomics/CarbProductionServlet?pageType=view&view=view&operationType=view&sideMenu=no&carbProductionId=carbSynthe_0190_D000_S66-1_05012002 |
| Te176 | S141 | http://www.functionalglycomics.org/glycomics/CarbProductionServlet?pageType=view&view=view&operationType=view&sideMenu=no&carbProductionId=carbSynthe_0199_D000_S141-1_12052002 |
| Tr43 | S98 | http://www.functionalglycomics.org/glycomics/CarbProductionServlet?pageType=view&view=view&operationType=view&sideMenu=no&carbProductionId=carbSynthe_0191_D000_S98-1_02222002 |
| Te323 | S310 | http://www.functionalglycomics.org/glycomics/CarbProductionServlet?pageType=view&view=view&operationType=view&sideMenu=no&carbProductionId=carbSynthe_1130_D000_S310-1_09012005 |
| Te224 | S229 | http://www.functionalglycomics.org/glycomics/CarbProductionServlet?pageType=view&view=view&operationType=view&sideMenu=no&carbProductionId=carbSynthe_0517_D000_S229-1_04212004 |
| Te259 | S239 | http://www.functionalglycomics.org/glycomics/CarbProductionServlet?pageType=view&view=view&operationType=view&sideMenu=no&carbProductionId=carbSynthe_480_D000_S239-1_05302004 |
| Te222 | S228 | http://www.functionalglycomics.org/glycomics/CarbProductionServlet?pageType=view&view=view&operationType=view&sideMenu=no&carbProductionId=carbSynthe_493_D000_S245-1_07152004 |
| Te258 | S238 | http://www.functionalglycomics.org/glycomics/CarbProductionServlet?pageType=view&view=view&operationType=view&sideMenu=no&carbProductionId=carbSynthe_481_D000_S238-1_05302004 |
| Te223 | S227 | http://www.functionalglycomics.org/glycomics/CarbProductionServlet?pageType=view&view=view&operationType=view&sideMenu=no&carbProductionId=carbSynthe_0503_D000_S227-2_04212004 |
| Te201 | S201 | http://www.functionalglycomics.org/glycomics/CarbProductionServlet?pageType=view&view=view&operationType=view&sideMenu=no&carbProductionId=carbSynthe_0555_D000_S201-1_01182004 |
| Te101 | S161 | http://www.functionalglycomics.org/glycomics/CarbProductionServlet?pageType=view&view=view&operationType=view&sideMenu=no&carbProductionId=carbSynthe_0255_D000_S161-3_04212004 |
| Te98 | S117 | http://www.functionalglycomics.org/glycomics/CarbProductionServlet?pageType=view&view=view&operationType=view&sideMenu=no&carbProductionId=carbSynthe_0192_D000_S147-1_02052004 |
| Tr260 | S243 | http://www.functionalglycomics.org/glycomics/CarbProductionServlet?pageType=view&view=view&operationType=view&sideMenu=no&carbProductionId=carbSynthe_477_D000_S243-1_05302004 |
| Te221 | S216 | http://www.functionalglycomics.org/glycomics/CarbProductionServlet?pageType=view&view=view&operationType=view&sideMenu=no&carbProductionId=carbSynthe_460_D000_S216-1_02182004 |
| Tr59 | S182 | http://www.functionalglycomics.org/glycomics/CarbProductionServlet?pageType=view&view=view&operationType=view&sideMenu=no&carbProductionId=carbSynthe_469_D000_S182-2_08182003 |
| Te78 | S137 | http://www.functionalglycomics.org/glycomics/CarbProductionServlet?pageType=view&view=view&operationType=view&sideMenu=no&carbProductionId=carbSynthe_0202_D000_S137-2_03032003 |
| Te79 | S112 | http://www.functionalglycomics.org/glycomics/CarbProductionServlet?pageType=view&view=view&operationType=view&sideMenu=no&carbProductionId=carbSynthe_0202_D000_S137-2_03032003 |
| Te75 | S207 | http://www.functionalglycomics.org/glycomics/CarbProductionServlet?pageType=view&view=view&operationType=view&sideMenu=no&carbProductionId=carbSynthe_0549_D000_S207-2_11122004 |
| Tr32 | S57 | http://www.functionalglycomics.org/glycomics/CarbProductionServlet?pageType=view&view=view&operationType=view&sideMenu=no&carbProductionId=carbSynthe_0555_D000_S201-1_01182004 |
| Tr55 | S116 | http://www.functionalglycomics.org/glycomics/CarbProductionServlet?pageType=view&view=view&operationType=view&sideMenu=no&carbProductionId=carbSynthe_0194_D000_S117-1_03142006 |
| Te119 | S139 | http://www.functionalglycomics.org/glycomics/CarbProductionServlet?pageType=view&view=view&operationType=view&sideMenu=no&carbProductionId=carbSynthe_0203_D000_S139-1_11112002 |
| Te97 | S121 | http://www.functionalglycomics.org/glycomics/CarbProductionServlet?pageType=view&view=view&operationType=view&sideMenu=no&carbProductionId=carbSynthe_0203_D000_S139-1_11112002 |
| Tr116 | S123 | http://www.functionalglycomics.org/glycomics/CarbProductionServlet?pageType=view&view=view&operationType=view&sideMenu=no&carbProductionId=carbSynthe_0171_D000_S123-2_10242002 |
| Tr117 | S134 | http://www.functionalglycomics.org/glycomics/CarbProductionServlet?pageType=view&view=view&operationType=view&sideMenu=no&carbProductionId=carbSynthe_0173_D000_S134-3_07072003 |
| Te134 | S147 | http://www.functionalglycomics.org/glycomics/CarbProductionServlet?pageType=view&view=view&operationType=view&sideMenu=no&carbProductionId=carbSynthe_0525_D000_S193-1_11062003 |
| Te135 | S179 | http://www.functionalglycomics.org/glycomics/CarbProductionServlet?pageType=view&view=view&operationType=view&sideMenu=no&carbProductionId=carbSynthe_467_D000_S179-1_11132003 |
| D9 | S23 | http://www.functionalglycomics.org/glycomics/CarbProductionServlet?pageType=view&view=view&operationType=view&sideMenu=no&carbProductionId=carbSynthe_0189_D000_S65-1_05012002 |
| D21 | S162 | http://www.functionalglycomics.org/glycomics/CarbProductionServlet?pageType=view&view=view&operationType=view&sideMenu=no&carbProductionId=carbSynthe_0169_D000_S162-2_05292003 |
| Tr57 | S140 | http://www.functionalglycomics.org/glycomics/CarbProductionServlet?pageType=view&view=view&operationType=view&sideMenu=no&carbProductionId=carbSynthe_0176_D000_S140-1_02052003 |
| D8 | S41 | http://www.functionalglycomics.org/glycomics/CarbProductionServlet?pageType=view&view=view&operationType=view&sideMenu=no&carbProductionId=carbSynthe_0180_D000_S54-1_03212002 |
| Te319 | S301 | http://www.functionalglycomics.org/glycomics/CarbProductionServlet?pageType=view&view=view&operationType=view&sideMenu=no&carbProductionId=carbSynthe_940_D000_S301-1_07142005 |
| Te118 | S178 | http://www.functionalglycomics.org/glycomics/CarbProductionServlet?pageType=view&view=view&operationType=view&sideMenu=no&carbProductionId=carbSynthe_0523_D000_S178-1_07142003 |
| Te212 | S193 | http://www.functionalglycomics.org/glycomics/CarbProductionServlet?pageType=view&view=view&operationType=view&sideMenu=no&carbProductionId=carbSynthe_0525_D000_S193-1_11062003 |
| Te72 | S68 | http://www.functionalglycomics.org/glycomics/CarbProductionServlet?pageType=view&view=view&operationType=view&sideMenu=no&carbProductionId=carbSynthe_0196_D000_S68-1_05012002 |
| Tr54 | S62 | http://www.functionalglycomics.org/glycomics/CarbProductionServlet?pageType=view&view=view&operationType=view&sideMenu=no&carbProductionId=carbSynthe_0196_D000_S68-1_05012002 |
| Te327 | S323 | http://www.functionalglycomics.org/glycomics/CarbProductionServlet?pageType=view&view=view&operationType=view&sideMenu=no&carbProductionId=carbSynthe_1073_D000_S323-1_08132005 |
| Tr62 | S115 | http://www.functionalglycomics.org/glycomics/CarbProductionServlet?pageType=view&view=view&operationType=view&sideMenu=no&carbProductionId=carbSynthe_0175_D000_S115-2_12102002 |
| Te102 | S160 | http://www.functionalglycomics.org/glycomics/CarbProductionServlet?pageType=view&view=view&operationType=view&sideMenu=no&carbProductionId=carbSynthe_0193_D000_S160-4_04082003 |
| Te100 | S113 | http://www.functionalglycomics.org/glycomics/CarbProductionServlet?pageType=view&view=view&operationType=view&sideMenu=no&carbProductionId=carbSynthe_467_D000_S179-1_11132003 |

To access the data concatenate URL as http://ligacloud.ca/searchLibInfo?f=0&b=0&d=20171003-87YGqcEV-SS

| LiGA | Composition details | Naïve composition | Experimental data | Used in |
|---|---|---|---|---|
| YG.xlsx | 56 glycans, with 4 displayed at 2-5 different densities **"LiGA-65"** | 20171003-87YGooOO-SS<br>20171016-87YGooOO-SS | 20171003-87YGqcEV-SS<br>20171003-87YGqdEV-SS<br>20171003-87YGqeEV-SS<br>20171003-87YGqgEV-SS<br>20171016-87YGafEV-SS<br>20171016-87YGbtEV-SS<br>20171016-87YGbtZQ-SS<br>20171016-87YGpgEV-SS<br>20171016-87YGpgZQ-SS<br>20171016-87YGqaEV-SS<br>20171016-87YGqaZQ-SS<br>20171016-87YGqbEV-SS<br>20171016-87YGqbZQ-SS<br>20171016-87YGqcZQ-SS<br>20171016-87YGqdZQ-SS<br>20171016-87YGqeZQ-SS<br>20171016-87YGqfEV-SS<br>20171016-87YGqfZQ-SS<br>20171016-87YGqgZQ-SS<br>20171016-87YGqjEV-SS<br>20171016-87YGqjZQ-SS<br>20171016-87YGqkEV-SS<br>20171016-87YGsaEV-SS<br>20171016-87YGsaZQ-SS | Fig. 3A |
| YU.xlsx | 9 glycans, each one is displayed at 5-6 densities **"LiGA 9x6"** | 20190703-87YUooOO-MS | 20190703-87YUbcBS-MS<br>20190703-87YUsbBS-MS<br>20190703-87YUbgBS-MS<br>20190319-87YUbgBS-MS<br>20190815-87YUqkBS-MS<br>20190815-87YUqkRC-MS<br>20190815-87YUpgBS-MS | Fig. 4B<br>Fig. 4C<br>Fig. 4D |
| YY.xlsx | Same as **"LiGA 9x6"** | 20180130-87YYooOO-JM | 20180222-87YYrfRD-MS<br>20180130-87YYrfRD-RD<br>20180222-87YYrdRD-MS<br>20180130-87YYrdRD-RD<br>20180222-87YYrfRC-MS<br>20180130-87YYrfRC-RD<br>20180222-87YYrdRC-MS<br>20180130-87YYrdRC-RD<br>20180222-87YYrfRB-MS<br>20180130-87YYrfRB-RD<br>20180222-87YYrdRB-MS<br>20180130-87YYrdRB-RD<br>20180222-87YYrfRA-MS<br>20180130-87YYrfRA-RD<br>20180222-87YYrdRA-MS<br>20180130-87YYrdRA-RD | Fig. 5D<br>Fig. S12 |
| YZ.xlsx | 66 glycans with 5 glycans displayed at 2 different densities **"LiGA-71"** | 20190815-87YZooOO-MS | 20190815-87YZsdBS-MS<br>20190815-87YZpgBS-MS<br>20190815-87YZrbRC-MS<br>20190815-87YZrcRC-MS<br>20191210-87YZrfRC-MS<br>20190815-87YZrdRC-MS<br>20191210-87YZvgAW-RR<br>20191210-87YZndAW-RR | Fig. 5A-C<br>Fig. S10<br>Fig. S11 |
| MK.xlsx | 6 glycans, all at 6 densities | 20180130-87MKooOO-MS | 20180130-87MKcaJT-MS | Fig. S9 |
| MJ.xlsx | Same as **"LiGA 9x6"** | 20180130-87MJooOO-MS | 20180130-87MJcaJT-MS<br>20180130-87MJcaKZ-MS | Fig. S9 |
| YT.xlsx | 63 glycans with 9 at 2-5 densities, 5 blank phage clones **"LiGA-75"** | 20181108-87YTooBO-DF | 20181108-87YTbgBS-DF<br>20181108-87YTbgBS-DV<br>20181108-87YTipBS-DV<br>20181108-87YTsbBS-DV | Fig. 2H |
| YO.xlsx | Different from LiGA-75 by 1 glycan | 20180711-87YOooPA-JM | 20180711-87YOrdRB-JM<br>20180711-87YOrfRB-JM | Fig. S11 |
| YX.xlsx | 46 glycans with 3 at 2 densities | 20180222-87YXooPA-JM | 20180222-87YXrdRC-RD<br>20180222-87YXrfRC-RD | Fig. S11 |

**Supplementary Table S3**. List of LiGA mixtures used in this manuscript.
References to the deep-sequencing data, instructions for online access by URL-concatenation, figures in which this data was used and alternative names of LiGA used in the Main Text. Green highlighted names correspond to names used in the Main Text to describe this LiGA.

**1.11** Methods that measure binding of LiGA components

**1.11.1** *Binding of glycosylated clones measured by ELISA.*
Lyophilized Concanavalin A (Sigma-Aldrich, #C2272) was dissolved in PBS at final concentration of 1 mg/mL. The solution was then diluted with PBS to afford final concentration of 20 μg/mL and 50 μL was added to each well of 96 well plate (Corning®, #CLS3369). The plate was covered with sealing tape (Thermo Scientific™, #15036) and incubated overnight at 4 °C. The following day, the wells were washed 3 times by adding washing buffer (200 μL, 0.1% Tween-20 in PBS) in the wells and discarding the solution by inverting the plate on top of a paper towel. Thereafter, blocking solution (100 μL, 20 μg/uL BSA in PBS) was added to the wells and incubated for 1 h at rt. The solution was discarded by inverting the plate, the wells were then washed three times with washing buffer. Thereafter, solutions of LiGA clones (50 μL in PBS) was added to the wells and incubated for 1 h at rt. The solution was discarded by inverting the plate, the wells were washed three times washing buffer (200 μL 0.1% Tween-20 in PBS) and probed with anti-M13-HRP conjugate.

**1.11.2** *Binding of LiGA to ConA or Antibodies immobilized on Plate.*
Plates were prepared as in previous Section **1.11.11**. After the incubation with blocking solution and triple washing 50 μL of either LiGA mixture MJ or MK ($2 \times 10^5$ PFU/mL in PBS) was added to the wells. The solution was incubated for 1 h at rt and discarded by inverting the plate. The wells were washed 2x with washing buffer and 1x with PBS (200 μL). To elute bound phage, 50 μL of HCl (pH 2.0) was added to the well, incubated for 9 min at rt, and the content of each well was transferred to an Eppendorf tube containing 25 μL of 5× Phusion HF buffer (NEB, #M0530S). The neutralized solution was used for titer and as DNA template for PCR and Illumina sequencing.
Binding of LiGA to anti-A and anti-B antibodies was performed in similar fashion.

**1.11.3** *Binding of LiGA to CD22-Fc immobilized on Protein G Magnetic Beads.*
Suspension of protein G coated magnetic beads (Promega #G7472) was vortexed for 2 seconds and 30 μL of suspension was combined with wash buffer (1 mL, 50 mM HEPES, 2 mM $CaCl_2$, pH 7.4 with 0.1% Tween-20 (v/v)). The tube was placed in a Dynabead™ MPC-S (Thermo Fisher, #A13346) for 30 seconds to capture the beads. The supernatant was aspirated and discarded, and the beads were resuspended in 1 mL of wash buffer. Stock solution of CD22-fc protein (1 mg/mL in PBS) was added to the beads suspension to afford 10 μg/mL final concentration. The mixture was incubated overnight at 4 °C with gentle mixing (Labquake®, Rotisserie Shaker). The following day, the beads suspension and other reagents were added to a 96-well Deepwell Plate (Thermo Fisher, #95040450) as follows:

Row A: Suspension of protein-coated beads (1 mL/well)
Row B: reserved for 12-tip Deepwell magnetic comb (Thermo Fisher, #97003500)
Row C: Wash Buffer (1 mL, 0.1 % Tween-20 (v/v) in HEPES buffer)
Row D: Blocking Buffer (1 mL, 2% BSA (w/v) in HEPES Buffer)
Row E: Solution of LiGA YZ (1 mL, $10^8$ PFU/mL in HEPES Buffer)
Row F: Wash Buffer (1 mL, 0.1 % Tween-20 (v/v) in HEPES buffer)
Row G: Wash Buffer (1 mL, 0.1 % Tween-20 (v/v) in HEPES buffer)
Row H: Wash Buffer (1 mL, HEPES buffer)

Following steps were performed using a KingFisher™ Duo Prime Purification System with a magnetic comb to transfer the beads. The program is as follows: a) collect comb from row B, b) collect beads from row A on comb, c) wash beads in row C – 30 s, d) block in row D – 1 h, e) phage binding in row E – 1.5 h, f) wash beads in row F – 1 min, g) wash beads in row G – 1 min, h) wash beads in row H – 1 min. At the end of the program, the protein-coated beads with bound phage were in wells in the row H. The content of each well from row H was transferred to individual Eppendorf™ tube and the tubes were placed in Dynabead™ MPC-S for 30 seconds to capture the beads. The supernatant was discarded, the beads were resuspended in Nuclease free $H_2O$ (30 µL); 2 µL of the suspension solution was sampled and combined with PBS (500 µL, pH 7.4) for titering. The remaining solution was incubated at 90 °C for 15 min, centrifuged at 21,000×g for 10 min, and 25 µL of the supernatant was used as template for PCR reaction as described in PCR Protocol Section.

**1.11.3** *Binding of LiGA to Lectins immobilized on Streptavidin Magnetic Beads.*
The steps in this protocol are identical to "Screening of LiGA YZ to CD22-FC immobilized on Protein G Magnetic Beads" with three exceptions: (i) Streptavidin coated magnetic beads (Promega #Z5481) were used instead of protein G coated magnetic beads. (ii) Biotinylated proteins (ConA, Galectin-3) were added to the beads solution to afford 10 µg/mL final concentration. (iii) LiGA YU (1 mL, $10^7$ PFU/mL in HEPES Buffer) was used.

**1.11.4** *Binding of LiGA YZ to CHO cells expressing CD22.*
Confluent CHO-CD22(+) and CHO-wt cells were detached from culture flask using PBS containing EDTA (5 mM) and centrifuged for 5 min at 300xg. The supernatant was decantated and the pellet was washed twice by resuspending it in PBS (5 mL) and centrifuged for 5 min at 300×g. After final wash the cell pellet was resuspended in incubation buffer (1% BSA in HEPES buffer) at 2 x$10^6$ cells/mL. The cells were aliquoted (500 µL) to FACS tube (Corning, #352054), which afforded 1 million cells per FACS tube. Thereafter LiGA was added to each FACS tube at $10^8$ PFU, which afford approximately $10^6$ PFU of each clone in the incubation solution. The solution was incubated for 1 h at 4 °C. After incubation, the cells were gently vortex (Speed 1, Fisher Vortex Genie 2™, #12-812) and 3 mL of Wash Buffer (0.1% BSA in HEPES buffer) was added to each FACS tube using a squirt bottle (Thermo Fisher ™, #2401-0125). The solution was centrifuged at 218 ×g for 5 min at 4 °C in a swinging bucket rotor. The supernatant was decanted by inverting the FACS tubes and blotting on Kimwipe. Two additional washes were performed: at each wash, the tube was filled with 3 mL of wash buffer, centrifuged and inverted to discard the supernatant in same manner as described above. After the last wash the pellet was resuspended in 1 mL of HEPES buffer, transferred Eppendorf™ tube and centrifuged for 5 min at 218 ×g at 4 °C in a swinging bucket rotor. The supernatant was discarded by pipetting and the pellet was resuspended in 30 µL Nuclease Free $H_2O$. 2 µL of the suspension solution was sampled and combined with PBS (500 µL, pH 7.4) for titering. The remaining solution was incubated at 90 °C for 15 min, centrifuged at 21,000×g for 10 min, and 25 µL of the supernatant was used as template for PCR reaction as described in PCR Protocol Section.

**1.11.5** *Binding of LiGA YZ to Fibroblasts cells expressing DC-SIGN.*
The Rat-6 fibroblast DC-SIGN(+) and DC-SIGN(-) fibroblast cells were detached from culture flask using TrypLE (ThermoFisher, # 12605036) and resuspended in incubation buffer (20 mM HEPES, 150 mM NaCl, 2 mM $CaCl_2$, pH 7.4, 1% BSA,) at 2x$10^6$ cells/mL. The cells were aliquoted (500 µL) to FACS tube (Corning, #352054), which afforded 1 million cells per FACS tube. Binding of LiGA to these cells was performed using the steps identical to the aforementioned protocol ("*Binding of LiGA YZ to CHO cells expressing CD22*").

**1.11.6** *Binding of LiGA YY to Fibroblasts cells expressing DC-SIGN.*
This protocol is analogous to "*Binding of LiGA YZ to Fibroblasts cells expressing DC-SIGN*" with two exceptions: (i) LiGA YY was used in place of YZ; (ii) Number of washed were varied from 1 to 4 and sequencing data from each wash optimization are available in Fig. S11.

**1.12** *Origin and maintenance of rat fibroblast cells expressing DC-SIGN*
The Rat-6 fibroblast DC(+) and DC(−) cell lines[11] were received as a gift Professor Kurt Drickamer at Department of Life Sciences, Imperial College London. Cell were maintained in DMEM-F12 medium (Thermo Fisher, #11330057) supplemented with 10% Fetal Bovine Serum (Thermo Fisher, #26140079) and Penicillin-Streptomycin (Thermo Fisher, #15140122).

**1.13** *Production and maintenance of CHO cells expressing CD22*
Full length human CD22, in the pcDNA5/FRT vector, was stably transfected into Chinese Hamster Ovary (CHO) cell line through Flp-In™ (Thermo Fisher, #K601001) system under selection with 0.5 mg/mL hygromycin-B (Thermo Fisher) for two weeks. The cells were passaged using 1 mM EDTA/PBS (Thermo Fisher #13151014) as a dissociation solution and maintained in DMEM-F12 medium (Thermo Fisher, # 11330057) supplemented with 10% Fetal Bovine Serum (Thermo Fisher, #26140079) and Penicillin-Streptomycin (Thermo Fisher, #15140122). Untransfected CHO cells were maintained in the same conditions and used as control.

**1.14** *PCR Protocol Section*
The 25 μL of DNA template solution in Nuclease free water was amplified in total volume of 50 μL with 1x Phusion® buffer, 50 μM of each dNTPs, 500 μM $MgCl_2$, 1 μM forward barcoded primer, 1 μM reverse barcoded primer and one unit of Phusion® High-Fidelity DNA Polymerase (NEB, #M0530S). A typical 50 μL reaction mixture contained:

| | |
|---|---|
| 1. 5x Phusion buffer | 10 μL |
| 2. 10 mM dNTPs | 1 μL |
| 3. Phusion® Polymeras | 0.5 μL |
| 4. Forward primer (10 μM) | 2.5 μL |
| 4. Reverse primer (10 μM) | 2.5 μL |
| 6. Template solution | 25 μL |
| 7. Nuclease free water | 8.5 μL |

Exceptions: in amplification of clonal phage and naïve libraries, volume of template (phage solution) was 2 μL. In panning against the intact cells, protein coated beads, and protein coated wells the volume of solution containing the template was 25 μL.

Cycling was performed using the following thermocycler settings:
a) 98°C 3 min,
b) 98°C 10 s,
c) 50 °C 20 s,
d) 72 °C 30 s,
e) repeat b)-d) for 10 cycles,
f) 98 °C 10 s,
g) 72 °C 30s,
h) repeat f)-g) for 20 cycles,

h) 72 °C 5 min, i) 4 °C hold

**1.15** *Illumina sequencing.*
The PCR products described in "1.14 PCR Section" were quantified by 2% (w/v) agarose gel in Tris-Borate-EDTA buffer at 100 volts for ~35 min using a low molecular weight DNA ladder as standard (NEB, #N3233S). PCR products that contain different indexing barcodes were pooled allowing 10 ng of each product in the mixture. The mixture was purified by eGel, quantified by quBit and sequenced using the Illumina NextSeq paired-end 500/550 High Output Kit v2.5 (2x75 Cycles). Data was automatically uploaded to BaseSpace™ Sequence Hub. Processing of the data is described in section "2.2 Processing of Illumina data".

**1.16** *Generation of Monoclonal anti-Galf$_4$ antibody*

**1.16.1** *Conjugation of Galf$_4$ to tetanus toxoid.*
Tetanus toxoid (TT) purchased from Statens Serum Institute (15 mL in saline) was exchanged to PBS buffer, pH 7.4 using an Amicon Ultra-15 Centrifugal Filter Unit (NMWL 10,000). Propargyl-N-hydroxy succinimidyl ester (4.0 mg) was added to the protein solution (6 mL) and the reaction was stirred at room temperature overnight. The protein was exchanged with fresh PBS buffer a few times and further purified by passing through a gel filtration column (Sephadex G-25, GE HealthCare). The propargylated tetanus toxoid was concentrated and its concentration was estimated using Micro BCA Protein Assay (Thermo Fisher Scientific). The number of amino group modifications on the protein was estimated by using both MALDI-TOF MS and TNBS amino group estimation assay (Sigma Aldrich). Next, propargylated tetanus toxoid (35 mg) and Gal*f*$_4$ (6 mg) were dissolved in 0.2 M Tris buffer, pH 8.0 (6 mL) with and copper powder (~1 mg) was added. The glass vial was sealed with open-top screw cap with a rubber septum and the vial was alternatively degassed and purged with argon a few times. The bathophenanthroline / $Cu^+$ catalyst (0.1 mL) was added to initiate the conjugation. After overnight incubation at room temperature, the reaction was quenched with 0.5 M EDTA, pH 8.0 and exchanged with fresh PBS buffer few times using an Amicon Ultra-15 Centrifugal Filter Unit (NMWL 10,000). The conjugate was then purified on a gel filtration column (Superdex S-200, GE HealthCare) in PBS. The TT conjugate was concentrated to 6 mL and as before its protein concentration was estimated using Micro BCA Protein Assay (Thermo Fisher). The degree of Gal*f*$_4$ units incorporated on the TT was 14/protein as determined by MALDI-TOF MS. The final conjugates were sterile filtered prior to injection.

**1.16.2** *Animals for generation of anti-Galf$_4$ antibody.*
Female balb/c mice (5 mice, Charles River, Canada, 6–8 weeks old) were used to raise the monoclonals. All the procedures and experiments involving animals were carried out using a protocol approved by Animal Care Committee: Biosciences, University of Alberta. The protocol was approved as per Canadian Council on Animal Care (CCAC) guidelines.

**1.16.3** *Immunizations.*
Animals were immunised three times (10.0 µg of TT-Gal*f*$_4$/mouse/injection) at intervals of 21 days. A total volume of 250 µL was injected on each mouse of which 150 µL was injected inter-peritoneally and the remaining 100 µL was injected subcutaneously. The conjugate was adsorbed on freshly prepared alum[12] on the night prior to injection. Pre bleeds were collected before immunisation and first test bleed was collected 10 days after the second dose. The animals were euthanized 10 days after the final injection and final bleed was collected.

**1.16.4** *Serum processing*.
After collection, blood was incubated at 37° C for 1 h and then spun at 1500 g for 10 min. Clear serum from the top of the tube was collected and stored at –20 °C until use.

**1.16.5** *Immunoassays*.
Antibody levels in the murine sera were studied using enzyme linked immuno-sorbent assay (ELISA). A published protocol[13] was followed with slight modifications. Polystyrene microtiter plates were incubated with BSA-Gal$f_4$ (1 µg/mL, 100 µL/well) at 4 °C overnight, then washed (5×) with PBST (0.05% Tween-20 in phosphate buffer saline, PBS). Then murine sera were added to the coated well in serial 10-fold dilutions (100 µL/well). The starting dilution was 1:100. After incubation at room temperature for 2 h, the plates were washed (5×) with PBST. Then the plate was incubated with 100 µL/well of 1: 5000 diluted goat anti-mouse IgG antibody, tagged with HRPO (KPL, 1.0 mg/mL stock) for 30 min at room temperature, then washed (5×) with PBST. A peroxidase substrate, 3,3',5,5'-Tetramethylbenzidine (TMB) with $H_2O_2$, was added. After 15 min the reaction was quenched by addition of phosphoric acid (1M, 100 µL/well). The plates were read at 450 nm and the data were processed using Excel and Origin softwares. The buffer, 0.1% BSA in PBST was used to dilute all sera. End point dilution ($x_0$) was recorded as the serum dilution giving an absorbance 0.20 above background and antibody titer was calculated as reciprocal of $x_0$.

**1.16.6** *Hybridoma fusion and cloning*.
The fusions were performed by following earlier published protocols[14] with little modification. Mouse myeloma cell line SP2/0 was fused with the splenocytes collected from the immunised mice, in the 1:5 ratio in the presence of 35% polyethylene glycol (PEG). Total time of exposure of cells to PEG was 8 min, which included addition of PEG to cells, incubation with swirling and finally spinning down of the cell pellet to remove the PEG. All the cell culture media, hybridoma selection media, growing conditions, plating and maintaining of the fused cells were performed as per the published protocol[14]. At days 10 post-fusion, the plates were scanned visually for any sign of cloud formation at the bottom of the well and also for any color change of media. Pre-screened wells were then observed under microscope to check for clone formation. The supernatant from the wells, with clone, were tested by ELISA using BSA-Gal$f_4$ as coating agent. The two positive clones were transferred to 12-well plate for expansion and later cloned twice using limiting dilution method of 1 cell/well. After each cloning the supernatants were screened as before.

**1.16.7** *Isotyping and specificity testing of the hybridomas*.
The culture supernatants of two final clones were isotyped using mouse monoclonal antibody isotyping kit (IsoStrip, Roche, supplied by Sigma Aldrich, Canada) by following the vendor's protocol. Both the clones were found to be IgG1 type.

**1.17** *Panning of LiGA in mice*.
All the procedures and experiments involving animals were carried out using a protocol approved by the Health Sciences Laboratory Animal Services (HSLAS), University of Alberta. The protocol was approved as per the Canadian Council on Animal Care (CCAC) guidelines. All mice were maintained in pathogen-free conditions at the University of Alberta breeding facility. To screen for sialoside ligands of hCD22, we created a mixed bone marrow chimera mouse by lethally irradiating the recipient mice and transplanting them with a 50:50 mixture of bone marrow collected from the femurs of mCD22KO (mCD22$^{-/-}$ hCD22$^{-}$ CD45.2$^{+/+}$) and hCD22 transgenic (mCD22$^{-/-}$ hCD22$^{+}$ CD45.2$^{+/+}$) mice. Four weeks after transplantation, mice were bled

to verify that the desired chimera had been established. Six weeks after irradiation, animals were injected with LiGA (0.2 m, 1 x $10^{11}$ PFU/mL in PBS). One hour post-injection mice were euthanized with $CO_2$. Internal organs (heart, liver, kidney, lung and spleen) were collected and stored in cold DMEM (Thermo Fisher).
Tissues were homogenized by grinding between 75 mm frosted microscope slides (Thermo Fisher). Homogenized tissues was transferred into 5 mL of RBC lysis buffer (155 mM ammonium chloride, 10 mM, EDTA, 100 mM Tris, pH 7.4), incubated for 7 min, centrifuged at 300 rcf for 5 min, and the resulting pellet was resuspended in 0.4 mL PBS + 0.1% BSA. A 2 mL sample from cell suspension was used to determine the amount of phage particles in specific organ by plaque forming (PFU) assay.
To sort the B cells, the spleen was homogenized and RBCs was lysed as described above. RBC-depleted splenocytes were stained with a cocktail of fluorescent antibodies: anti-mouse/human CD45R/B220-APC-Cy7 (1 µg/mL, Biolegend), Brilliant anti-mouse CD19-BV605 (1 µg/mL, Biolegend), anti-mouse CD22-AF647 (2 µg/mL, Biolegend), anti-human CD22-BV421 (2 µg/mL, Biolegend), anti-mouse CD45.1- PE/Cy7 (1 µg/mL, Biolegend), and anti-mouse CD45.2-BV421 (1 µg/mL, Biolegend). Dead cells were excluded with propidium iodide (1 µg/mL, Thermo Fisher). The stained cells were sorted on FACS Aria cell sorter. Viable singlet B cells ($CD19^{+}B220^{+}$) were divided into $CD45.1^{+/+}$ ($mCD22^{-/-}hCD22^{-}$) and $CD45.2^{+/+}$ ($mCD22^{-/-}hCD22^{+}$). Both populations were verified to be lack mCD22 expression and the $CD45.2^{+/+}$ cells were verified to express hCD22. One million cells from each population were collected and these cells were analyzed using PFU-assay.

### 2. Data Processing Methods

**2.1** *General data processing methods*.
Data analysis was performed in Python, Matlab or R/Bioconductor. Core scripts are available as part of the supporting information or on GitHub. Other scripts are available upon request.
Data storage cloud http://ligacloud.ca/ was implemented by 48Hour Discovery Inc (Edmonton, Canada) as Amazon Web Server (AWS)-backed cloud in Linux-Apache-MySQL-Python (LAMP) architecture. Details of the implementation are beyond the scope of this report; details will be disclosed in our later reports.

For statistical analysis of titering assays described in main text Fig. 2, Fig. 6, and Supplementary Figure 8, we employed two tailed Student's t-tests. Curve fitting of ELISA (Supplementary Figure 7) was performed using R (Make_fig_SI7b.r and Make_fig_SI7c.r).

Comparison and testing differences for significance in LiGA data was performed essentially as differential enrichment analyses of phage displayed library sequencing described in our previous reports[12-13] and based on differential expression (DE) analysis implemented in edgeR[14-15]. In DE-analysis three factors were considered: (i) modeling of the observed counts using a negative binomial model; (ii) Benjamini–Hochberg (BH) adjustment to control the false discovery rate (FDR) at $\alpha = 0.05$[16]; (iii) normalization of data across multiple samples using the Trimmed Mean of M-values (TMM) normalization[17].

To assess the significance of a glycan binding in a specific experiment the differential enrichment of the levels of the DNA barcode associated with that glycan in "test" sets of DNA read was compared to the levels of the same read in "control" sets. For example, in cell-based experiments, the "test" dataset was association of LiGA with receptor (+) cell line, whereas the control was the dataset was association of the identical LiGA with isogenic cell line that contained no target receptor. In binding to lectins, the "control" dataset was association of LiGA with blank carriers (beads or plate). Prior to DE-analysis, "test" and "control" data sets were retrieved from the http://ligacloud.ca/ server as tables of glycans, DNA, and raw sequencing counts. DNA reads that could not be mapped to any entries in LiGA dictionary were discarded.

**2.2** *Processing of Illumina data.*

The Gzip compressed FASTQ files were downloaded from BaseSpace™ Sequence Hub. The files were converted into tables of DNA sequences and their counts per experiment, essentially as previously described.[18-20] Briefly, FASTQ files were parsed into separate files based on unique multiplexing barcodes within the reads. Reads that did not contain an identifiable multiplex barcode were discarded. Reads that contained a Phred=0 quality score in any position were also discarded. No other filtering by Phred score was performed because we previously have shown[12, 21] that filtering deep-sequencing output of phage display libraries of short peptides by Phred score does not increase the quality of the data.[12] Instead, quality control was based on:

(i) mapping the forward and reverse barcoding regions allowing no more than one base substitution each.
(ii) alignment of the forward and reverse read-end overlap, allowing no mismatches between forward and reverse read in the overlap region
Reads that did not match criteria (i) and (ii) were discarded. The two endds of read-pairs that pass the filtering criteria were joined and trimmed to the DNA sequences located between the priming regions; the reads were organized in a tab-delimited text file containing the unique DNA sequences and their copy numbers. Technical replicates were often combined in the same file.

Using an SDB-lookup table, we mapped DNA sequences to SDB. Any reads that were one substitution or less (H≤1) away from the expected SDB sequence were assigned to the parent SDB sequence. Once SDB were mapped, a LiGA-specific lookup table ("LiGA dictionary") was used to convert the identified SDB to glycans and display density. Abbreviated names of glycans are based on those used on CFG website:
http://www.functionalglycomics.org/static/consortium/resources/resourcecored2.shtml

Translated files with raw DNA reads, raw counts, and mapped glycans were uploaded to http://ligacloud.ca/ server.

**2.3** *Maintenance of parsed sequencing data.*

All LiGA sequencing data is publicly available at http://ligacloud.ca/ server. Each experiment has a unique alphanumeric name (e.g., **20180711-87YOrdRB-JM**) and unique static URL (Highlighted portion of the URL is constant for all files):
http://ligacloud.ca/searchLibInfo?f=0&b=0&d=20180711-87YOrdRB-JM

**Screenshot** below described the data page of **20180711-87YOrdRB-JM** dataset:

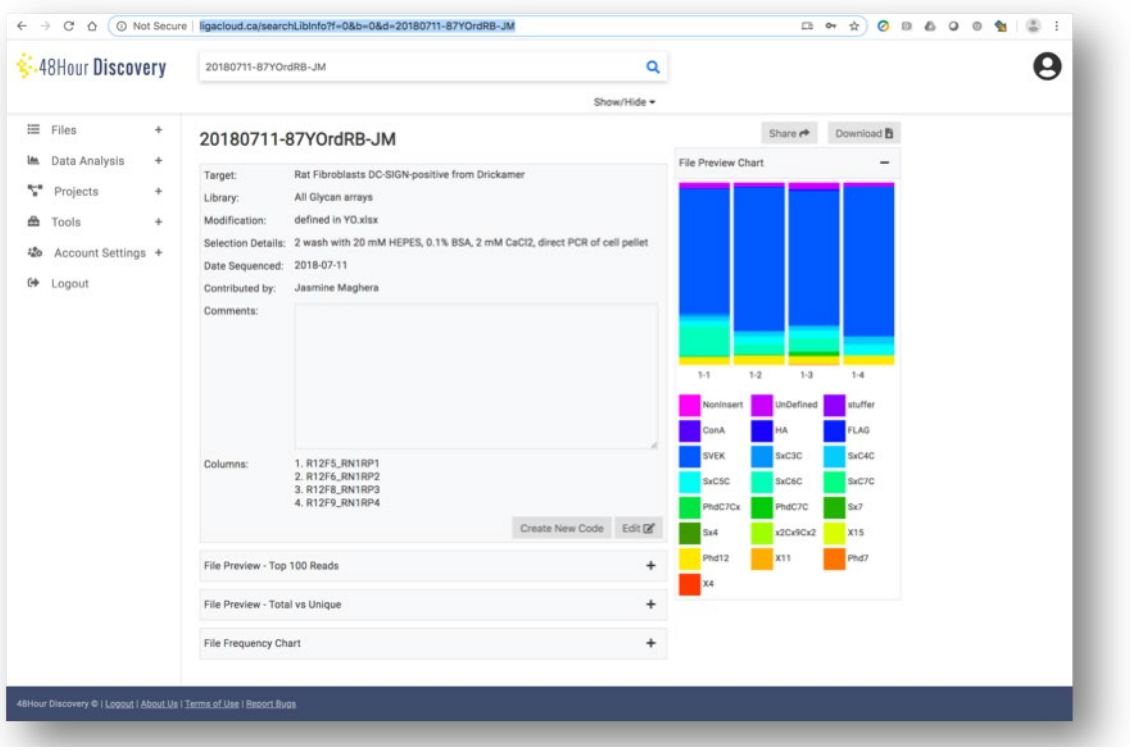

The data can be viewed using "File Preview - Top 100 Reads" option. The names of data files relevant to this manuscript are listed in **Supplementary Table S3**. Data files can also be searched for by any metadata associated with the experiment (e.g., target name, experimental conditions).

**2.4** *Access to CFG data*.

The script *download_CFG.py* downloads the publicly available CFG data for a particular lectin. It enters the given term in the search tool at http://www.functionalglycomics.org . The results for the search are then used to extract PSIDs or 'prime screen ID' used by CFG website used to keep track of the data. The PSIDs can be used to download the available data on the given lectin. Automated approach conveniently downloaded any data that can be found through the manual "search page" at: http://www.functionalglycomics.org/glycomics/publicdata/primaryscreen.jsp It also downloaded calibration data that was not directly available through the "Search" page.

Lectin binding data originating from different arrays was aligned by *align_CFG.py* script. It combines three data storing formats used in CFG array version 3.0-5.2 and converts the glycan names of all the old formats to IUPAC condensed using *IUPAC_converter.py*. The three glycan lists were merged to form a fixed order for all three formats found in *order.xlsx*. Each CFG data file is then parsed, its format is identified, glycan array data is extracted and appended to this order. Heat maps for the combined data were generated using MatLab.

Synchronization of the nomenclature of glycans and linkers was critical for comparing LiGA data with CFG glass array data. To aid this comparison, LiGA dictionary contained multiple alternative spelling: condensed and short-form IUPAC notation, GlyTouCan notation, common name and CGF compound number.

**2.5** *Automated drawing of glycan symbol nomenclature from glycan names*.

For main text and supporting figures other than MALDI analysis figures, glycan drawings were made by *draw_glycans.py* which uses a previously reported DrawGlycan-SNFG tool[15]. In many

cases, the resulting vector images were manually post-processed in Adobe Illustrator to detail the linkage information.
One exception: in automated MALDI processing, we wrote a drawGlycan.m MatLab script to automatically calculate the molecular weights of glycans and generated symbolic CFG representation of glycans, linkers from glycan names in a short-form IUPAC notation used by CGF at:

Glycans: http://www.functionalglycomics.org/static/consortium/resources/resourcecored2.shtml
Linkers: http://www.functionalglycomics.org/static/consortium/resources/resourcecoreh8.shtml

The automated analysis by drawGlycan.m converted short-form UIPAC notation to GlycanLeaf object, which contained all information about monomers, linkages and linkers. Summary of identities, abbreviations, CFG representation, and molecular weights of glycan monomers and linkers is available as part of @GlycanLeaf/monomers.xlsx.

We implemented class GlycanLeaf as MatLab data structure specialized for non-cyclic, non-modified oligosaccharides primarily of mammalian origin. It is a simpler alternative to existing tools such as GlyContainer[16] that aim to represent all classes of monomers, modification, and ambiguous linkage information. Specialization of GlycanLeaf class made it possible to incorporate a simple autocorrect for common errors observed in IUPAC notation in public repositories. Currently GlycanLeaf can handle names with wrong shapes of brackets, wrong case, trailing or internal blanks, inconsistent use of anomeric linkages, inconsistent spelling of building blocks such as Neu5Ac vs. NeuAc, the use of repeat units, the use of sub-units such as LN, etc. Such autocorrect would be significantly more challenging to implement for data structures that capture all possible glycans of all possible origins.

**2.6** *Semi-automated processing of MALDI data*.
Critical part of LiGA and LiGA-dictionary was the copy number of the each glycan in the mixture; these numbers were determined prospectively for each clone by MALDI. We implemented an automated pipeline for processing of raw MALDI *.txt files to images and integration data. This task was performed by plotMALDI.m MatLab script, available in SI/MatLab folder.
The script calculated the anticipated MW of species anticipated to be observed in MALDI: (i) pVIII protein; (ii) DBCO-pVIII; (iii) glycosylated pVIII; (iv) glycosylated pVIII with lost sialic acid (for sialylated glycans only). It fit the peaks in the areas around those masses to equation `'a1*exp(-((x-b1)/c1)^2)+e1*x+f1` (Gaussian function with linear baseline correction). The fit parameter `a1` was used as a simple estimate of the peak height. The relative heights of peaks that correspond to pVIII and glycosylated pVIII were used as an estimate fraction of glycosylated pVIII proteins. The absolute copy number of glycosylated pVIII proteins was calculated based on the assumption that the phage M13KE contains 2700 copies of pVIII. We avoided more complex algorithms for integration and baseline correction due to to relatively high noise of MALDI data and high uncertainty of ionization of glycosylated vs. non-glycosylated pVIII. In other words, we believe that height serves as a sufficiently accurate estimate of relative copy numbers; higher accuracy of estimation of peak area in MALDI is unlikely to yield a higher accuracy estimate of the true copy number of the glycans on phage.

### 3. Synthetic Methods

**3.1** *General Synthetic Methods.*
All reagents were purchased from commercial sources and used without further purification. Oven-dried glassware was used for all reactions. Reaction solvents were dried by passage through columns of alumina and copper under nitrogen. All reactions, unless stated otherwise, were carried out at room temperature under positive pressure of argon. Organic solutions were concentrated under vacuum below 40 °C. Reaction progress was monitored by TLC on Silica Gel 60 $F_{254}$ (0.25 mm, E. Merck). Visualization of the TLC spots was done either under UV light or charring the TLC plates with acidified *p*-anisaldehyde solution in ethanol or phosphomolybdic acid stain. Column chromatography was performed using Silica Gel 40–60 μM. Optical rotations $[\alpha]_D$ measurement were carried out at 22 °C and reported in deg·cm$^2$ dm$^{-1}$·g$^{-1}$. $^1$H NMR spectra were recorded using 500 MHz or 400 MHz instruments, and the data are reported as if they were first order. $^{13}$C NMR (APT) spectra were recorded at 125 MHz. Assignments of data were made using $^1$H–$^1$H COSY and HMQC experiments. In oligosaccharides, proton and carbon assignments of the non-reducing end residue (that attached to the 8-azido-octyl group) are reported without any prime and others are assigned as X′, X′′and X′′′ for each residue from the non-reducing to reducing end. Electrospray mass spectra were recorded on samples suspended in THF/MeOH mixture with added NaCl.

**3.2** *General procedure for Zémplen deacylations.*
The acylated compound was dissolved in 3:1 MeOH–$CH_2Cl_2$ (20–50 mL). To that solution NaOMe in MeOH (0.1 M) was added dropwise until the solution reached pH 12 as determined by wet pH paper. The reaction mixture was stirred from 4–8 h at room temperature with monitoring by TLC. After completion, the solution was neutralized by the addition of prewashed Amberlite-15 ($H^+$) cation exchange resin. After filtration and concentration of the solution, the crude residue was purified by chromatography.

**3.3** *General procedure for removal of levulinoyl protecting groups.*
The levulinoyl protected sugar (1 equivalent) was dissolved in 3:1 $CH_2Cl_2$–MeOH (10 mL) and hydrazine acetate (2 equivalents) was added. The solution was stirred for 2–4 h and the reaction was monitored by TLC. When the reaction was complete, the solvent was concentrated under vacuum, and the residue was diluted with EtOAc. The organic solution was then washed with a saturated solution of $NaHCO_3$ and brine, and then dried over anhydrous $Na_2SO_4$ filtered and concentrated. The crude residue was purified by chromatography.

**Scheme S1:** Synthesis of **Gal*f*1, S5** and **S6**

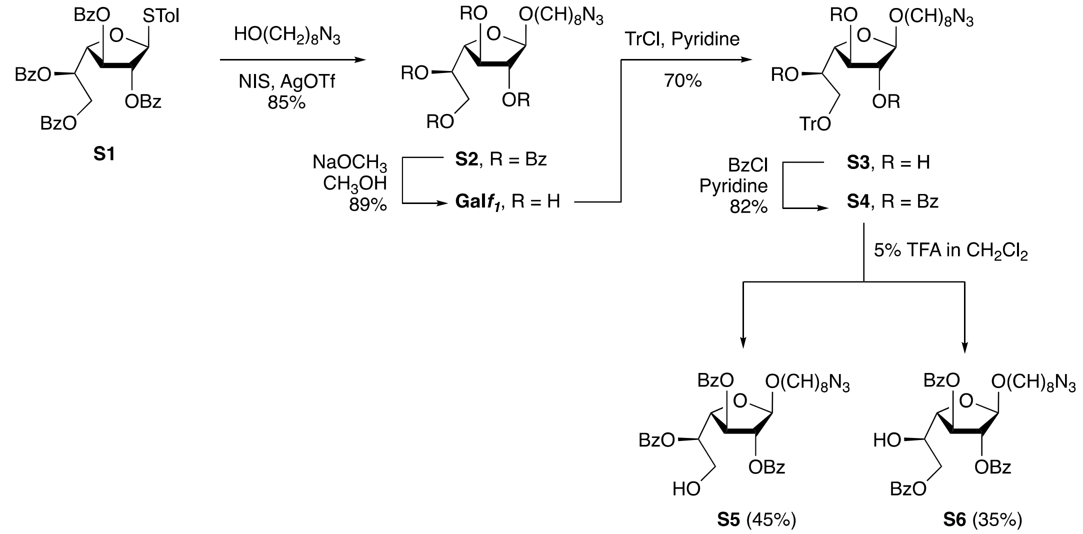

**3.4** *Procedures for synthesis of* ***Galf1, Galf2, Galf3, Galf4.***
The levulinoyl protected sugar (1 equivalent) was dissolved in 3:1 $CH_2Cl_2$–MeOH (10 mL) and hydrazine acetate (2 equivalents) was added. The solution was stirred for 2–4 h and the reaction was monitored by TLC. When the reaction was complete, the solvent was concentrated under vacuum, and the residue was diluted with EtOAc. The organic solution was then washed with a saturated solution of $NaHCO_3$ and brine, and then dried over anhydrous $Na_2SO_4$ filtered and concentrated. The crude residue was purified by chromatography.

**8-Azido-octyl 2,3,5,6-tetra-*O*-benzoyl-β-D-galactofuranoside (S2).** Thioglycoside **S1**[17] (2.0 g, 2.84 mmol) and 8-azido-octanol[18] (320 mg, 1.89 mmol) were dissolved in dry $CH_2Cl_2$ (20 mL) in the presence of 300 mg activated, powdered 4 Å molecular sieves and the mixture was cooled to 0 °C. The solution was stirred for 15 min followed by the addition of *N*-iodosuccinimide (NIS, 635 mg, 2.84 mmol) and AgOTf (120 mg, 0.47 mmol). The reaction mixture was stirred for another 30 min and then $Et_3N$ was added before being diluted with $CH_2Cl_2$ and filtered through a Celite pad. The filtrate was washed with a saturated $Na_2S_2O_3$ solution followed by water and brine, and then dried over anhydrous $Na_2SO_4$ and filtered. The filtrate was concentrated and the residue was purified by chromatography (4:1 hexane–EtOAc) to give **S2** (1.13 g, 85%) as a colorless syrup. $R_f$ 0.32 (4:1 hexanes–EtOAc); $[\alpha]_D$ –119.7 (*c* 0.4, $CH_2Cl_2$); $^1H$ NMR (500 MHz, $CDCl_3$, $\delta_H$): 8.20–7.85 (m, 6 H, Ar), 7.63–7.35 (m, 14 H, Ar), 6.14 (app dt, 1 H, $J$ = 7.8, 3.4 Hz, H-5), 5.66 (d, 1 H, $J$ = 5.2 Hz, H-3), 5.49 (d, 1 H, $J$ = 1.2 Hz, H-2), 5.32 (s, 1 H, H-1), 4.84–4.72 (m, 2 H, H-6a,b), 4.66 (dd, 1 H, $J$ = 5.3, 3.5 Hz, H-4), 3.78 (td, 1 H, $J$ = 9.6, 6.7 Hz, octyl O$CH_2$), 3.56 (td, 1 H, $J$ = 9.5, 6.3 Hz, octyl O$CH_2$), 3.25 (t, 2 H, $J$ = 6.9 Hz, octyl $CH_2N_3$), 1.64–1.60 (m, 4 H, octyl $CH_2$), 1.44–1.29 (m, 8 H, octyl $CH_2$); $^{13}C$ NMR (125 MHz, $CDCl_3$, $\delta_C$): 166.1 (C=O), 165.7 (C=O), 165.6 (C=O), 165.5 (C=O), 133.4 (Ar), 133.3 (Ar), 133.2 (Ar), 133.1 (Ar), 129.9 (Ar), 129.9 (Ar), 129.8 (Ar), 129.7 (Ar), 129.6 (Ar), 129.5 (Ar), 129.1(Ar), 129.0 (Ar), 128.4 (Ar), 128.4 (Ar), 128.4 (Ar), 105.6 (C-1), 82.1 (C-2), 81.2 (C-4), 76.8 (C-3), 70.3(C-5), 67.6 (octyl O$CH_2$), 63.6 (C-6), 51.5 (octyl $CH_2N_3$), 29.5 (octyl $CH_2$), 29.3 (octyl $CH_2$), 29.1 (octyl $CH_2$), 28.8 (octyl $CH_2$), 26.7 (octyl $CH_2$), 26.1 (octyl $CH_2$). HR ESIMS: *m/z* $[M+Na^+]$ calcd for $C_{42}H_{43}N_3NaO_{10}$: 772.2841. Found: 772.2833.

**8-Azido-octyl β-D-galactofuranoside (Gal*f*1).** Compound **S2** (1.1 g, 1.5 mmol) was debenzoylated according to the general deacylation procedure. Removal of the solvent afforded an oily residue that was purified by chromatography (9:1 $CH_2Cl_2$–MeOH) to give **Gal*f*1** (444 mg, 89%) as a colorless oil. $R_f$ 0.4 (5:1 $CH_2Cl_2$–MeOH); $[\alpha]_D$ –49.5 (*c* 0.4, $CH_2Cl_2$); $^1H$ NMR (500 MHz, $CDCl_3$, $\delta_H$): 4.98 (s, 1 H, H-1), 4.13–3.97 (m, 4 H, H-2, H-3, H-4, H-5), 3.82–3.64 (m, 3 H, H-6a,b, octyl C$H_2$), 3.44 (dt, 1 H, $J$ = 9.6, 6.7 Hz, octyl C$H_2$), 3.28 (t, $J$ = 6.9 Hz, 2 H, octyl C$H_2N_3$), 1.62–1.58 (m, 4 H, octyl C$H_2$), 1.36–1.21 (m, 8 H, octyl C$H_2$); $^{13}C$ NMR (125 MHz, $CDCl_3$, $\delta_C$): 107.9 (C-1), 85.6 (C-4), 79.9 (C-3), 78.2 (C-5), 71.1 (C-2), 68.0 (octyl O$CH_2$), 64.2 (C-6), 51.5 (octyl $CH_2N_3$), 29.5 (octyl $CH_2$), 29.3 (octyl $CH_2$), 29.1 (octyl $CH_2$), 28.8 (octyl $CH_2$), 26.7 (octyl $CH_2$), 25.9 (octyl $CH_2$); HR ESIMS: *m/z* [M+Na$^+$] calcd for $C_{14}H_{27}N_3NaO_6$: 356.1792. Found: 356.1790.

**8-Azido-octyl 6-*O*-trityl-β-D-galactofuranoside (S3).** To a solution of **Gal*f*1** (1.2 g, 3.6 mmol) in pyridine (35 mL), trityl chloride (1.5 g, 5.31 mmol) was added. The reaction mixture was stirred at 50 ºC for 48 h. After cooling to room temperature, the solvent was evaporated; co-evaporation with toluene removed pyridine. The residue was then purified by chromatography (19:1 $CH_2Cl_2$–MeOH) to give **S3** (1.44 g, 70%) as a colorless oil. $R_f$ 0.49 (19:1 $CH_2Cl_2$–MeOH); $[\alpha]_D$ –63.6 (*c* 0.5, $CH_2Cl_2$); $^1H$ NMR (500 MHz, $CDCl_3$, $\delta_H$) 7.49–7.42 (m, 5 H, Ar), 7.37–7.25 (m, 10 H, Ar), 4.96 (s, 1 H, H-1), 4.10–4.01 (m, 2 H, H-2, H-3), 3.97–3.91 (m, 2 H, H-4, H-5), 3.68 (dt, 1 H, $J$ = 9.6, 6.7 Hz, octyl OC$H_2$), 3.45–3.33 (m, 3 H, octyl $CH_2$, H-6a,b), 3.25 (t, 2 H, $J$ = 7.0 Hz, octyl C$H_2N_3$), 1.57–1.52 (m, 5 H, octyl C$H_2$), 1.39–1.27 (m, 7 H, octyl C$H_2$); $^{13}C$ NMR (125 MHz, $CDCl_3$, $\delta_C$) 143.6 (Ar), 128.6 (Ar), 127.9 (Ar), 127.3 (Ar), 108.3 (C-1), 87.2 ($Ph_3C$) 86.9 (C-4), 79.1 (C-2), 78.3 (C-3), 70.8 (C-5), 67.6 (C-6), 64.7 (octyl O$CH_2$), 51.5 (octyl $CH_2N_3$), 29.4 (octyl $CH_2$), 29.2 (octyl $CH_2$), 29.0 (octyl $CH_2$), 28.8 (octyl $CH_2$), 26.6 (octyl $CH_2$), 26.0 (octyl $CH_2$); HR ESIMS: *m/z* [M+Na$^+$] calcd for $C_{33}H_{41}N_3NaO_6$: 598.2888. Found: 598.2882.

**8-Azido-octyl 2,3,5-tri-*O*-benzoyl-6-*O*-trityl-β-D-galactofuranoside (S4).** A solution of **S3** (1.2 g, 2.08 mmol) in pyridine (50 mL) was cooled to 0 ºC. Benzoyl chloride (1.68 g, 12 mmol) was then added slowly. The reaction mixture was stirred overnight and was then poured into an ice water mixture and extracted with $CH_2Cl_2$. The organic layer was washed with brine, dried over anhydrous $Na_2SO_4$, filtered and concentrated to give a crude residue that was purified by chromatography (3:1 Hexane–EtOAc) to give **S4** (1.45g, 82%) as a white solid, $R_f$ 0.39 (3:1 Hexane–EtOAc); $[\alpha]_D$ –51.2 (*c* 0.5 $CH_2Cl_2$); $^1H$ NMR (500 MHz, $CDCl_3$, $\delta_H$) 8.22–8.10 (m, 2 H, Ar), 8.08 (d, 2 H, $J$ = 8.2 Hz, Ar), 7.90 (d, 2 H, $J$ = 8.2 Hz, Ar), 7.67–7.18 (m, 24 H, Ar), 5.89–5.77 (app dt, 1 H, $J$ = 10.0, 5.1 Hz, H-5), 5.46 (d, 1 H, $J$ = 4.8 Hz, H-3), 5.38 (s, 1 H, H-1), 5.27 (d, 1 H, $J$ = 4.9 Hz, H-2), 4.74 (dd, 1 H, $J$ = 5.3, 4.8 Hz, H-4), 3.82–3.71 (m, 2 H, H-6a,b), 3.62 (dd, 1 H, $J$ = 9.7, 6.0 Hz, octyl OC$H_2$), 3.53 (ddt, 1 H, $J$ = 18.5, 9.8, 5.6 Hz, octyl OC$H_2$), 3.23 (t, 3 H, $J$ = 7.0 Hz, octyl C$H_2N_3$), 1.71–1.54 (m, 4 H, octyl C$H_2$), 1.37–1.28 (m, 8 H, octyl C$H_2$); $^{13}C$ NMR (125 MHz, $CDCl_3$, $\delta_C$) 165.7 (C=O), 165.5 (C=O), 165.5 (C=O), 143.6 (Ar), 133.4 (Ar), 133.2 (Ar), 133.1 (Ar), 130.0 (Ar), 129.9 (Ar),129.9 (Ar), 129.8 (Ar), 129.3 (Ar), 129.1 (Ar), 128.6 (Ar), 128.4 (Ar), 128.3 (Ar), 127.8 (Ar), 127.0 (Ar), 105.4 (C-1), 86.9 ($Ph_3C$), 82.3 (C-4), 80.9 (C-2), 76.5 (C-3), 72.1 (C-5), 67.4 (C-6), 62.7 (octyl O$CH_2$), 51.4 (octyl $CH_2N_3$), 29.5 (octyl $CH_2$), 29.3 (octyl $CH_2$), 29.1 (octyl $CH_2$), 28.8 (octyl $CH_2$), 26.7 (octyl $CH_2$), 26.1 (octyl $CH_2$); HR ESIMS: *m/z* [M+Na$^+$] calcd for $C_{54}H_{53}N_3NaO_9$: 910.3814. Found: 910.3815.

**8-Azido-octyl 2,3,5-tri-*O*-benzoyl-β-D-galactofuranoside (S5)** and **8-azidooctyl 2,3,6-tri-*O*-benzoyl-β-D-galactofuranoside (S6)**. A solution of compound **S4** (1.3 g, 1.46 mmol) in $CH_2Cl_2$ (35 mL) was cooled to 0 °C and 5% trifluoroacetic acid in $CH_2Cl_2$ (20 mL) was added dropwise.

The reaction mixture was stirred for 24 h, concentrated, and then co-evaporated with toluene. The residue was purified by chromatography (4:1 Hexane–EtOAc) to give **S5** (423 mg, 45%) and **S6** (329 mg, 35%) both as colourless syrup. $R_f$ 0.28 (**S5**) and $R_f$ 0.35 (**S6**) (3:1 Hexane–EtOAc). Data for **S5**: $[\alpha]_D$ –20.5 (*c* 0.4, $CH_2Cl_2$); $^1$H NMR (400 MHz, $CDCl_3$, $\delta_H$) 8.11–8.03 (m, 3 H, Ar), 8.02–7.95 (m, 2 H, Ar), 7.64–7.48 (m, 3 H, Ar), 7.50–7.26 (m, 7 H, Ar), 5.64 (app dt, 1 H, 7.2, *J* = 4.9 Hz, H-5), 5.60 (d, 1 H, *J* = 5.1 Hz, H-3), 5.48 (d, 1 H, *J* = 1.0 Hz, H-2), 5.31 (s, 1 H, H-1), 4.68–4.60 (m, 1 H, H-4), 4.08 (br s, 2 H, H-6a,b), 3.76–3.74 (m, 1 H, octyl $OCH_2$), 3.58–3.55 (m, 1 H, octyl $OCH_2$), 3.24 (t, 2 H, *J* = 7.0 Hz, octyl $CH_2N_3$), 1.56–1.48 (m, 4 H, octyl $CH_2$), 1.40–1.24 (m, 8 H, octyl $CH_2$); $^{13}$C NMR (125 MHz, $CDCl_3$, $\delta_C$) 166.4 (C=O), 165.8 (C=O), 165.5 (C=O), 133.6 (Ar), 133.5 (Ar), 133.3 (Ar), 130 (Ar), 129.9 (Ar), 129.7 (Ar), 129.6 (Ar), 129.1 (Ar), 129.1 (Ar), 128.6 (Ar), 128.5 (Ar), 128.5 (Ar), 128.4 (Ar), 105.7 (C-1), 82.2 (C-2), 81.9 (C-4), 77.6 (C-5), 73.5 (C-3), 67.6 (octyl $OCH_2$), 62.7 (C-6), 51.0 (octyl $CH_2N_3$), 29.5 (octyl $CH_2$), 29.3 (octyl $CH_2$), 29.1 (octyl $CH_2$), 28.8 (octyl $CH_2$), 26.7 (octyl $CH_2$), 26.1 (octyl $CH_2$); HR ESIMS: *m/z* [M+Na$^+$] calcd for $C_{35}H_{39}N_3NaO_9$: 668.2579. Found: 668.2575. Data for **S6** $[\alpha]_D$ +2.9 (*c* 0.4, $CH_2Cl_2$); $^1$H NMR (400 MHz, $CDCl_3$, $\delta_H$) 8.11–8.02 (m, 6 H, Ar), 7.63–7.52 (m, 3 H, Ar), 7.49–7.39 (m, 6 H, Ar), 5.64 (d, 1 H, *J* = 4.8 Hz, H-3), 5.52 (d, 1 H, *J* = 1.3 Hz, H-2), 5.27 (s, 1 H, H-1), 4.60 (dd, 1 H, *J* = 12.6, 8.0 Hz, H-6a), 4.55–4.44 (m, 2 H, H-5, H-6a), 4.37 (dd, 1 H, *J* = 4.8, 2.2 Hz, H-4), 3.72 (dt, 1 H, *J* = 9.5, 6.7 Hz, octyl $OCH_2$), 3.51 (dt, 1 H, *J* = 9.5, 6.2 Hz, octyl $OCH_2$), 3.21 (t, 2 H, *J* = 7.0 Hz, octyl $CH_2N_3$), 2.40 (s, 1 H, OH), 1.68–1.43 (m, 4 H, octyl $CH_2$), 1.30–1.19 (m, 8 H, octyl $CH_2$); $^{13}$C NMR (125 MHz, $CDCl_3$, $\delta_C$) 166.5 (C=O), 166.0 (C=O), 165.4 (C=O), 133.6 (Ar), 133.1 (Ar), 129.9 (Ar), 129.9 (Ar), 129.8 (Ar), 129.7 (Ar), 129.2 (Ar), 129.1 (Ar), 128.6 (Ar), 128.5 (Ar), 128.4 (Ar), 105.7 (C-1), 83.1 (C-4), 81.5 (C-2), 78.2 (C-3), 69.1 (C-5), 67.6 (C-6), 66.2 (octyl $OCH_2$), 51.4 (octyl $CH_2N_3$), 29.4 (octyl $CH_2$), 29.2 (octyl $CH_2$), 29.1 (octyl $CH_2$), 28.8 (octyl $CH_2$), 26.6 (octyl $CH_2$), 26.0 (octyl $CH_2$); HR ESIMS: *m/z* [M+Na$^+$] calcd for $C_{35}H_{39}N_3NaO_9$: 668.2579. Found: 668.2563.

**Scheme S2**: Synthesis of **Gal*f*$_2$**

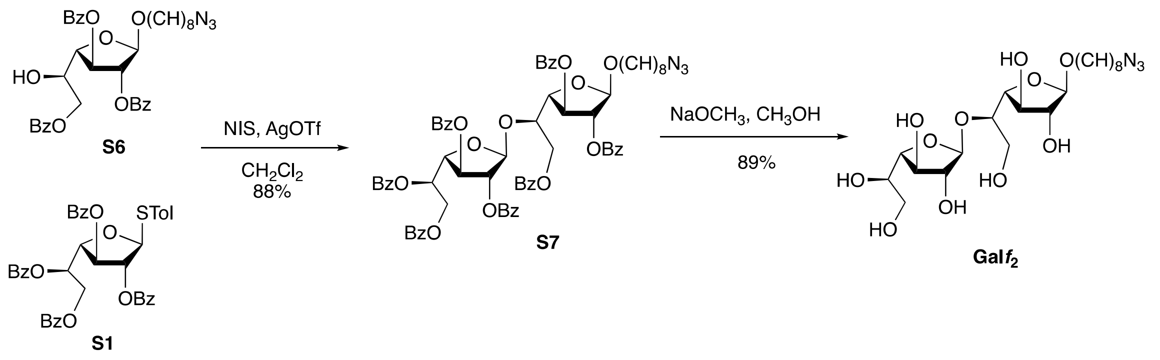

**8-Azido-octyl 2,3,5,6-tetra-*O*-benzoyl-β-D-galactofuranosyl)-(l→5)-2,3,6-tri-*O*-benzoyl-β-D-galactofuranoside (S7).** Acceptor **S6** (110 mg, 0.17 mmol), donor **S1** (176 mg, 0.26 mmol) and activated powdered 4 Å molecular sieves (425 mg) were added to $CH_2Cl_2$ (25 mL). The reaction mixture was cooled to 0 ºC and stirred for 15 min before the addition of NIS (55 mg, 0.23 mmol) and AgOTf (10 mg, 0.042 mmol). After 30 min, $Et_3N$ was added and the mixture was then diluted with $CH_2Cl_2$ and filtered through Celite. The organic layer was washed with a saturated solution of $Na_2S_2O_3$ then brine, dried over $Na_2SO_4$, filtered, concentrated and purified by chromatography to afford disaccharide **S7** (182 mg, 88%) as a colourless oil (2:1 Hexanes–EtOAc). $R_f$ 0.31 (2:1 Hexane–EtOAc); $[\alpha]_D$ –67.2 (*c* 1.5, $CH_2Cl_2$); $^1$H NMR (400 MHz, $CDCl_3$,

$\delta_H$□7.99–7.84 (m, 11 H, Ar), 7.83 (d, 4 H, *J* = 7.8 Hz, Ar), 7.65–7.01 (m, 20 H, Ar), 6.11 (app dt, 1 H, *J* = 7.0, 3.6 Hz, H-5′), 5.86 (dd, 1 H, *J* = 5.1, 1.3 Hz, H-3), 5.82 (s, 1 H, H-1′), 5.72 (s, 1 H, H-2′), 5.63 (dd, 1 H, *J* = 5.1, 1.1 Hz, H-3′), 5.51 (s, 1 H, H-2), 5.22 (s, 1 H, H-1), 5.08 (dd, 1 H, *J* = 5.0, 3.5 Hz, H-4′), 4.75–4.64 (m, 5 H, H-6a, H-6a′, H-5, H-6b, H-6b′), 4.56–4.42 (dd, 1 H, *J* = 5.1, 1.0 Hz, H-4), 3.75 (ddd, 1 H, *J* = 10.0, 6.0, 6.0 Hz, octyl OC$H_2$), 3.51 (ddd, 1 H, *J* = 10.0, 6.0, 6.0 Hz, octyl OC$H_2$), 3.20 (t, 2 H, *J* = 7.0 Hz, octyl C$H_2N_3$), 1.57–1.50 (m, 4 H, octyl OC$H_2$), 1.37–1.22 (m, 8 H, octyl OC$H_2$); $^{13}$C NMR (125 MHz, $CDCl_3$, $\delta_C$) 166.1 (C=O), 166.0 (C=O), 165.6 (C=O), 165.6 (C=O), 165.5 (C=O), 165.4 (C=O), 165.2 (C=O), 133.4 (Ar), 133.4 (Ar), 133.3 (Ar), 133.2 (Ar), 133.1 (Ar), 133.0 (Ar), 132.9 (Ar), 129.9 (Ar), 129.9 (Ar), 129.8 (Ar), 129.8 (Ar), 129.8 (Ar), 129.7 (Ar), 129.8 (Ar), 129.7 (Ar), 129.6 (Ar), 129.5 (Ar), 129.0 (Ar), 128.9 (Ar), 128.9 (Ar), 128.8 (Ar), 128.5 (Ar), 128.4 (Ar), 128.4 (Ar), 128.3 (Ar), 128.2 (Ar), 128.1 (Ar), 105.5 (C-1′), 105.2 (C-1), 82.4 (C-2′), 82.1 (C-2), 82.0 (C-4′), 81.7 (C-4), 77.9 (C-3), 77.5 (C-3′), 77.3 (C-5), 77.1 (C-5′), 70.5 (C-6′), 67.5 (octyl O*C*$H_2$), 64.6 (C-6), 51.4 (octyl *C*$H_2N_3$), 29.4 (octyl *C*$H_2$), 29.2 (octyl *C*$H_2$), 29.1 (octyl *C*$H_2$), 28.8 (octyl *C*$H_2$), 26.6 (octyl *C*$H_2$), 26.1 (octyl *C*$H_2$); HR ESIMS: *m/z* [M+$Na^+$] calcd. for $C_{69}H_{65}N_3O_{18}Na$: 1246.4155. Found: 1246.4158.

**8-Azido-octyl β-D-galactofuranosyl-(1→5)-β-D-galactofuranoside (Gal*f*$_2$).** Application of the Zémplen deacylation procedure to disaccharide **S7** (100 mg, 0.08 mmol) afforded **Gal*f*$_2$** (35 mg, 89%) as a colourless syrup. Purification of the residue was done by chromatography (5:1 $CH_2Cl_2$–MeOH) followed by further purification using $C_{18}$ silica gel ($H_2O$–MeOH) as the eluent. $R_f$ 0.41 (4:1 $CH_2Cl_2$–MeOH); $[\alpha]_D$ –124.2 (*c* 0.5, MeOH); $^1$H NMR (500 MHz, $CD_3OD$, $\delta_H$) 5.17 (br s, 1 H, H-1′), 4.82 (br s, 1 H, H-1), 4.15 (dd, 1 H, *J* = 5.9, 3.7 Hz, H-3′), 4.12–3.97 (m, 3 H, H-2, H-2′, H-3), 3.91–3.87 (m, 1 H, H-4′), 3.81–3.77 (m, 2 H, H-4′, H-5′), 3.75–3.69 (m, 3 H, H-5, H-6a, H-6b), 3.70–3.61 (m, 3 H, H-6′a, H-6′b, octyl OC$H_2$), 3.40 (td, 1 H, *J* = 9.6, 6.5 Hz, octyl OC$H_2$), 3.27 (t, 2 H, *J* = 6.9 Hz, octyl C$H_2N_3$), 1.57–1.48 (m, 4 H, octyl C$H_2$), 1.40–1.32 (m, 8 H, octyl C$H_2$); $^{13}$C NMR (125 MHz, $CD_3OD$, $\delta_C$) 109.3 (C-1′), 109.1 (C-1), 84.8 (C-2′), 83.6 (C-2), 83.4 (C-4′), 82.7 (C-4), 78.7 (C-3), 78.6 (C-3′), 77.1 (C-5), 72.2 (C-5′), 68.9 (C-6), 64.2 (octyl O*C*$H_2$), 62.8 (C-6′), 52.4 (octyl *C*$H_2N_3$), 30.6 (octyl *C*$H_2$), 30.3 (octyl *C*$H_2$), 30.2 (octyl *C*$H_2$), 29.9 (octyl *C*$H_2$), 27.7 (octyl *C*$H_2$), 27.1 (octyl *C*$H_2$); HR ESIMS: *m/z* [M+$Na^+$] calcd. for $C_{20}H_{37}N_3NaO_{11}$: 518.2320. Found: 518.2317.

**Scheme S3**: Synthesis of **Gal*f*3** and **Gal*f*4**

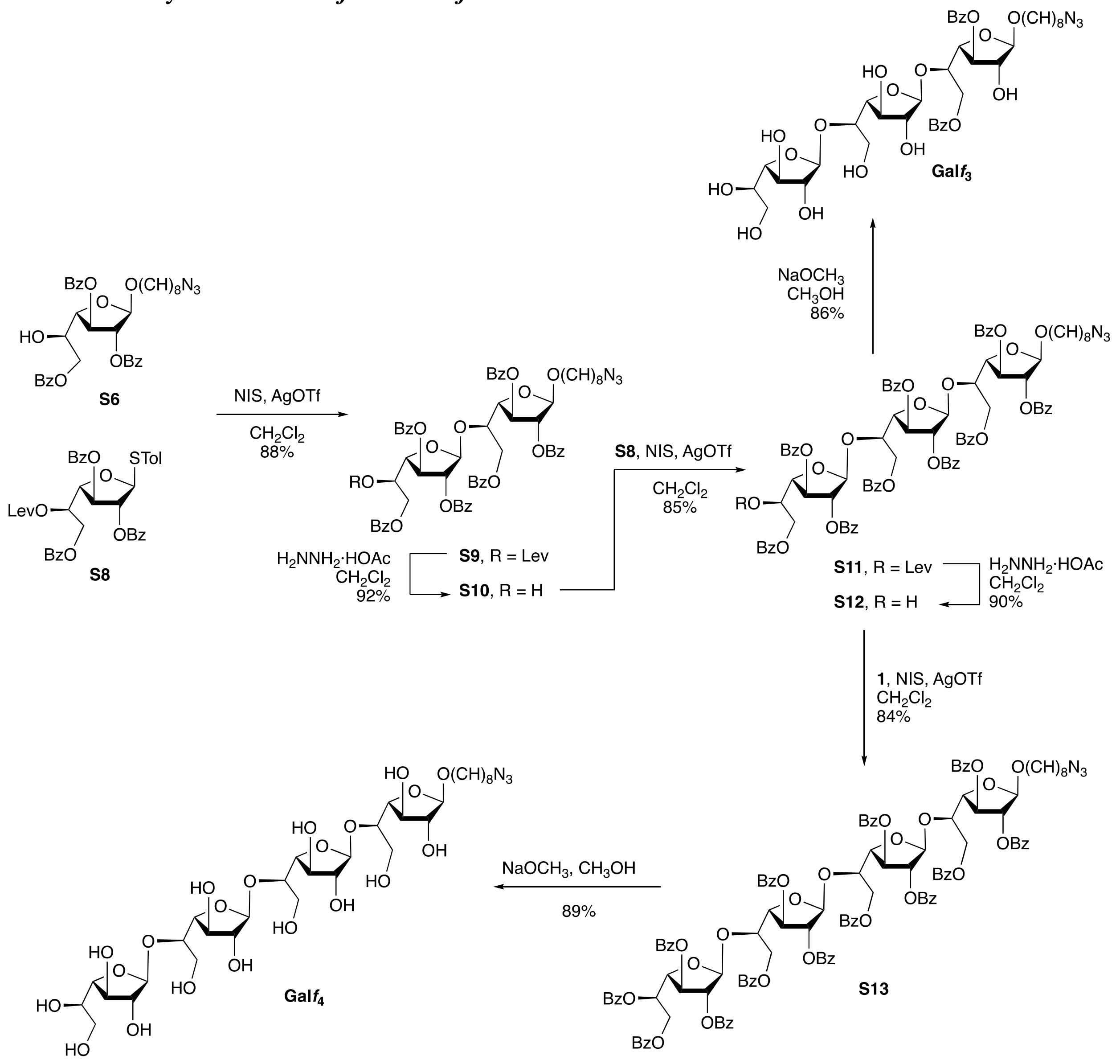

**8-Azido-octyl 2,3,6-tri-*O*-benzoyl-5-*O*-levulinoyl-β-D-galactofuranosyl-(1→5)-2,3,6-tri-*O*-benzoyl-β-D-galactofuranoside (S9).** Acceptor **S6** (100 mg, 0.15 mmol), donor **S8**[19] (160 mg, 0.23 mmol) and activated powdered 4 Å molecular sieves (380 mg) were added to $CH_2Cl_2$ (30 mL) and the reaction mixture was cooled to 0 ºC. The reaction mixture was stirred for 15 min before the addition of NIS (55 mg, 0.23 mmol) and AgOTf (10 mg, 0.034 mmol). After 30 min, $Et_3N$ was added. The mixture was then diluted with $CH_2Cl_2$ and filtered through Celite. The organic layer was washed with a saturated solution of $Na_2S_2O_3$ then brine, dried over anhydrous $Na_2SO_4$, filtered and concentrated. The obtained residue was purified by chromatography (2:1 Hexane–EtOAc) to give **S9** (160 mg, 88%) as a colorless oil $R_f$ 0.28 (2:1 Hexane–EtOAc); $[\alpha]_D$ –0.7 (*c* 0.4, $CH_2Cl_2$); $^1H$ NMR (500 MHz, $CDCl_3$, $\delta_H$) 8.08–7.97 (m, 11 H, Ar), 7.90–7.86 (m, 2 H, Ar), 7.59–7.56 (m, 2 H, Ar), 7.53–7.40 (m, 11 H, Ar), 7.38–7.32 (m, 4 H, Ar), 5.84 (app dt, 1 H, *J* = 7.1, 3.3 Hz, H-5′), 5.81–5.76 (m, 2 H, H-1′, H-3), 5.74 (dd, 1 H, *J* = 1.8, 0.6 Hz, H-2′), 5.56 (dd, 1 H, *J* = 5.2, 1.6 Hz, H-3′), 5.50 (d, 1 H, *J* = 1.1 Hz, H-2), 5.24 (s, 1 H, H-1), 4.91 (dd, 1 H, *J* = 5.1, 4.2 Hz, H-4′), 4.84–4.74 (m, 1 H, H-5), 4.76–4.66 (m, 3 H, H-4, H-6′a,b), 4.58–4.48 (m, 2 H, H-6a,b), 3.74 (td, 1 H, *J* = 9.5, 6.7 Hz, octyl OC$H_2$), 3.51 (td, 1 H, *J* = 9.6, 6.3 Hz, octyl O$CH_2$), 3.23 (t, 2 H, *J* = 6.9 Hz, octyl $CH_2N_3$), 2.67–2.55 (m, 1 H, Lev-$CH_2$) 2.52–2.45 (m, 1 H, Lev-$CH_2$), 2.04 (s, 3 H, Lev-$CH_3$), 1.65–1.54 (m, 4 H, octyl $CH_2$), 1.41–1.26 (m, 8 H, octyl $CH_2$); $^{13}$C NMR (125 MHz, $CDCl_3$, $\delta_C$) 205.9 (Lev C=O), 171.9 (C=O), 166.1 (C=O), 165.9 (C=O), 165.7 (C=O), 165.5 (C=O), 165.4 (C=O), 165.2 (C=O), 133.5 (Ar), 133.3 (Ar), 133.1 (Ar), 129.8 (Ar), 129.1 (Ar), 128.9 (Ar), 128.4 (Ar), 105.5 (C-1′), 105.3 (C-1), 82.3 (C-2), 81.8 (C-2′), 77.4 (C-4), 77.1 (C-4′), 76.9 (C-3), 73.2 (C-3′), 70.1 (C-5′), 69.8 (C-5), 67.5 (octyl O$CH_2$), 64.6 (C-6), 63.5 (C-6′), 51.4 (octyl $CH_2N_3$), 37.9 (Lev-$CH_2$), 29.6 (Lev-$CH_3$), 29.3 (octyl $CH_2$), 29.1 (octyl $CH_2$), 28.8 (octyl $CH_2$), 27.9 (octyl $CH_2$), 26.7 (octyl $CH_2$), 26.1 (octyl $CH_2$); HR ESIMS: *m/z* [M+Na$^+$] calcd for $C_{67}H_{67}N_3NaO_{19}$: 1240.4261. Found: 1240.4260.

**8-Azido-octyl 2,3,6-tri-*O*-benzoyl-β-D-galactofuranosyl-(1→5)-2,3,6-tri-*O*-benzoyl-β-D-galactofuranoside (S10).** Disaccharide **S9** (100 mg, 0.08 mmol) was subjected to the general procedure for the removal of the levulinoyl protecting group. After purification by chromatography (2:1 Hexane–EtOAc) compound **S10** (82 mg, 92%) was obtained as a colourless oil. $R_f$ 0.34 (2:1 Hexane–EtOAc); $[\alpha]_D$ –0.5 (*c* 0.4, $CH_2Cl_2$); $^1$H NMR (500 MHz, $CDCl_3$, $\delta_H$) 8.05–7.90 (m, 10 H, Ar), 7.86 (dd, 2 H, *J* = 8.1, 1.0 Hz, Ar), 7.60–7.28 (m, 16 H, Ar), 7.22 (t, 2 H, *J* = 7.9 Hz, Ar), 5.81–5.78 (m, 1 H, H-5), 5.75 (s, 1 H, H-1′), 5.72 (d, 1 H, *J* = 1.6 Hz, H-2′), 5.65 (app dt, 1 H, *J* = 7.2, 3.2 Hz, H-5′), 5.45 (d, 1 H, *J* = 1.3 Hz, H-2), 5.17 (s, 1 H, H-1), 4.78–4.60 (m, 5 H, H-3, H-3′, H-4′, H-6a′,b′), 4.49–4.38 (m, 3 H, H-4, H-6a,b), 4.38 (s, 1 H, O*H*), 3.67 (dt, 1 H, *J* = 9.5, 6.6 Hz, octyl O$CH_2$), 3.45 (dt, 1 H, *J* = 9.6, 6.3 Hz, octyl O$CH_2$), 3.20 (t, 2 H, *J* = 7.0 Hz, octyl $CH_2N_3$), 1.55–1.48 (m, 4 H, octyl $CH_2$), 1.39–1.19 (m, 8 H, octyl $CH_2$); $^{13}$C NMR (125 MHz, $CDCl_3$, $\delta_C$) 166.4 (C=O), 166.1 (C=O), 165.9 (C=O), 165.8 (C=O), 165.5 (C=O), 165.2 (C=O), 133.5 (Ar), 133.5 (Ar), 133.3 (Ar), 133.1 (Ar), 132.9 (Ar), 129.9 (Ar), 129.8 (Ar), 129.8 (Ar), 129.7 (Ar), 129.1 (Ar), 129 (Ar), 128.9 (Ar), 128.9 (Ar), 128.6 (Ar), 128.5 (Ar), 128.4 (Ar), 128.3 (Ar), 128.2 (Ar), 105.6 (C-1′), 105.5 (C-1), 83.6 (C-2′), 81.8 (C-4), 81.7 (C-2), 78.0 (C-4′), 73.4 (C-3), 72.3 (C-5), 70.2 (C-5′), 69.4 (C-3′), 67.5 (octyl O$CH_2$), 66.2 (C-6), 64.6 (C-6′), 51.4 (octyl $CH_2N_3$), 29.3 (octyl O$CH_2$), 29.2 (octyl O$CH_2$), 29.0 (octyl O$CH_2$), 28.8 (octyl O$CH_2$), 26.6 (octyl O$CH_2$), 26.0 (octyl O$CH_2$); HR ESIMS: *m/z* [M+Na$^+$] calcd for $C_{62}H_{61}N_3NaO_{17}$: 1142.3893. Found: 1142.3883.

**8-Azido-octyl 2,3,6-tri-*O*-benzoyl-5-*O*-levulinoyl-β-D-galactofuranosyl-(1→5)-2,3,6-tri-*O*-benzoyl-β-D-galactofuranosyl-(1→5)-2,3,6-tri-*O*-benzoyl-β-D-galactofuranoside (S11)**. Acceptor **S10** (100 mg, 0.09 mmol) was coupled with donor **S8** (78 mg, 0.11 mmol) in the presence of NIS (25 mg, 0.11 mmol) and AgOTf (5 mg, 0.017 mmol) as described for the synthesis of **S9**. After purification by chromatography (2:1 Hexane–EtOAc) **S11** (129 mg, 85%) was obtained as a colorless syrup, $R_f$ 0.29; $[\alpha]_D$ –3.1 (*c* 1.3, $CH_2Cl_2$); $^1$H NMR (500 MHz, $CDCl_3$, $\delta_H$) 8.06–7.86 (m, 14 H, Ar), 7.78–7.71 (m, 4 H, Ar), 7.59–7.11 (m, 27 H, Ar), 5.81–5.76 (m, 1 H, H-3′′), 5.74 (s, 1 H, H-1′), 5.73 (s, 1 H, H-1′′), 5.71 (m, 1 H, H-3′), 5.68 (d, 1 H, *J* = 1.9 Hz, H-2′), 5.67 (d, 1 H, *J* = 1.7 Hz, H-2′′), 5.49 (dd, 1 H, *J* = 7.1, 3.3 Hz, H-5′′), 5.46 (d, 1 H, *J* = 1.2 Hz, H-2), 5.19 (s, 1 H, H-1), 4.89–4.78 (m, 2 H, H-4′, H-4′′), 4.77–4.58 (m, 8 H, H-3, H-5, H-5′, H-6a, H-6a′,b′, H-6a′′,b′′), 4.52–4.41 (m, 2 H, H-4, H-6b), 3.68 (dt, 1 H, *J* = 9.5, 6.7 Hz, octyl O$CH_2$), 3.46 (dt, 1 H, *J* = 9.6, 6.2 Hz, octyl O$CH_2$), 3.19 (t, 2 H, *J* = 7.0 Hz, octyl $CH_2N_3$), 2.64–2.28 (m, 4 H, Lev-$CH_2$), 1.98 (s, 3 H, Lev-$CH_3$), 1.55–1.39 (m, 4 H, octyl $CH_2$), 1.38–1.24 (m, 8 H, octyl $CH_2$); $^{13}$C NMR (125 MHz, $CDCl_3$, $\delta_C$) 205.5 (Lev-C=O), 171.5 (C=O), 166.2 (C=O), 165.9 (C=O), 165.9 (C=O), 165.7 (C=O), 165.6 (C=O), 165.5 (C=O), 165.4 (C=O), 165.3 (C=O), 165.1 (C=O), 133.5 (Ar), 133.5 (Ar), 133.4 (Ar), 133.4 (Ar), 133.2 (Ar), 132.9 (Ar), 132.9 (Ar), 129.9 (Ar), 129.9 (Ar), 129.8 (Ar), 129.8 (Ar), 129.8 (Ar), 129.7 (Ar), 129.6 (Ar), 129.1 (Ar), 129.1 (Ar), 129.0 (Ar), 128.9 (Ar), 128.8 (Ar), 128.8 (Ar), 128.6 (Ar), 128.6 (Ar), 128.5 (Ar), 128.4 (Ar), 128.3 (Ar), 128.3 (Ar), 128.2 (Ar), 128.1 (Ar), 105.4 (C-1′, C-1′′),

105.3 (C-1), 83.2 (C-4′, C-4′′), 82.6 (C-2′), 81.9 (C-2), 81.8 (C-2′′), 81.7 (C-4), 77.2 (C-5), 77.1 (C-5′′), 73.2 (C-3), 73.1 (C-3′), 72.9 (C-3′′), 70.1 (C-5′), 67.4 (octyl O*C*$H_2$), 65.0 (C-6), 64.4 (C-6′), 63.7 (C-6′′), 51.4 (octyl *C*$H_2N_3$), 37.8 (Lev-*C*$H_2$), 29.6 (Lev-*C*$H_3$), 29.4 (Lev-*C*$H_2$), 29.3 (octyl *C*$H_2$), 29.1 (octyl *C*$H_2$), 28.8 (octyl *C*$H_2$), 27.9 (octyl *C*$H_2$), 26.7 (octyl *C*$H_2$), 26.0 (octyl *C*$H_2$); HR ESIMS: *m/z* [M+Na$^+$] calcd for $C_{94}H_{89}N_3NaO_{27}$: 1714.5576. Found: 1714.5564.

**8-Azido-octyl 2,3,6-tri-*O*-benzoyl-β-D-galactofuranosyl-(1→5)-2,3,6-tri-*O*-benzoyl-β-D-galactofuranosyl-(1→5)-2,3,6-tri-*O*-benzoyl-β-D-galactofuranoside (S12).** Application of the general procedure for the removal of the levulinoyl group to **S11** (80 mg, 0.047 mmol) gave the product, which was purified by chromatography (2:1 Hexane–EtOAc) to give **S12** (67 mg, 90%) as a colourless syrup, $R_f$ 0.29 (2:1 Hexane–EtOAc); [α]$_D$ –1.2 (*c* 0.5, $CH_2Cl_2$); $^1$H NMR (500 MHz, $CDCl_3$, $\delta_H$) 8.04–7.99 (m, 3 H, Ar), 7.96–7.81 (m, 12 H, Ar), 7.56 (dd, 1 H, *J* = 14.7, 7.1 Hz, Ar), 7.33–7.10 (m, 29 H, Ar), 5.82–5.80 (m, 2 H, H-2, H-2′), 5.73 (d, 2 H, *J* = 2.8 Hz, H-3, H-3′), 5.72 (s, 1 H, H-2′,′) 5.67 (s, 1 H, H-1′′), 5.64–5.59 (m, 1 H, H-3′′), 5.47 (s, 1 H, H-1′), 5.20 (s, 1 H, H-1), 4.87 (dd, 1 H, *J* = 5.0, 3.1 Hz, H-5′′), 4.77–4.60 (m, 7 H, H-4, H-4′, H-6a,b, H-6′a,b, H-6′′a), 4.50–4.48 (m, 1 H, H-4′′), 4.45 (d, 2 H, *J* = 5.4 Hz, H-5, H-5′), 4.39–4.32 (m, 2 H, H-6′′b, OH), 3.76–3.63 (m, 1 H, octyl O*C*$H_2$), 3.47 (td, 1 H, *J* = 9.6, 6.3 Hz, octyl OC$H_2$), 3.19 (t, 2 H, *J* = 6.9 Hz, octyl C$H_2N_3$), 1.55–1.38 (m, 4 H, octyl C$H_2$), 1.41–1.22 (m, 8 H, octyl C$H_2$); $^{13}$C NMR (125 MHz, $CDCl_3$, $\delta_C$) 166.3 (C=O), 166.1 (C=O), 165.9 (C=O), 165.8 (C=O), 165.7 (C=O), 165.6 (C=O), 165.4 (C=O), 165.2 (C=O), 165.1 (C=O), 133.7 (Ar), 133.4 (Ar), 133.4 (Ar), 133.4 (Ar), 133.3 (Ar), 133.2 (Ar), 133.2 (Ar), 132.9 (Ar), 132.9 (Ar), 132.8 (Ar), 129.9 (Ar), 129.8 (Ar), 129.8 (Ar), 129.8 (Ar), 129.7 (Ar), 129.7 (Ar), 129.6 (Ar), 129.6 (Ar), 129.6 (Ar), 129.0 (Ar), 129.0 (Ar), 128.9 (Ar), 128.9 (Ar), 128.7 (Ar), 128.6 (Ar), 128.6 (Ar), 128.5 (Ar), 128.4 (Ar), 128.3 (Ar), 128.3 (Ar), 128.2 (Ar), 128.2 (Ar), 128.1 (Ar), 105.6 (C-1′′), 105.4 (C-1′), 105.2 (C-1), 83.5 (C-4′), 82.8 (C-4′′), 82.5 (C-2′), 81.9 (C-2), 81.0 (C-2′′), 81.6 (C-4), 77.9 (C-5), 77.1 (C-5′′), 73.1 (C-3, C-3′′), 73.0 (C-3′), 69.5 (C-5′), 67.4 (octyl O*C*$H_2$), 66.2 (C-6), 65.0 (C-6′), 64.4 (C-6′′), 51.4 (octyl *C*$H_2N_3$), 29.4 (octyl *C*$H_2$), 29.2 (octyl *C*$H_2$), 29.0 (octyl *C*$H_2$), 28.7 (octyl *C*$H_2$), 26.6 (octyl *C*$H_2$), 26.0 (octyl *C*$H_2$); HR ESIMS: *m/z* [M+Na$^+$] calcd for $C_{89}H_{83}N_3NaO_{25}$: 1616.5208. Found: 1616.5195.

**8-Azido-octyl 2,3,5,6-tetra-*O*-benzoyl-β-D-galactofuranosyl-(1→5)-2,3,6-tri-*O*-benzoyl-β-D-galactofuranosyl-(1→5)-2,3,6-tri-*O*-benzoyl-β-D-galactofuranosyl-(1→5)-2,3,6-tri-*O*-benzoyl β-D-galactofuranoside (S13).** Trisaccharide acceptor **S12** (60 mg, 0.038 mmol) and thioglycoside donor **S1** (33 mg, 0.047 mmol) were dissolved in $CH_2Cl_2$ (20 mL). The reaction mixture was stirred for 15 min before the addition of NIS (10 mg, 0.038 mmol) and AgOTf (2 mg, 0.006 mmol). After 30 min, $Et_3N$ was added. The mixture was diluted with $CH_2Cl_2$ and filtered through Celite. The organic layer was washed with a saturated solution of $Na_2S_2O_3$ then brine, dried over $Na_2SO_4$, filtered, and concentrated. The product was purified by chromatography (2:1 Hexane–EtOAc) to give **S13** (70 mg, 84%) as a colorless syrup. $R_f$ 0.35 (2:1 Hexane–EtOAc); [α]$_D$ –8.8 (*c* 0.3, $CH_2Cl_2$); $^1$H NMR (500 MHz, $CDCl_3$, $\delta_H$) 8.05–7.87 (m, 22 H, Ar), 7.76–7.70 (m, 6 H, Ar), 7.51–7.06 (m, 37 H, Ar), 6.01 (m, 1 H, *J* = 6.3, 3.5 Hz, H-3′′), 5.85 (ddd, 2 H, *J* = 8.8, 5.0, 1.1 Hz, H-3, H-3′), 5.80 (d, 1 H, *J* = 4.9 Hz, H-3′′′), 5.78 (d, 1 H, *J* = 3.7 Hz, H-2′), 5.74 (s, 2 H, H-1′′, H-1′′′), 5.70 (dd, 2 H, *J* = 5.1, 1.5 Hz, H-2, H-5′′′), 5.66 (d, 1 H, *J* = 1.4 Hz, H-2′′′), 5.63–5.60 (m, 2 H, H-3, H-5′′′), 5.59 (dd, 1 H, *J* = 5.2, 1.2 Hz, H-2′′), 5.45 (d, 1 H, *J* = 1.2 Hz, H-1′), 5.19 (s, 1 H, H-1), 5.00 (dd, 1 H, *J* = 5.1, 3.5 Hz, H-6a), 4.90–4.83 (m, 4 H, H-5, H-5′, H-5′′, H-6b), 4.77–4.59 (m, 9 H, H-4, H-4′, H-4′′′, H-6′a,b, H-6′′a,b, H-6′′′a,b), 4.47 (dd, 1 H, *J* = 4.8, 3.6 Hz, H-4′′), 3.67 (td, 1 H, *J* = 9.6, 6.7 Hz, octyl OC$H_2$), 3.45 (td, 1 H, *J* = 9.6, 6.2 Hz, octyl C$H_2N_3$), 3.19 (dd, 2 H, *J* = 9.2, 4.8 Hz, octyl OC$H_2$), 1.59–1.49 (m, 4 H, octyl C$H_2$), 1.33–1.23 (m, 8 H, octyl C$H_2$); $^{13}$C NMR (125 MHz, $CDCl_3$, $\delta_C$)

166.1 (C=O), 165.9 (C=O), 165.9 (C=O), 165.8 (C=O), 165.6 (C=O), 165.6 (C=O), 165.5 (C=O), 165.4 (C=O), 165.2 (C=O), 165.2 (C=O), 165.1 (C=O), 133.4 (Ar), 133.4 (Ar), 133.3 (Ar), 133.2 (Ar), 133.1 (Ar), 133.0 (Ar), 132.9 (Ar), 132.9 (Ar), 132.8 (Ar), 132.7 (Ar), 129.9 (Ar), 129.8 (Ar), 129.7 (Ar), 129.6 (Ar), 129.6 (Ar), 129.0 (Ar), 128.9 (Ar), 128.9 (Ar), 128.7 (Ar), 128.6 (Ar), 128.6 (Ar), 128.4 (Ar), 128.3 (Ar), 128.2 (Ar), 128.2 (Ar), 128.1 (Ar), 128.0 (Ar), 105.4 (C-1′′′, C-1′′), 105.2 (C-1′), 105.2 (C-1), 83.5 (C-4′), 83.2 (C-4′′′), 82.6 (C-4′′), 82.0 (C-2′), 82.0 (C-2′′), 81.8 (C-2), 81.8 (C-2′′′), 81.6 (C-4), 77.8 (C-5), 77.7 (C-5′′), 77.0, (C-5′′′), 76.9 (C-5′), 73.1 (C-3), 72.9 (C-3′), 72.7 (C-3′′′), 70.5 (C-3′′), 67.3 (octyl O*C*$H_2$), 65.3 (C-6), 64.9 (C-6′′′), 64.5 (C-6′), 63.9 (C-6′′), 51.4 (octyl *C*$H_2N_3$), 29.4 (octyl *C*$H_2$), 29.2 (octyl *C*$H_2$), 29.0 (octyl *C*$H_2$), 28.7 (octyl *C*$H_2$), 26.6 (octyl *C*$H_2$), 26.0 (octyl *C*$H_2$); HR ESIMS: *m/z* [M+Na$^+$] calcd for $C_{123}H_{109}N_3NaO_{34}$: 2194.6785. Found: 2194.6767.

**8-Azido-octyl β-D-galactofuranosyl-(1→5)-β-Dgalactofuranosyl-(1→5)-β-D-galacto-furanoside (Gal*f*3).** Application of the general Zémplen deprotection procedure to **S11** (1.14 g, 0.673 mmol), gave a product that was was purified using $C_{18}$ silica gel with a $H_2O$–MeOH gradient as the eluent to give **Gal*f*3** (0.647 g, 86%) as a fluffy solid, $R_f$ 0.03 (9:1 $CH_2Cl_2$–MeOH); $^1$H NMR (700 MHz, $D_2O$, $\delta_H$): 5.22 (d, *J* = 2.0 Hz, 1H, H-1′′), 5.19 (d, *J* = 1.9 Hz, 1H, H-1′), 4.97 (d, *J* = 2.4 Hz, 1H, H-1), 4.18–4.14 (m, 3H, H-2′, H-2′′, H-3), 4.13–4.06 (m, 4H, H-4, H-4′′, H-4′, H-3′), 4.05–4.02 (m, 2H, H-2, H-3′′), 3.98–3.95 (m, 1H, H-5′), 3.94–3.91 (m, 1H, H-5′′), 3.86–3.83 (m, 1H, H-5), 3.82–3.77 (m, 4H, H-6′′a,b, H-6′a,b), 3.75–3.71 (m, 2H, H-6a, octyl OC*$H_2$*), 3.68 (dd, *J* = 11.6, 7.4 Hz, 1H, H-6b), 3.58 (dt, *J* = 10.0, 6.5 Hz, 1H, octyl OC*$H_2$*), 3.32 (t, *J* = 6.9 Hz, 2H, octyl C*$H_2$*$N_3$), 1.64–1.58 (m, 4H, octyl C*$H_2$*), 1.40–1.32 (m, 8H octyl C*$H_2$*). $^{13}$C NMR (125 MHz, $D_2O$, $\delta_C$): 108.1 (2 C, C-1′, C-1′′), 107.9 (C-1), 83.6, 82.5, 82.4, 82.3, 82.2, 82.0, 77.5, 77.36, 77.3, 77.1 (C-5′′), 76.5 (C-5′), 71.5 (C-5), 69.6 (octyl O*C*$H_2$), 63.8 (C-6), 62.2 (C-6′), 62.1 (C-6′′), 52.2 (octyl *C*$H_2N_3$), 29.6 (octyl *C*$H_2$), 29.2 (octyl *C*$H_2$), 29.1 (octyl *C*$H_2$), 28.9 (octyl *C*$H_2$), 26.8 (octyl *C*$H_2$), 26.0 (octyl *C*$H_2$). ESI-MS *m/z* calcd. for [M + Na]$^+$ $C_{26}H_{47}N_3O_{16}Na$: 680.2849. Found 680.2852.

**8-Azido-octyl β-D-galactofuranosyl-(1→5)-β-D-galactofuranosyl-(1→5)-β-D galactofuranosyl- (1→5)-β-D-galactofuranoside (Gal*f*4).** Application of the general Zémplen deprotection procedure to **S13** (30 mg, 0.013 mmol) gave a product that was initially purified by chromatography (3:1 $CH_2Cl_2$–MeOH). The solvent was evaporated and the residue was dissolved in water and was filtered through $C_{18}$ silica gel using $H_2O$–MeOH as eluent to give **Gal*f*4** (9 mg, 87%) as a colouless syrup, $R_f$0.42 (3:1 $CH_2Cl_2$–MeOH); $[\alpha]_D$ –35.6 (*c* 0.5, MeOH); $^1$H NMR (500 MHz, $CD_3OD$, $\delta_H$) 5.20 (d, 1 H, *J* = 2.0 Hz, H-1′′′), 5.17 (s, 2 H, H-1′, H-1′′), 4.94 (d, 1 H, *J* = 2.3 Hz, H-1), 4.15–4.00 (m, 10 H), 3.97–3.88 (m, 4 H), 3.85–3.80 (m, 2 H), 3.80–3.74 (m, 8 H), 3.73–3.63 (m, 1 H, octyl OC*$H_2$*), 3.55 (d, 1 H, *J* = 9.9 Hz, octyl OC*$H_2$*), 3.30 (t, 2 H, *J* = 6.9 Hz, octyl C*$H_2$*$N_3$), 1.62–1.55 (m, 4 H, octyl *C*$H_2$), 1.35–1.30 (m, 8 H, octyl *C*$H_2$); $^{13}$C NMR (125 MHz, $CD_3OD$, $\delta_C$) 109.2 (C-1′′′), 109.0 (C-1), 108.9 (C-1′′), 108.7 (C-1′′′), 85.1, 84.5, 83.6, 83.5, 83.1, 83.0, 82.7, 78.8, 78.7, 78.6, 78.5, 77.2, 71.0, 70.6, 68.9 (octyl O*C*$H_2$), 64.2, 63.5, 62.7, 52.4 (octyl *C*$H_2N_3$), 30.6 (octyl *C*$H_2$), 30.4 (octyl *C*$H_2$), 30.2 (octyl *C*$H_2$), 29.9 (octyl *C*$H_2$), 27.8 (octyl *C*$H_2$), 27.1 (octyl *C*$H_2$); HR ESIMS: *m/z* [M+Na$^+$] calcd for $C_{32}H_{57}N_3NaO_{21}$: 842.3377. Found: 842.3374.

**3.5** *Procedures for synthesis of* ***DiN₃, TriAN₃*** *and* ***TriBN₃***

**Scheme S4.** Synthesis of **S15**

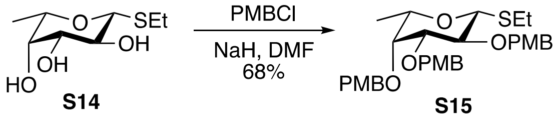

**Scheme S5**. Synthesis of **DiN₃**, **TriAN₃** and **TriBN₃**

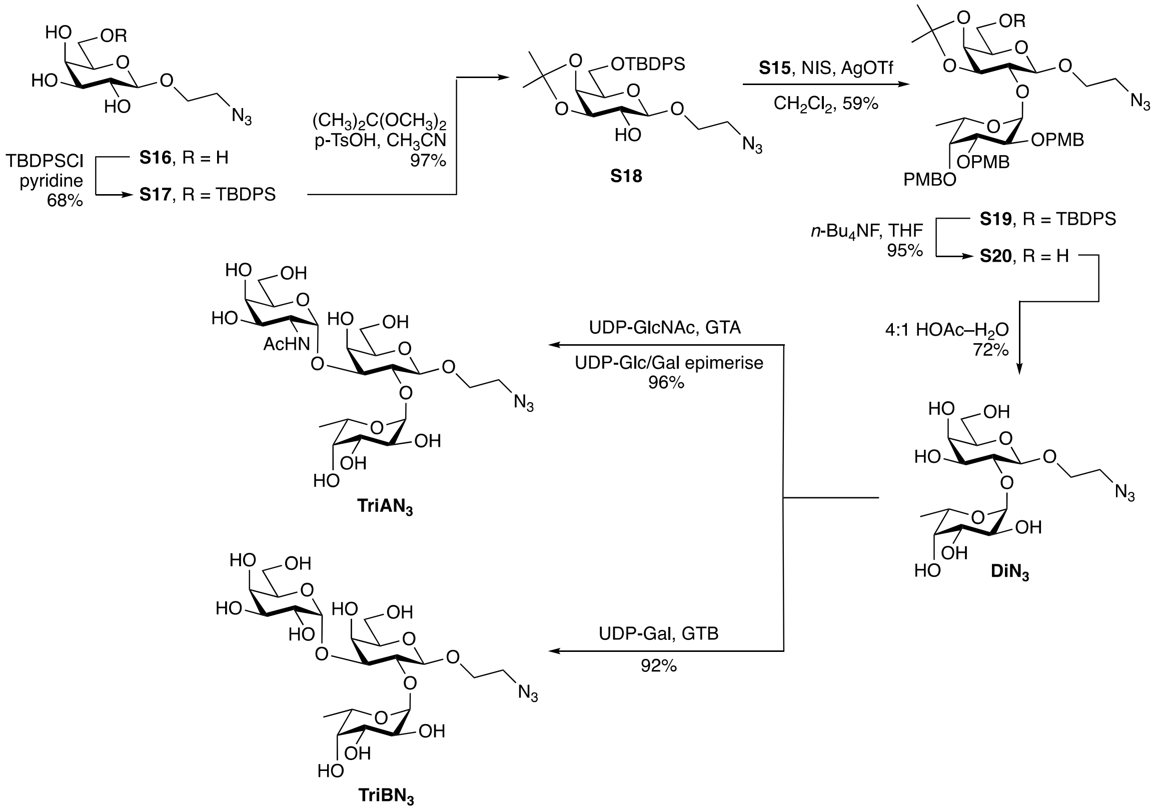

**Ethyl 2,3,4-tri-*O*-*p*-methoxybenzyl-1-thio-β-L-fucopyranoside (S15)**. To a stirred solution of ethyl 1-thio-β-L-fucopyranoside (**S14**)[20] (2.658 g, 12.77 mmol) in dry DMF (50 mL) was added sodium hydride (3.07 g, 76.64 mmol, 60% dispersion in mineral oil) portion-wise at 0 °C. After stirring for 30 min, *p*-methoxybenzyl chloride (6.24 mL, 46 mmol) was added dropwise. Then reaction mixture was warmed to room temperature and stirred for 12 h. After the reaction was complete, $CH_3OH$ was added at 0 °C, the solution was concentrated and the crude residue was purified by chromatography (Hexane–EtOAc, 5:1) to afford **S15** (4.87 g, 68%) as a white solid: $R_f$ = 0.35 (Hexane–EtOAc, 4:1); $[\alpha]_D^{25}$ = –28.6 (*c* 2.1, $CH_2Cl_2$); $^1$H NMR (500 MHz, $CDCl_3$, $\delta_H$) 7.35–7.27 (m, 6 H, Ar), 6.91–6.84 (m, 6 H, Ar), 4.90 (d, 1 H, *J* = 11.5 Hz, ArC$H_2$O), 4.81 (d, 1 H, *J* = 10.0 Hz, ArC$H_2$O), 4.73 (d, 1 H, *J* = 10.0 Hz, ArC$H_2$O), 4.70 (d, 1 H, *J* = 11.5 Hz, ArC$H_2$O), 4.66 (d, 1 H, *J* = 11.5 Hz, ArC$H_2$O), 4.63 (d, 1 H, *J* = 11.5 Hz, ArC$H_2$O), 4.36 (d, 1 H, *J* = 9.5 Hz, H-1), 3.82 (s, 3 H, OC$H_3$), 3.81 (s, 3 H, OC$H_3$), 3.80 (s, 3 H, OC$H_3$), 3.77 (t, 1 H, *J* = 9.5 Hz, H-2), 3.55 (dd, 1 H, *J* = 3.0, 1.0 Hz, H-4), 3.51 (dd, 1 H, *J* = 9.5, 3.0 Hz, H-3), 3.44 (qd, 1 H, *J* = 6.0, 1.0 Hz, H-5), 2.76 (dq, 1 H, *J* = 12.5, 7.5 Hz, SC$H_2$CH$_3$), 2.69 (dq, 1 H, *J* =

12.5, 7.5 Hz, $SCH_2CH_3$), 1.29 (t, 3 H, *J* = 7.5 Hz, $SCH_2CH_3$), 1.16 (d, 3 H, *J* = 6.0 Hz, H-6); $^{13}C$ NMR (125 MHz, $CDCl_3$, $\delta_C$) 159.4 (Ar), 159.3 (Ar), 159.2 (Ar), 131.0 (Ar), 130.9 (Ar), 130.8 (Ar), 130.2 (Ar), 130.0 (Ar), 129.3 (Ar), 114.0 (Ar), 113.9 (Ar), 113.6 (Ar), 85.1 (*C*-1), 84.4 (*C*-3), 78.2 (*C*-2), 76.1 (*C*-4), 75.4 ($ArCH_2O$), 74.7 (*C*-5), 74.1 ($ArCH_2O$), 72.7 ($ArCH_2O$), 55.41 ($OCH_3$ × 2), 55.39 ($OCH_3$), 24.8 ($SCH_2CH_3$), 17.4 (*C*-6), 15.1 ($SCH_2CH_3$); HR ESIMS: *m*/*z* $[M+NH_4]^+$ calcd for $C_{32}H_{44}NO_7S$: 586.2833. Found: 586.2838; $[M+Na]^+$ calcd for $C_{32}H_{40}NaO_7S$: 591.2387. Found: 591.2381.

**Azidoethyl 6-*O*-*tert*-butyldiphenylsilyl-β-D-galactopyranoside (S17)**. To a solution of **S16**[21] (1.97 g, 7.93 mmol) in dry pyridine (30 mL) was added *tert*-butyldiphenylchlorosilane (2.68 mL, 10.31 mmol) at room temperature. After stirring for 12 h, $CH_3OH$ was added, the solution was concentrated and the crude residue was purified by chromatography ($CH_2Cl_2$–$CH_3OH$, 30:1) to give **S17** (2.57 g, 68%) as a colorless syrup: $R_f$ = 0.35 ($CH_2Cl_2$–$CH_3OH$, 30:1); $[\alpha]_D^{25}$ = –9.2 (*c* 1.0, $CH_3OH$); $^1H$ NMR (500 MHz, $CD_3OD$, $\delta_H$) $\delta$ 7.72–7.70 (m, 4 H, Ar), 7.46–7.39 (m, 6 H, Ar), 4.24 (d, 1 H, *J* = 7.6 Hz, H-1), 3.95–3.85 (m, 4 H, $OCH_2CH_2N_3$, H-6a, H-4, H-6b), 3.69 (ddd, 1 H, *J* = 10.8, 5.8, 4.9 Hz, $OCH_2CH_2N_3$), 3.62 (ddd, 1 H, *J* = 6.5, 5.6, 0.9 Hz, H-5), 3.53 (dd, 1 H, *J* = 9.7, 7.6 Hz, H-2), 3.46 (dd, 1 H, *J* = 9.7, 3.4 Hz, H-3), 3.45–3.42 (m, 2 H, $OCH_2CH_2N_3$), 1.05 (s, 9 H, $SiC(CH_3)_3$); $^{13}C$ NMR (125 MHz, $CD_3OD$, $\delta_C$) 136.7 (Ar), 134.7 (Ar), 134.6 (Ar), 130.92 (Ar), 130.91 (Ar), 128.83 (Ar), 128.82 (Ar), 105.1 (*C*-1), 76.9 (*C*-5), 75.0 (*C*-3), 72.5 (*C*-2), 70.3 (*C*-4), 69.2 ($OCH_2CH_2N_3$), 64.6 (*C*-6), 52.1 ($OCH_2CH_2N_3$), 27.3 ($SiC(CH_3)_3$), 20.0 ($SiC(CH_3)_3$); HR ESIMS: *m*/*z* $[M+NH_4]^+$ calcd for $C_{24}H_{37}N_4O_6Si$: $[M+NH_4]^+$ 505.2477. Found: 505.2483; $[M+Na]^+$ calcd for $C_{24}H_{33}N_3NaO_6Si$: 510.2031. Found: 510.2026.

**2-Azidoethyl 3,4-*O*-isopropylidene-6-*O*-*tert*-butyldiphenylsilyl-β-D-galactopyranoside (S18)**. To a stirred solution of triol **S17** (2.57 g, 5.28 mmol) in dry $CH_3CN$ (30 mL) were added 2,2-dimethoxypropane (0.97 mL, 7.91 mmol) and *p*-toluenesulfonic acid monohydrate (100 mg, 528 μmol) successively. The reaction mixture was stirred at room temperature for 2 h and then triethylamine was added. The solution was concentrated and the crude residue was purified by flash column chromatography (Hexane–EtOAc, 2:1) to give **S18** (2.70 g, 97%) as a colorless syrup: $R_f$ = 0.31 (Hexane–EtOAc, 2:1); $[\alpha]_D^{25}$ = –1.0 (*c* 1.1, $CH_2Cl_2$); $^1H$ NMR (500 MHz, $CDCl_3$, $\delta_H$) 7.71–7.68 (m, 4 H, Ar), 7.46–7.36 (m, 6 H, Ar), 4.27 (dd, 1 H, *J* = 5.5, 2.1 Hz, H-4), 4.21 (d, 1 H, *J* = 8.3 Hz, H-1), 4.08 (dd, 1 H, *J* = 7.4, 5.5 Hz, H-3), 4.04 (ddd, 1 H, *J* = 10.6, 5.0, 3.8 Hz, $OCH_2CH_2N_3$), 3.99 (dd, 1 H, *J* = 10.0, 7.0 Hz, H-6a), 3.93 (dd, 1 H, *J* = 10.0, 6.2 Hz, H-6b), 3.88 (ddd, 1 H, *J* = 7.0, 6.2, 2.1 Hz, H-5), 3.68 (ddd, 1 H, *J* = 10.6, 8.2, 3.6 Hz, $OCH_2CH_2N_3$), 3.57 (ddd, 1 H, *J* = 8.3, 7.4, 2.4 Hz, H-2), 3.53 (ddd, 1 H, *J* = 13.1, 8.2, 3.8 Hz, $OCH_2CH_2N_3$), 3.32 (ddd, 1 H, *J* = 13.1, 5.0, 3.6 Hz, $OCH_2CH_2N_3$), 2.43 (d, 1 H, *J* = 2.4 Hz, 2-O*H*), 1.52 (s, 3 H, $CH_3C$), 1.35 (s, 3 H, $CH_3C$), 1.06 (s, 9 H, $SiC(CH_3)_3$); $^{13}C$ NMR (125 MHz, $CDCl_3$, $\delta_C$) 135.8 (Ar), 135.7 (Ar), 133.6 (Ar), 133.4 (Ar), 129.9 (Ar), 127.9 (Ar), 127.8 (Ar), 110.3 ($C(CH_3)_2$), 102.7 (*C*-1), 78.7 (*C*-3), 74.0 (*C*-5), 73.9 (*C*-2), 73.3 (*C*-4), 68.7 ($OCH_2CH_2N_3$), 62.8 (*C*-6), 50.9 ($OCH_2CH_2N_3$), 28.3 ($CH_3C$), 26.9 ($SiC(CH_3)_3$), 26.4 ($CH_3C$), 19.4 ($SiC(CH_3)_3$); HR ESIMS: *m*/*z* $[M+NH_4]^+$ calcd for $C_{27}H_{41}N_4O_6Si$: 545.2790. Found: 545.2796. $[M+Na]^+$ calcd for $C_{27}H_{37}N_3NaO_6Si$: 550.2344. Found: 550.2345.

**2-Azidoethyl 2,3,4-tri-*O*-*p*-methoxybenzyl-α-L-fucopyranosyl-(1→2)-3,4-*O*-isopropylidene-6-*O*-(*tert*-butyldiphenylsilyl)-β-D-galactopyranoside (S19)**. To a stirred solution of acceptor **S18** (521.8 mg, 1.07 mmol) and donor **S15** (670 mg, 1.18 mmol) in dry $CH_2Cl_2$ (30 mL) was added molecular sieves (3.0 g, 4Å, powder). After stirring for 30 min at room temperature, the reaction mixture was allowed to cool to 0 °C, and then *N*-iodosuccinimide (345 mg, 1.53 mmol) and trifluoromethanesulfonic acid (14.2 μL, 161 μmol) were added successively. The resulting solution was stirred for 1 h at room temperature before triethylamine was added. The solution was filtered and the filtrate was washed with saturated $NaHCO_3$ (aq.). The aqueous layer was extracted with $CH_2Cl_2$ (20 mL × 3), dried over $Na_2SO_4$, filtered, concentrated and the residue was purified by chromatography (Hexane–EtOAc, 4:1) to give **S19** (652 mg, 59%) as a colorless syrup: $R_f$ = 0.44 (Hexane–EtOAc, 3:1); $[\alpha]_D^{25}$ = –63.0 (*c* 0.8, $CH_2Cl_2$); $^1$H NMR (500 MHz, $CDCl_3$, $\delta_H$) 7.73–7.70 (m, 4 H, Ar), 7.46–7.35 (m, 10 H, Ar), 7.28–7.26 (m, 2 H, Ar), 6.93–6.91 (m, 2 H, Ar), 6.88–6.84 (m, 4 H, Ar), 5.50 (d, 1 H, *J* = 3.7 Hz, H-1′), 4.90 (d, 1 H, *J* = 11.4 Hz, ArC*H*₂O), 4.81 (d, 1 H, *J* = 11.2 Hz, ArC*H*₂O), 4.74 (s, 2 H, ArC*H*₂O), 4.67 (d, 1 H, *J* = 11.2 Hz, ArC*H*₂O), 4.60 (d, 1 H, *J* = 11.4 Hz, ArC*H*₂O), 4.37 (d, 1 H, *J* = 8.3 Hz, H-1), 4.32 (dd, 1 H, *J* = 6.6, 5.4 Hz, H-3), 4.25 (dd, 1 H, *J* = 5.4, 1.3 Hz, H-4), 4.21 (qd, 1 H, *J* = 6.6, 1.5 Hz, H-5′), 4.04 (dd, 1 H, *J* = 10.2, 3.7 Hz, H-2′), 4.01–3.92 (m, 4 H, H-6a, $OCH_2CH_2N_3$, H-6b, H-3′), 3.85–3.82 (m, 10 H, $OCH_3$, $OCH_3$, $OCH_3$, H-5), 3.80 (t, 1 H, *J* = 8.0 Hz, H-2), 3.66 (d, 1 H, *J* = 1.5 Hz, H-4′), 3.55 (dt, 1 H, *J* = 11.5, 5.8 Hz, $OCH_2CH_2N_3$), 3.39–3.37 (m, 2 H, $OCH_2CH_2N_3$), 1.51 (s, 3 H, $CH_3C$), 1.38 (s, 3 H, $CH_3C$), 1.09 (s, 9 H, $SiC(CH_3)_3$), 1.07 (d, 3 H, *J* = 6.6 Hz, H-6′); $^{13}$C NMR (125 MHz, $CDCl_3$, $\delta_C$) 159.3 (Ar), 159.22 (Ar), 159.16 (Ar), 135.8 (Ar), 135.7 (Ar), 133.6 (Ar), 133.4 (Ar), 131.5 (Ar), 131.2 (Ar), 130.9 (Ar), 130.2 (Ar), 129.87 (Ar), 129.85 (Ar), 129.2 (Ar), 127.84 (Ar), 127.78 (Ar), 113.8 (Ar), 113.6 (Ar), 110.2 ($C(CH_3)_2$), 101.2 (*C*-1), 95.9 (*C*-1′, $J_{C\text{-}H}$ = 173.0 Hz), 80.2 (*C*-3), 79.3 (*C*-3), 77.5 (*C*-4′), 76.0 (*C*-2′), 75.0 (*C*-2), 74.4 ($ArCH_2O$), 73.53 (*C*-5), 73.49 (*C*-4), 73.1 ($ArCH_2O$), 72.7 ($ArCH_2O$), 67.2 ($OCH_2CH_2N_3$), 66.3 (*C*-5′), 62.8 (*C*-6), 55.41 ($OCH_3$), 55.40 ($OCH_3$), 55.36 ($OCH_3$), 51.1 ($OCH_2CH_2N_3$), 28.1 ($CH_3C$), 26.9 ($SiC(CH_3)_3$), 26.7 ($CH_3C$), 19.3 ($SiC(CH_3)_3$), 16.6 (C-6′); HR ESIMS: *m/z* $[M+NH_4]^+$ calcd for $C_{57}H_{75}N_4O_{13}Si$: 1051.5094. Found: 1051.5109. $[M+Na]^+$ calcd for $C_{57}H_{71}N_3NaO_{13}Si$: 1056.4648. Found: 1056.4649.

**2-Azidoethyl 2,3,4-tri-*O*-*p*-methoxybenzyl-α-L-fucopyranosyl-(1→2)-3,4-*O*-isopropylidene-β-D-galactopyranoside (S20)**. To a stirred solution of disaccharide **S19** (1.50 g, 1.45 mmol) in THF (50 mL) was added tetrabutylammonium fluoride (4.35 mL, 4.35 mmol, 1.0 M in THF) at room temperature. After stirrinng for 8 h, the solvent was evaporatted and the crude residue was purified by chromatography ($CH_2Cl_2$–$CH_3OH$, 50:1) to give **S20** (1.10 g, 95%) as a colorless syrup: $R_f$ = 0.54 ($CH_2Cl_2$–$CH_3OH$, 30:1); $[\alpha]_D^{25}$ = –71.4 (*c* 0.7, $CH_2Cl_2$); $^1$H NMR (500 MHz, $CDCl_3$, $\delta_H$) 7.33–732 (m, 4 H, Ar), 7.25–7.23 (m, 2 H, Ar), 6.90–6.88 (m, 2 H, Ar), 6.84–6.82 (m, 4 H, Ar), 5.45 (d, 1 H, *J* = 3.7 Hz, H-1′), 4.88 (d, 1 H, *J* = 11.4 Hz, $ArCH_2O$), 4.77 (d, 1 H, *J* = 11.3 Hz, $ArCH_2O$), 4.72 (d, 1 H, *J* = 12.2 Hz, $ArCH_2O$), 4.69 (d, 1 H, *J* = 12.2 Hz, $ArCH_2O$), 4.64 (d, 1 H, *J* = 11.3 Hz, $ArCH_2O$), 4.57 (d, 1 H, *J* = 11.4 Hz, $ArCH_2O$), 4.41 (d, 1 H, *J* = 8.3 Hz, H-1), 4.33 (dd, 1 H, *J* = 6.6, 6.0 Hz, H-3), 4.18 (qd, 1 H, *J* = 6.5, 2.0 Hz, H-5′), 4.13 (dd, 1 H, *J* = 6.0, 1.4 Hz, H-4), 4.03–3.95 (m, 3 H, H-2′, $OCH_2CH_2N_3$, H-6a), 3.91 (dd, 1 H, *J* = 10.2, 2.6 Hz, H-3′), 3.82–3.77 (m, 12 H, H-6b, $OCH_3$, $OCH_3$, $OCH_3$, H-5, H-2), 3.63 (d, 1 H, *J* = 2.0 Hz, H-4′), 3.59 (dt, 1 H, *J* = 11.2, 5.9 Hz, $OCH_2CH_2N_3$), 3.41–3.39 (m, 2 H, $OCH_2CH_2N_3$), 2.14 (br s, 1 H, 6-O*H*), 1.49 (s, 3 H, $CH_3C$), 1.35 (s, 3 H, $CH_3C$), 1.05 (d, 3 H, *J* = 6.5 Hz, H-6′); $^{13}$C NMR (175 MHz, $CDCl_3$, $\delta_C$) 159.3 (Ar), 159.24 (Ar), 159.18 (Ar), 131.4 (Ar), 131.1 (Ar), 130.9 (Ar), 130.2 (Ar), 129.8 (Ar), 129.2 (Ar), 113.85 (Ar), 113.84 (Ar), 113.6 (Ar), 110.7 ($C(CH_3)_2$), 101.2 (*C*-1), 95.9 (*C*-1′), 80.3 (*C*-3), 79.3 (*C*-3′), 77.4 (*C*-4′), 76.0 (*C*-2′), 74.8 (*C*-2), 74.4 ($ArCH_2O$), 74.3 (*C*-4), 73.4 (*C*-5), 73.1 ($ArCH_2O$), 72.8 ($ArCH_2O$), 67.4 ($OCH_2CH_2N_3$), 66.4 (*C*-5′), 62.7 (*C*-6), 55.42 ($OCH_3$), 55.40 ($OCH_3$), 55.37 ($OCH_3$), 51.1 ($OCH_2CH_2N_3$), 28.0 ($CH_3C$), 26.7 ($CH_3C$), 16.6 (*C*-6′); HR ESIMS: *m/z* $[M+NH_4]^+$ Calcd for $C_{41}H_{57}N_4O_{13}$: 813.3917. Found: 813.3934. $[M+Na]^+$ Calcd for $C_{41}H_{53}N_3NaO_{13}$ 818.3471. Found: 818.3469.

**2-Azidoethyl α-L-fucopyranosyl-(1→2)-β-D-galactopyranoside (DiN$_3$)**. Disaccharide **S20** (335 mg, 421 μmol) was dissolved in a solution of acetic acid (10 mL, 80% in water, v/v) and heated to 80 °C for 4 h. After cooling to room temperature, the reaction mixture was concentrated and co-concentrated with dry toluene three times. The residue was then purified by chromatography ($CH_2Cl_2$–$CH_3OH$, 2:1) and reverse-phase chromatography ($C_{18}$, $CH_3OH$–$H_2O$, 1:1) to give the **DiN$_3$** (120 mg, 72%) as a white foam: $R_f$ = 0.30 ($CH_2Cl_2$–$CH_3OH$, 2:1); $[\alpha]_D^{25}$ = –84.9 (*c* 0.8, $CH_3OH$); $^1$H NMR (500 MHz, $CD_3OD$, $\delta_H$) 5.21 (d, 1 H, *J* = 3.2 Hz, H-1′), 4.38 (d, 1 H, *J* = 7.3 Hz, H-1), 4.32 (qd, 1 H, *J* = 6.6, 2.1 Hz, H-5′), 4.02 (ddd, 1 H, *J* = 10.3, 6.2, 3.8 Hz, $OCH_2CH_2N_3$), 3.82 (d, 1 H, *J* = 2.0 Hz, H-4), 3.79–3.66 (m, 8 H, H-3′, H-2′, H-6a, H-6b, $OCH_2CH_2N_3$, H-2, H-3, H-4′), 3.54-3.49 (m, 2 H, H-5, $OCH_2CH_2N_3$), 3.44 (ddd, 1 H, *J* = 13.1, 7.1, 3.8 Hz, $OCH_2CH_2N_3$), 1.22 (d, 3 H, *J* = 6.6 Hz, H-6′); $^{13}$C NMR (125 MHz, $CD_3OD$, $\delta_C$) 103.6 (*C*-1), 101.6 (*C*-1′), 78.9, 76.7, 75.7, 73.8, 71.8, 70.6, 70.4, 68.8 ($OCH_2CH_2N_3$), 67.9, 62.5, 52.2 ($OCH_2CH_2N_3$), 16.7; HR ESIMS: *m/z* $[M+Na]^+$ Calcd for $C_{14}H_{25}N_3NaO_{10}$: 418.1432. Found: 418.1429.

**2-Azidoethyl α-L-fucopyranosyl-(1→2)-[2-acetamido-2-deoxy-α-D-galactopyranosyl-(1→3)]-β-D-galactopyranoside (TriAN$_3$)**. A solution of **DiN$_3$** (21.6 mg, 54.6 μmol), UDP-GlcNAc (53.4 mg, 81.9 μmol), UDP-Glc/Gal 4-epimerase (1.5 mL, 1.5 mg/mL, from *C. jejuni* strain NCTC 11168), blood group A transferase (2.0 μL, 2.5 mg/mL), and alkaline phosphatase (2 units) in a buffer (5×, 875 μL) (35 mM MOPS, pH 7.0, 20 mM $MnCl_2$ and 1 mg/mL BSA) in Milli-Q water (1.0 mL) was incubated at 37 °C. After 24 h, TLC ($CHCl_3$–$CH_3OH$–$H_2O$, 30:15:2) showed the disappearance of **DiN$_3$** and the formation of a slower moving product. After centrifugation, the supernatant was loaded onto the $C_{18}$ Sep-Pak cartridge using water as the eluent. The fractions containing the trisaccharide product were collected and lyophilized to give **TriAN$_3$** (31.4 mg, 96%) as a white solid: $R_f$ = 0.25 ($CHCl_3$–$CH_3OH$–$H_2O$, 30:15:2); $[\alpha]_D^{25}$ = +24.3 (*c* 0.4, $CH_3OH$); $^1$H NMR (700 MHz, $CD_3OD$, $\delta_H$) 5.30 (d, 1 H, *J* = 3.9 Hz, H-1′), 5.15 (d, 1 H, *J* = 3.6 Hz, H-1″), 4.52 (q, 1 H, *J* = 6.5 Hz), 4.42 (d, 1 H, *J* = 7.6 Hz, H-1), 4.33 (dd, 1 H, *J* = 11.0, 3.6 Hz), 4.19 (dd, 1 H, *J* = 7.1, 4.4 Hz), 4.12 (d, 1 H, *J* = 2.7 Hz), 4.05 (ddd, 1 H, *J* = 10.9, 5.7, 3.3 Hz), 3.98 (dd, 1 H, *J* = 9.7, 8.0 Hz), 3.93 (dd, 1 H, *J* = 9.8, 3.1 Hz), 3.90 (d, 1 H, *J* = 2.5 Hz), 3.82 (dd, 1 H, *J* = 10.9, 3.0 Hz), 3.80–3.68 (m, 8 H), 3.54 (ddd, 1 H, *J* = 13.4, 5.7, 3.2 Hz), 3.49 (t, 1 H, *J* = 6.0 Hz), 3.45 (ddd, 1 H, *J* = 13.4, 7.6, 3.3 Hz), 2.01 (s, 3H), 1.21 (d, 3 H, *J* = 6.5 Hz); $^{13}$C NMR (175 MHz, $CD_3OD$, $\delta_C$) 174.5, 103.4 (*C*-1, $J_{C\text{-}H}$ = 160.3 Hz), 100.1 (*C*-1′, $J_{C\text{-}H}$ = 174.0 Hz), 93.7 (*C*-1″, $J_{C\text{-}H}$ = 172.1 Hz), 78.0, 76.5, 73.9, 73.3, 72.7, 71.8, 70.6, 70.2, 70.0, 68.6, 67.7, 64.9, 63.4, 62.5, 52.1, 51.3, 22.7, 16.7; HR ESIMS: *m/z* $[M+Na]^+$ Calcd for $C_{22}H_{38}N_4NaO_{15}$: 621.2226. Found: 621.2229.

**2-Azidoethyl α-L-fucopyranosyl-(1→2)-[α-D-galactopyranosyl-(1→3)]-β-D-galactopyranoside (TriBN$_3$)**. A solution of **DiN$_3$** (33.4 mg, 84.5 μmol), UDP-Gal (67.1 mg, 109.9 μmol), blood group B transferase (1.0 mL, 2.5 mg/mL), alkaline phosphatase (2 units), and buffer (5×, 500 μL) (35 mM MOPS, pH 7.0, 20 mM $MnCl_2$ and 1 mg/mL BSA) in Milli-Q water (1.0 mL) was incubated at 37 °C, after 24 h, TLC ($CHCl_3$–$CH_3OH$–$H_2O$, 30:15:2) showed the disappearance of the disaccharide acceptor and the formation of a slower moving product. After centrifugation, the supernatant was loaded onto the $C_{18}$ Sep-Pak cartridge using water as the eluent. The fractions containing the trisaccharide product were collected and lyophilized to give **TriBN$_3$** (43.4 mg, 92%) as a white solid: $R_f$ = 0.23 ($CHCl_3$–$CH_3OH$–$H_2O$, 30:15:2); $[\alpha]_D^{25}$ = +11.2 (*c* 0.6, $CH_3OH$); $^1$H NMR (700 MHz, $CD_3OD$, $\delta_H$) 5.30 (d, 1 H, *J* = 1.8 Hz, H-1′), 5.15 (d, 1 H, *J* = 1.5 Hz, H-1″), 4.50 (q, 1 H, *J* = 6.2 Hz), 4.43 (d, 1 H, *J* = 6.6 Hz, H-1), 4.15–4.14 (m, 2 H), 4.05–

4.04 (m, 1 H), 3.98–3.90 (m, 3H), 3.85–3.69 (m, 10 H), 3.55–3.52 (m, 2 H), 3.46–3.43 (m, 1 H), 1.20 (d, 3 H, $J$ = 6.2 Hz); $^{13}$C NMR (125 MHz, $CD_3OD$, $\delta_C$) 103.4 ($C$-1″, $J_{C\text{-}H}$ = 169.5 Hz), 100.1 ($C$-1′, $J_{C\text{-}H}$ = 172.9 Hz), 96.1 ($C$-1, $J_{C\text{-}H}$ = 161.4 Hz), 79.9, 76.2, 73.9, 73.4, 73.1, 71.8, 71.4, 71.3, 70.04, 70.03, 68.5, 67.6, 65.6, 63.4, 62.5, 52.1, 16.6; HR ESIMS: $m/z$ $[M+Na]^+$ Calcd for $C_{20}H_{35}N_3NaO_{15}$: 580.1960. Found: 580.1958.

**3.6** *Synthesis of azido-peg-betagalactoside (**S24**)*

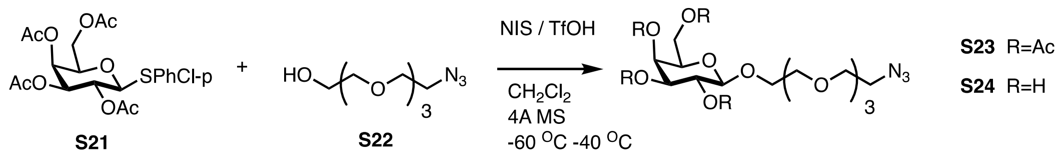

To a solution of thiogalactoside **S21** (150 mg, 0.32 mmol) and alcohol **S22** (50 mg, 0.23 mmol) in anhydrous $CH_2Cl_2$ (3.0 mL), we added 4Å molecular sieve (0.2 g), and the solution was cooled to -60 °C under argon and stirred for 1 h. NIS (120 mg, 0.54 mmol) was then added and the mixture was stirred for 10 minutes. Triflic acid (15 µL, 0.1 mmol) was added dropwise, and the reaction mixture was allowed to warm up slowly to -40°C and stirred for 30 min. $NEt_3$ (0.5 mL) was added to quench the reaction, and the reaction mixture was diluted with EtOAc (~30 mL), filtered off and extracted with a 3% $Na_2S_2O_3$ aqueous solution (~10 mL). The organic layer was dried over anhydrous $Na_2SO_4$, filtered and concentrated. The crude mixture was purified column chromatography on silica gel using a gradient of EtOAc–hexanes (30% → 50%), to yield the desired compound **S23** (63 mg, 50% yield). $^1$H NMR (400 MHz, $CDCl_3$) $\delta$ 5.38 (dd, $J$ = 3.4, 1.0 Hz, 1H, H-4), 5.20 (dd, $J$ = 10.5, 8.0 Hz, 1H, H-2), 5.01 (dd, $J$ = 10.5, 3.4 Hz, 1H, H-3), 4.57 (d, $J$ = 8.0 Hz, 1H, H-1), 4.17 (dd, $J$ = 6.6, 11.2 Hz, 1H, H-6a), 4.12 (dd, $J$ = 6.8, 11.2 Hz, 1H, H-6b), 3.95 (ddd, $J$ = 4.2, 4.6 11.0 Hz, 1H, OCHa), 3.91 (ddd, $J$ = 1.1, 5.6, 5.6 Hz, 1H, H-5), 3.75 (ddd, $J$ = 4.2, 6.8, 11.0 Hz, 1H, OCHb), 3.59 - 3.71 (m, 12H, 6 × $OCH_2$), 3.39 (t, $J$ = 5.3 Hz, 2H, $CH_2N_3$), 2.15 (s, 3H, OAc), 2.06 (s, 3H, OAc), 2.04 (s, 3H, OAc), 1.98 (s, 3H, OAc). Compound **3** (50 mg, 0.091 mmol) was dissolved in anhydrous MeOH (3.0 mL), and a solution of NaOMe in methanol (1.0 M, ~20 µL) was added. After stirring for 1 h, the mixture was neutralized with Amberlite IR-120 ($H^+$), and evaporated to afford the desired compound **S24** which was freeze-dried as a colorless white solid (34 mg, ~quantitative yield). $^1$H NMR (400 MHz, $CD_3OD$) $\delta$ 4.26 (d, $J$ = 7.5 Hz, 1H, H-1), 4.02 (ddd, $J$ = 10.2, 5.0, 3.0 Hz, 1H, OCHa), 3.83 (dd, $J$ = 3.2, ~1.0 Hz, 1H, H-4), 3.79 – 3.64 (m, 15H, 6 × $OCH_2$ + H-5 + H-6a + H-6b), 3.56 – 3.49 (m, 2H, H-2 + OCHb), 3.47 (dd, $J$ = 9.7, 3.3 Hz, 1H, H-3), 3.39 (t, $J$ = 5.0 Hz, 2H, $CH_2N_3$).

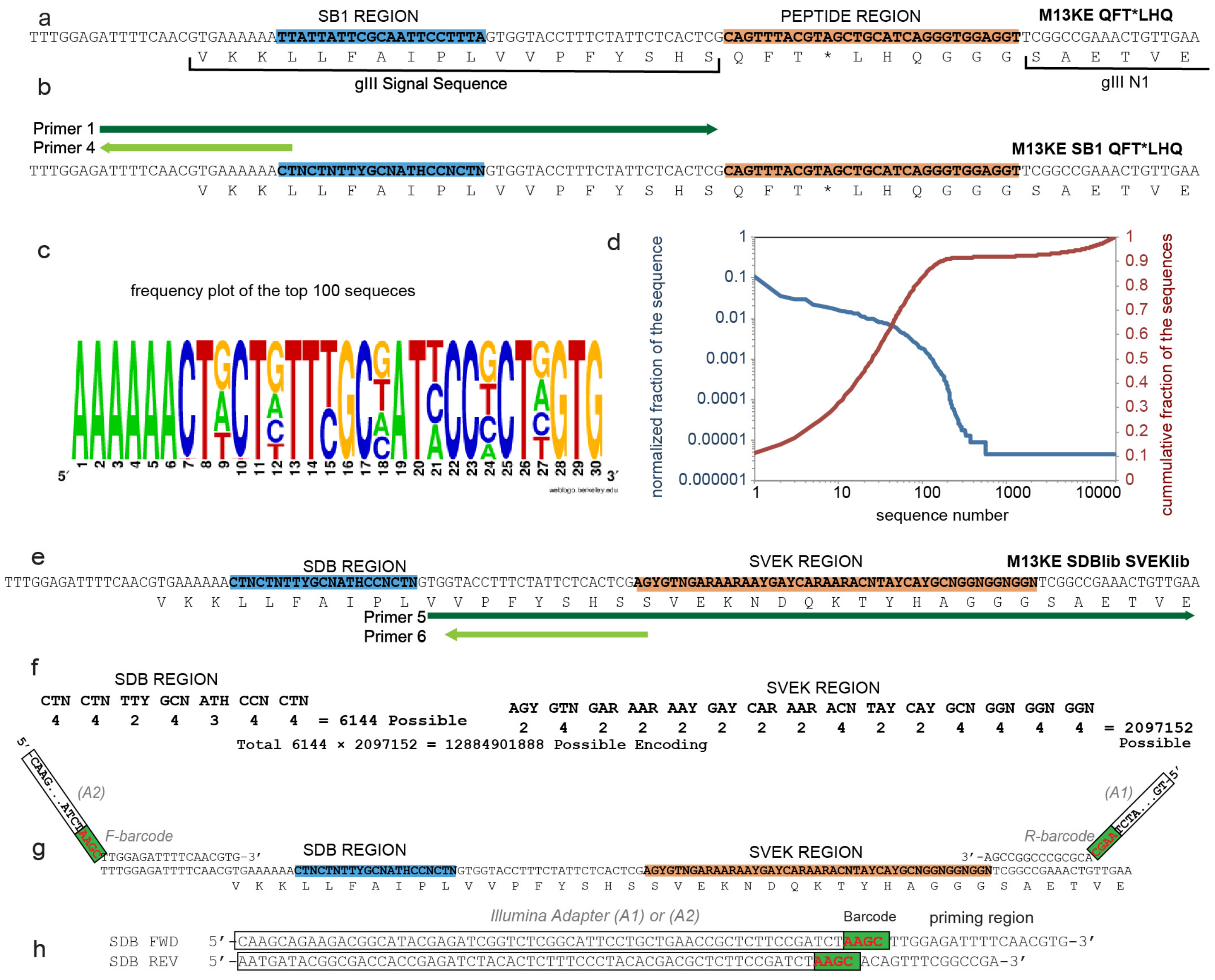

**Fig. S1**. Cloning of silent double barcode (SDB) regions.
**a**, Location of SDB and Peptide region on initial template. **b**, Primer 1 and 4 were used to convert M13KE QFT*LHQ to M13KE SDBlib QFT*LHQ. **c**, Frequency plot analysis of the top leader region of 100 unique sequences observed in deep sequencing of M13KE SDBlib vector; notably nucleotide C was completely suppressed in position 8 even though it was possible by design (CTN codon in Primer 1); **d**, Distribution and cumulative distribution of the unique sequences in deep sequencing of M13KE SDBlib vector. From 6144 possible sequences the most abundant ~200 unique sequences dominated 90% of the available diversity. **e**, Primer 5 and 6 were used to convert M13KE SDB QFT*LHQ to M13KE SDBlib SVEKlib. **f**, Summary of Degenerate sites in M13KE SDB QFT*LHQ. (G) Location of Illumina Sequencing Primers. **h**, Sequences of the Illumina sequencing primers.

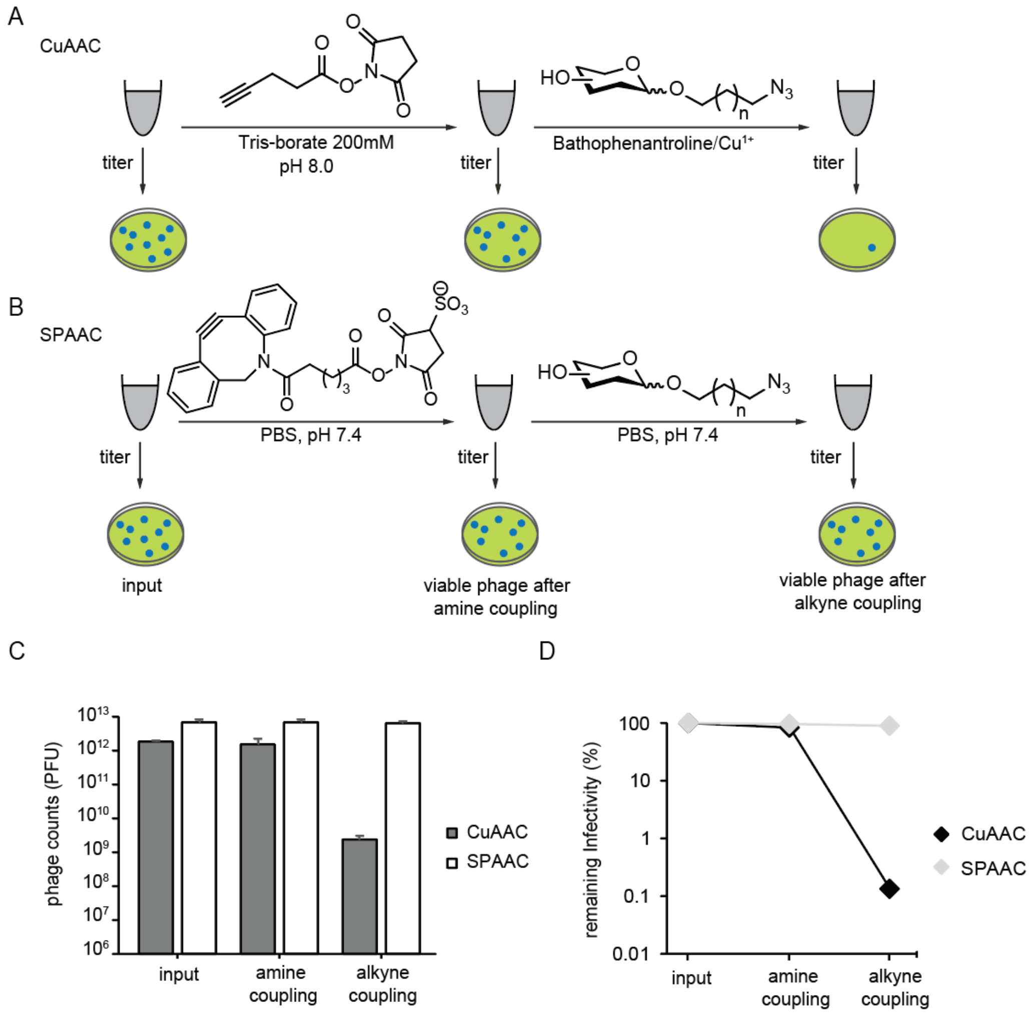

**Fig. S2**. Toxicity of the chemical glycosylation to phage.
**a**, General steps of the procedure to incorporate the glycans via CuAAC. **b**, Glycosylation of phages using copper-free (SPAAC) chemistry. **c**, Phage titers as total counts of plaque-forming units (PFU) from chemical glycosylation experiments, n=3. **d**, Phage viability results represented as percentage of remaining infectivity observed after first and second coupling steps amine and alkyne respectively, relative to the unmodified phage mixture (input).

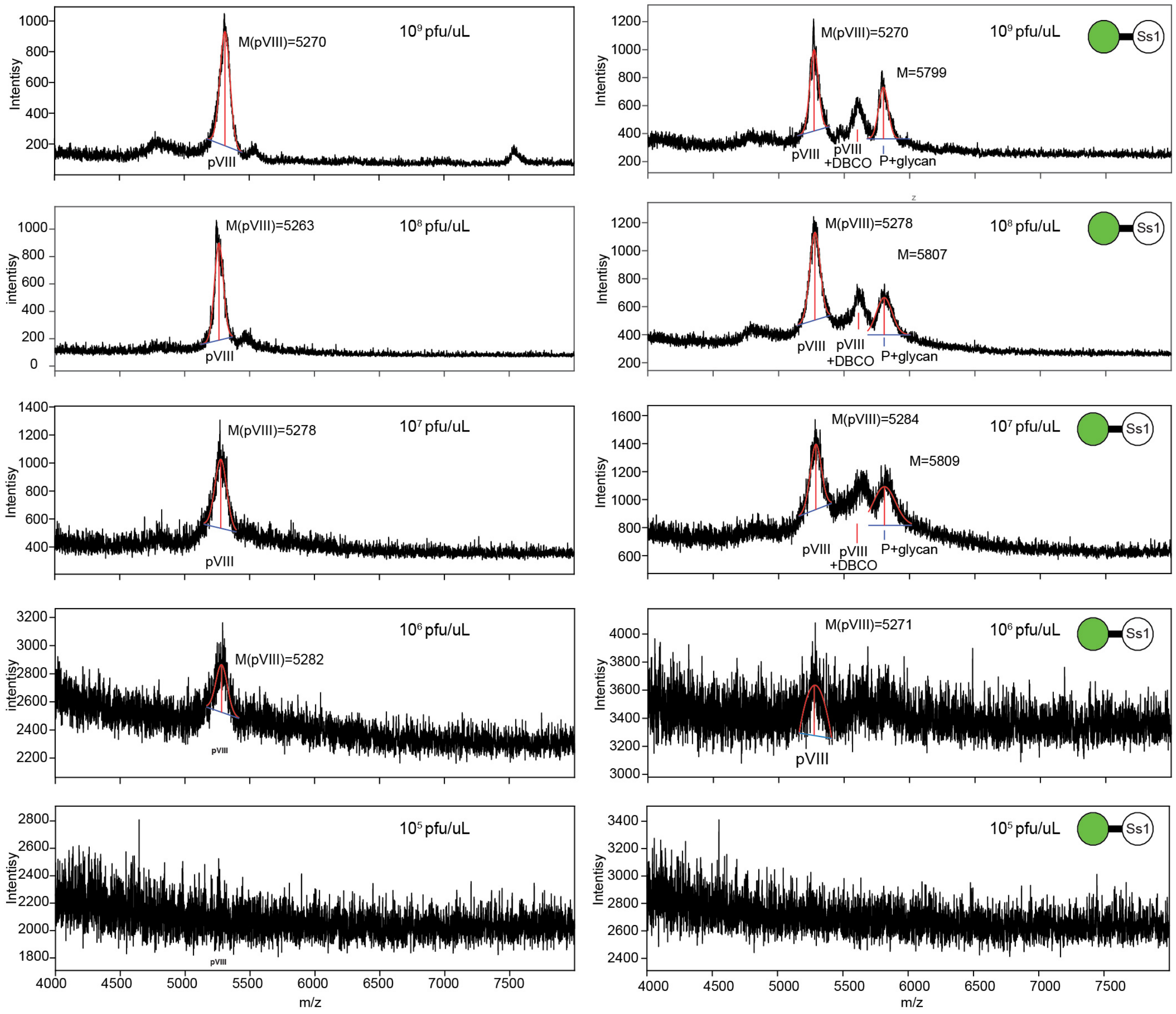

**Fig. S3**. MALDI spectra of phage at different concentrations.
Spectra were acquired as described in section "*1.7 Analysis of glycosylation of phage samples by MALDI-TOF MS*" using indicated concentrations of unmodified phage (left) or (right) phage that was partially acylated by DBCO-NHS and then partially modified beta-thiomethylene-mannoside (main text Fig. 1d).

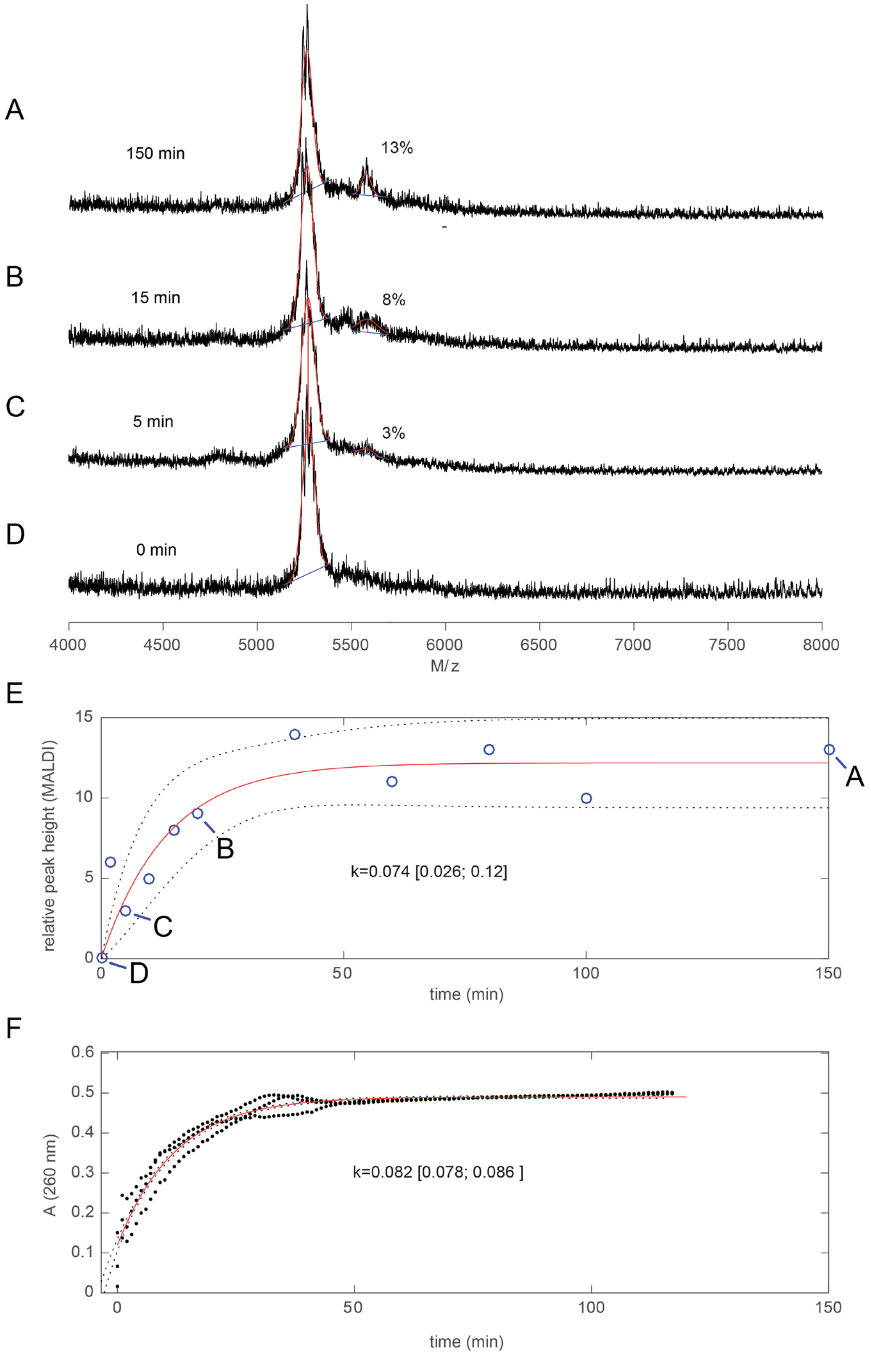

**Fig. S4**. Monitoring of the acylation of phage particles by MALDI.
**a-d**, Phage was exposed to DBCO-NHS for indicated time. The reaction was quenched and analyzed by MALDI. **e**, The kinetic profile of the reaction, rate constant ($M^{-1}$ $s^{-1}$) and 95% bounds of the fit. **f**, Hydrolysis of DBCO-NHS in the same buffer monitored by change in UV absorbance of DBCO-NHS solution indicated that hydrolysis is also completed at ~50 minutes. Similarity of the kinetic profiles of the acylation and hydrolysis reaction suggests that the fraction of the acylated N-termini is limited by the rate of the hydrolysis of DBCO-NHS. In such conditions, the best practice for controlling the number of acylated sites is to vary the concentration of DBCO-NHS but maintain the time of the reaction constant (>1 hour), allowing consumptions of all DBCO-NHS either due to reaction with N-terminus or due to hydrolysis.

a

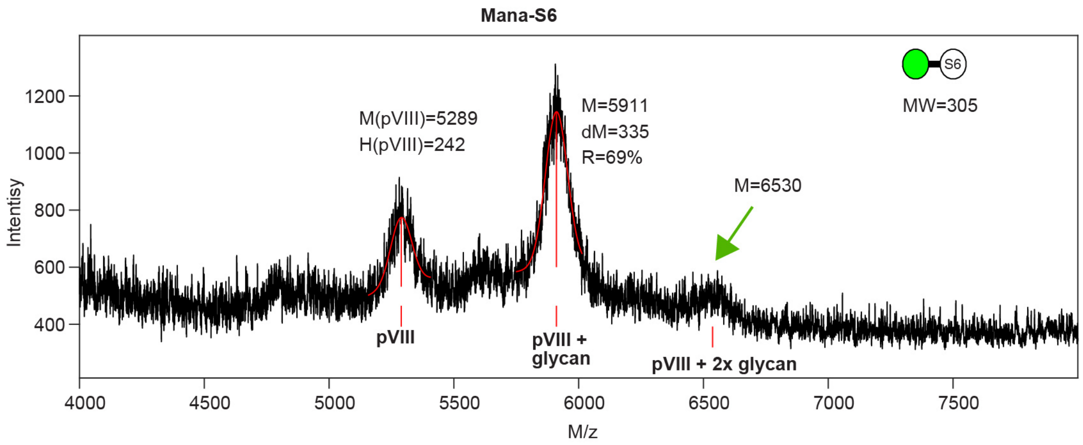

b

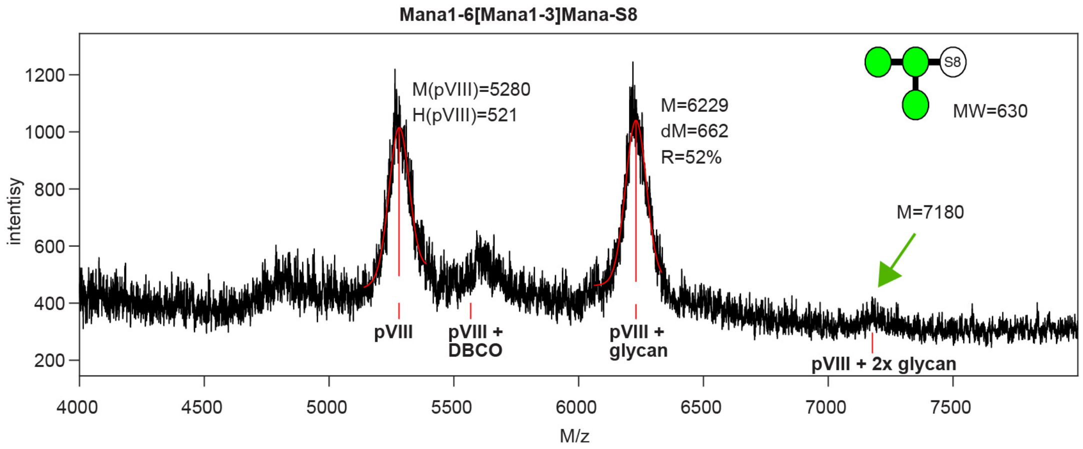

**Fig. S5.** MALDI-TOF analysis of double-modified pVIII protein.
Spectra of the phage particles in which over 50% of the pVIII residues have been modified with (**a**) αMan-S6 (Manα-$(CH_2)_6$-) or (**b**) $Man_3$-S6 (Manα1-6[Manα1-3]Manα-$(CH_2)_6$-) indicates presence of ~5% of species that corresponds to the M/z of pVIII protein modified by two glycans (presumably on the N-terminus and Lys8 of pVIII).

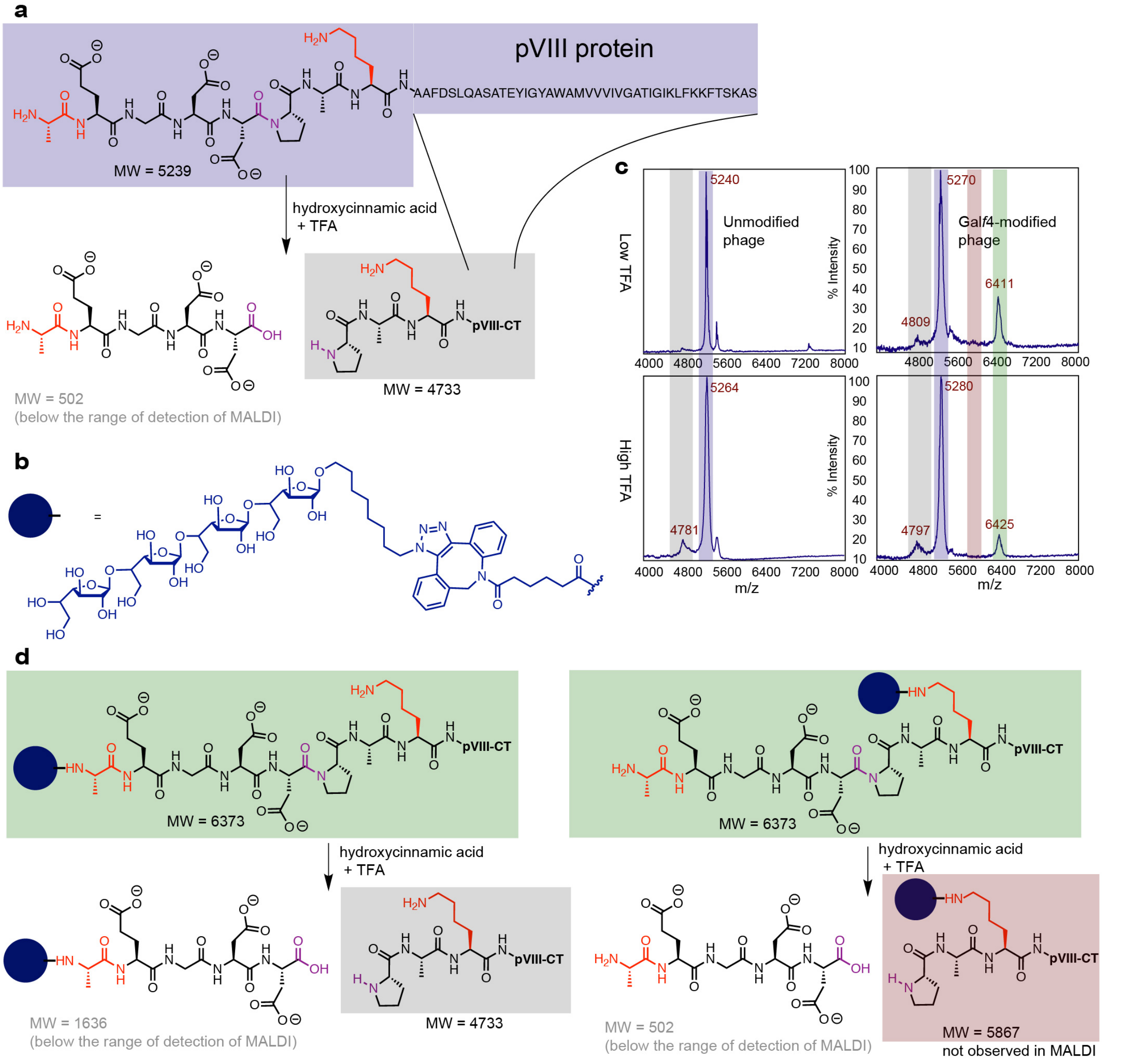

**Fig. S6**. Regioselectivity of the modification of pVIII protein.
**a**, Scheme of acid induced cleavage of Asp-5-Pro-6 (D5-P6)[3] bond in pVIII to yield pVIII[1-5] and pVIII[6-55] fragments. **b**, Scheme of the modification of the phage clone with the azido-glycan **Gal*f*4** (Galfβ1-5Galfβ1-5Galfβ1-5Galfβ-($CH_2$)-$N_3$). Modified phage was purified by Zeba column to remove unreacted glycan. **c**, MALDI-TOF analysis of modified and unmodified phage clones after incubation in 70% (v/v) TFA for 1 h ("High TFA") or without any pre-treatment with TFA ("Low TFA"). Analysis detects unmodified pVIII[6-55] fragment but does not detect presence of glycosylated pVIII[6-55] fragment. Glycosylation, thus, occurs preferentially at the N-terminus residue. **d**, Scheme of the cleavage of the pVIII modified with the azido-glycan **Gal*f*4** on the N-terminus and ε-amine of Lys-8 residue.

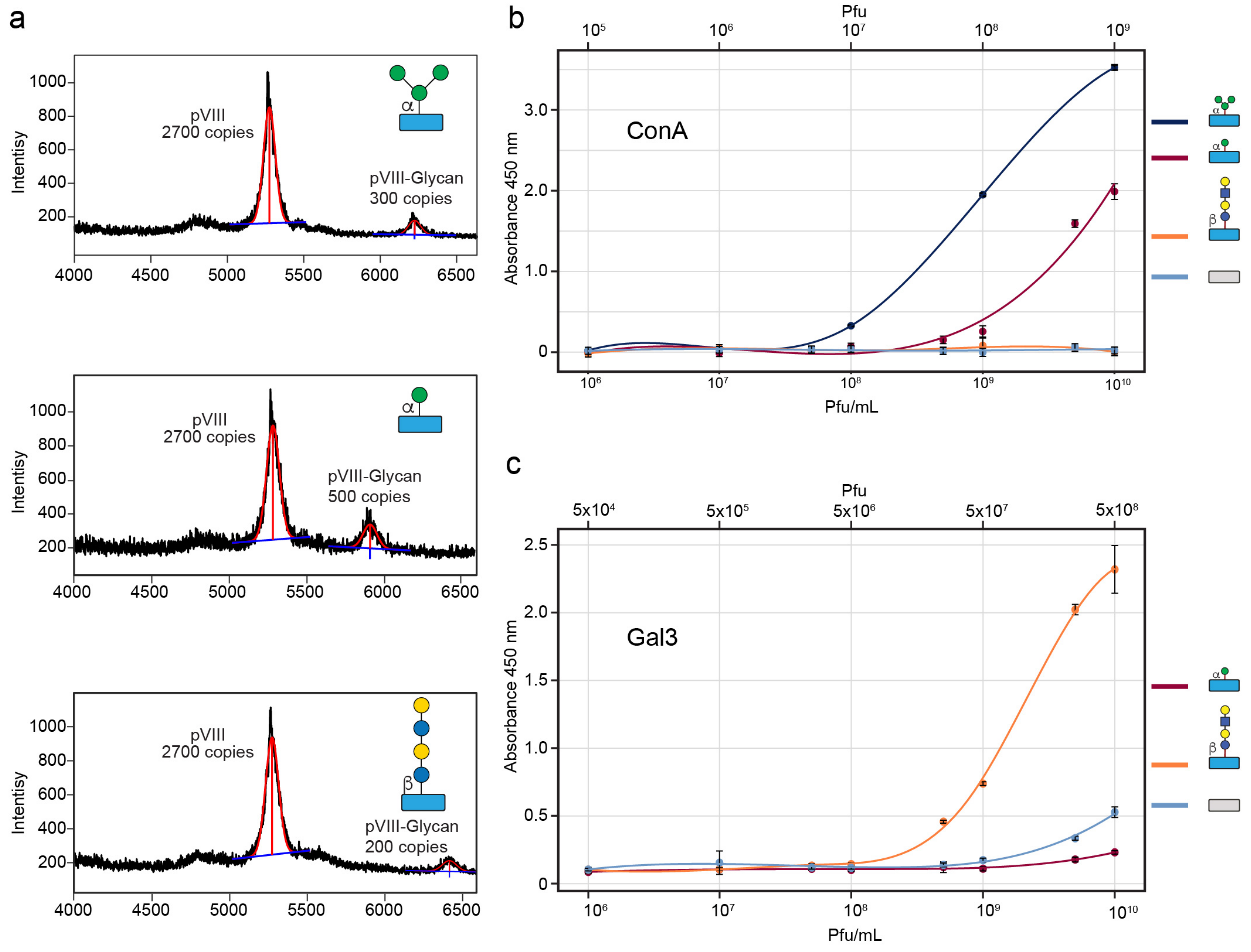

**Fig. S7.** ELISA measuring the binding of glycosylated phage to lectins.
**a**, MALDI characterization of the tested phage constructs. **b**, In binding to ConA-coated wells, phage displaying branched trimannoside (Manα1-6[Manα1-3]Manα-s6) exhibited the highest signal, the clone displaying a similar copy number of Manα glycan exhibited a 10x lower binding, while phage conjugated with LNT and unconjugated blank phage showed no detectable binding. **c**, In binding to Galectin 3 (Gal3)-coated wells, the phage modified with LNT produced the highest signal. Minor non-specific binding of unmodified phage was observed at $10^{10}$ pfu/mL concentration. Interestingly, Manα-decorated clone showed a significantly lower non-specific binding to Gal3 at the same concentration. Data from (**b**-**c**) are represented as mean ± s.d. (n=3).

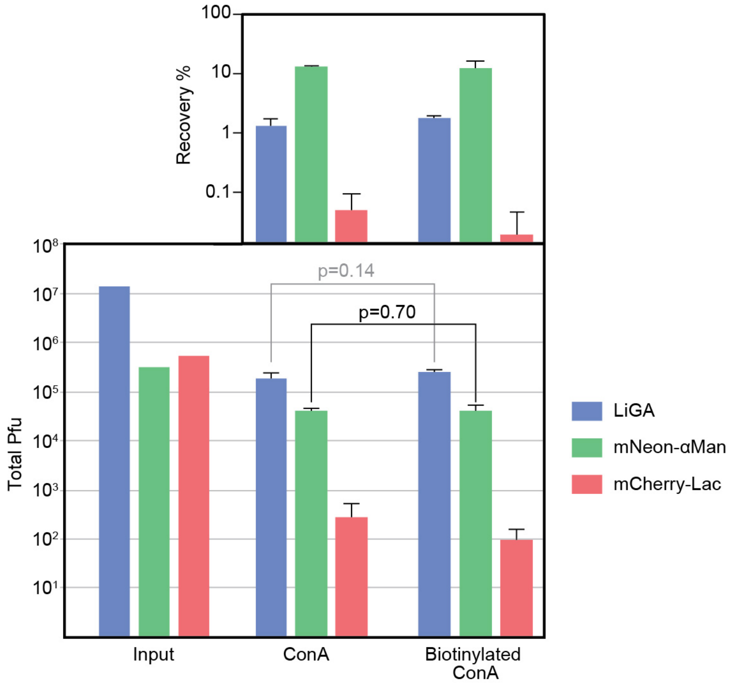

**Fig. S8**. Binding activity of ConA and Biotinylated-ConA.
Reporter phage controls can be used to test the effect of biotinylation on ability of ConA to recognize glycoslylated phage clones. The polystyrene wells were coated by similar concentrations of ConA or biotinylated-ConA. LiGA that contained reporter phage were added to the wells, incubated for 1 hour, rinsed 3x with PBS, eluted with HCl pH 2.0 and titered using PFU-assay. Phage transducing mNeon (modified with αMan) showed statistically indistinguishable recoveries of 13.1% and 12.3% on ConA or biotinylated-ConA respectively as calculated by Student's t-test. PFU data is represented as mean (input, n=2) or mean + s.d. (output, n=4). Recovery data represented as mean + s.d. propagated from s.d. of input and output PFU.

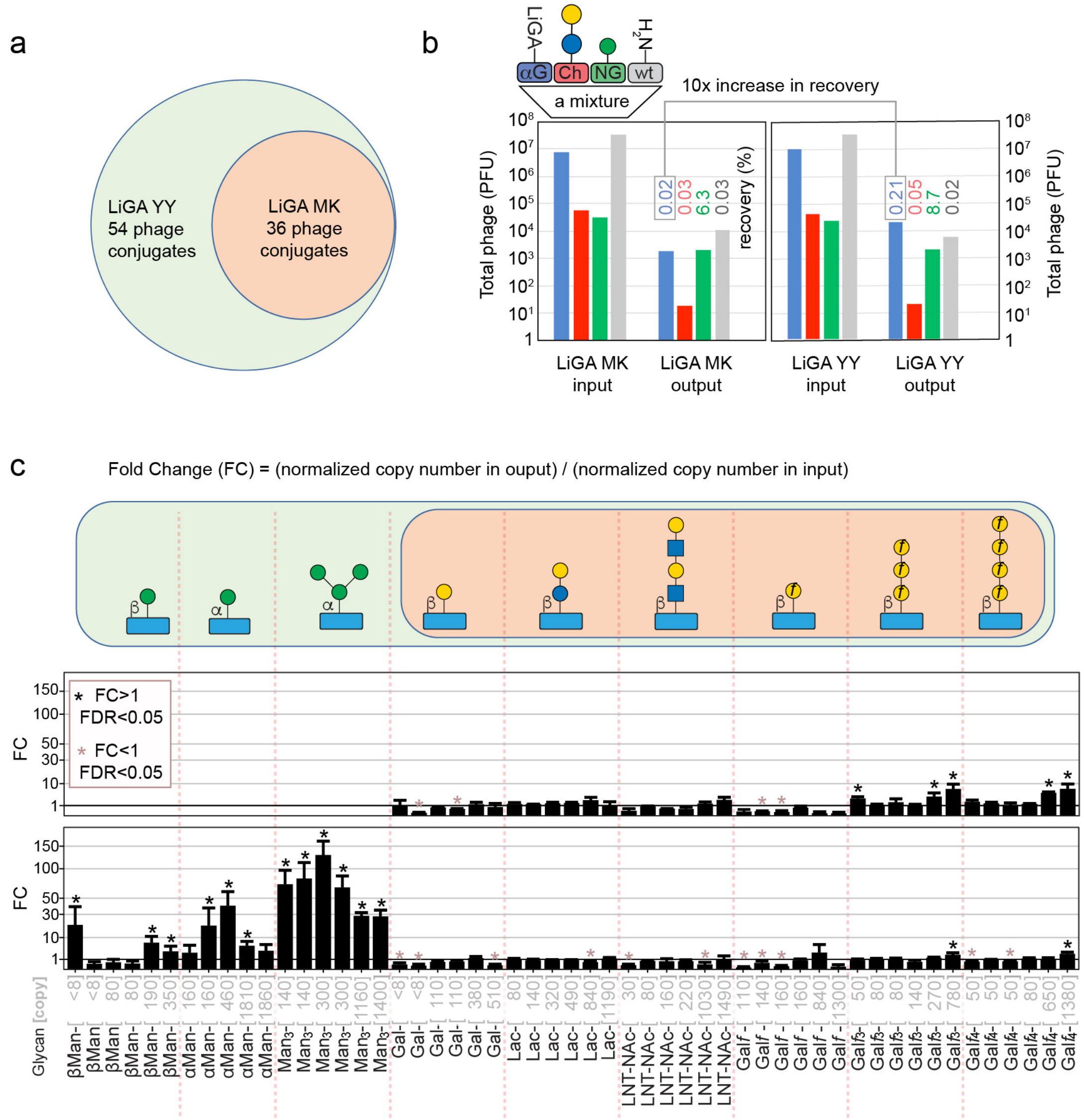

**Fig. S9**. Binding of LiGA with and without ConA-binding clones to ConA.
**a**, Venn diagram of LiGA YY with putative ConA-binding glycosylated phage and LiGA YM without these ConA-binding glycosylated phage. **b**, PFU assay determine that recovery of LiGA YY is 10 times higher than recovery of LiGA YM. Results are an average of two experiments and significance cannot be reliably determined. **c**, Deep sequencing of YM and YY datasets after the selection. Fold change (FC) values were determined using as differential enrichment (DE) analysis using copy numbers in the output and input LiGA mixtures. Error bars describe a propagated dispersion of the input and output reads. Stars denote significantly enriched (black) and depleted (pink) reads in DE-analysis with FDR≤0.05. n=2 top, n=4 bottom.

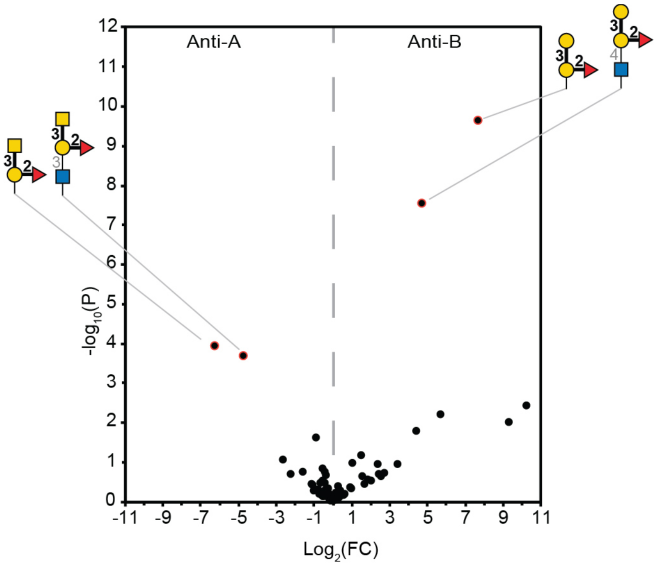

**Fig. S10**: Binding of LiGA to anti-A and anti-B antibodies.
Murine monoclonal anti-A IgM and anti-B IgM (generous gift of Lori West, University of Alberta) were coated on polystyrene 96-well plate (5 and 6 replicates respectively) and LiGA YZ (**Table S3**) was added to the well. After 1 h incubation, the wells were rinsed 3 times with PBS 0.1% Tween before phage were eluted by treatment with HCl, pH 2. Eluted phage solution was PCR amplified and deep-sequenced as described in sections 1.14-1.15. The data was analyzed using EdgeR differential enrichment analysis (see section 2.1) using anti-A and anti-B as two sets. The volcano plot describes fold change (FC) difference and *p*-value of glycans enriched on anti-A (FC<0) and anti-B (FC>0) antibodies, n=4. Red circles denote four DNA barcodes that exhibited enrichment with FDR < 0.05. The structures of the glycans associated with these barcodes are known blood group glycans targeted by anti-A and anti-B antibodies.

Data files are available at:
Naïve YZ LiGA http://ligacloud.ca/searchLibInfo?f=0&b=0&d=20191210-87YZooPA-RR
Binding to Anti-A http://ligacloud.ca/searchLibInfo?f=0&b=0&d=20191210-87YZvgAW-RR
Binding to Anti-B http://ligacloud.ca/searchLibInfo?f=0&b=0&d=20191210-87YZndAW-RR

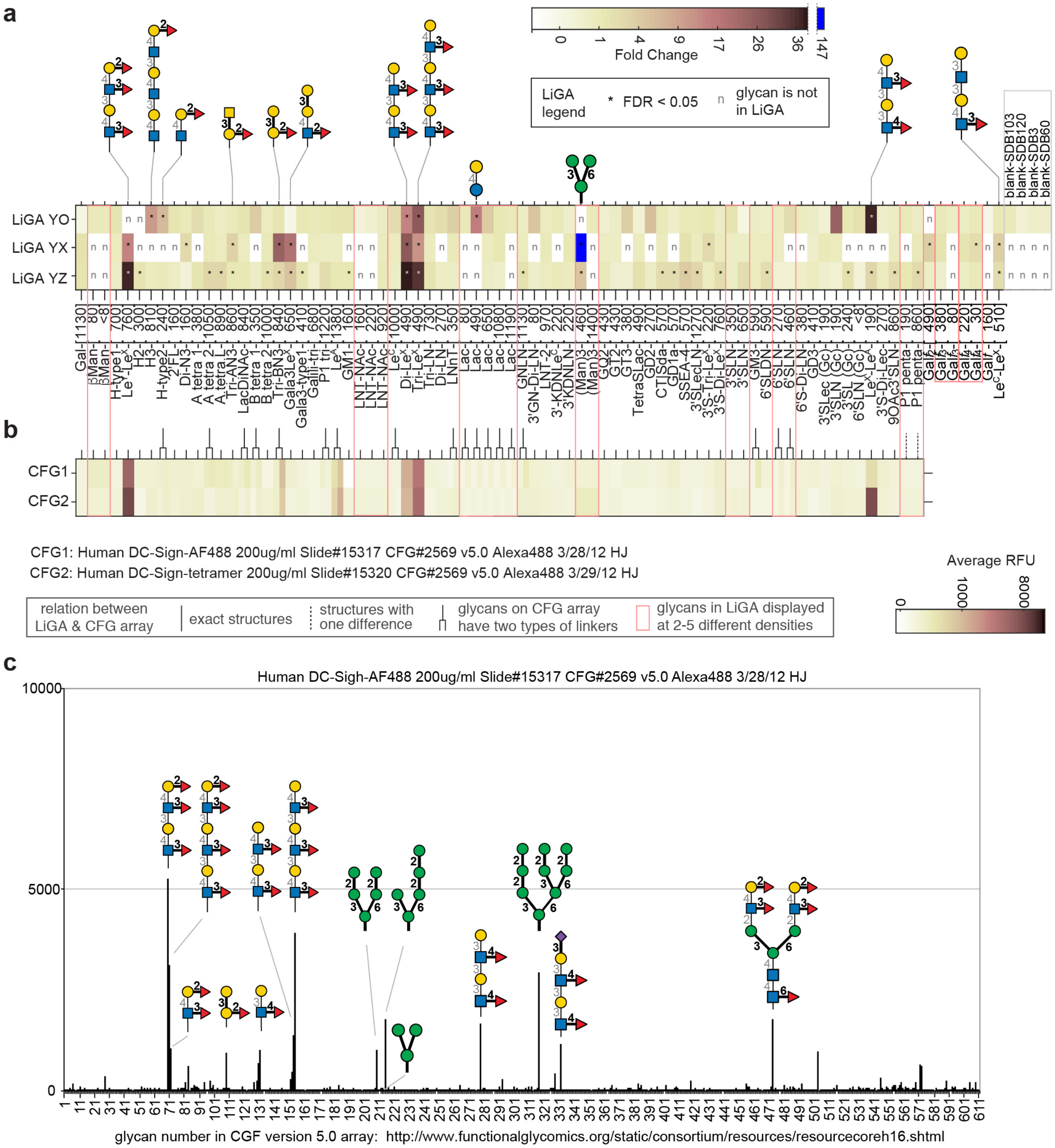

**Fig. S11**: Binding of DC-SIGN to LiGA and glass-based glycan array.

**a**, Binding of LiGA of three different compositions (YO, YX and YZ) to DC-SIGN on the surface of rat fibroblasts measured as differential enrichment of LiGA pulled down by DC-SIGN(+) fibroblasts and DC-SIGN(-) fibroblasts. LiGA YZ data in heat map is mirrored from bar chart in Fig. 5c for consistency. n=3,2,4 for YO, YX and YZ, respectively. **b**, Alignment to data describing binding of fluorescently labeled monomeric and tetrametic DC-SIGN protein to glass based glycan array (publically available CFG data under CFG request #2569 and searchable as "primscreen_5273" at: http://www.functionalglycomics.org/static/index.shtml). **c**, Overview of the full CFG array describing the binding of monomeric DC-SIGN protein and structures of the top binding glycans. *FDR<0.05. Bars represent an average response fluorescent unit (FRU, n=6).

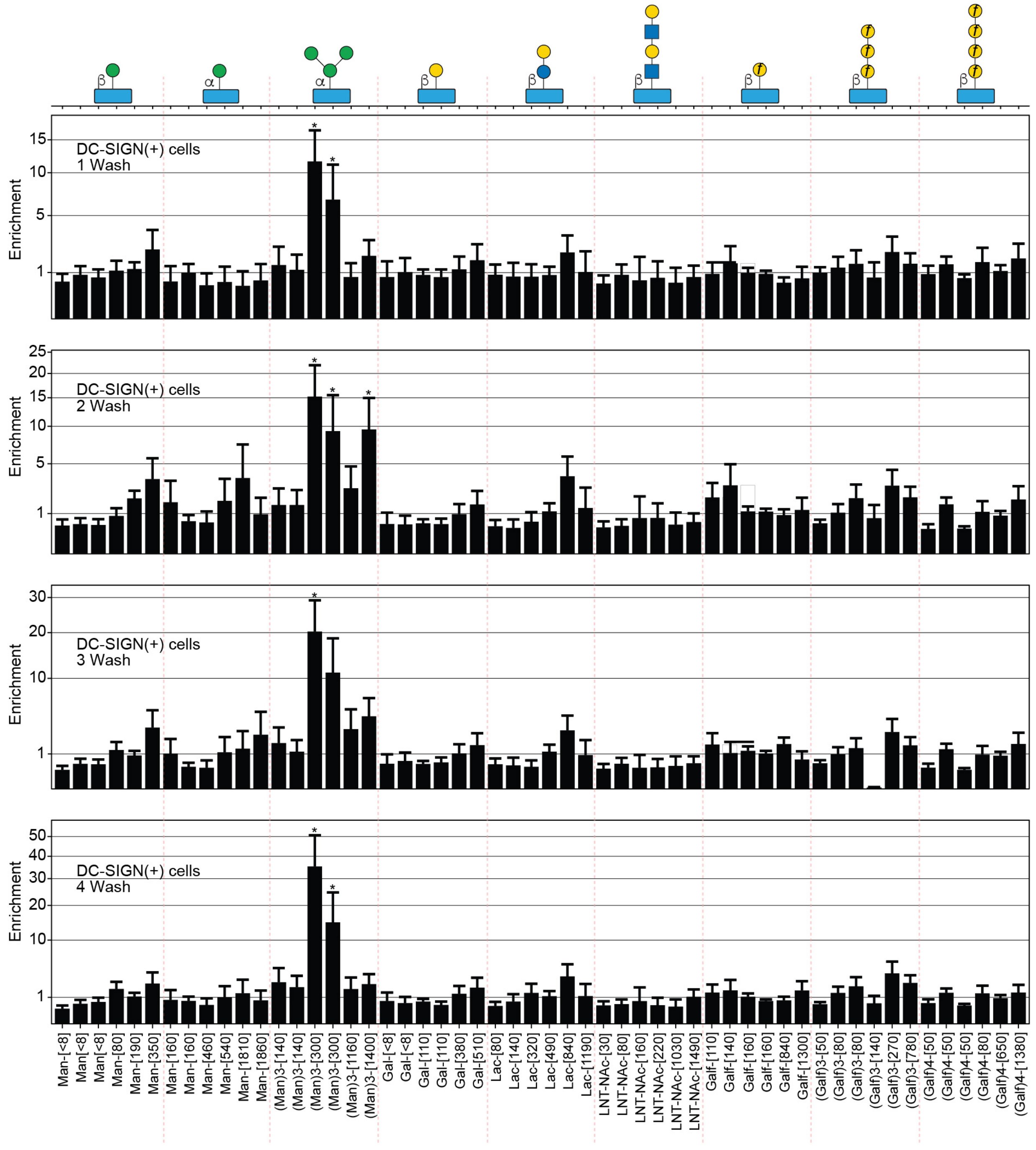

**Fig. S12**: Screening of LiGA 3x3 with DC-SIGN Cells.
Effect of washes on the binding of LiGA to cells. Enrichment was determined using as ratio of normalized average copy numbers in LiGA associated with DC-SIGN(+) and DC-SIGN(-) cells, n=3.

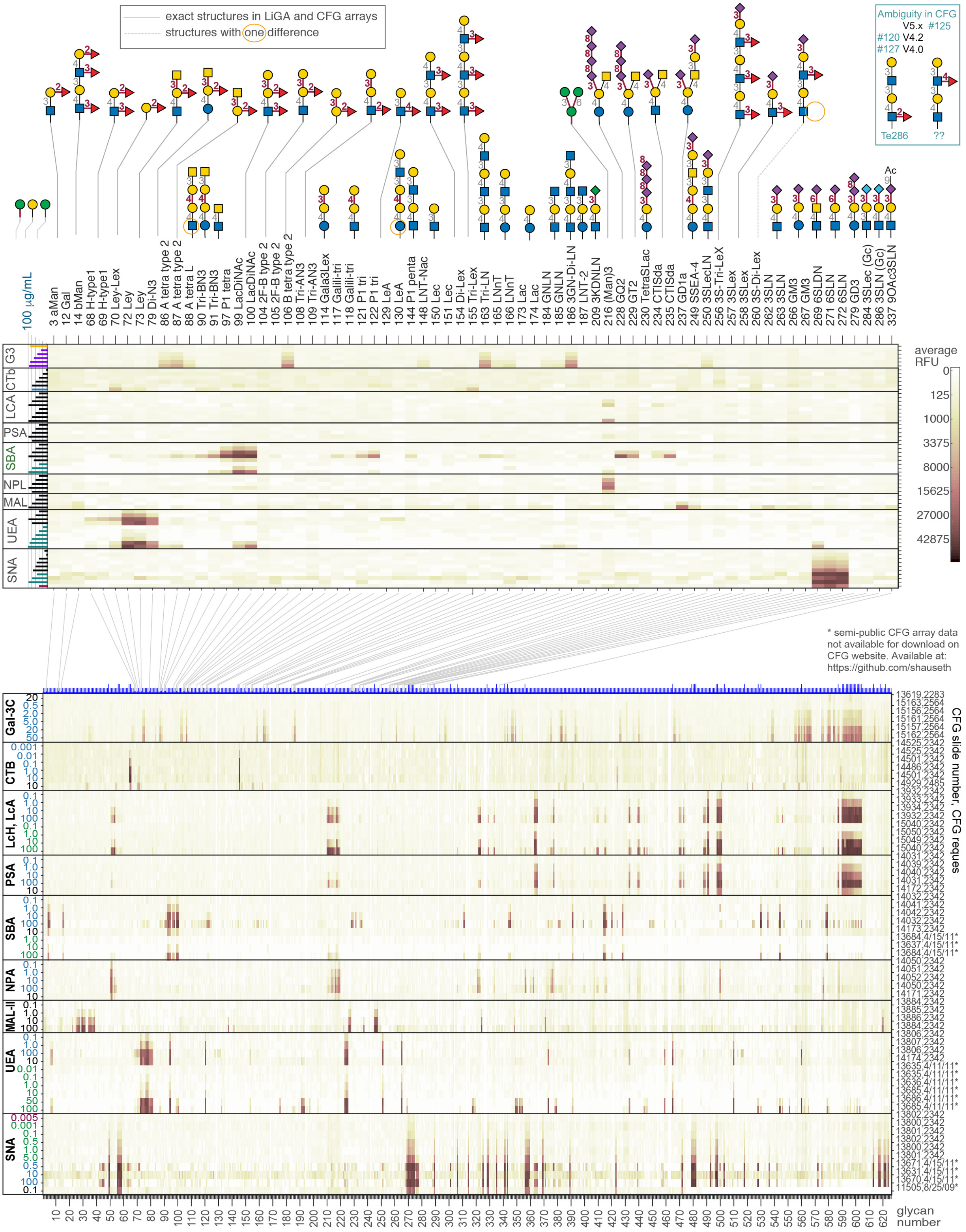
exact structures in LiGA and CFG arrays
structures with one difference
Ambiguity in CFG
V5.x #125
#120 V4.2
#127 V4.0
Te286
??
100 µg/mL
3 aMan
12 Gal
14 bMan
68 H-type1
69 H-type1
70 Ley-Lex
72 Ley
73 Ley
79 Di-N3
86 A tetra type 2
87 A tetra type 2
88 A tetra L
90 Tri-BN3
91 Tri-BN3
97 P1 tetra
99 LacDiNAc
100 LacDiNAc
104 2F-B type 2
105 2F-B type 2
106 B tetra type 2
108 Tri-AN3
109 Tri-AN3
114 Gala3Lex
117 Galili-tri
118 Galili-tri
121 P1 tri
122 P1 tri
129 LeA
130 LeA
144 P1 penta
148 LNT-Nac
150 Lec
151 Lec
154 Di-Lex
155 Tri-Lex
163 Tri-LN
165 LNnT
166 LNnT
173 Lac
174 Lac
184 GNLN
185 GNLN
186 3GN-Di-LN
187 LNT-2
209 3KDNLN
216 (Man)3
228 GQ2
229 GT2
230 TetraSLac
234 CTISda
235 CTISda
237 GD1a
249 SSEA-4
250 3SLecLN
256 3S-Tri-LeX
257 3SLex
258 3SLex
260 3S-Di-Lex
262 3SLN
263 3SLN
266 GM3
267 GM3
269 6SLDN
271 6SLN
272 6SLN
279 GD3
284 3SLec (Gc)
286 3SLN (Gc)
337 9OAc3SLN
G3
CTb
LCA
PSA
SBA
NPL
MAL
UEA
SNA
average RFU
0
125
1000
3375
8000
15625
27000
42875
* semi-public CFG array data
not available for download on
CFG website. Available at:
https://github.com/shauseth
Gal-3C
CTB
LcH, LcA
PSA
SBA
NPA
MAL-II
UEA
SNA
CFG slide number, CFG reques
glycan number

*(continued from the previous page)*

**Fig. S13**: Expanded view of the CFG data used in this manuscript

Top: Focused heatmap describing the summary of the binding of nice glycan binding protein to 65 glycans that were identical or nearly identical to the glycans used to construct LiGA. The data originates from 60 different glycan arrays (version 4.0, 4.1 and 5.0).

Bottom: Heat map describing 60 complete arrays composed of 626 glycans. The lines connecting the top and bottom heatmap describe the location of the 65 glycans in the list of 626 glycans, n=6.

Each lectin was tested at a few specific concentrations, which is displayed as a log-scale bar chart in (A) or as a number in (B). Heat map in (B), indexed heat map in (A) and the connecting lines were generated by MatLab script plotCFG2_allLectins.m and post-processed in Adobe Illustrator to fine tune the appearance of the axis labels. We calculated the cubic root of RFU data prior to applying the linear color scale to achieve the optimal display the data on heat map.

Each line of the heat map contains two references to the public CFG data (slide number and CFG request). Files marked with the * do not have a CFG request number but can be found by slide number and experiment date. All data was downloaded from CFG website using automated python script as described in section "*Access to CFG data*". Each dataset can be accessed manually via CFG website
http://www.functionalglycomics.org/glycomics/publicdata/primaryscreen.jsp
(i) make a selection (e.g., "plan lectins") or no selection; (ii) press "submit selections"; (ii) search for specific CFG request number on the page (e.g., in a browser window: Edit → Find, type "2342", press Enter)

For readers' convenience, the downloaded concatenated data for each glycan binding protein can be found in Data/CFG data in these files:

UEA_all.xlsx
SNL_all.xlsx
PSA_all.xlsx
LCA_all.xlsx
CTB_all.xlsx
ACG_all.xlsx
MAL_all.xlsx
NPL_all.xlsx
SBA_all.xlsx
Gal3_all.xlsx

### References

1. Bailey, J.J. & Bundle, D.R. Synthesis of high-mannose 1-thio glycans and their conjugation to protein. *Org. Biomol. Chem.* **12**, 2193-2213 (2014).
2. Lipinski, T., Kitov, P.I., Szpacenko, A., Paszkiewicz, E. & Bundle, D.R. Synthesis and immunogenicity of a glycopolymer conjugate. *Bioconjug. Chem.* **22**, 274-281 (2011).
3. Crimmins, D.L., Mische, S.M. & Denslow, N.D. Chemical Cleavage of Proteins in Solution. *Current Protocols in Protein Science* **41**, 11.14.11-11.14.11 (2005).
4. Ng, S. *et al.* Genetically encoded fragment-based discovery of glycopeptide ligands for carbohydrate-binding proteins. *J. Am. Chem. Soc.* **137**, 5248-5251 (2015).
5. Twibanire, J.D.K., Paul, N.K. & Grindley, T.B. Synthesis of novel types of polyester glycodendrimers as potential inhibitors of urinary tract infections. *New J. Chem.* **39**, 4115-4127 (2015).
6. Gaitonde, V. & Sucheck, S.J. Synthesis of beta-Glycosyl Amides from N-Glycosyl Dinitrobenzenesulfonamides. *J. Carbohydr. Chem.* **31**, 353-370 (2012).
7. Percec, V. *et al.* Modular Synthesis of Amphiphilic Janus Glycodendrimers and Their Self-Assembly into Glycodendrimersomes and Other Complex Architectures with Bioactivity to Biomedically Relevant Lectins. *J. Am. Chem. Soc.* **135**, 9055-9077 (2013).
8. Vasiliu, D. *et al.* Large-scale chemoenzymatic synthesis of blood group and tumor-associated poly-N-acetyllactosamine antigens. *Carbohydr. Res.* **341**, 1447-1457 (2006).
9. Blixt, O. *et al.* Chemoenzymatic synthesis of 2-azidoethyl-ganglio-oligosaccharides GD3, GT3, GM2, GD2, GT2, GM1, and GD1a. *Carbohydr. Res.* **340**, 1963-1972 (2005).
10. Zhang, P. *et al.* Unexpected structure of a C. difficile toxin A ligand necessitates an annotation correction in a popular screening library. *Chem. Commun. (Camb.)* **47**, 12397-12399 (2011).
11. Guo, Y. *et al.* Structural basis for distinct ligand-binding and targeting properties of the receptors DC-SIGN and DC-SIGNR. *Nat. Struct. Mol. Biol.* **11**, 591-598 (2004).
12. Lipinski, T. *et al.* A beta-mannan trisaccharide conjugate vaccine aids clearance of Candida albicans in immunocompromised rabbits. *Vaccine* **30**, 6263-6269 (2012).
13. Bundle, D.R. *et al.* Oligosaccharides and Peptide Displayed on an Amphiphilic Polymer Enable Solid Phase Assay of Hapten Specific Antibodies. *Bioconjug. Chem.* **25**, 685-697 (2014).
14. Bundle, D.R., Gidney, M.A., Kassam, N. & Rahman, A.F. Hybridomas specific for carbohydrates; synthetic human blood group antigens for the production, selection, and characterization of monoclonal typing reagents. *J. Immunol.* **129**, 678-672 (1982).
15. Cheng, K., Zhou, Y. & Neelamegham, S. DrawGlycan-SNFG: a robust tool to render glycans and glycopeptides with fragmentation information. *Glycobiology* **27**, 200-205 (2017).
16. Tsuchiya, S., Yamada, I. & Aoki-Kinoshita, K.F. GlycanFormatConverter: a conversion tool for translating the complexities of glycans. *Bioinformatics* **35**, 2434-2440 (2019).
17. Completo, G.C. & Lowary, T.L. Synthesis of galactofuranose-containing acceptor substrates for mycobacterial galactofuranosyltransferases. *J. Org. Chem.* **73**, 4513-4525 (2008).
18. Cao, B., White, J.M. & Williams, S.J. Synthesis of glycoconjugate fragments of mycobacterial phosphatidylinositol mannosides and lipomannan. *Beilstein J. Org. Chem.* **7**, 369-376 (2011).
19. Li, X.H., He, P., Liu, X.Y., Chao, R.B. & Wang, F.P. Synthesis and cardiac activity evaluation of the proposed structures of fuzinoside. *Tetrahedron* **71**, 8661-8668 (2015).

20. Daly, R., Vaz, G., Davies, A.M., Senge, M.O. & Scanlan, E.M. Synthesis and biological evaluation of a library of glycoporphyrin compounds. *Chemistry (Easton)* **18**, 14671-14679 (2012).
21. Fazio, F., Bryan, M.C., Blixt, O., Paulson, J.C. & Wong, C.H. Synthesis of sugar arrays in microtiter plate. *J. Am. Chem. Soc.* **124**, 14397-14402 (2002).
