## Supplementary material for "Genetically Encoded, Multivalent Liquid Glycan Array (LiGA)": MakeFigure3B.m

```
clear all;
close all;
%%%%%%%%%%%%%%%%%%%%%%%%%%%%%%%%%%%%%%%%%%%%%%%%%%%%%%%%%%%%%%%%%%%%%%%%%%%
% open a window of a perfect size: this is one option that might have
% to be changed on another computer with another screen resolution
%%%%%%%%%%%%%%%%%%%%%%%%%%%%%%%%%%%%%%%%%%%%%%%%%%%%%%%%%%%%%%%%%%%%%%%%%%%
fh = figure(1);
S = get(groot, 'ScreenSize');
set(fh, 'Position', [S(3)/2 S(4)/2 S(3)/2 S(4)/2])

VerticalScale=1.4; % vertical scaling factor for subplots
kink = 3.5;

drawnow;


% replace with another directory
coreD = pwd;
CFG_D = 'CFG data';
LiGA_D = 'LiGA data for figures';
D = fullfile(coreD, CFG_D);
D2 = fullfile(coreD, LiGA_D);

XLS = 'YG CFG PairingVerticallyAligned.xlsx';

F = {
    'CTB_all.xlsx';     [1:5 13];
    'LCA_all.xlsx';     0;
    'PSA_all.xlsx';     0;
    'SBA_all.xlsx';     0;
    'NPL_all.xlsx';     [2:5, 1];
    'MAL_all.xlsx';     13:16;       
    'UEA_all.xlsx';     [1:4, 9:14];
    'SNL_all.xlsx';     [1:6 8 7 9 16];
};

savename = 'AllLectins.pdf';


[~, XLS_names] = xlsread(fullfile(D,XLS));

FID = fopen(fullfile(D2, 'Figure3A_CTB-YG-knownGlycans.txt'));
disp(fgetl(FID));
TESTT= textscan(FID, ['%s' '%*[^\n\r]']);
fclose(FID);

HEAT_names = TESTT{1};

cHEAT_names = char(HEAT_names);
cHEAT_names(cHEAT_names=='.')=' ';
HEAT_names = cellstr(cHEAT_names);

YG_names_IUPAC  = XLS_names(2:end,4);
YG_names_common = XLS_names(2:end,3);
YG_names_alpha  = XLS_names(2:end,2);
YG_names_CFG    = XLS_names(2:end,6);

for i=1:numel(YG_names_IUPAC)
    YG_names_IUPAC{i} = strtrim(YG_names_IUPAC{i});
end


for i=1:numel(YG_names_common)
    parent = find(YG_names_common{i} == char(39) );
    
    if numel(parent) == 1
        parent = [];
    end
    notblanks = setdiff(1:numel(YG_names_common{i}), parent);
    YG_names_common{i} = YG_names_common{i}(notblanks);

    % remove front and back blanks
    YG_names_common{i} = strtrim(YG_names_common{i});

end

for i=1:numel(YG_names_alpha)
    blanks = find(YG_names_alpha{i} == ' ' );
    parent = find(YG_names_alpha{i} == char(39) );
    
    notblanks = setdiff(1:numel(YG_names_alpha{i}), union(blanks, parent));
    YG_names_alpha{i} = YG_names_alpha{i}(notblanks);
end

for i=1:numel(HEAT_names)
    HEAT_names{i} = strtrim ( HEAT_names{i});
end

uHEAT_names = HEAT_names;


for i=1:numel(YG_names_common)

    IX = regexpi(uHEAT_names,  [YG_names_common{i} '-[']);
    for j=1:numel(IX)
        if ~isempty(IX{j})
            
                disp([YG_names_common{i} ' was found in ' ...
                      uHEAT_names{j}]);
            
        end
    end
    
    
end

% for each name in the Heatmap, find 1,2 or more corresponding CFG names

temp1 = {};
temp2 = {};


for i=1:numel(uHEAT_names)
    
    IX = regexp(uHEAT_names{i}, '-[');
    
    searchFor = uHEAT_names{i}(1:IX(1)-1);
    IX = find(strcmpi( searchFor, YG_names_common));
    
    if isempty(IX)
        disp(['Something is wrong.... '  searchFor ]);
        return
    end
    
    for j=1:numel(IX)
        
        disp(['Heatmap entry ' uHEAT_names{i} ...
              ' maps to '  YG_names_common{IX(j)} ...
              ' and CFG entry ' YG_names_CFG{IX(j)}    ]);
          
        temp1 = [temp1; uHEAT_names(i) ...
                       YG_names_common(IX(j)) ...
                       YG_names_CFG(IX(j)) ];
                   
        temp2 = [temp2; uHEAT_names{i} ...
                       YG_names_common{IX(j)} ...
                       YG_names_CFG{IX(j)} ];            
        
    end
    
end

[~, IX] = unique(temp2);

IX = sort(IX, 'ascend');

temp1 = temp1(IX,:);

% go through the entries and count
MAX = 0;
temp3 = [];
XtickSparce  = [0];

for i=1:size(temp1,1)
    
    N = numel( find ( strcmp(temp1{i}, temp1(:,1)) ) );
    if N>MAX
        MAX=N;
    end
    
    if N==1
        % duplicate the entry
        temp3 = [temp3; temp1(i, :); temp1(i, :) ];
        XtickSparce  = [XtickSparce ; XtickSparce(end)+1.5;  XtickSparce(end)+1.51 ];
    elseif N==2
        temp3 = [temp3; temp1(i, :) ];
        XtickSparce = [XtickSparce ;  XtickSparce(end)+1 ];
    else
        disp('???');
    end
    
    
end    
XtickSparce  = XtickSparce(2:end);

for i=1:size(temp3,1)
    
    disp([uHEAT_names{ceil(i/2)} ' ' temp3{i,1} ' ' temp3{i,3} ]);
end


%%%%%%%%%%%%%%%%%%%%%%%%%%%%%%%%%%%%%%%%%%%%%%%%%%%%%%%%%%%%%%%%%%%%%%%%%%%
disp('******************************************************************');
disp('******************************************************************');
being_plotted = 1:numel(uHEAT_names);


DEBUG = 1; 

LigaIX = [];
Data = [];
Labels = {};


for i=1:2:numel(F)
    [temp1,temp2] = xlsread(fullfile(D,F{i}));
    
    CFGNamesBigTable = temp2(2:end,3);
    
    % extract glycan numbers from the first column
    GlycanNumber = temp1(:,1);
    % frame shift to exclude the three columns with names in them
    temp1 = temp1(:,5:end);
    
    tempLabel = temp2(1,4:end);
    
    NANS = nan(size(temp1,1),1);
    
    % if indices are defined for which columns to use; use them
    if F{i+1}

        Labels = [Labels tempLabel(F{i+1}) ];
        Data = [Data temp1(:,F{i+1}) ];
    else 
        % otherwise load all columns
        Labels = [Labels tempLabel];
        Data = [Data temp1];
    end

end

for i = 1:numel(CFGNamesBigTable)
    CFGNamesBigTable{i} = strtrim(CFGNamesBigTable{i});
end

CFGName = temp3(:,3);
CommonName = temp3(:,1);

for i = 1:numel(CFGName)
    CFGName{i} = strtrim(CFGName{i});
end

for i=1:numel(CFGName)
    
    IX = find(strcmp(CFGNamesBigTable, CFGName{i} ));
    if isempty(IX)
        disp([CFGName{i} ' no match' ]);
        LigaIX(i) = 0;
    elseif numel(IX)>1
        disp([CFGName{i} ' too many matches, taking the first']);
        LigaIX(i) = IX(1);
    else
        LigaIX(i) = IX(1);
    end
    
    % cleanup delimiter ' characters (ASCII 39) from the common names
    CommonNameU{i} = CommonName{i}(CommonName{i}~=char(39));
end

% eliminate the 0 entry
nonZero = find(LigaIX);
LigaIX = LigaIX(nonZero);
CommonNameU = CommonNameU(nonZero);

% sort the liga-CFG indices in the ascending order
[ LigaIX, IXX] = sort(LigaIX,'ascend');
CommonNameU = CommonNameU(IXX);

% define the labels for the focused X axis
XLabels = {};
for i=1:numel(LigaIX)
    XLabels{i} = [ num2str(LigaIX(i)) ' ' CommonNameU{i} ]; 
end;

% define color scheme and labels for Y-axes
bone2 = flipud(pink);
bone2(1,:) = [173 219 215]/255;
bone2(1,:) = [1 1 1];
%%%%%%%%%%%%%%%%

% remove data values that are small or negative
IX = find(Data<1);
IX2 = find(isnan(Data));
b=Data;
b(IX)=0; % use non zero value here if you want to use log scale

% transfom the data by cubic root to help visualizing data better
PlotData = b.^(1/3);

% make non-existent entry distincly different from others
PlotData(IX2) = -0.7;


%%%%%%%%%%%%%%%%%%%%%%%% draw a small CFG subplot
subplot(2,1,1);

imagesc(PlotData(LigaIX,:)', [ min(min(PlotData)) max(max(PlotData))*1.05 ]);

colormap(bone2);
colorbar

ax = gca;
ax.YTick = 1:numel(Labels);
if DEBUG
    ax.YTickLabel = Labels;
else 
    ax.YTickLabel = ''; 
end

ax.XAxisLocation = 'top';

ax.XTick = 1.5:2:numel(LigaIX);
ax.XTickLabel = XLabels(  1:2:numel(LigaIX)  );

ax.TickDir = 'out';
ax.FontName = 'arial';
ax.FontSize = 10;

xtickangle(ax,90)

 
ax_pos = ax.Position;  % position of first axes

%%%%%%%%%%%%%%%%% draw a large CFG plot
subplot(2,1,2);

imagesc(PlotData', [ min(min(PlotData)) max(max(PlotData))*1.05 ]);
colormap(bone2);
colorbar
drawnow

ax = gca;               % make a current axis object

ax1_pos = ax.Position;  % position of first axes

% draw a second X-axis on the large CFG heatmap
ax2 = axes('Position',ax1_pos,...
    'XAxisLocation','top',...
    'YAxisLocation','right',...
    'Color','none');

% draw a the clone of the original bottom-X and left Y axis
ax1 = axes('Position',ax1_pos,...
    'XAxisLocation','bottom',...
    'YAxisLocation','left',...
    'Color','none');

% define the left axes and bottom axes that range from 1 to 620
ax1.XLim = ax.XLim;
ax1.YLim = ax.YLim;
ax1.YTick = 1:numel(Labels);
ax1.FontName = 'arial';
ax1.FontSize = 10;

% use Y-labels only if debugging is necessary
if DEBUG
    ax1.YTickLabel = fliplr(Labels);
else 
    ax1.YTickLabel = ''; 
end

ax1.XTick = 10:10:size(PlotData,1);
ax1.TickDir = 'out';
xtickangle(ax1,90);

ax1.XAxis.MinorTick = 'on'; % Must turn on minor ticks if they are off
ax1.XAxis.MinorTickValues = 1:1:size(PlotData,1);

% define the righ axes and top axes that display only 65 selected ticks
ax2.XLim = ax.XLim;
ax2.YLim = ax.YLim;
ax2.YTick = 1:numel(Labels);
ax2.YTickLabel = ' ';
ax2.YAxis.Visible = 'on';
ax2.FontName = 'arial';
ax2.FontSize = 10;


ax2.XTick = unique(LigaIX);

ax2.XTickLabel = XLabels; % use for debugging to make sure labels match
%ax2.XTickLabel = ' ';

ax2.TickDir = 'out';
ax2.XTickMode = 'manual';
xtickangle(ax2,90);


% this part displays the ticks where most highly binding glycans are
% it also turns on the ticks for positions where LiGA overlap is
% you can have only one or the other. Toggle display_hits to change it. 

display_hits = 1;

% define the hits here

MaxData = max(PlotData,[],2);
[~,HITS] = sort(MaxData,'descend');

if display_hits 
    ax3 = ax2;
    
    % major ticks are in the location of the top 50 hits
    ax3.XTick = sort(HITS(1:50),'ascend');
    ax3.XTickLabel = ' ';
    ax3.XColor = [0 0 1];
    
    Minor = 1;
    if Minor
        
        ax3.XAxis.MinorTick = 'on'; % Must turn on minor ticks if they are off

        % values for minor ticks on top are the same as majot ticks on the
        % bottom
        ax3.XAxis.MinorTickValues = ax1.XAxis.MinorTickValues; 
    end
end
drawnow;
% hide the default axis
ax.XAxis.Visible = 'off';
ax.YAxis.Visible = 'off';


for i=1:numel(LigaIX)
    
    HorizontalScale = size(PlotData,1) / numel(LigaIX);
    
    if ~isempty(find(LigaIX(i)==HITS(1:50)))
        % clover green shade
        COLOR = [62 160 85]/255;
    else
        COLOR = 0.8*[1 1 1];
    end
    
    line([LigaIX(i) LigaIX(i)], numel(Labels)+[1 kink ],...
         'color', COLOR);
     
    line([LigaIX(i) HorizontalScale*i-kink],...
          numel(Labels)+[kink  VerticalScale*numel(Labels)],...
         'color', COLOR);
end

set(ax1,'Clipping','Off');

fh.PaperOrientation = 'landscape';
print(fh, '-dpdf', fullfile(D,savename));
```
