## Supplementary material for "Genetically Encoded, Multivalent Liquid Glycan Array (LiGA)": MakeFigureS11.m

```
clear all
DIR = fullfile(pwd, 'LiGA data for figures');

PWD = pwd;
DIR_CFG = 'CFG data';

% the name of the file that contains CFG data and colums for this figure
CFG_File = { 'DC_SIGN_all.xlsx';    [1 3]; };

savename = [CFG_File{1}(1:end-4) 'pdf'];

AllF = {
    'FigureS11_DCSIGNcells-YO 3rep-knownGlycans.txt'
    'FigureS11_DCSIGNcells-YX-knownGlycans.txt'
    'FigureS11_DCSIGNcells-YZ-knownGlycans.txt'
};

XLS = 'dict_table+CFG.xlsx';

[~,~,temp] = xlsread(fullfile(pwd, DIR_CFG, XLS));

axis_name = temp(:,3);

AllGycans = {};

% correct mismatches in names and small mimatches in density betweeen 
% different record files / tables
Rename = {
    'Tri-Man-[140].SDB18',  '(Man)3-[140].SDB18';
    '(Galf)2-[490].SDB131', '(Galf)2-[500].SDB131';
    'LNT-Nac-[160].SDB68',  'LNT-NAc-[160].SDB68';
    'LNT-Nac-[220].SDB69',  'LNT-NAc-[220].SDB69'; 
    'LNT-Nac-[920].SDB42',  'LNT-NAc-[920].SDB42';
    'Lac-peg4-[1080].SDB9', 'Lac-[1080].SDB9';
    'Ley-Lex-[800].SDB22',  'Ley-Lex-[760].SDB22'; 
    '3''SLex-[60].SDB65', '3''SLex-[80].SDB65'
    'none-[<8].SDB174', 'blank-[<8].SDB174';
    'Blank-[].SDB114', 'blank-[<8].SDB114';
    'Blank-[].SDB123', 'blank-[<8].SDB123';
    'Blank-[].SDB8', 'blank-[<8].SDB8';
    '(Galf)2-[500].SDB131', '(Galf)2-[490].SDB131';
    };

% low copy number sparce reads in test, control and naive samples
IGNORE = {
    '(Man)3-[0140].SDB18';
    'Ley-Di-Lex-[0320].SDB40'
    'Gal-[0110].SDB130';
    };

for j=1:numel(AllF)
    
    F=AllF{j};
    
    YLabel{j}=F(9:end-17);
    
fid = fopen(fullfile(DIR,F));

TITLE = fgetl(fid);
col = numel(regexp(TITLE, '\S+'));

FORM = '%s ';

for i=1:col-1
    FORM = [FORM '%f '];
end

AllVar = textscan(fid, FORM);
fclose(fid);

glycans{j} = AllVar{1};

% rename some mis-named glycans   
for m = 1:size(Rename,1)
    IX = find(strcmp(Rename{m,1}, glycans{j}));
    if ~isempty(IX)
        glycans{j}{IX} = Rename{m,2};
    end
end


Fr =[];
for i=2:col
    Fr = [Fr AllVar{i}];
end

FC{j} = (2.^(-Fr(:, end-2)));
Pval{j} = Fr(:, end-1);
Qval{j} = Fr(:, end-0);
IX = find(Qval{j}<0.05);

figure(100)
if j==1
    plot(log2(FC{j}), -log10(Pval{j}),'.k'); hold on;
    plot(log2(FC{j}(IX)), -log10(Pval{j}(IX)),'or');
end

% pad the numbers in glycan densities with zeros to improve sorting
for m = 1:numel(glycans{j})
    IX1 = find(glycans{j}{m}=='[');
    IX2 = find(glycans{j}{m}==']');
    insert = char ( '0'* ones(1, 5-(IX2-IX1) ) );
    TEMPglycans{j}{m} = [ glycans{j}{m}(1:IX1) insert glycans{j}{m}(IX1+1:end) ];
end

AllGycans = union(AllGycans, TEMPglycans{j});


end


clear IX

% remove the glycans that need to be ignored
for i=1:numel(IGNORE)
    if ~isempty(find( strcmp(IGNORE{i} , AllGycans) ))
        IX(i) = find( strcmp(IGNORE{i} , AllGycans) );
    else
        IX(i) = 0;
    end
end

keep = setdiff(1:numel(AllGycans), IX);
AllGycans = AllGycans(keep);
%%%%%%%%%%%%%%%%%%%%%%%


for i=1:numel(AllGycans)
    for j = 1:numel(TEMPglycans)
        
        IX = find( strcmp(AllGycans{i} , TEMPglycans{j}) );
        
        if ~isempty(IX)
            
            AllFC(i,j) = FC{j}(IX);
        else
            AllFC(i,j) = NaN;
        end
    end
end


% remove the numbers in glycan densities with zeros to allow searching

for i=1:numel(AllGycans)
    
    [IX1, IX2] = regexp(AllGycans{i},'[0+.','start', 'end');
    if ~isempty(IX1)
        AllGycans{i} = AllGycans{i}( [1:IX1 IX2:end] );
    end
    
    
    % is it in the comparrison table?
    IX = regexp(AllGycans{i},'].','start');
    if isempty(IX)
        disp(['something is wrong with ' AllGycans{i}]);
        return       
    end
    
    name = AllGycans{i}(1:IX(1));
    name(name=='.')=' ';
    
    
    IX2 = find ( strcmp(name,  axis_name));
    if isempty(IX2)
        disp(['Cannot find ' AllGycans{i}]); 
            CFGN(i,1) = -1; 
            CFGN(i,2) = -1; 
    else
        if isnumeric(temp{IX2(1),20}) 
            CFGN(i,1) = temp{IX2(1),20};
            if isnumeric(temp{IX2(1),21})
               CFGN(i,2) = temp{IX2(1),21}; 
            end
        elseif temp{IX2(1),20}(1)=='N'
            CFGN(i,1) = 0; 
            CFGN(i,2) = 0; 
        end
    end
end

found = find(CFGN(:,1)>0);
not_found = find(CFGN(:,1)<=0);

[~,IX1] = sort(CFGN(found,1));
[~,IX2] = sort(AllGycans(not_found));

order = [found(IX1); not_found(IX2)];

LigaIX = [];
for i =1:numel(found)
    LigaIX = [LigaIX; CFGN(order(i),:)'];
end

% plotting %%%%%%%%%%%%%%%%%%%%%%%%%%%%%%%%%%%%%%%%%%%%%%%%%%%%%%%%%%%%%%%

% trim values above the max value
% BE CAREFULL HERE; IT TRIMS THE DATA
MAX = 41; % this value works only for this data
AllFC(AllFC > MAX) = MAX;
% THERE SHOULD ONLY BE ONE DATA POINT

% use the order
PlotData = (AllFC(order,:).^(1/2))';
IX2 = find(isnan(PlotData));
PlotData(IX2) = -0.7;

fh = figure(1);
S = get(groot, 'ScreenSize');
set(fh, 'Position', [S(3)/2 S(4)/1.5 S(3)/2 S(4)/6]);

imagesc(PlotData)
ax = gca;
ax.XTickLabelRotation = 90;
ax.XTick = 1:numel(AllGycans);

% use the order
XLabels = AllGycans(order);
% trim the SDB
for i=1:numel(XLabels)
    IX = regexp(XLabels{i},'.SDB','start');
    IX2 = regexp(XLabels{i},'blank','end');
    if isempty(IX2)
        XLabels{i}=XLabels{i}(1:IX(1)-1);
    else
        XLabels{i}=XLabels{i}([1:IX2(1)+1, IX(1)+1:end]);
    end

end

% pad the numbers in glycan densities with 2x spaces to improve appearence
for m = 1:numel(XLabels)
    IX1 = find(XLabels{m}=='[');
    IX2 = find(XLabels{m}==']');
    if ~isempty(IX1)
        insert = char ( ' '* ones(1, 2*( 5-(IX2-IX1) )) );
        XLabels{m} = [ XLabels{m}(1:IX1) insert XLabels{m}(IX1+1:end) ];
    end
end

ax.XTickLabel = XLabels;
ax.FontSize = 8;
ax.TickDir = 'out';

ax.YTick = 1:numel(YLabel);
ax.YTickLabel = YLabel;

COLORMAP = flipud(pink);
COLORMAP(1,:) = [173 219 215]/255;
COLORMAP(1,:) = [1 1 1];
COLORMAP(end,:) = [0 0 1];

colormap(COLORMAP);
colorbar;

% add the stars where Q-values are <0.05
Qval_plot = [];

for kk = 1:size( PlotData,2 )
    
    for jj = 1:size(PlotData,1)
        
        %find the name of the glycan in the original dataset
        IX = find ( strcmp( AllGycans(order(kk)), glycans{jj}));
        if ~isempty(IX)          
            Qval_plot(kk,jj) = Qval{jj}(IX);
        else
            Qval_plot(kk,jj) = 2;
        end
    end
    
end

for kk = 1:size( Qval_plot,1 )    
    for jj = 1:size(Qval_plot,2)
        if Qval_plot(kk,jj)<0.05 & PlotData(jj,kk)>1
            text(kk,jj,'*', 'VerticalAlignment', 'middle',...
                  'HorizontalAlignment', 'center');
        elseif Qval_plot(kk,jj)==2
            text(kk,jj,'n', 'VerticalAlignment', 'middle',...
                  'HorizontalAlignment', 'center', 'color', 0.5*[1 1 1]);
        end
    end
end
            
            
  

figure(2);

colormap(hot);
bar(PlotData(1,:)', 'edgecolor', 'k');
ax = gca;
ax.XTickLabelRotation = 90;
% use the order
ax.XTick = 1:numel(AllGycans);
ax.XTickLabel = XLabels;

xlim([0 numel(AllGycans)+1]);
ax.FontSize = 8;
ax.TickDir = 'out';

CFG = temp(:,19);

fh = figure(3);
S = get(groot, 'ScreenSize');
set(fh, 'Position', [S(3)/2 S(4)/3 S(3)/2 S(4)/6]);
drawnow;
 
 %%%%%%%%%%%%%%%%%%%%%%%%%%%%%%%%%%%%%%%%%%%%%%%%%%%%%%%%%%%%%%%%%%%%%%%%%%%
disp('******************************************************************');
disp('******************************************************************');

uHEAT_names = AllGycans(order(1:numel(found)) );

being_plotted = 1:numel( uHEAT_names );

%set(fh, 'Position', [S(3)/2 S(4)/2 S(3)/2 S(4)/2])
VerticalScale=1.8;  % vertical scaling factor for subplots

DEBUG = 1; 

Data = [];
Labels = {};


for i=1:2:numel(CFG_File)
    [temp1,temp2] = xlsread(fullfile(PWD,'CFG data',CFG_File{i}));
    
    CFGNamesBigTable = temp2(2:end,3);
    
    % extract glycan numbers from the first column
    GlycanNumber = temp1(:,1);
    % frame shift to exclude the three columns with names in them
    temp1 = temp1(:,5:end);
    
    tempLabel = temp2(1,4:end);
    
    NANS = nan(size(temp1,1),1);
    
    % if indices are defined for which columns to use; use them
    if CFG_File{i+1}

        Labels = [Labels tempLabel(CFG_File{i+1}) ];
        Data = [Data temp1(:,CFG_File{i+1}) ];
    else 
        % otherwise load all columns
        Labels = [Labels tempLabel];
        Data = [Data temp1];
    end

end

for i = 1:numel(CFGNamesBigTable)
    CFGNamesBigTable{i} = strtrim(CFGNamesBigTable{i});
end

% remove data values that are small or negative
IX = find(Data<1);
IX2 = find(isnan(Data));
b=Data;
b(IX)=0; % use non zero value here if you want to use log scale

% transfom the data by cubic root to help visualizing data better
PlotData = b.^(1/3);

% make non-existent entry distincly different from others
PlotData(IX2) = -0.7;

%%%%%%%%%%%%%%%%%%%%%%%% draw a CFG subplot

imagesc(PlotData(LigaIX,:)', [ min(min(PlotData)) max(max(PlotData))*1.05 ]);


COLORMAP = flipud(pink);
COLORMAP(1,:) = [173 219 215]/255;
COLORMAP(1,:) = [1 1 1];

colormap(COLORMAP);
colorbar

ax = gca;
ax.YTick = 1:numel(Labels);
if DEBUG
    ax.YTickLabel = Labels;
else 
    ax.YTickLabel = ''; 
end

ax.XAxisLocation = 'top';


ax.XTick = 1.5:2:numel(LigaIX);
ax.XTickLabel = XLabels;

ax.TickDir = 'out';
ax.FontName = 'arial';
ax.FontSize = 10;

xtickangle(ax,90)
```
