## Supplementary material for "Genetically Encoded, Multivalent Liquid Glycan Array (LiGA)": Maldi.pdf

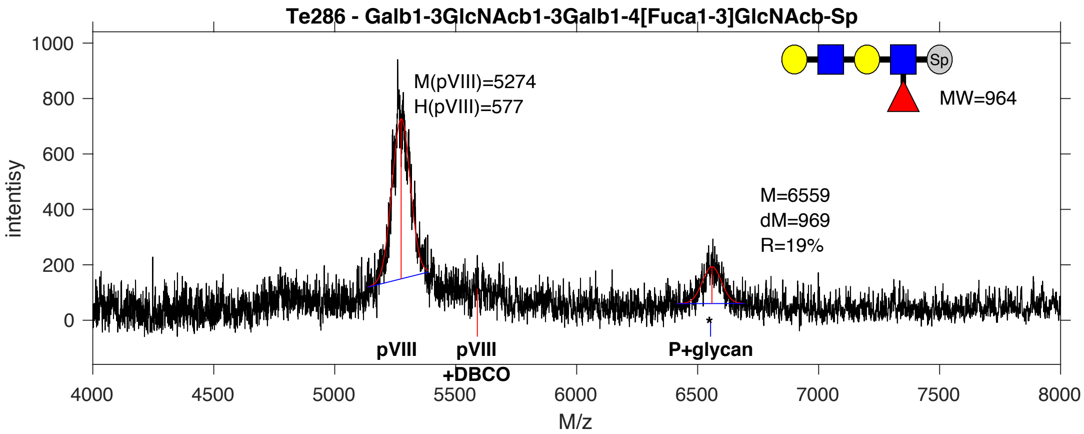


**SDB Number:** SDB1

**Sequencing File:** http://ligacloud.ca/searchLibInfo?f=0&b=0&d=20190703-87CLooOOKFFV-DF

**Barcode:**
CTGCTGTTCGCAATACCACTCAGTGTGGAGAAGAATGATCAGAAGACTTATCATGCGGGTGGAGGT

**Axis Name:** Lec-LeX-[510]

**IUPAC:** Gal(b1-3)GlcNac(b1-3)Gal(b1-4)[Fuc(a1-3)]GlcNac(b1-Sp

**CFG Name:** Galb1-3GlcNacb1-3Galb1-4[Fuca1-3]GlcNAcb-Sp

**Common Name:** Lec-LeX

**Glytoucan ID:** G69522TI

**Compound Number:** Te286

**Maldi File:** 1.txt

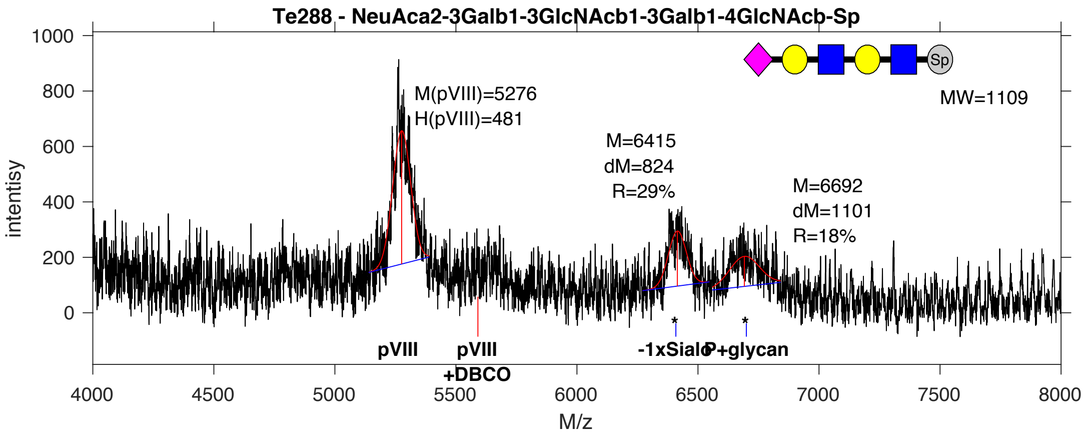


**SDB Number:** SDB2

**Sequencing File:** http://ligacloud.ca/searchLibInfo?f=0&b=0&d=20171128-87CLooOOCIWZ-SS

**Barcode:**
CTTCTATTCGCAATTCCGCTCAGTGTGGAGAAGAATGATCAGAAGACTTATCATGCGGGTGGAGGT

**Axis Name:** 3'SLecLN-[1270]

**IUPAC:** Neu5Ac(a2-3)Gal(b1-3)GlcNAc(b1-3)Gal(b1-4)GlcNAc(b1-Sp

**CFG Name:** Neu5Aca2-3Galb1-3GlcNAcb1-3Galb1-4GlcNAcb-Sp

**Common Name:** 3'SLecLN

**Glytoucan ID:** G78959US

**Compound Number:** Te288

**Maldi File:** 2.txt

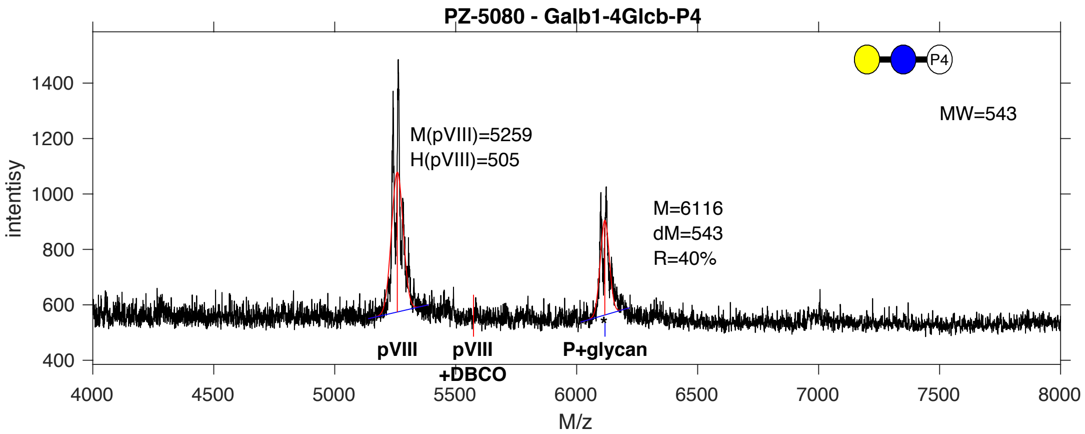


**SDB Number:** SDB9

**Sequencing File:** http://ligacloud.ca/searchLibInfo?f=0&b=0&d=20171128-87CLooOOKAQU-SS

**Barcode:**
CTTCTTTTTGCAATTCCTCTAAGTGTGGAGAAGAATGATCAGAAGACTTATCATGCGGGTGGAGGT

**Axis Name:** Lac-peg4-[1080]

**IUPAC:** Gal(b1-4)Glc(b1-p4

**CFG Name:** Galb1-4Glcb-P4

**Common Name:** Lac-peg4

**Glytoucan ID:** G94144EF

**Compound Number:** PZ-5080

**Maldi File:** 4.txt

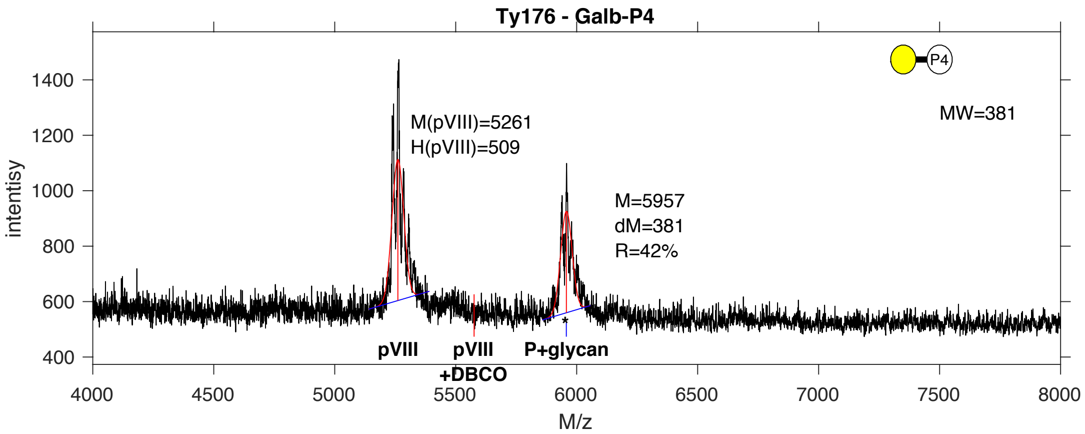


**SDB Number:** SDB10

**Sequencing File:** http://ligacloud.ca/searchLibInfo?f=0&b=0&d=20171128-87CLooOOPTIT-SS

**Barcode:**
CTACTGTTTGCTATACCGCTGAGTGTGGAGAAGAATGATCAGAAGACTTATCATGCGGGTGGAGGT

**Axis Name:** Gal-[1130]

**IUPAC:** Gal(b1-P4

**CFG Name:** Galb-P4

**Common Name:** Gal

**Glytoucan ID:** G65889KE

**Compound Number:** Ty-176

**Maldi File:** 5.txt

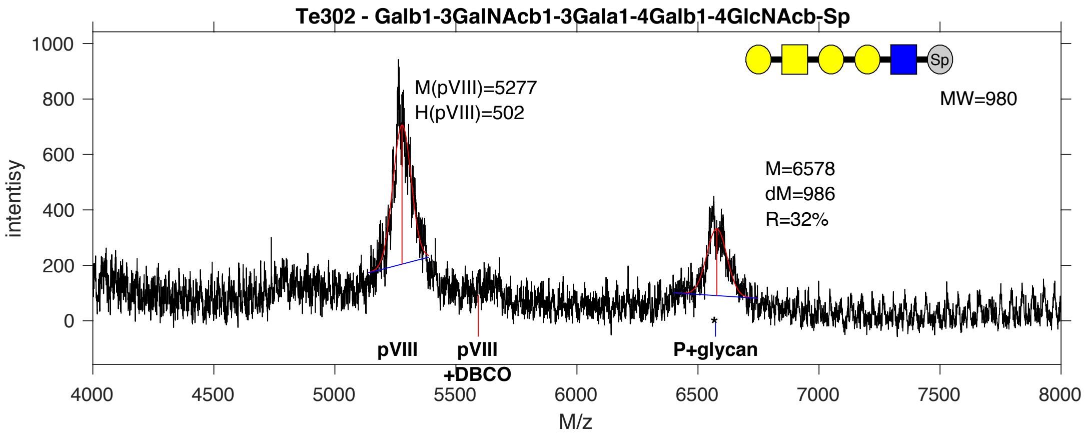


**SDB Number:** SDB13

**Sequencing File:** http://ligacloud.ca/searchLibInfo?f=0&b=0&d=20171128-87CLooOOMKLB-SS

**Barcode:**
CTACTTTTCGCAATTCCTCTGAGTGTGGAGAAGAATGATCAGAAGACTTATCATGCGGGTGGAGGT

**Axis Name:** P1 penta-[860]

**IUPAC:** Gal(b1-3)GalNAc(b1-3)Gal(a1-4)Gal(b1-4)GlcNAc(b1-Sp

**CFG Name:** Galb1-3GalNAcb1-3Gala1-4Galb1-4GlcNAcb-Sp

**Common Name:** P1 penta

**Glytoucan ID:** G28005AF

**Compound Number:** Te302

**Maldi File:** 6.txt

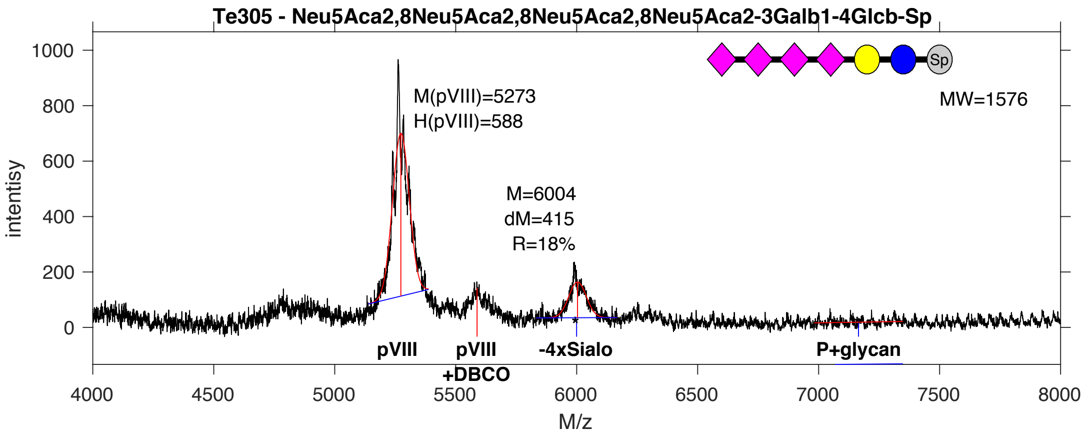


**SDB Number:** SDB15

**Sequencing File:** http://ligacloud.ca/searchLibInfo?f=0&b=0&d=20171128-87CLooOOBQBB-SS

**Barcode:**
CTGCTGTTCGCCATACCCCTTAGTGTGGAGAAGAATGATCAGAAGACTTATCATGCGGGTGGAGGT

**Axis Name:** TetraSLac-[490]

**IUPAC:** Neu5Ac(a2-8)Neu5Ac(a2-8)Neu5Ac(a2-8)Neu5Ac(a2-3)Gal(b1-4)Glc(b1-Sp

**CFG Name:** Neu5Aca2-8Neu5Aca2-8Neu5Aca2-8Neu5Aca2-3Galb1-4Glcb-Sp

**Common Name:** TetraSLac

**Glytoucan ID:** not registered

**Compound Number:** Te305

**Maldi File:** 8.txt

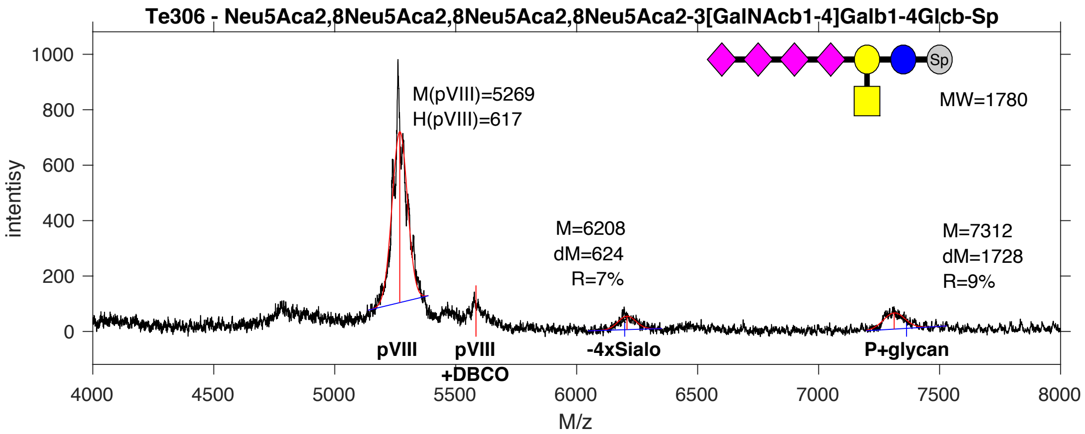


**SDB Number:** SDB16

**Sequencing File:** http://ligacloud.ca/searchLibInfo?f=0&b=0&d=20171128-87CLooOOIEEY-SS

**Barcode:**
CTGCTGTTCGCAATCCCGCTGAGTGTGGAGAAGAATGATCAGAAGACTTATCATGCGGGTGGAGGT

**Axis Name:** GQ2-[430]

**IUPAC:** Neu5Ac(a2-8)Neu5Ac(a2-8)Neu5Ac(a2-8)Neu5Ac(a2-3)[GalNAc(b1-4)]Gal(b1-4)Glc(b1-Sp

**CFG Name:** Neu5Aca2-8Neu5Aca2-8Neu5Aca2-8Neu5Aca2-3[GalNAcb1-4]Galb1-4Glcb-Sp

**Common Name:** GQ2

**Glytoucan ID:** G69728IZ

**Compound Number:** Te306

**Maldi File:** 9.txt

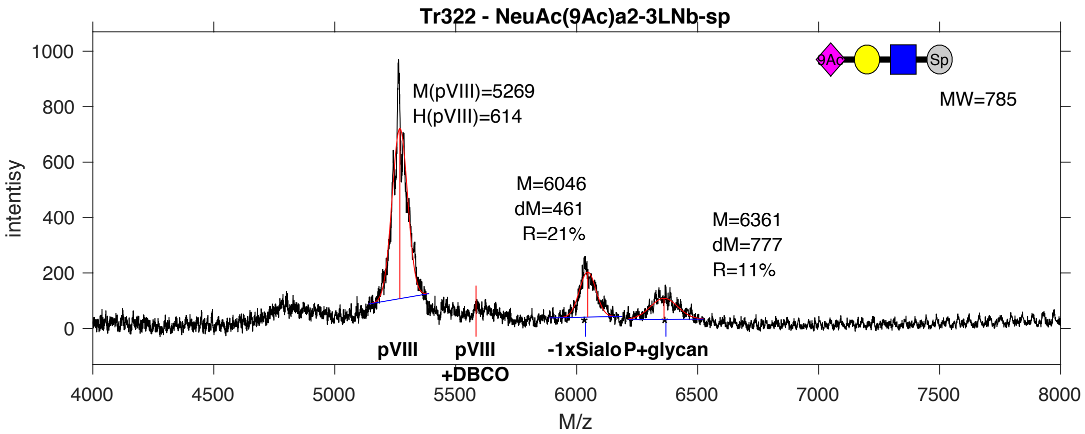


**SDB Number:** SDB17

**Sequencing File:** http://ligacloud.ca/searchLibInfo?f=0&b=0&d=20171128-87CLooOOKVTV-SS

**Barcode:**
CTACTCTTCGCGATTCCGCTTAGTGTGGAGAAGAATGATCAGAAGACTTATCATGCGGGTGGAGGT

**Axis Name:** 9OAc3'SLN-[860]

**IUPAC:** Neu5,9Ac(a2-3)Gal(b1-4)GlcNAc(b1-Sp

**CFG Name:** Neu5Ac(9Ac)a2-3Galb1-4GlcNAcb-Sp

**Common Name:** 9OAc3'SLN

**Glytoucan ID:** G63851XY

**Compound Number:** Tr322

**Maldi File:** 10.txt

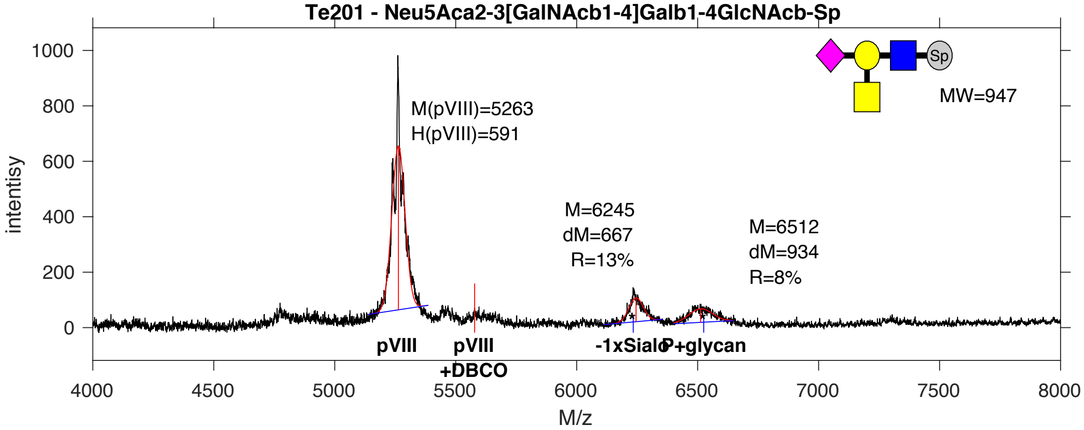


**SDB Number:** SDB21

**Sequencing File:** http://ligacloud.ca/searchLibInfo?f=0&b=0&d=20171128-87CLooOOEHIN-SS

**Barcode:**
CTACTCTTTGCAATTCCCCTTAGTGTGGAGAAGAATGATCAGAAGACTTATCATGCGGGTGGAGGT

**Axis Name:** CT|Sda-[570]

**IUPAC:** Neu5Ac(a2-3)[GalNAc(b1-4)]Gal(b1-4)GlcNAc(b1-Sp

**CFG Name:** Neu5Aca2-3[GalNAcb1-4]Galb1-4GlcNAcb-Sp

**Common Name:** CT|Sda

**Glytoucan ID:** G41052GC

**Compound Number:** Te201

**Maldi File:** 11.txt

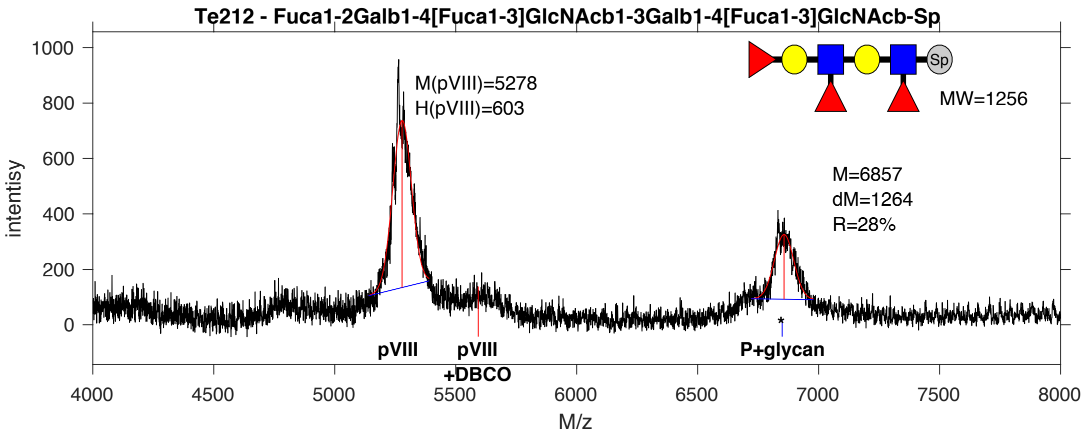


**SDB Number:** SDB22

**Sequencing File:** http://ligacloud.ca/searchLibInfo?f=0&b=0&d=20171128-87CLooOOTVNE-SS

**Barcode:**
CTACTGTTTGCTATCCCACTTAGTGTGGAGAAGAATGATCAGAAGACTTATCATGCGGGTGGAGGT

**Axis Name:** Ley-Lex-[760]

**IUPAC:** Fuc(a1-2)Gal(b1-4)[Fuc(a1-3)]GlcNAc(b1-3)Gal(b1-4)[Fuc(a1-3)]GlcNAc(b1-Sp

**CFG Name:** Fuca1-2Galb1-4[Fuca1-3]GlcNAcb1-3Galb1-4[Fuca1-3]GlcNAcb-Sp

**Common Name:** Ley-Lex

**Glytoucan ID:** G74917JX

**Compound Number:** Te212

**Maldi File:** 12.txt

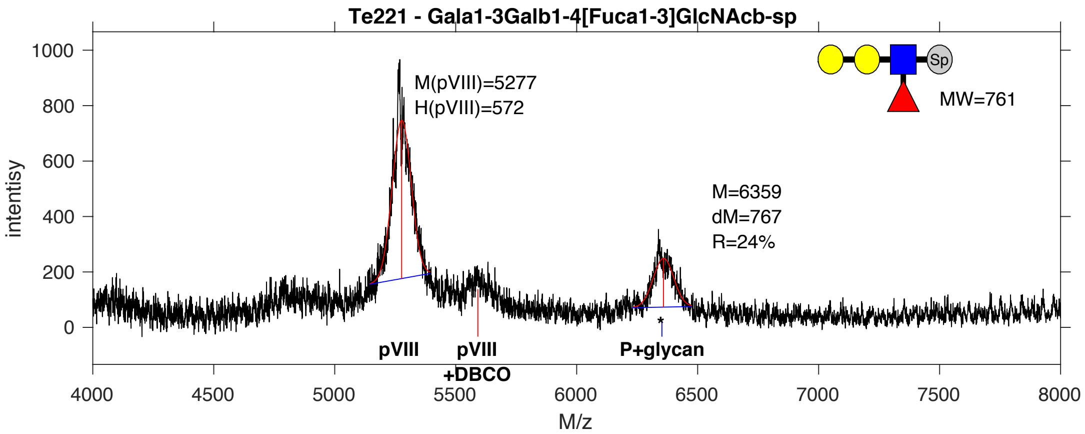


**SDB Number:** SDB23

**Sequencing File:** http://ligacloud.ca/searchLibInfo?f=0&b=0&d=20171128-87CLooOONHEY-SS

**Barcode:**
CTGCTCTTTGCAATACCTCTTAGTGTGGAGAAGAATGATCAGAAGACTTATCATGCGGGTGGAGGT

**Axis Name:** Gala3Lex-[650]

**IUPAC:** Gal(a1-3)Gal(b1-4)[Fuc(a1-3)]GlcNAc(b1-Sp

**CFG Name:** Gala1-3Galb1-4[Fuca1-3]GlcNAcb-Sp

**Common Name:** Gala3Lex

**Glytoucan ID:** G86393KI

**Compound Number:** Te221

**Maldi File:** 13.txt

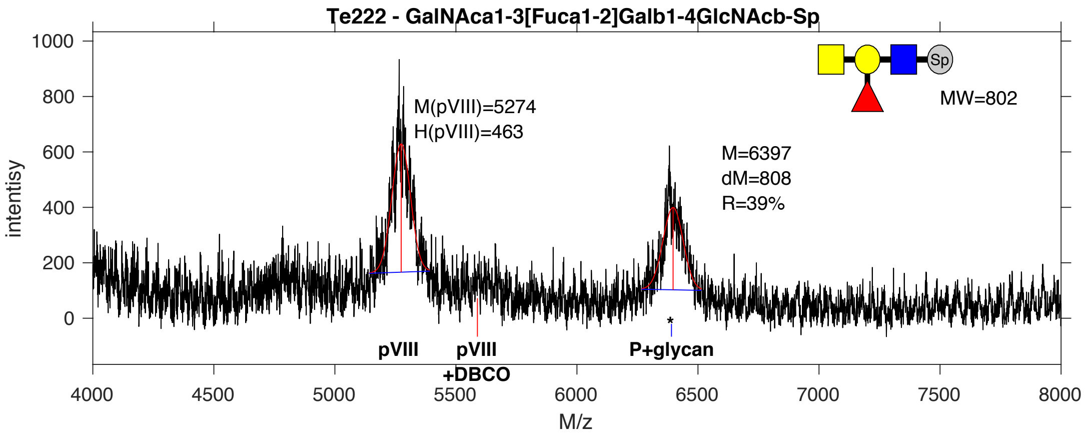


**SDB Number:** SDB24

**Sequencing File:** http://ligacloud.ca/searchLibInfo?f=0&b=0&d=20171128-87CLooOOJDTS-SS

**Barcode:**
CTACTATTCGCGATCCCGCTCAGTGTGGAGAAGAATGATCAGAAGACTTATCATGCGGGTGGAGGT

**Axis Name:** A tetra type 2-[1050]

**IUPAC:** Gal(a1-3)Gal(b1-4)[Fuc(a1-3)]GlcNAc(b1-Sp

**CFG Name:** GalNAca1-3[Fuca1-2]Galb1-4GlcNAcb-Sp

**Common Name:** A tetra type 2

**Glytoucan ID:** G86393KI

**Compound Number:** Te222

**Maldi File:** 14.txt

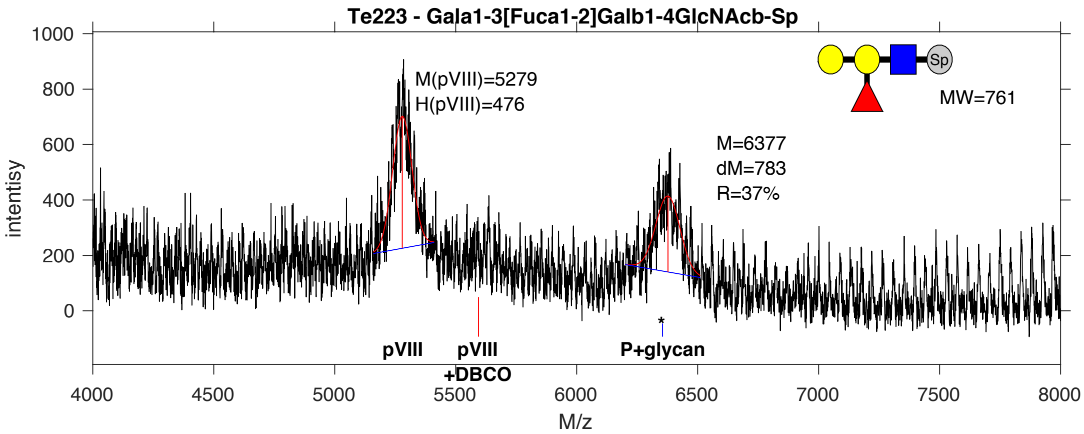


**SDB Number:** SDB26

**Sequencing File:** http://ligacloud.ca/searchLibInfo?f=0&b=0&d=20190703-87CLooOOVHRU-DF

**Barcode:**
CTGCTATTCGCTATCCCACTCAGTGTGGAGAAGAATGATCAGAAGACTTATCATGCGGGTGGAGGT

**Axis Name:** B tetra type 2-[1000]

**IUPAC:** Gal(a1-3)[Fuc(a1-2)]Gal(b1-4)GlcNAc(b1-Sp

**CFG Name:** Gala1-3[Fuca1-2]Galb1-4GlcNAcb-Sp

**Common Name:** B tetra type 2

**Glytoucan ID:** G45802NQ

**Compound Number:** Te223

**Maldi File:** 15.txt

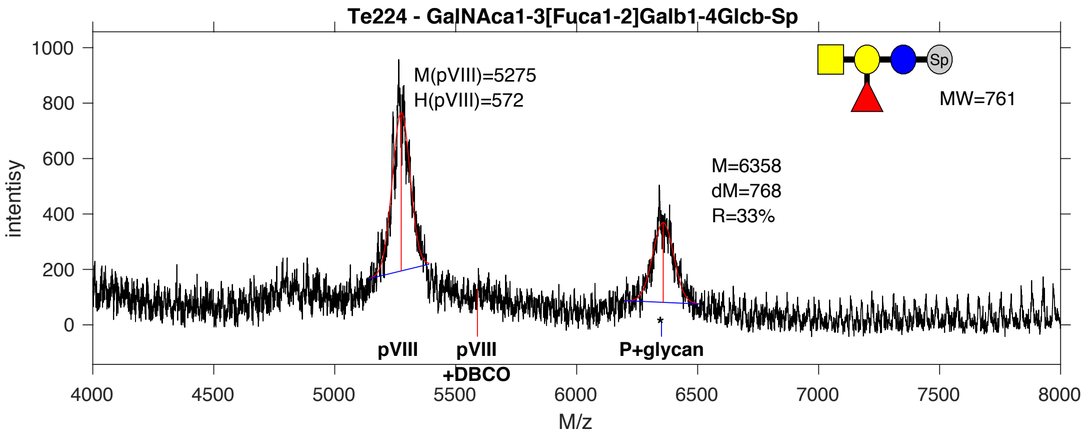


**SDB Number:** SDB29

**Sequencing File:** http://ligacloud.ca/searchLibInfo?f=0&b=0&d=20171128-87CLooOOSNHI-SS

**Barcode:**
CTGCTATTTGCGATCCCGCTGAGTGTGGAGAAGAATGATCAGAAGACTTATCATGCGGGTGGAGGT

**Axis Name:** A tetra L-[890]

**IUPAC:** GalNAc(a1-3)[Fuc(a1-2)]Gal(b1-4)Glc(b1-Sp

**CFG Name:** GalNAca1-3[Fuca1-2]Galb1-4Glcb-Sp

**Common Name:** A tetra L

**Glytoucan ID:** G19524AS

**Compound Number:** Te224

**Maldi File:** 16.txt

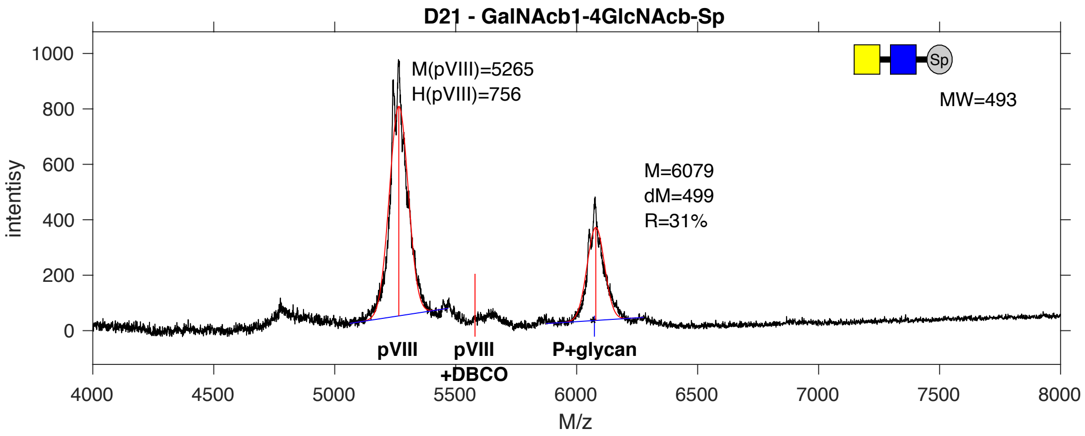


**SDB Number:** SDB34

**Sequencing File:** http://ligacloud.ca/searchLibInfo?f=0&b=0&d=20171128-87CLooOOTLAJ-SS

**Barcode:**
CTTCTTTTTGCGATTCCGCTGAGTGTGGAGAAGAATGATCAGAAGACTTATCATGCGGGTGGAGGT

**Axis Name:** LacDiNAc-[840]

**IUPAC:** GalNAc(b1-4)GlcNAc(b1-Sp

**CFG Name:** GalNAcb1-4GlcNAcb-Sp

**Common Name:** LacDiNAc

**Glytoucan ID:** G33129LQ

**Compound Number:** D21

**Maldi File:** 17.txt

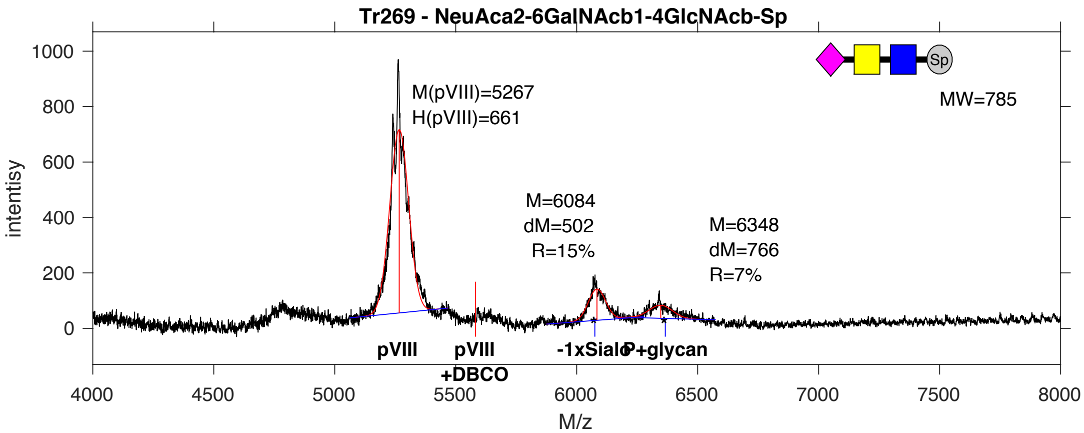


**SDB Number:** SDB35

**Sequencing File:** http://ligacloud.ca/searchLibInfo?f=0&b=0&d=20171128-87CLooOOBBJH-SS

**Barcode:**
CTGCTCTTCGCTATTCCACTTAGTGTGGAGAAGAATGATCAGAAGACTTATCATGCGGGTGGAGGT

**Axis Name:** 6’SLDN-[590]

**IUPAC:** Neu5Ac(a2-6)GalNAc(b1-4)GlcNAc(b1-Sp

**CFG Name:** Neu5Aca2-6GalNAcb1-4GlcNAcb-Sp

**Common Name:** 6’SLDN

**Glytoucan ID:** G54118CT

**Compound Number:** Tr269

**Maldi File:** 18.txt

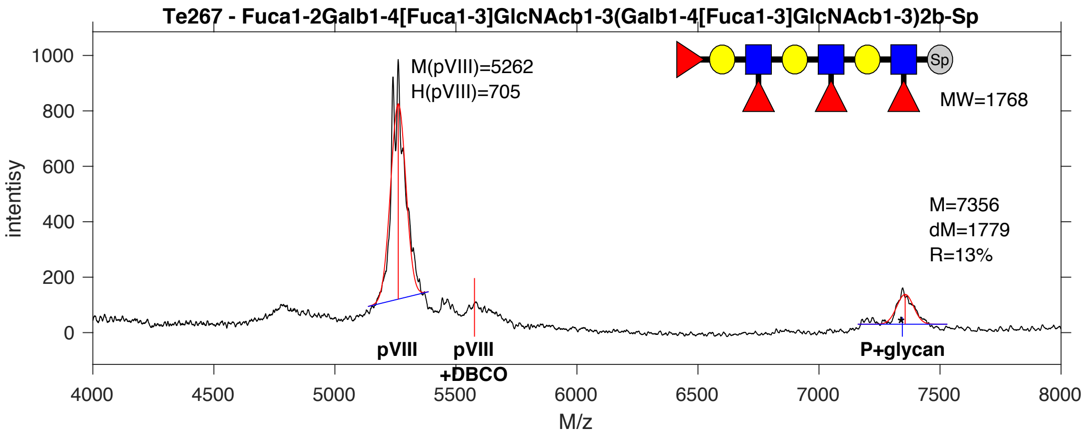


**SDB Number:** SDB40

**Sequencing File:** http://ligacloud.ca/searchLibInfo?f=0&b=0&d=20180809-87CLooPA1-1JM7

**Barcode:**
CTTCTGTTCGCTATTCCGCTTAGCGTGGAAAAGAACGATCAAAAGACCTATCACGCCGGGGGAGGG

**Axis Name:** Ley-Di-Lex-[320]

**IUPAC:** Fuca1-2Galb1-4[Fuca1-3]GlcNAcb1-3(Galb1-4[Fuca1-3]GlcNAcb1-3)2b-SpSp

**CFG Name:** Fuca1-2Galb1-4[Fuca1-3]GlcNAcb1-3(Galb1-4[Fuca1-3]GlcNAcb1-3)2b-Sp

**Common Name:** Ley-Di-Lex

**Glytoucan ID:** not registered

**Compound Number:** Te267

**Maldi File:** 19.txt

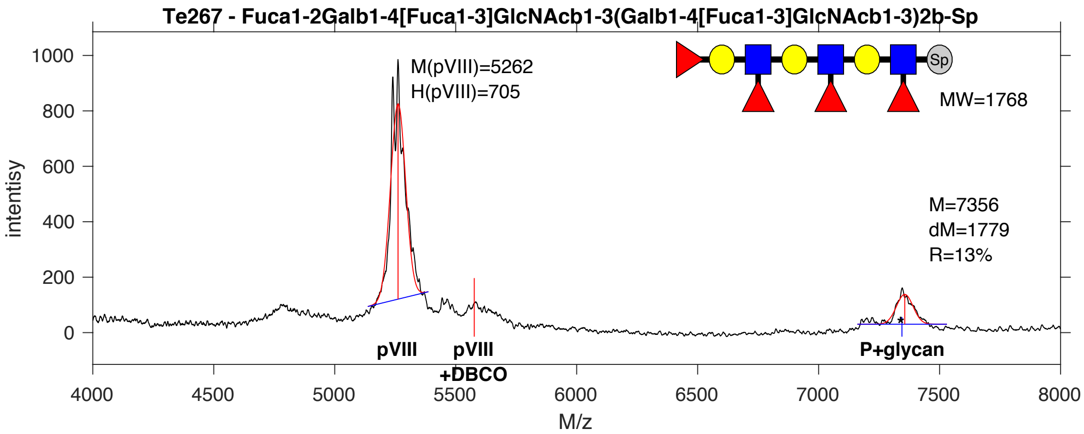


**SDB Number:** SDB40

**Sequencing File:** http://ligacloud.ca/searchLibInfo?f=0&b=0&d=20180809-87CLooPA1-1JM7

**Barcode:**
CTTCTGTTCGCTATTCCGCTTAGCGTGGAAAAGAACGATCAAAAGACCTATCACGCCGGGGGAGGG

**Axis Name:** Ley-Di-Lex-[320]

**IUPAC:** Fuca1-2Galb1-4[Fuca1-3]GlcNAcb1-3(Galb1-4[Fuca1-3]GlcNAcb1-3)2b-SpSp

**CFG Name:** Fuca1-2Galb1-4[Fuca1-3]GlcNAcb1-3(Galb1-4[Fuca1-3]GlcNAcb1-3)2b-Sp

**Common Name:** Ley-Di-Lex

**Glytoucan ID:** not registered

**Compound Number:** Te267

**Maldi File:** 19.txt

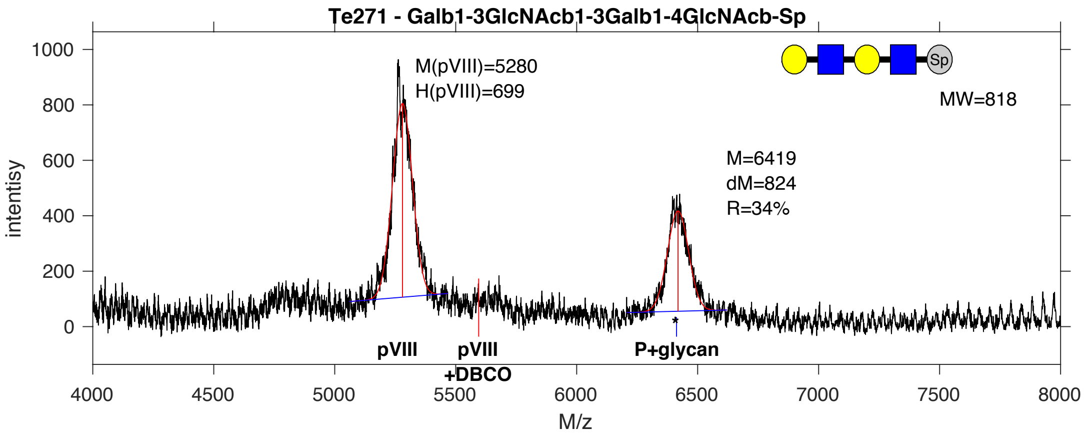


**SDB Number:** SDB42

**Sequencing File:** http://ligacloud.ca/searchLibInfo?f=0&b=0&d=20180809-87CLooPA1-1JM9

**Barcode:**
CTTCTGTTTGCAATACCCCTGAGCGTGGAGAAGAATGATCAAAAGACGTACCATGCGGGGGGAGGG

**Axis Name:** LNT-Nac-[920]

**IUPAC:** Gal(b1-3)GlcNAc(b1-3)Gal(b1-4)GlcNAc(b1-Sp

**CFG Name:** Galb1-3GlcNAcb1-3Galb1-4GlcNAcb-Sp

**Common Name:** LNT-Nac

**Glytoucan ID:** G09681LD

**Compound Number:** Te271

**Maldi File:** 20.txt

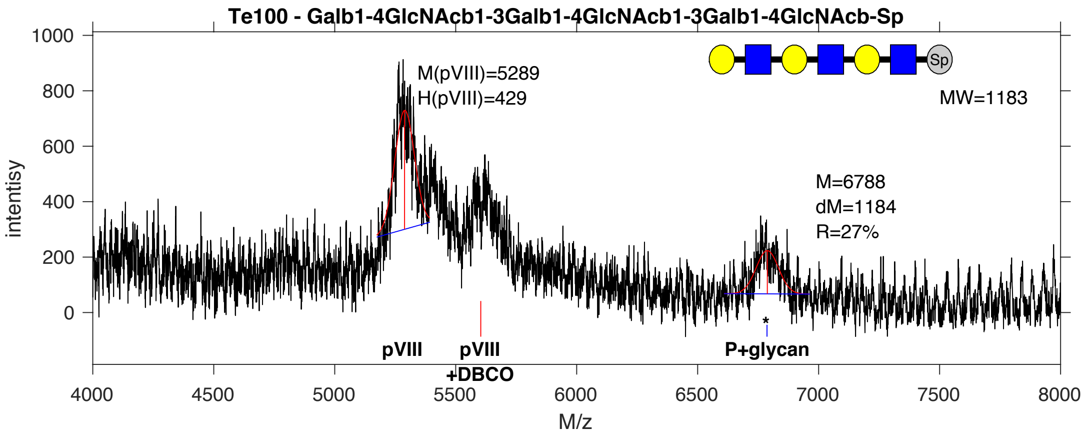


**SDB Number:** SDB47

**Sequencing File:** http://ligacloud.ca/searchLibInfo?f=0&b=0&d=20180809-87CLooPA1-1JM14

**Barcode:**
TTATTATTCGCAATACCGCTAAGTGTTGAGAAGAATGACCAGAAGACGTATCACGCCGGGGGAGGG

**Axis Name:** Tri-LN-[730]

**IUPAC:** Gal(b1-4)GlcNAc(b1-3)Gal(b1-4)GlcNAc(b1-3)Gal(b1-4)GlcNAc(b1-Sp

**CFG Name:** (Galb1-4GlcNAcb1-3)3b-Sp

**Common Name:** Tri-LN

**Glytoucan ID:** G46055MA

**Compound Number:** Te100

**Maldi File:** 22.txt

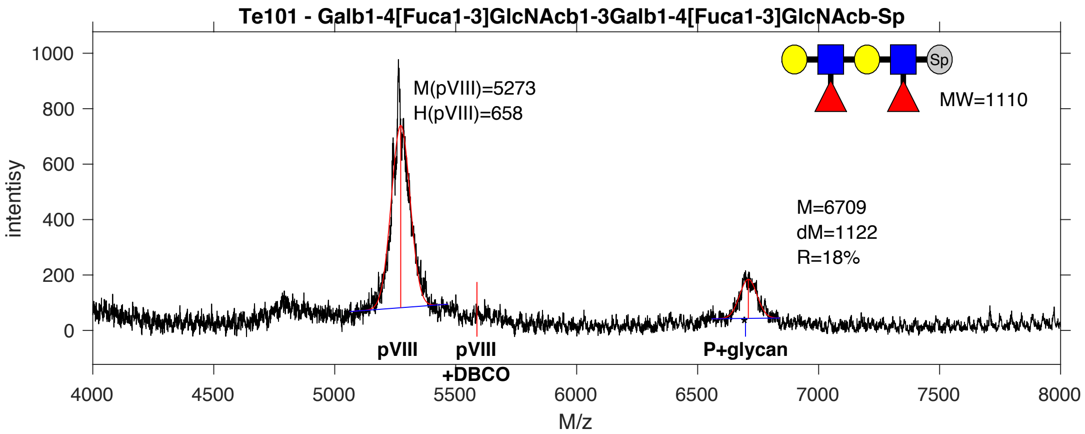


**SDB Number:** SDB48

**Sequencing File:** http://ligacloud.ca/searchLibInfo?f=0&b=0&d=20180809-87CLooPA1-1JM15

**Barcode:**
TTATTATTCGCAATTCCTTTAAGCGTAGAGAAAAACGACCAGAAGACCTATCACGCGGGAGGAGGT

**Axis Name:** Di-Lex-[490]

**IUPAC:** Gal(b1-4)[Fuc(a1-3)]GlcNAc(b1-3)Gal(b1-4)[Fuc(a1-3)]GlcNAc(b1-Sp

**CFG Name:** (Galb1-4[Fuca1-3]GlcNAcb1-3)2b-Sp

**Common Name:** Di-Lex

**Glytoucan ID:** G79535HZ

**Compound Number:** Te101

**Maldi File:** 23.txt

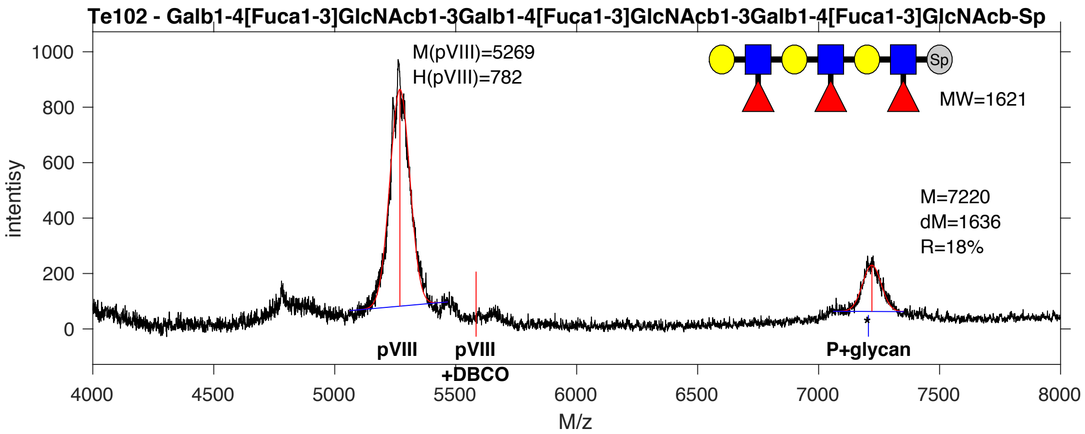


**SDB Number:** SDB49

**Sequencing File:** http://ligacloud.ca/searchLibInfo?f=0&b=0&d=20180809-87CLooPA1-1JM16

**Barcode:**
CTACTGTTCGCTATCCCGCTGAGTGTGGAAAAGAATGATCAGAAGACTTACCACGCTGGTGGTGGG

**Axis Name:** Tri-Lex-[490]

**IUPAC:** Gal(b1-4)[Fuc(a1-3)]GlcNAc(b1-3)Gal(b1-4)[Fuc(a1-3)]GlcNAc(b1-3)Gal(b1-4)[Fuc(a1-3)]GlcNAc(b1-Sp

**CFG Name:** (Galb1-4[Fuca1-3]GlcNAcb1-3)3b-Sp

**Common Name:** Tri-Lex

**Glytoucan ID:** G57884AX

**Compound Number:** Te102

**Maldi File:** 24.txt

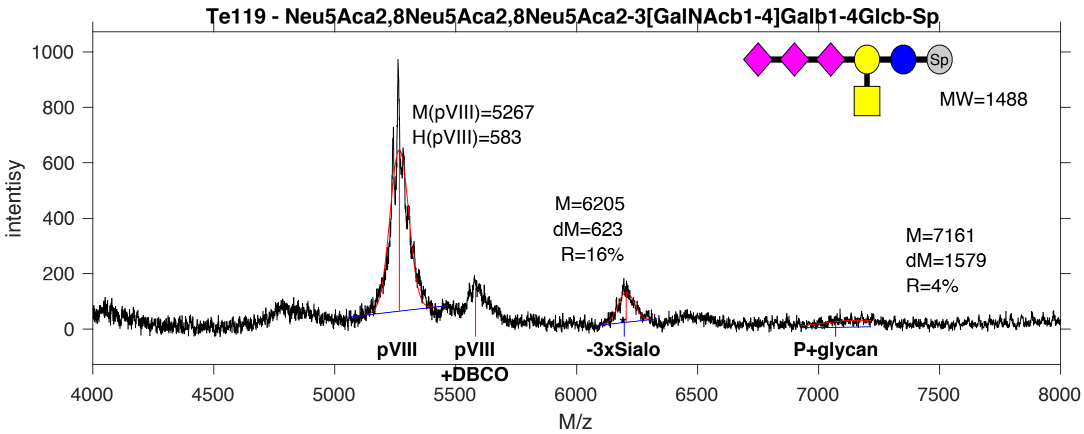


**SDB Number:** SDB51

**Sequencing File:** http://ligacloud.ca/searchLibInfo?f=0&b=0&d=20180809-87CLooPA1-1JM18

**Barcode:**
CTGCTCTTTGCTATACCCCTAAGCGTCGAGAAAAATGATCAAAAAACCTATCACGCCGGGGGGGGA

**Axis Name:** GT2-[430]

**IUPAC:** Neu5Ac(a2-8)Neu5Ac(a2-8)Neu5Ac(a2-3)[GalNAc(b1-4)]Gal(b1-4)Glc(b1-Sp

**CFG Name:** Neu5Aca2-8Neu5Aca2-8Neu5Aca2-3[GalNAcb1-4]Galb1-4Glcb-Sp

**Common Name:** GT2

**Glytoucan ID:** G35228ZS

**Compound Number:** Te119

**Maldi File:** 25.txt

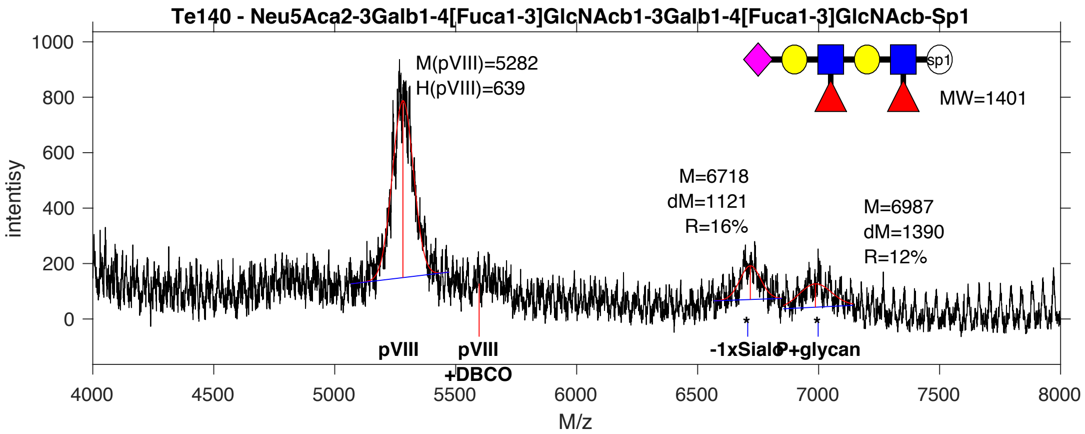


**SDB Number:** SDB52

**Sequencing File:** http://ligacloud.ca/searchLibInfo?f=0&b=0&d=20171128-87CLooOORRZY-SS

**Barcode:**
CTACTGTTTGCTATCCCGCTGAGTGTAGAGAAGAATGACCAAAAGACATATCATGCGGGAGGAGGT

**Axis Name:** 3'S-Di-Lex-[760]

**IUPAC:** Neu5Ac(a2-3)Gal(b1-4)[Fuc(a1-3)]GlcNAc(b1-3)Gal(b1-4)[Fuc(a1-3)]GlcNAc(b1-Sp

**CFG Name:** Neu5Aca2-3(Galb1-4[Fuca1-3]GlcNAcb1-3)2b-Sp

**Common Name:** 3'S-Di-Lex

**Glytoucan ID:** G19619KS

**Compound Number:** Te140

**Maldi File:** 26.txt

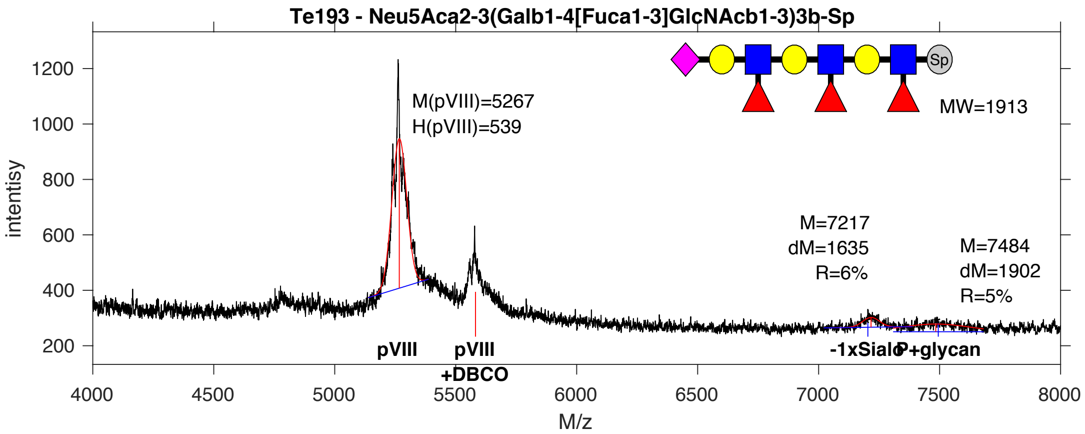


**SDB Number:** SDB53

**Sequencing File:** http://ligacloud.ca/searchLibInfo?f=0&b=0&d=20171128-87CLooOOWGUJ-SS

**Barcode:**
CTGCTGTTTGCGATCCCGCTCAGTGTGGAGAAAAATGACCAAAAGACCTATCATGCTGGAGGGGGT

**Axis Name:** 3'S-Tri-LeX-[220]

**IUPAC:** Neu5Ac(a2-3)Gal(b1-4)[Fuc(a1-3)]GlcNAc(b1-3)Gal(b1-4)[Fuc(a1-3)]GlcNAc(b1-3)Gal(b1-4)[Fuc(a1-3)]GlcNAc(b1-Sp

**CFG Name:** Neu5Aca2-3(Galb1-4[Fuca1-3]GlcNAcb1-3)3b-Sp

**Common Name:** 3'S-Tri-LeX

**Glytoucan ID:** G70868ZC

**Compound Number:** Te193

**Maldi File:** 27.txt

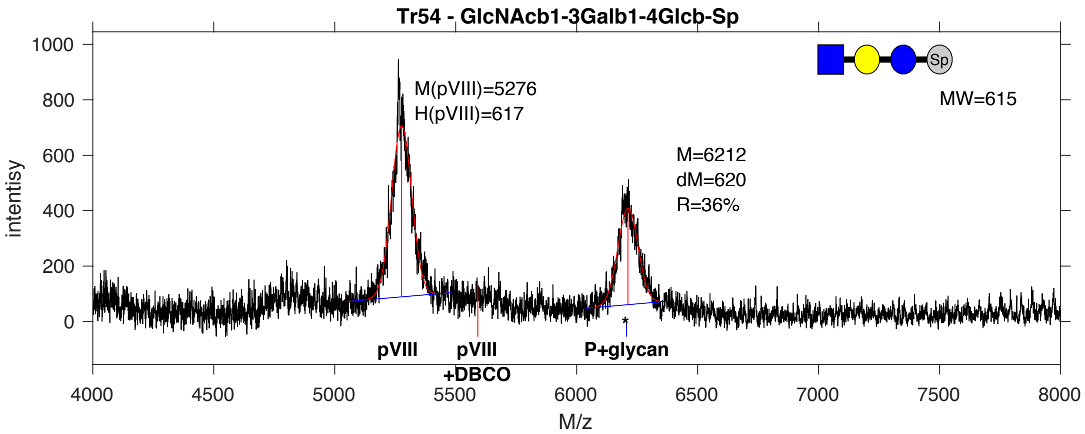


**SDB Number:** SDB54

**Sequencing File:** http://ligacloud.ca/searchLibInfo?f=0&b=0&d=20171128-87CLooOOBRPI-SS

**Barcode:**
TTATTATTCGCAATTCCTTTAAGTGTCGAAAAAAACGATCAGAAGACCTATCACGCAGGCGGGGGC

**Axis Name:** LNT-2-[970]

**IUPAC:** GlcNAc(b1-3)Gal(b1-4)Glc(b1-Sp

**CFG Name:** GlcNAcb1-3Galb1-4Glcb-Sp

**Common Name:** LNT-2

**Glytoucan ID:** G64979HM

**Compound Number:** Tr54

**Maldi File:** 28.txt

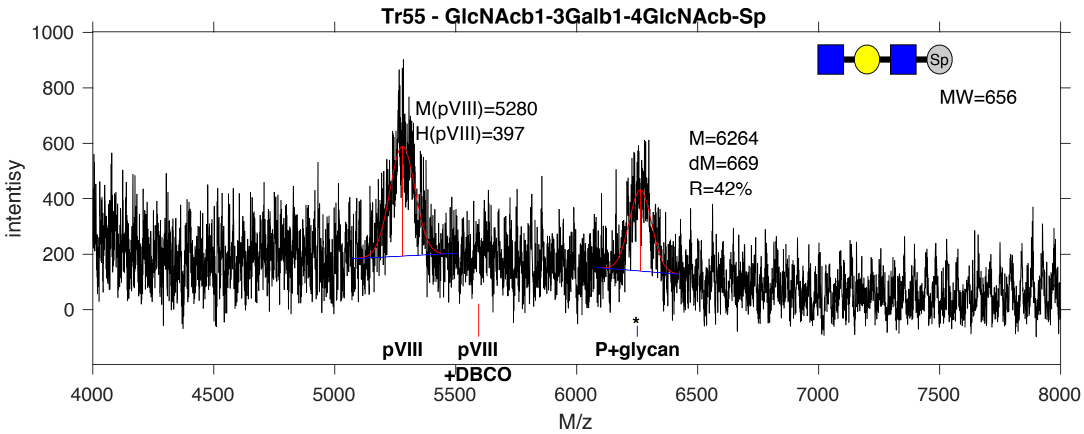


**SDB Number:** SDB55

**Sequencing File:** http://ligacloud.ca/searchLibInfo?f=0&b=0&d=20180809-87CLooPA1-1JM22

**Barcode:**
CTGCTCTTTGCAATTCCGCTGAGTGTCGAAAAAAATGACCAAAAGACTTATCACGCTGGTGGTGGA

**Axis Name:** GNLN-[1130]

**IUPAC:** GlcNAc(b1-3)Gal(b1-4)GlcNAc(b1-Sp

**CFG Name:** GlcNAcb1-3Galb1-4GlcNAcb-Sp

**Common Name:** GNLN

**Glytoucan ID:** G29487JS

**Compound Number:** Tr55

**Maldi File:** 29.txt

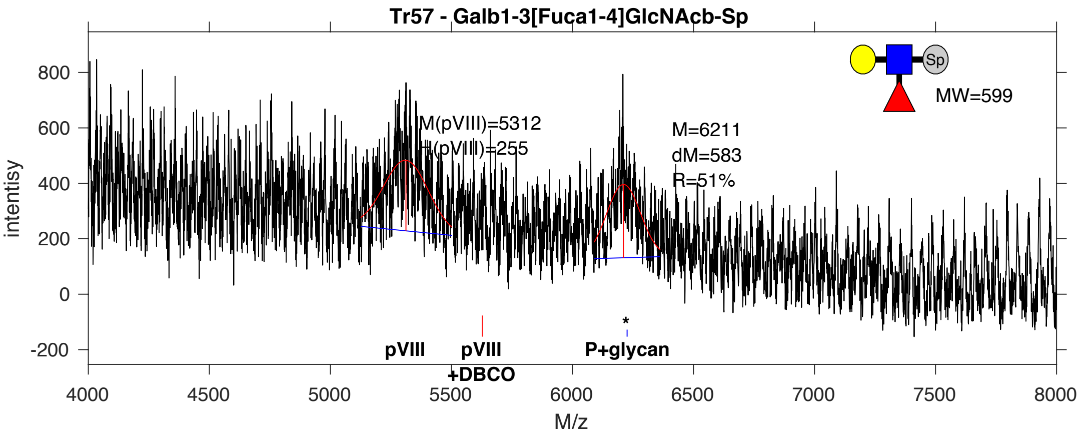


**SDB Number:** SDB56

**Sequencing File:** http://ligacloud.ca/searchLibInfo?f=0&b=0&d=20180809-87CLooPA1-1JM23

**Barcode:**
CTACTGTTTGCTATACCTCTTAGCGTTGAAAAAAATGACCAGAAAACTTACCATGCGGGTGGAGGT

**Axis Name:** LeA-[1380]

**IUPAC:** Gal(b1-3)[Fuc(a1-4)]GlcNAc(b1-Sp

**CFG Name:** Galb1-3[Fuca1-4]GlcNAcb-Sp

**Common Name:** LeA

**Glytoucan ID:** G39023AU

**Compound Number:** Tr57

**Maldi File:** 30.txt

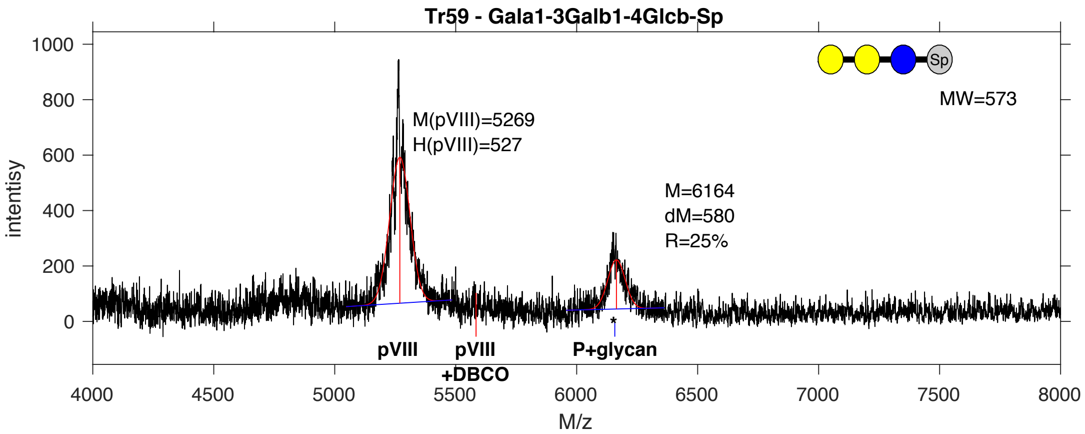


**SDB Number:** SDB57

**Sequencing File:** http://ligacloud.ca/searchLibInfo?f=0&b=0&d=20171128-87CLooOORPVC-SS

**Barcode:**
TTATTATTCGCAATTCCTTTAAGCGTCGAGAAGAACGACCAAAAAACCTATCACGCAGGTGGCGGG

**Axis Name:** Galili-tri-[680]

**IUPAC:** Gal(a1-3)Gal(b1-4)Glc(b1-Sp

**CFG Name:** Gala1-3Galb1-4Glcb-Sp

**Common Name:** Galili-tri

**Glytoucan ID:** G78074HX

**Compound Number:** Tr59

**Maldi File:** 31.txt

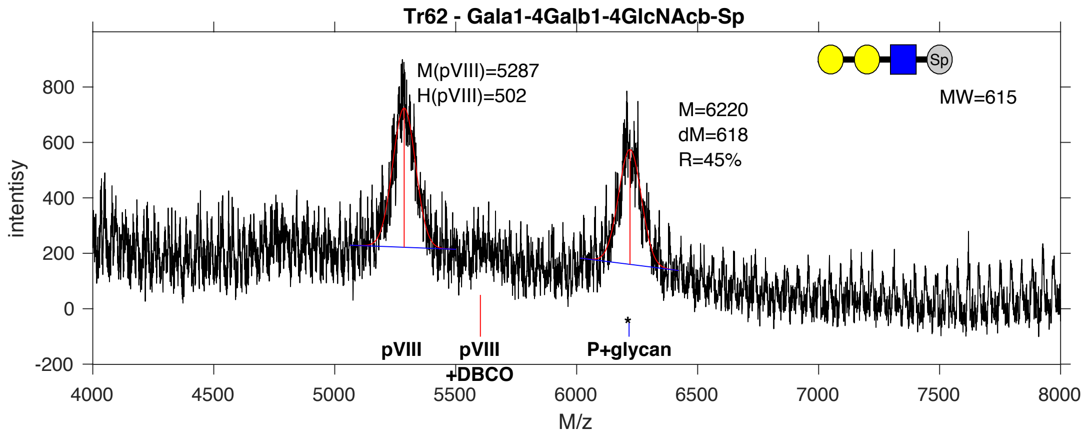


**SDB Number:** SDB58

**Sequencing File:** http://ligacloud.ca/searchLibInfo?f=0&b=0&d=20171128-87CLooOOQAMP-SS

**Barcode:**
CTACTCTTCGCTATACCCCTCAGCGTGGAGAAAAACGACCAAAAGACCTACCATGCAGGTGGTGGC

**Axis Name:** P1 tri-[1220]

**IUPAC:** Gal(a1-4)Gal(b1-4)GlcNAc(b1-Sp

**CFG Name:** Gala1-4Galb1-4GlcNAcb-Sp

**Common Name:** P1 tri

**Glytoucan ID:** G00076PK

**Compound Number:** Tr62

**Maldi File:** 32.txt

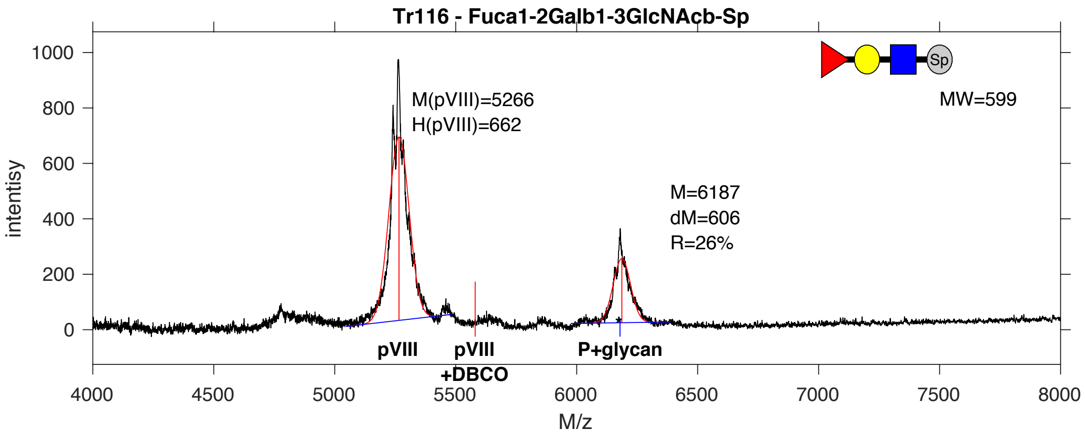


**SDB Number:** SDB59

**Sequencing File:** http://ligacloud.ca/searchLibInfo?f=0&b=0&d=20190703-87CLooOOJPTG-DF

**Barcode:**
TTATTATTCGCAATTCCTTTAAGTGTCGAAAAGAATGACCAAAAAACTTATCATGCAGGTGGGGGA

**Axis Name:** H-type1-[700]

**IUPAC:** Fuc(a1-2)Gal(b1-3)GlcNAc(b1-Sp

**CFG Name:** Fuca1-2Galb1-3GlcNAcb-Sp

**Common Name:** H-type1

**Glytoucan ID:** G43418EX

**Compound Number:** Tr116

**Maldi File:** 33.txt

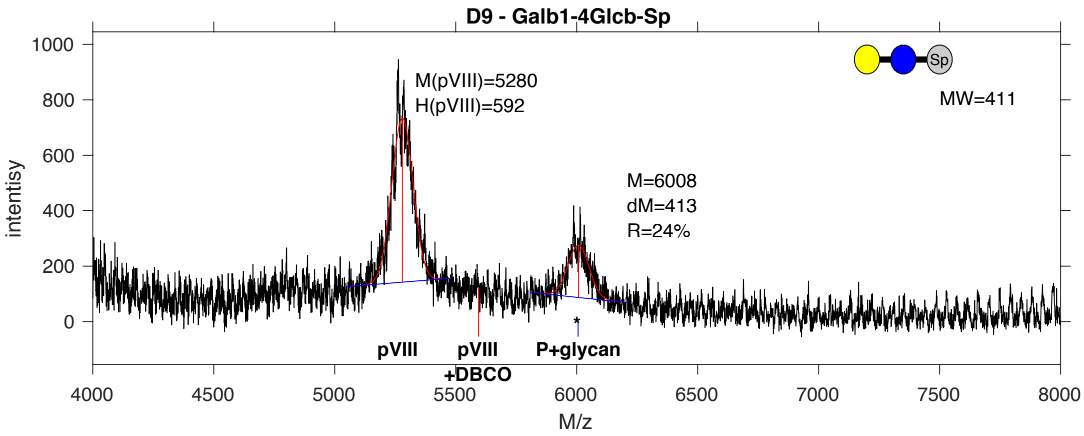


**SDB Number:** SDB66

**Sequencing File:** http://ligacloud.ca/searchLibInfo?f=0&b=0&d=20190703-87CLooORORN-DF

**Barcode:**
TTATTATTCGCAATTCCTTTAAGTGTCGAGAAGAACGATCAGAAGACCTACCACGCAGGGGGGGGT

**Axis Name:** Lac-[650]

**IUPAC:** Gal(b1-4)Glc(b1-Sp

**CFG Name:** Galb1-4Glcb-Sp

**Common Name:** Lac

**Glytoucan ID:** G94144EF

**Compound Number:** D9

**Maldi File:** 34.txt

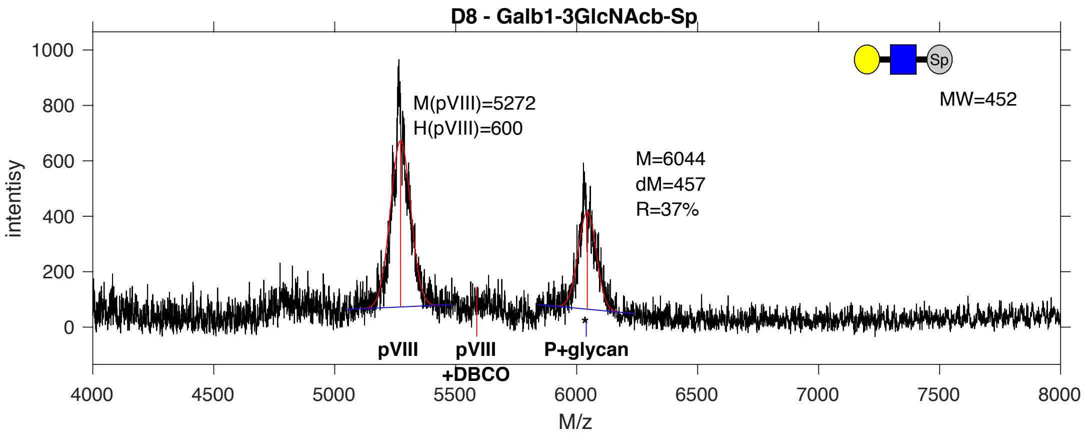


**SDB Number:** SDB67

**Sequencing File:** http://ligacloud.ca/searchLibInfo?f=0&b=0&d=20171128-87CLooOOSXNG-SS

**Barcode:**
CTTCTGTTCGCTATACCTCTCAGCGTGGAGAAGAATGACCAGAAAACTTATCACGCAGGAGGTGGA

**Axis Name:** Lec-[1000]

**IUPAC:** Gal(b1-3)GlcNAc(b1-Sp

**CFG Name:** Galb1-3GlcNAcb-Sp

**Common Name:** Lec

**Glytoucan ID:** G16384KS

**Compound Number:** D8

**Maldi File:** 35.txt

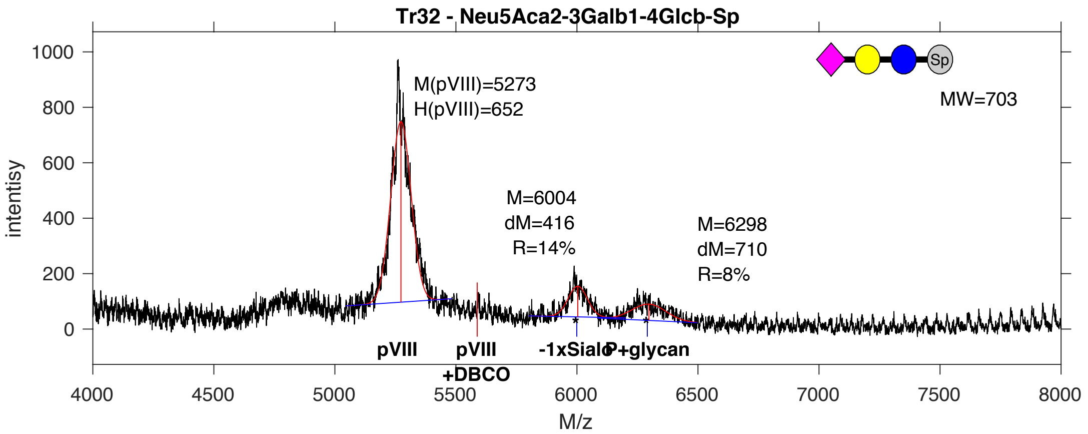


**SDB Number:** SDB70

**Sequencing File:** http://ligacloud.ca/searchLibInfo?f=0&b=0&d=20171128-87CLooOOJSQJ-SS

**Barcode:**
CTACTGTTCGCAATCCCGCTCAGTGTTGAAAAAAACGATCAAAAAACGTATCATGCTGGTGGAGGT

**Axis Name:** GM3-[590]

**IUPAC:** Neu5Ac(a2-3)Gal(b1-4)Glc(b1-Sp

**CFG Name:** Neu5Aca2-3Galb1-4Glcb-Sp

**Common Name:** GM3

**Glytoucan ID:** G91237TK

**Compound Number:** Tr32

**Maldi File:** 36.txt

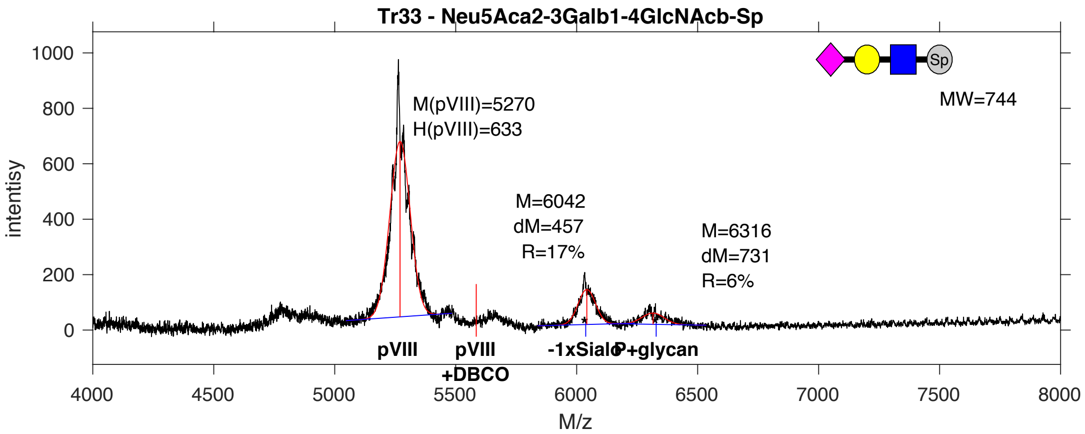


**SDB Number:** SDB71

**Sequencing File:** http://ligacloud.ca/searchLibInfo?f=0&b=0&d=20171128-87CLooOOBZUS-SS

**Barcode:**
CTGCTTTTTGCTATTCCTCTGAGTGTTGAAAAAAACGATCAGAAGACTTATCACGCGGGGGGCGGG

**Axis Name:** 3’SLN-[650]

**IUPAC:** Neu5Ac(a2-3)Gal(b1-4)GlcNAc(b1-Sp

**CFG Name:** Neu5Aca2-3Galb1-4GlcNAcb-Sp

**Common Name:** 3’SLN

**Glytoucan ID:** G04307DP

**Compound Number:** Tr33

**Maldi File:** 37.txt

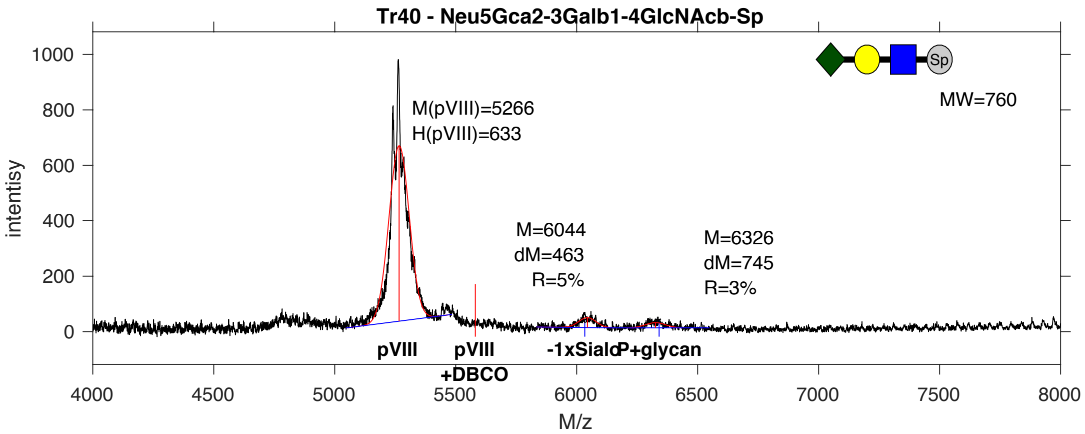


**SDB Number:** SDB73

**Sequencing File:** http://ligacloud.ca/searchLibInfo?f=0&b=0&d=20190703-87CLooOOPUOD-DF

**Barcode:**
TTATTATTCGCAATTCCTTTAAGCGTTGAAAAAAACGATCAAAAGACATATCACGCTGGCGGAGGA

**Axis Name:** 3'SLN (Gc)-[190]

**IUPAC:** Neu5Gc(a2-3)Gal(b1-4)GlcNAc(b1-Sp

**CFG Name:** Neu5Gca2-3Galb1-4GlcNAcb-Sp

**Common Name:** 3'SLN (Gc)

**Glytoucan ID:** G08169IN

**Compound Number:** Tr40

**Maldi File:** 38.txt

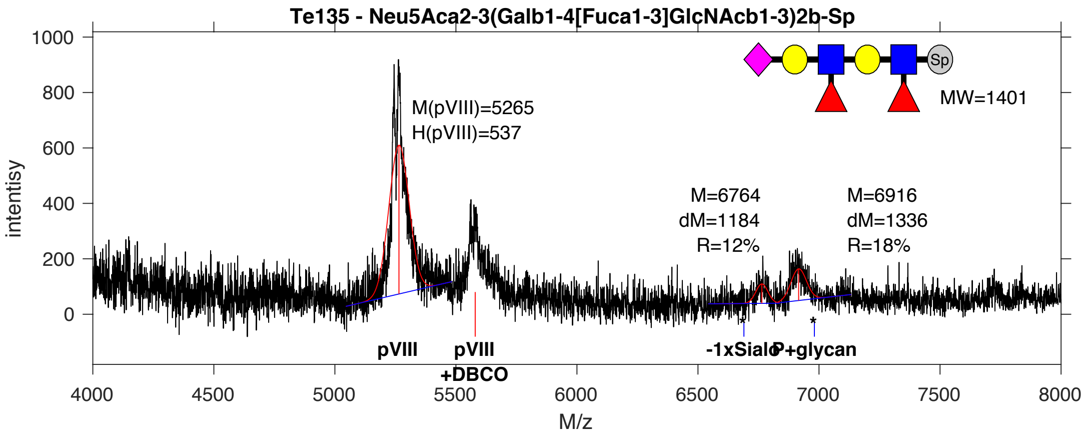


**SDB Number:** SDB77

**Sequencing File:** http://ligacloud.ca/searchLibInfo?f=0&b=0&d=20180522-87YJooOO1-1JM2

**Barcode:**
CTGCTATTTGCCATCCCGCTGAGCGTCGAGAAAAACGATCAAAAGACATACCATGCCGGCGGAGGA

**Axis Name:** H3-[810]

**IUPAC:** Fuc(a1-2)Gal(b1-4)GlcNAc(b1-3)Gal(b1-4)GlcNAc(b1-3)Gal(b1-4)GlcNAc(b1-Sp

**CFG Name:** Fuca1-2(Galb1-4GlcNAcb1-3)3b -Sp

**Common Name:** H3

**Glytoucan ID:** G85213RR

**Compound Number:** Te135

**Maldi File:** 40.txt

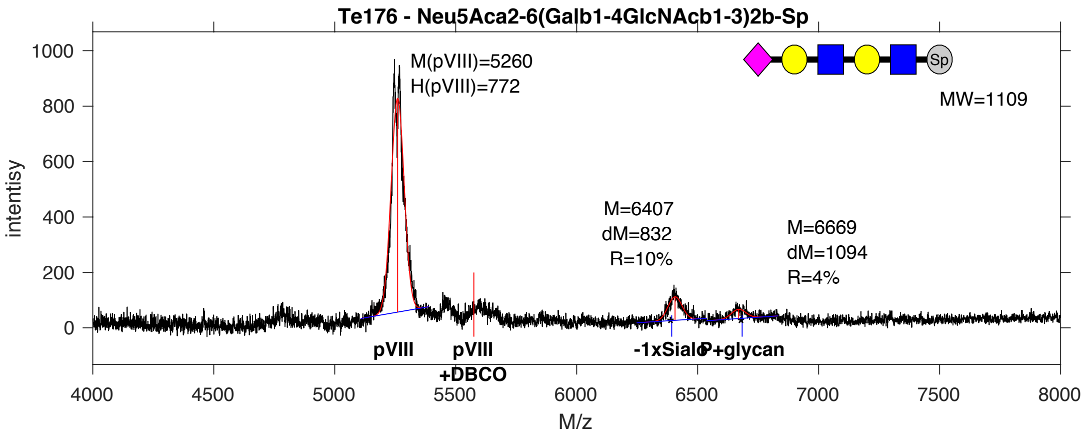


**SDB Number:** SDB79

**Sequencing File:** http://ligacloud.ca/searchLibInfo?f=0&b=0&d=20180522-87YJooOO1-1JM5

**Barcode:**
CTACTGTTCGCAATACCTCTCAGCGTGGAGAAAAACGACCAGAAAACGTATCATGCTGGTGGTGGA

**Axis Name:** 6’S-Di-LN-[380]

**IUPAC:** Neu5Ac(a2-6)Gal(b1-4)GlcNAc(b1-3)Gal(b1-4)GlcNAc(b1-Sp

**CFG Name:** Neu5Aca2-6(Galb1-4GlcNAcb1-3)2b-Sp

**Common Name:** 6’S-Di-LN

**Glytoucan ID:** G60430KO

**Compound Number:** Te176

**Maldi File:** 42.txt

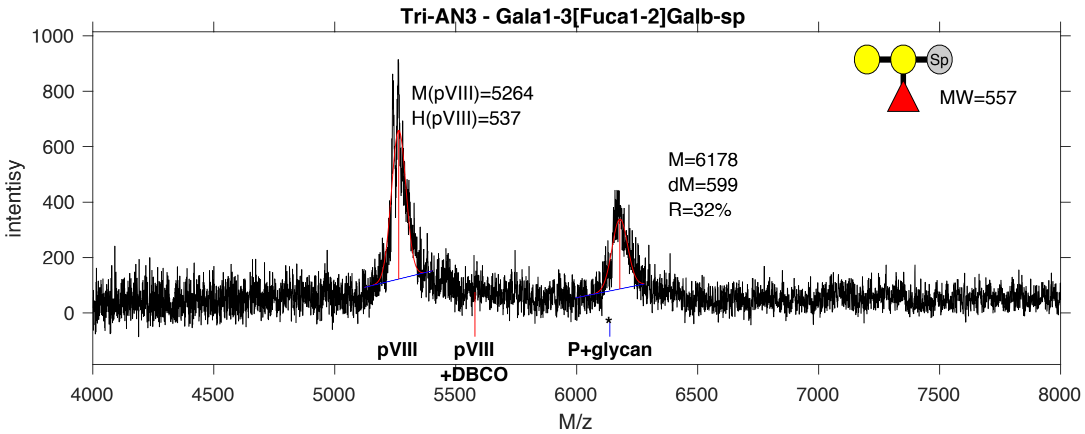


**SDB Number:** SDB80

**Sequencing File:** http://ligacloud.ca/searchLibInfo?f=0&b=0&d=20171128-87CLooOOINAX-SS

**Barcode:**
CTTCTGTTCGCGATCCCCCTAAGTGTAGAGAAGAATGATCAGAAAACGTATCATGCGGGAGGGGGT

**Axis Name:** Tri-AN3-[860]

**IUPAC:** GalNAc(a1-3)[Fuc(a1-2)]Gal(b1-Sp

**CFG Name:** Gala1-3[Fuca1-2]Galb-Sp

**Common Name:** Tri-AN3

**Glytoucan ID:** G34704BH

**Compound Number:** Tri-AN3

**Maldi File:** 43.txt

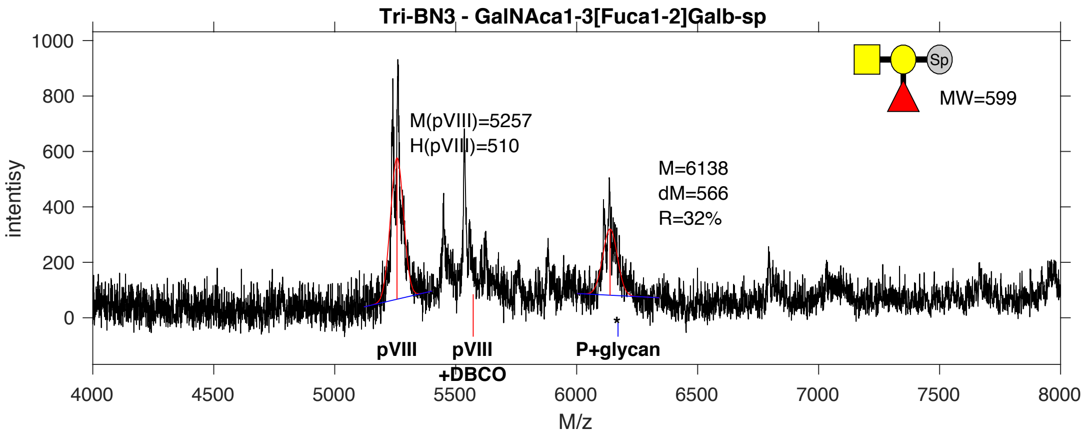


**SDB Number:** SDB81

**Sequencing File:** http://ligacloud.ca/searchLibInfo?f=0&b=0&d=20171128-87CLooOOIQKI-SS

**Barcode:**
CTACTCTTTGCTATTCCGCTTAGCGTGGAGAAGAATGATCAGAAAACGTACCACGCCGGTGGTGGC

**Axis Name:** Tri-BN3-[840]

**IUPAC:** Gal(a1-3)[Fuc(a1-2)]Gal(b1-Sp

**CFG Name:** GalNAca1-3[Fuca1-2]Galb-Sp

**Common Name:** Tri-BN3

**Glytoucan ID:** G47548GA

**Compound Number:** Tri-BN3

**Maldi File:** 44.txt

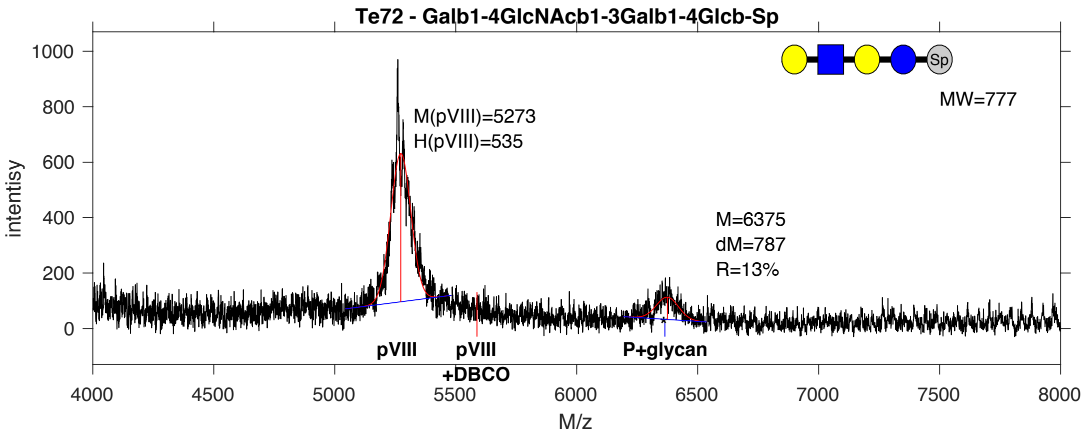


**SDB Number:** SDB88

**Sequencing File:** http://ligacloud.ca/searchLibInfo?f=0&b=0&d=20171128-87CLooOORWFR-SS

**Barcode:**
CTGCTGTTCGCGATTCCTCTAAGTGTAGAGAAGAATGACCAGAAGACCTACCACGCGGGCGGGGGG

**Axis Name:** LNnT-[350]

**IUPAC:** Gal(b1-4)GlcNAc(b1-3)Gal(b1-4)Glc(b1-Sp

**CFG Name:** Galb1-4GlcNAcb1-3Galb1-4Glcb-Sp

**Common Name:** LNnT

**Glytoucan ID:** G14139DE

**Compound Number:** Te72

**Maldi File:** 47.txt

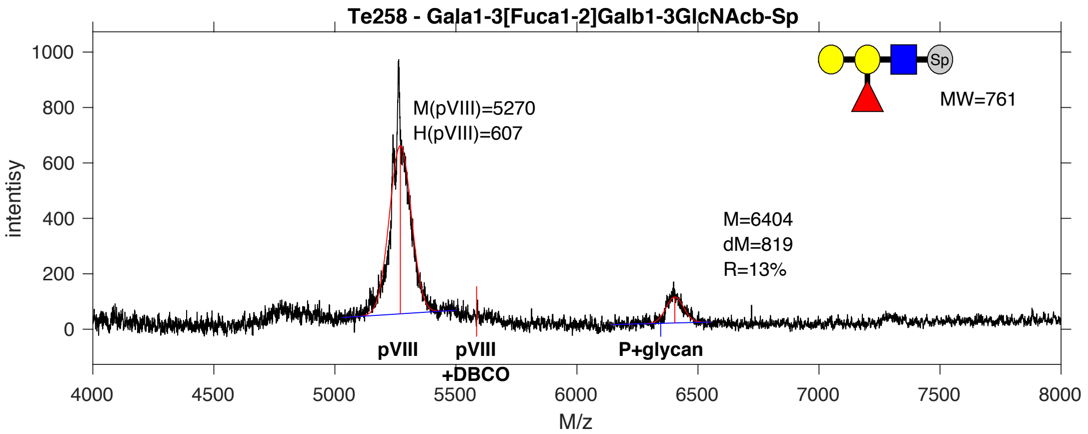


**SDB Number:** SDB105

**Sequencing File:** http://ligacloud.ca/searchLibInfo?f=0&b=0&d=NA

**Barcode:**
CTGCTTTTTGCTATTCCTCTTAGTGTCGAGAAGAATGATCAAAAAACTTACCACGCGGGTGGTGGA

**Axis Name:** B tetra type 1-[350]

**IUPAC:** Gal(a1-3)[Fuc(a1-2)]Gal(b1-3)GlcNAc(b1-Sp

**CFG Name:** Gala1-3[Fuca1-2]Galb1-3GlcNAcb-Sp

**Common Name:** B tetra type 1

**Glytoucan ID:** G31734OS

**Compound Number:** Te258

**Maldi File:** 48.txt

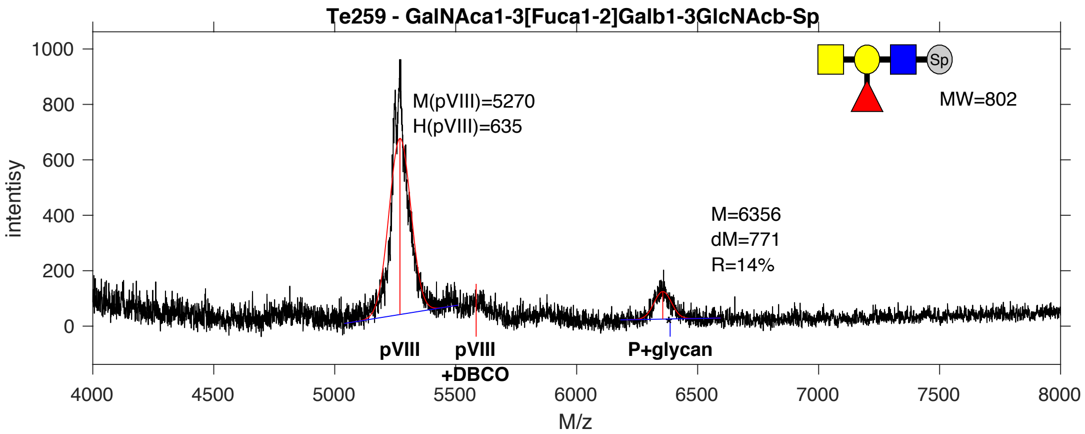


**SDB Number:** SDB106

**Sequencing File:** http://ligacloud.ca/searchLibInfo?f=0&b=0&d=NA

**Barcode:**
CTACTGTTCGCAATCCCGCTAAGTGTAGAAAAAAACGATCAGAAGACTTACCATGCTGGTGGTGGG

**Axis Name:** A tetra type 1-[380]

**IUPAC:** GalNAc(a1-3)[Fuc(a1-2)]Gal(b1-3)GlcNAc(b1-Sp

**CFG Name:** GalNAca1-3[Fuca1-2]Galb1-3GlcNAcb-Sp

**Common Name:** A tetra type 1

**Glytoucan ID:** G66163TI

**Compound Number:** Te259

**Maldi File:** 49.txt

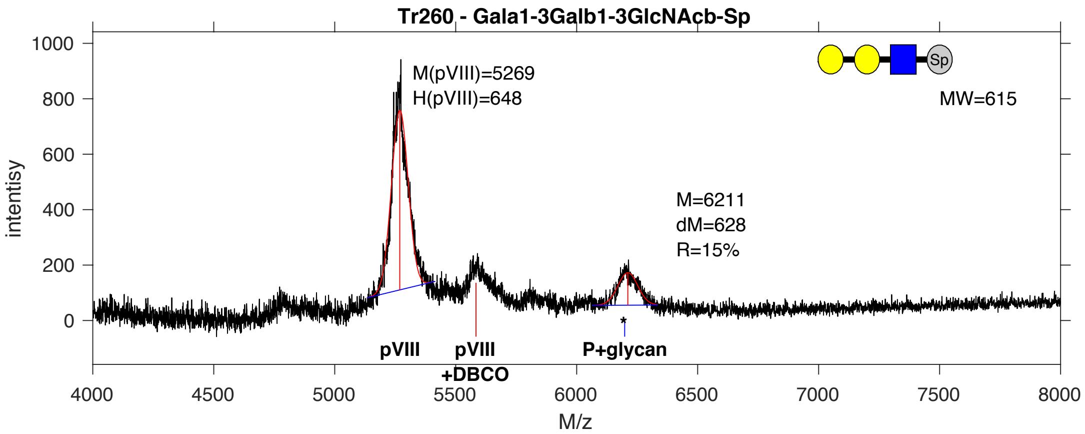


**SDB Number:** SDB112

**Sequencing File:** http://ligacloud.ca/searchLibInfo?f=0&b=0&d=NA

**Barcode:**
CTACTGTTCGCGATCCCACTGAGTGTAGAAAAAAATGACCAGAAGACGTATCACGCAGGCGGAGGC

**Axis Name:** Gala3-type1-[410]

**IUPAC:** Gal(a1-3)Gal(b1-3)GlcNAc(b1-Sp

**CFG Name:** Gala1-3Galb1-3GlcNAcb-Sp

**Common Name:** Gala3-type1

**Glytoucan ID:** G34356AA

**Compound Number:** Tr260

**Maldi File:** 52.txt

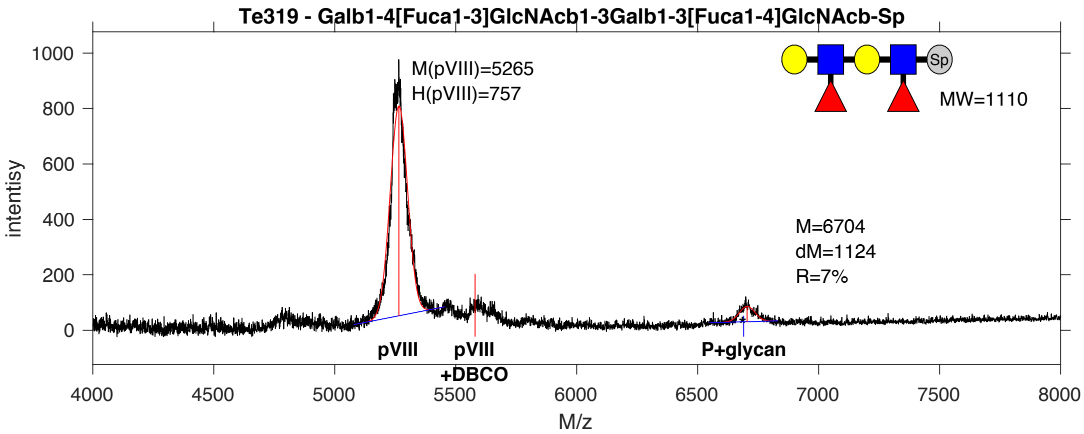


**SDB Number:** SDB113

**Sequencing File:** http://ligacloud.ca/searchLibInfo?f=0&b=0&d=20190703-87CLooORORN-DF

**Barcode:**
CTACTGTTTGCCATTCCCCTGAGTGTCGAAAAAAATGATCAGAAAACTTATCACGCAGGAGGGGGG

**Axis Name:** Lex-LeA-[190]

**IUPAC:** Gal(b1-4)[Fuc(a1-3)]GlcNAc(b1-3)Gal(b1-3)[Fuc(a1-4)]GlcNAc(b1-Sp

**CFG Name:** Galb1-4[Fuca1-3]GlcNAcb1-3Galb1-3[Fuca1-4]GlcNAcb-Sp

**Common Name:** Lex-LeA

**Glytoucan ID:** G59754DB

**Compound Number:** Te319

**Maldi File:** 53.txt

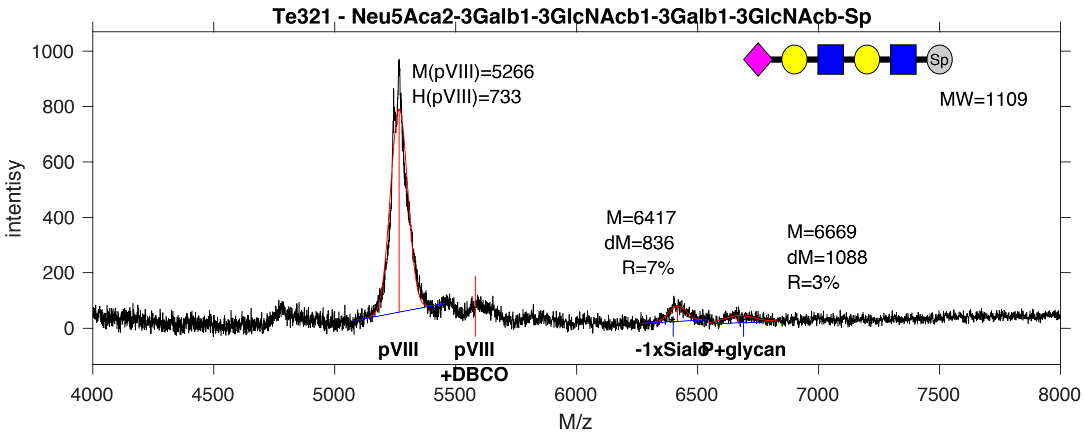


**SDB Number:** SDB115

**Sequencing File:** http://ligacloud.ca/searchLibInfo?f=0&b=0&d=20190703-87CLooOOWRCE-DF

**Barcode:**
CTACTGTTCGCTATCCCGCTGAGTGTCGAAAAAAATGACCAGAAAACGTATCATGCGGGGGGGGGT

**Axis Name:** 3'S-Di-Lec-[270]

**IUPAC:** Neu5Ac(a2-3)Gal(b1-3)GlcNAc(b1-3)Gal(b1-3)GlcNAc(b1-Sp

**CFG Name:** Neu5Aca2-3Galb1-3GlcNAcb1-3Galb1-3GlcNAcb-Sp

**Common Name:** 3'S-Di-Lec

**Glytoucan ID:** G11998FO

**Compound Number:** Te321

**Maldi File:** 54.txt

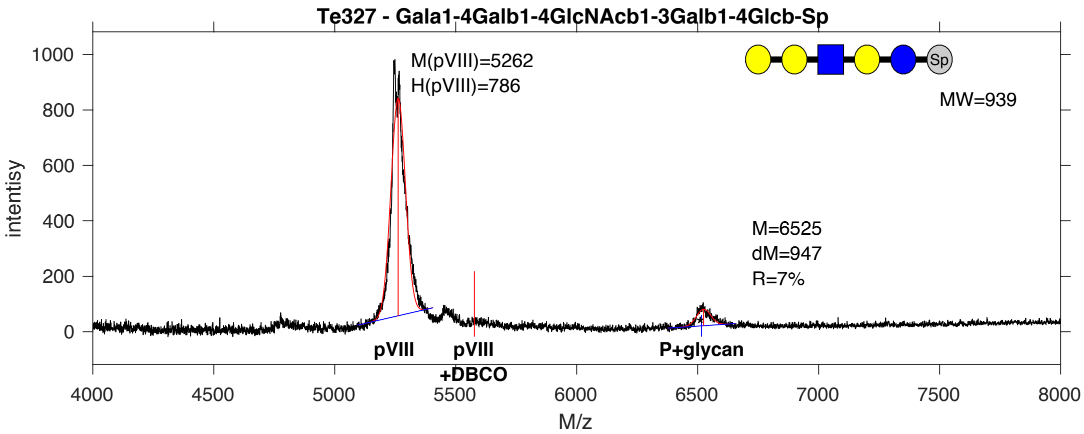


**SDB Number:** SDB118

**Sequencing File:** http://ligacloud.ca/searchLibInfo?f=0&b=0&d=20180530-87CLooOO1-1JM6

**Barcode:**
CTACTCTTCGCGATACCCCTCAGTGTCGAAAAGAATGATCAGAAGACTTATCATGCAGGTGGTGGT

**Axis Name:** P1 penta-[190]

**IUPAC:** Gal(a1-4)Gal(b1-4)GlcNAc(b1-3)Gal(b1-4)Glc(b1-Sp

**CFG Name:** Gala1-4Galb1-4GlcNAcb1-3Galb1-4Glcb-Sp

**Common Name:** P1 penta

**Glytoucan ID:** G12460DL

**Compound Number:** Te327

**Maldi File:** 55.txt

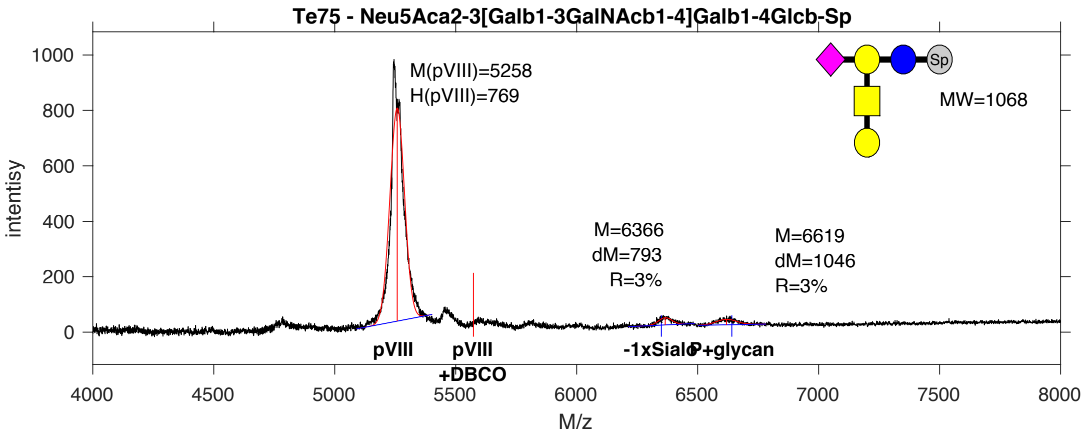


**SDB Number:** SDB121

**Sequencing File:** http://ligacloud.ca/searchLibInfo?f=0&b=0&d=20190703-87CLooOOJPTG-DF

**Barcode:**
CTGCTGTTTGCGATACCGCTTAGTGTGGAGAAAAACGATCAGAAGACCTATCATGCTGGGGGTGGA

**Axis Name:** GM1-[160]

**IUPAC:** Neu5Ac(a2-3)[Gal(b1-3)GalNAc(b1-4)]Gal(b1-4)Glc(b1-Sp

**CFG Name:** Neu5Aca2-3[Galb1-3GalNAcb1-4]Galb1-4Glcb-Sp

**Common Name:** GM1

**Glytoucan ID:** G40306MD

**Compound Number:** Te75

**Maldi File:** 56.txt

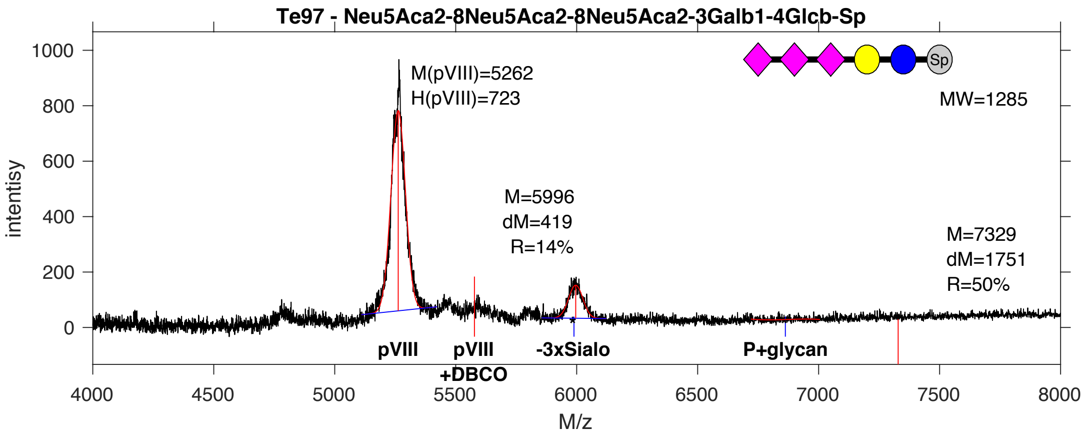


**SDB Number:** SDB124

**Sequencing File:** http://ligacloud.ca/searchLibInfo?f=0&b=0&d=20190703-87CLooOOQJTN-DF

**Barcode:**
CTACTGTTTGCGATCCCCCTCAGCGTTGAAAAAAACGATCAAAAAACATATCATGCGGGAGGCGGG

**Axis Name:** GT3-[380]

**IUPAC:** Neu5Ac(a2-8)Neu5Ac(a2-8)Neu5Ac(a2-3)Gal(b1-4)Glc(b1-Sp

**CFG Name:** Neu5Aca2-8Neu5Aca2-8Neu5Aca2-3Galb1-4Glcb-Sp

**Common Name:** GT3

**Glytoucan ID:** G93899SO

**Compound Number:** Te97

**Maldi File:** 58.txt

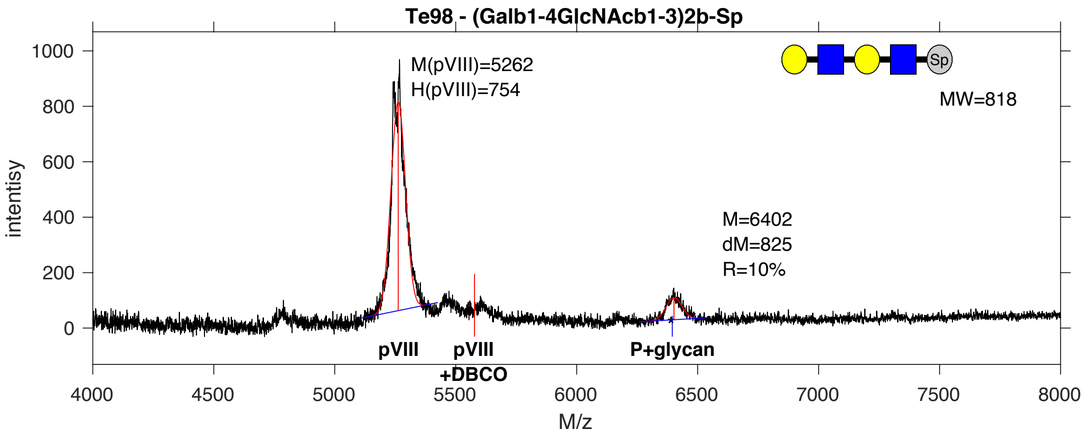


**SDB Number:** SDB149

**Sequencing File:** http://ligacloud.ca/searchLibInfo?f=0&b=0&d=NA

**Barcode:**
CTACTCTTTGCAATCCCGCTTAGCGTGGAGAAGAACGACCAGAAGACTTACCACGCGGGTGGTGGG

**Axis Name:** Di-LN-[270]

**IUPAC:** Gal(b1-4)GlcNAc(b1-3)Gal(b1-4)GlcNAc(b1-Sp

**CFG Name:** (Galb1-4GlcNAcb1-3)2b-Sp

**Common Name:** Di-LN

**Glytoucan ID:** G64789TL

**Compound Number:** Te98

**Maldi File:** 59.txt

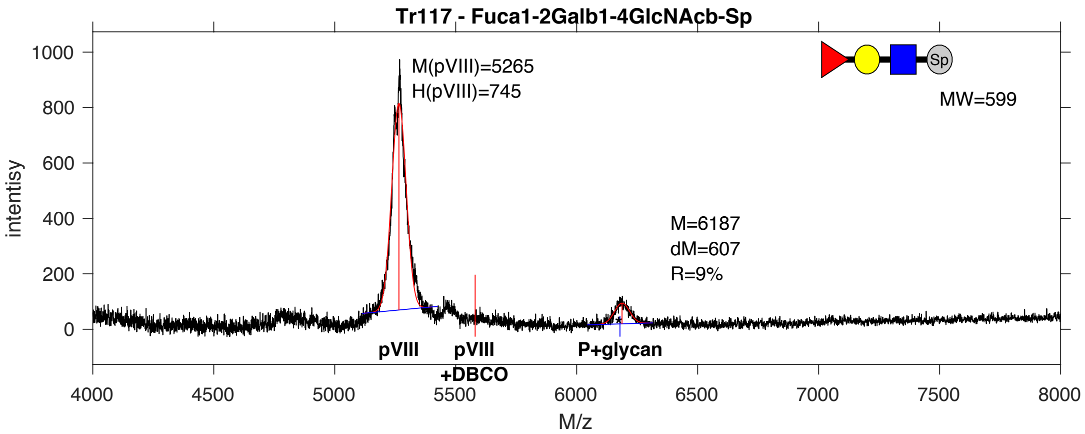


**SDB Number:** SDB152

**Sequencing File:** http://ligacloud.ca/searchLibInfo?f=0&b=0&d=NA

**Barcode:**
CTGCTTTTTGCTATACCTCTCAGCGTAGAAAAGAATGACCAGAAGACGTACCATGCCGGTGGGGGG

**Axis Name:** H-type2-[240]

**IUPAC:** Fuc(a1-2)Gal(b1-4)GlcNAc(b1-Sp

**CFG Name:** Fuca1-2Galb1-4GlcNAcb-Sp

**Common Name:** H-type2

**Glytoucan ID:** G17510DO

**Compound Number:** Tr117

**Maldi File:** 61.txt

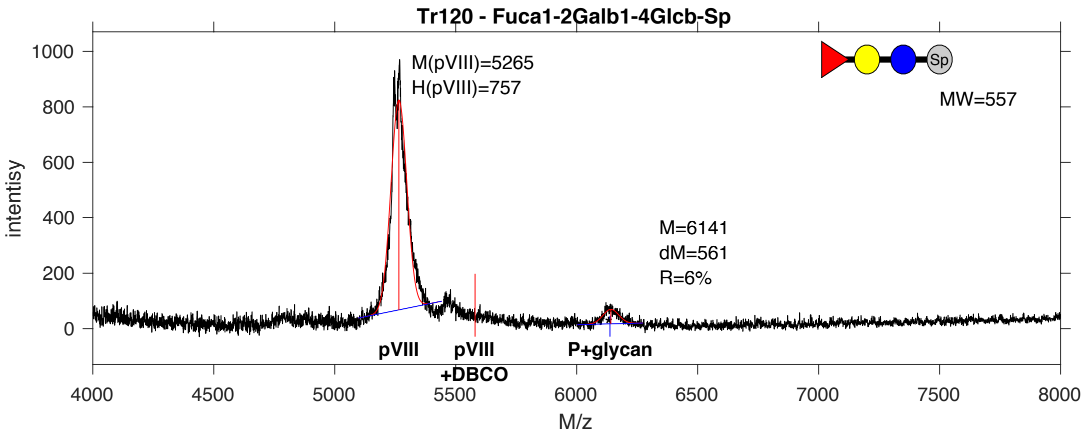


**SDB Number:** SDB153

**Sequencing File:** http://ligacloud.ca/searchLibInfo?f=0&b=0&d=NA

**Barcode:**
CTACTATTCGCTATTCCTCTGAGTGTCGAGAAAAACGATCAAAAGACCTATCATGCGGGTGGGGGT

**Axis Name:** 2’FL-[160]

**IUPAC:** Fuc(a1-2)Gal(b1-4)Glc(b1-Sp

**CFG Name:** Fuca1-2Galb1-4Glcb-Sp

**Common Name:** 2’FL

**Glytoucan ID:** G50519XG

**Compound Number:** Tr120

**Maldi File:** 62.txt

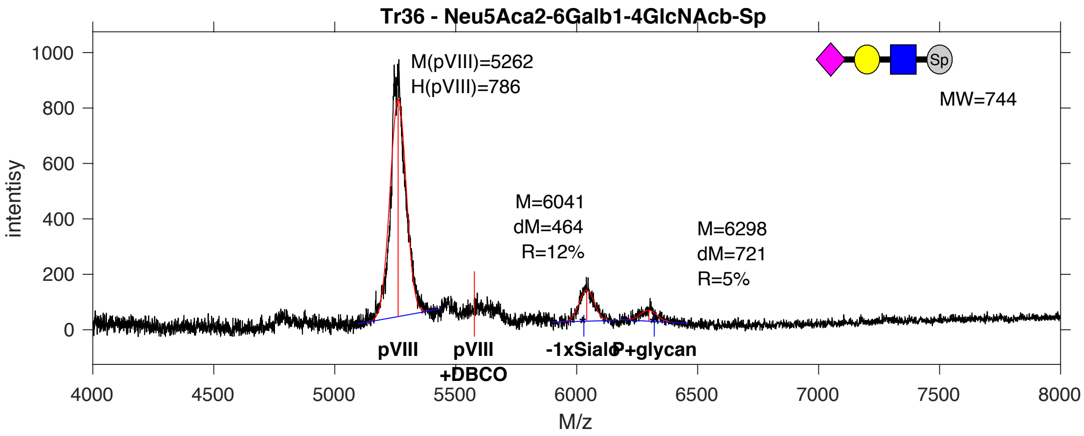


**SDB Number:** SDB160

**Sequencing File:** http://ligacloud.ca/searchLibInfo?f=0&b=0&d=NA

**Barcode:**
TTATTATTCGCAATTCCTTTAAGTGTCGAAAAAAATGACCAGAAAACTTACCATGCTGGAGGGGGC

**Axis Name:** 6’SLN-[460]

**IUPAC:** Neu5Ac(a2-6)Gal(b1-4)GlcNAc(b1-Sp

**CFG Name:** Neu5Aca2-6Galb1-4GlcNAcb-Sp

**Common Name:** 6’SLN

**Glytoucan ID:** G73578JC

**Compound Number:** Tr36

**Maldi File:** 63.txt

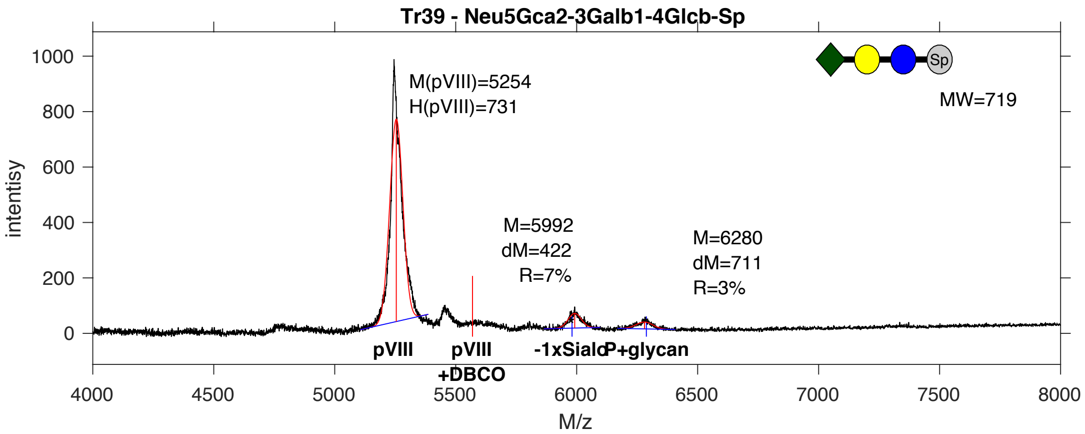


**SDB Number:** SDB161

**Sequencing File:** http://ligacloud.ca/searchLibInfo?f=0&b=0&d=NA

**Barcode:**
CTGCTATTCGCTATCCCCCTGAGTGTTGAGAAGAATGACCAAAAGACATATCACGCCGGGGGAGGG

**Axis Name:** 3’SL (Gc)-[240]

**IUPAC:** Neu5Gc(a2-3)Gal(b1-4)Glc(b1-Sp

**CFG Name:** Neu5Gca2-3Galb1-4Glcb-Sp

**Common Name:** 3’SL (Gc)

**Glytoucan ID:** G61420QM

**Compound Number:** Tr39

**Maldi File:** 64.txt

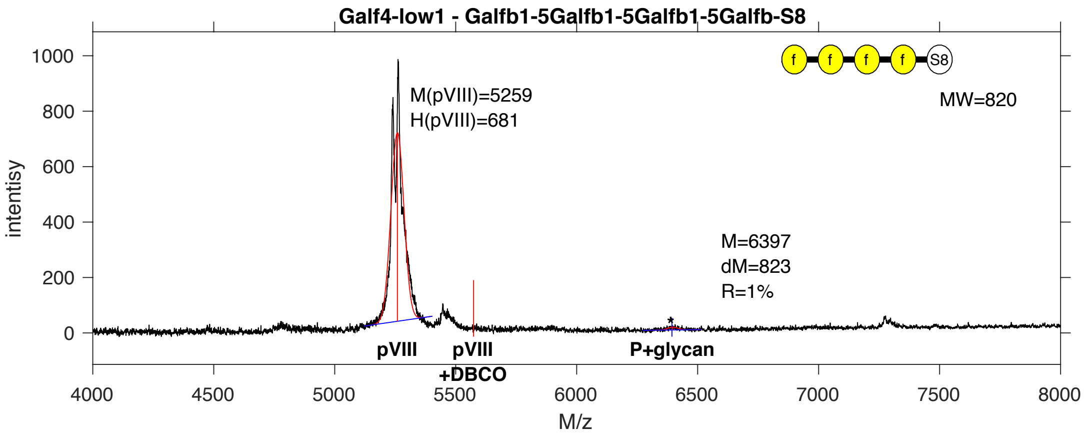


**SDB Number:** SDB125

**Sequencing File:** http://ligacloud.ca/searchLibInfo?f=0&b=0&d=20171128-87CLooOOEYOY-SS

**Barcode:**
CTGCTATTCGCTATCCCGCTGAGTGTAGAAAAAACGACCAGAAAACCTATCATGCCGGTGGCGGG

**Axis Name:** (Galf)4-[30]

**IUPAC:** Galf(b1-5)Galf(b1-5)Galf(b1-5)Galf(b1-S8

**CFG Name:** Galfb1-5Galfb1-5Galfb1-5Galfb-S8

**Common Name:** (Galf)4

**Glytoucan ID:** G92890KE

**Compound Number:** Galf4_low1

**Maldi File:** 65.txt

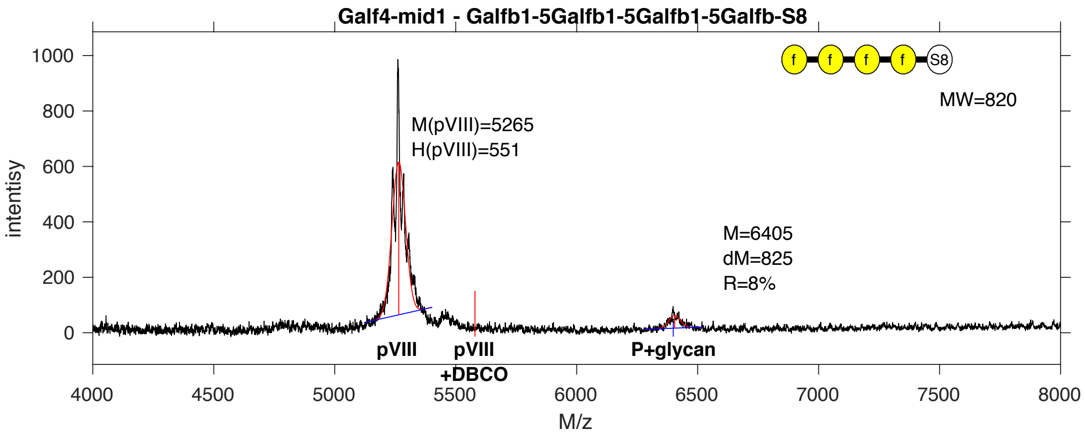


**SDB Number:** SDB126

**Sequencing File:** http://ligacloud.ca/searchLibInfo?f=0&b=0&d=20171128-87CLooOOXEGX-SS

**Barcode:**
CTGCTCTTTGCAATTCCGCTGAGTGTAGAAAAAAATGATCAGAAAACATACCATGCTGGTGGCGGA

**Axis Name:** (Galf)4-[220]

**IUPAC:** Galf(b1-5)Galf(b1-5)Galf(b1-5)Galf(b1-S8

**CFG Name:** Galfb1-5Galfb1-5Galfb1-5Galfb-S8

**Common Name:** (Galf)4

**Glytoucan ID:** G92890KE

**Compound Number:** Galf4-mid1

**Maldi File:** 66.txt

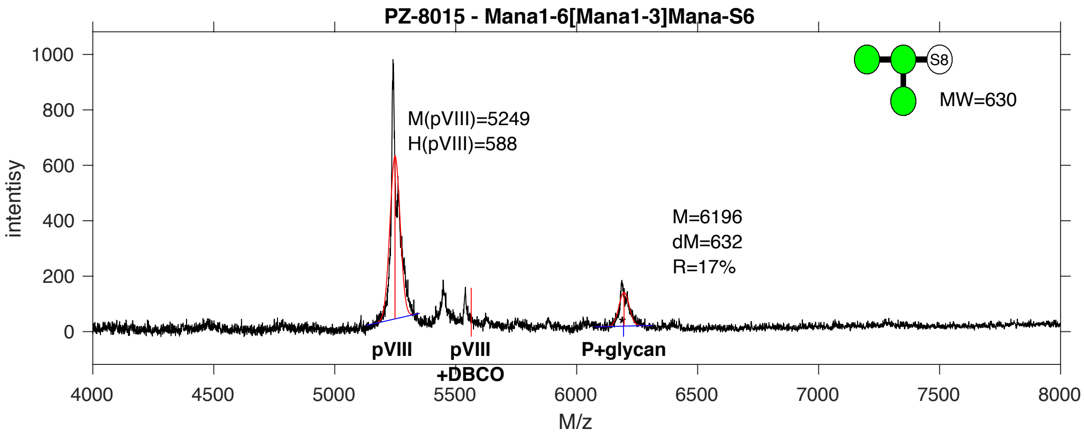


**SDB Number:** SDB127

**Sequencing File:** http://ligacloud.ca/searchLibInfo?f=0&b=0&d=20171128-87CLooOOZZNY-SS

**Barcode:**
CTGCTATTTGCCATCCCACTAAGTGTTGAGAAGAACGATCAGAAAACGTATCATGCAGGTGGTGGA

**Axis Name:** (Man)3-[460]

**IUPAC:** Man(a1-6)[Man(a1-3)]Man(a1-s6

**CFG Name:** Mana1-6[Mana1-3]Mana-S6

**Common Name:** (Man)3

**Glytoucan ID:** G80926OA

**Compound Number:** PZ-8015

**Maldi File:** 67.txt

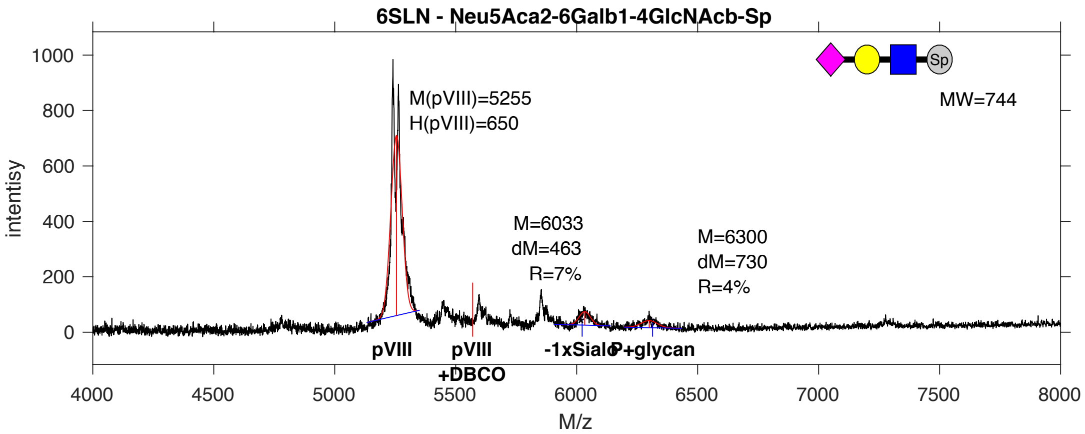


**SDB Number:** SDB128

**Sequencing File:** http://ligacloud.ca/searchLibInfo?f=0&b=0&d=20171128-87CLooOONWSB-SS

**Barcode:**
TTATTATTCGCAATTCCTTTAAGCGTTGAAAAGAATGACCAGAAAACATATCACGCTGGGGGAGGG

**Axis Name:** 6’SLN-[270]

**IUPAC:** Neu5Ac(a2-6)Gal(b1-4)GlcNAc(b1-Sp

**CFG Name:** Neu5Aca2-6Galb1-4GlcNAcb-Sp

**Common Name:** 6’SLN

**Glytoucan ID:** G73578JC

**Compound Number:** 6’SLN

**Maldi File:** 68.txt

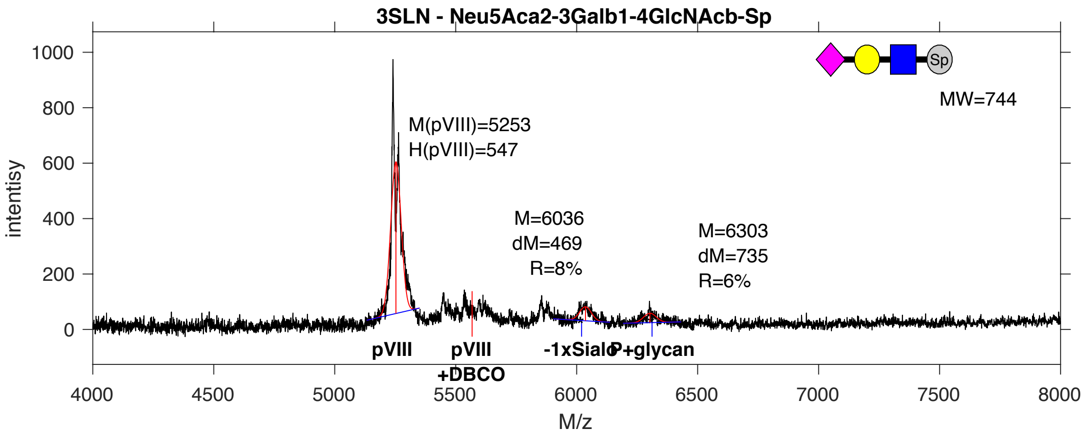


**SDB Number:** SDB129

**Sequencing File:** http://ligacloud.ca/searchLibInfo?f=0&b=0&d=20171128-87CLooOOOUNV-SS

**Barcode:**
CTTCTGTTCGCGATACCGCTGAGCGTCGAAAAGAATGATCAGAAAACTTATCATGCGGGGGGCGGG

**Axis Name:** 3’SLN-[350]

**IUPAC:** Neu5Ac(a2-3)Gal(b1-4)GlcNAc(b1-Sp

**CFG Name:** Neu5Aca2-3Galb1-4GlcNAcb-Sp

**Common Name:** 3’SLN

**Glytoucan ID:** G04307DP

**Compound Number:** 3’SLN

**Maldi File:** 69.txt

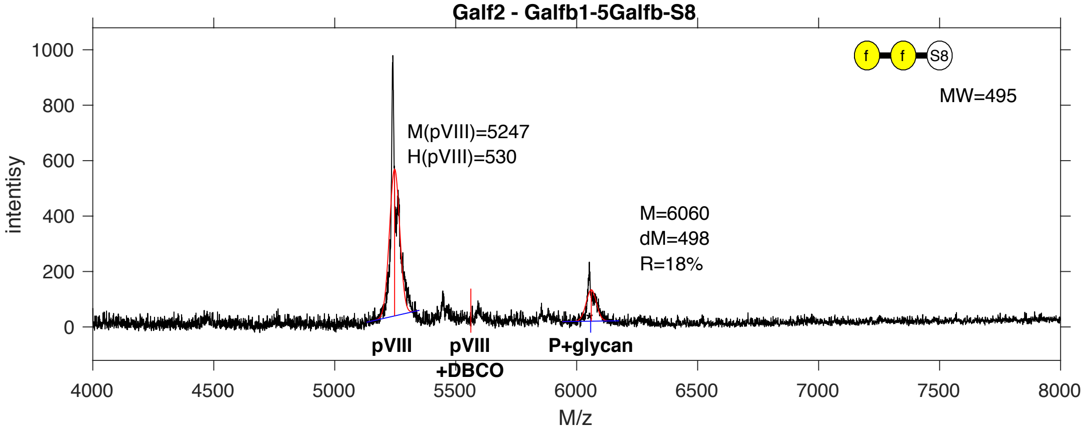


**SDB Number:** SDB131

**Sequencing File:** http://ligacloud.ca/searchLibInfo?f=0&b=0&d=NA

**Barcode:**
CTGCTGTTTGCGATTCCCCTGAGTGTCGAAAAGAATGACCAAAAGACGTACCATGCGGGGGGGGG

**Axis Name:** (Galf)2-[490]

**IUPAC:** Galf(b1-5)Galf(b1-S8

**CFG Name:** Galfb1-5Galfb-S8

**Common Name:** (Galf)2

**Glytoucan ID:** G98230LX

**Compound Number:** Galf2

**Maldi File:** 71.txt

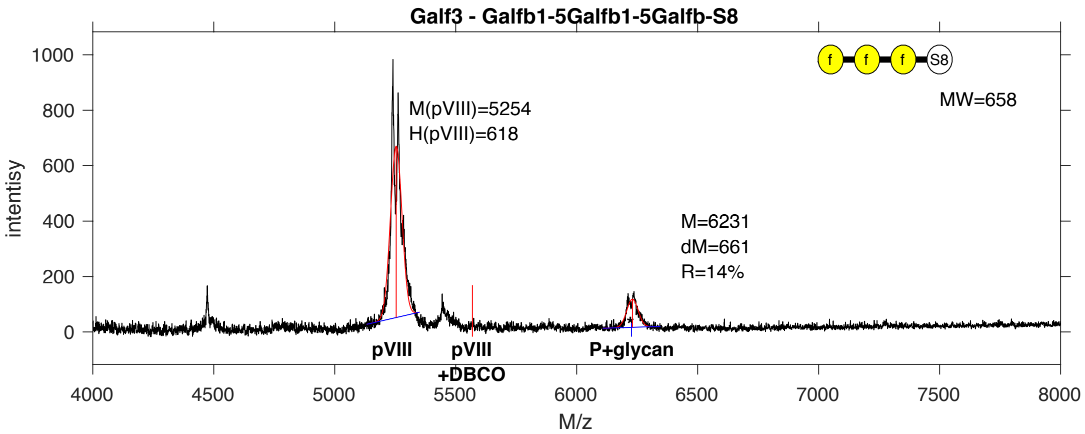


**SDB Number:** SDB132

**Sequencing File:** http://ligacloud.ca/searchLibInfo?f=0&b=0&d=20171128-87CLooOOTUHU-SS

**Barcode:**
CTACTGTTCGCAATACCTCTCAGCGTAGAAAAAAATGATCAAAAAACATATCACGCAGGTGGCGGA

**Axis Name:** (Galf)3-[380]

**IUPAC:** Galf(b1-5)Galf(b1-5)Galf(b1-S8

**CFG Name:** Galfb1-5Galfb1-5Galfb-S8

**Common Name:** (Galf)3

**Glytoucan ID:** G99590HR

**Compound Number:** Galf3

**Maldi File:** 72.txt

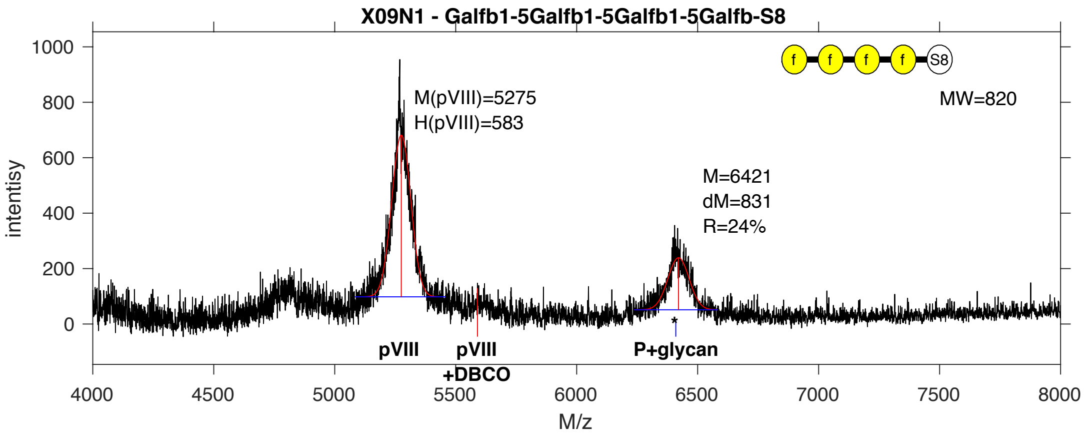


**SDB Number:** SDB2

**Sequencing File:** http://ligacloud.ca/searchLibInfo?f=0&b=0&d=20171128-87CLooOOCIWZ-SS

**Barcode:**
CTTCTATTCGCAATTCCGCTCAGTGTGGAGAAGAATGATCAGAAGACTTATCATGCGGGTGGAGGT

**Axis Name:** (Galf)4-[650]

**IUPAC:** Galf(b1-5)Galf(b1-5)Galf(b1-5)Galf(b1-S8

**CFG Name:** Galfb1-5Galfb1-5Galfb1-5Galfb-S8

**Common Name:** (Galf)4

**Glytoucan ID:** G92890KE

**Compound Number:** X09N1

**Maldi File:** 73.txt

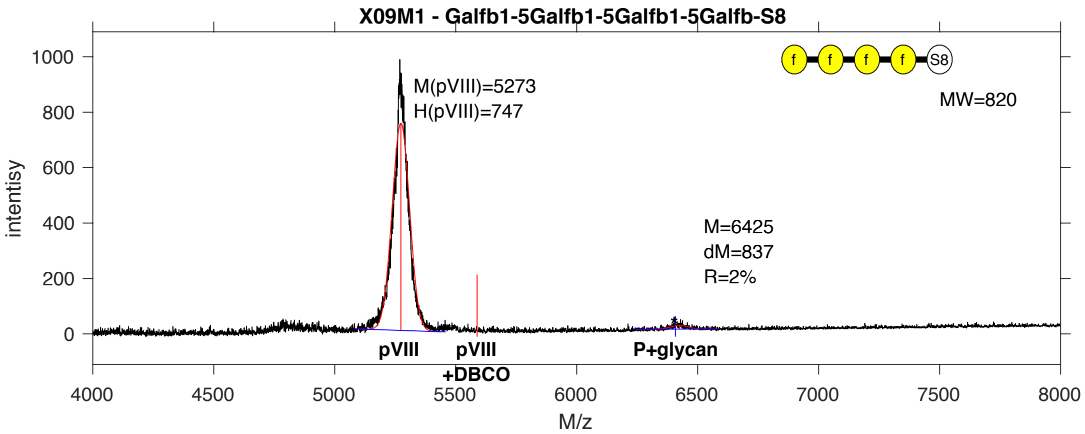


**SDB Number:** SDB8

**Sequencing File:** http://ligacloud.ca/searchLibInfo?f=0&b=0&d=20171128-87CLooOOOWCC-SS

**Barcode:**
CTGCTTTTTGCAATACCCCTCAGTGTGGAGAAGAATGATCAGAAGACTTATCATGCGGGTGGAGGT

**Axis Name:** (Galf)4-[50]

**IUPAC:** Galf(b1-5)Galf(b1-5)Galf(b1-5)Galf(b1-S8

**CFG Name:** Galfb1-5Galfb1-5Galfb1-5Galfb-S8

**Common Name:** (Galf)4

**Glytoucan ID:** G92890KE

**Compound Number:** X09M1

**Maldi File:** 75.txt

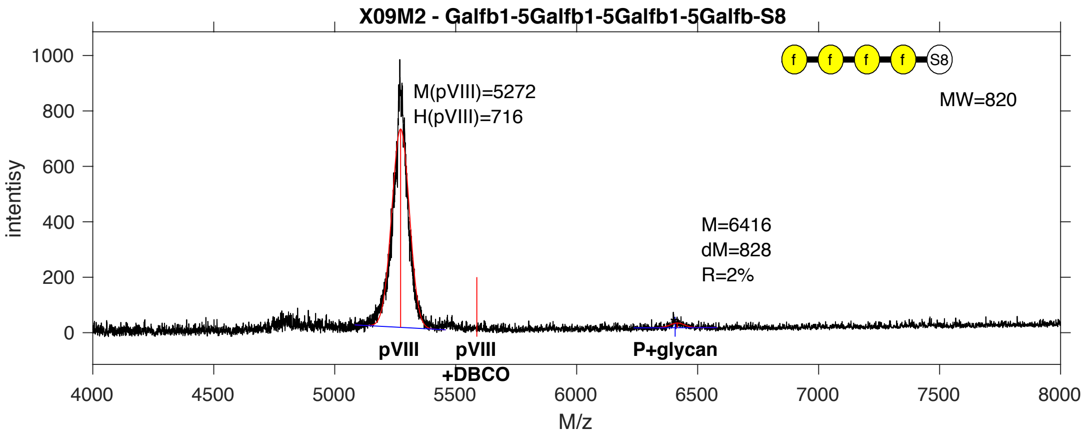


**SDB Number:** SDB9

**Sequencing File:** http://ligacloud.ca/searchLibInfo?f=0&b=0&d=20171128-87CLooOOKAQU-SS

**Barcode:**
CTTCTTTTGCAATTCCTCTAAGTGTGGAGAAGAATGATCAGAAGACTTATCATGCGGGTGGAGGT

**Axis Name:** (Galf)4-[50]

**IUPAC:** Galf(b1-5)Galf(b1-5)Galf(b1-5)Galf(b1-S8

**CFG Name:** Galfb1-5Galfb1-5Galfb1-5Galfb-S8

**Common Name:** (Galf)4

**Glytoucan ID:** G92890KE

**Compound Number:** X09M2

**Maldi File:** 76.txt

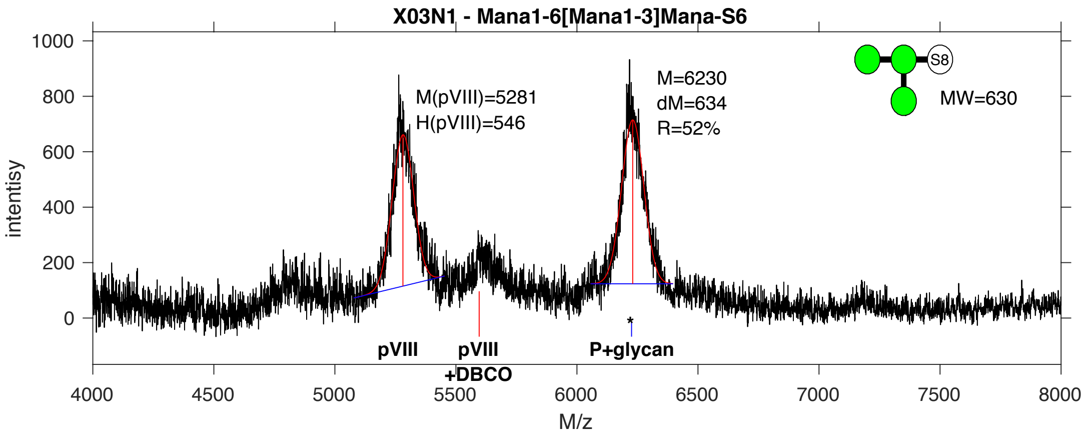


**SDB Number:** SDB12

**Sequencing File:** http://ligacloud.ca/searchLibInfo?f=0&b=0&d=20171128-87CLooOOHMWZ-SS

**Barcode:**
CTTCTGTTCGCGATACCTCTAAGTGTGGAGAAGAATGATCAGAAGACTTATCATGCGGGTGGAGGT

**Axis Name:** (Man)3-[1400]

**IUPAC:** Man(a1-6)[Man(a1-3)]Man(a1-S6

**CFG Name:** Mana1-6[Mana1-3]Mana-S6

**Common Name:** (Man)3

**Glytoucan ID:** G80926OA

**Compound Number:** X03N1

**Maldi File:** 79.txt

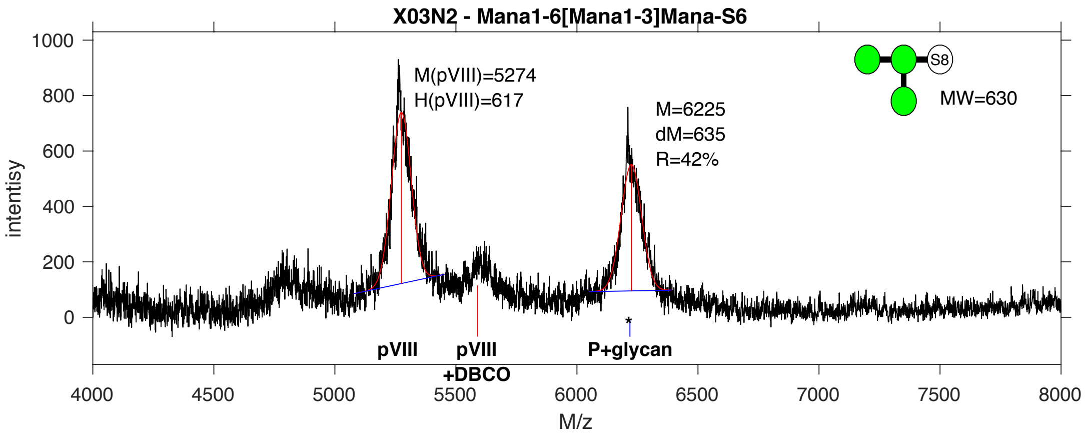


**SDB Number:** SDB13

**Sequencing File:** http://ligacloud.ca/searchLibInfo?f=0&b=0&d=20171128-87CLooOOMKLB-SS

**Barcode:**
CTACTTTTCGCAATTCCTCTGAGTGTGGAGAAGAATGATCAGAAGACTTATCATGCGGGTGGAGGT

**Axis Name:** (Man)3-[1160]

**IUPAC:** Man(a1-6)[Man(a1-3)]Man(a1-S6

**CFG Name:** Mana1-6[Mana1-3]Mana-S6

**Common Name:** (Man)3

**Glytoucan ID:** G80926OA

**Compound Number:** X03N2

**Maldi File:** 80.txt

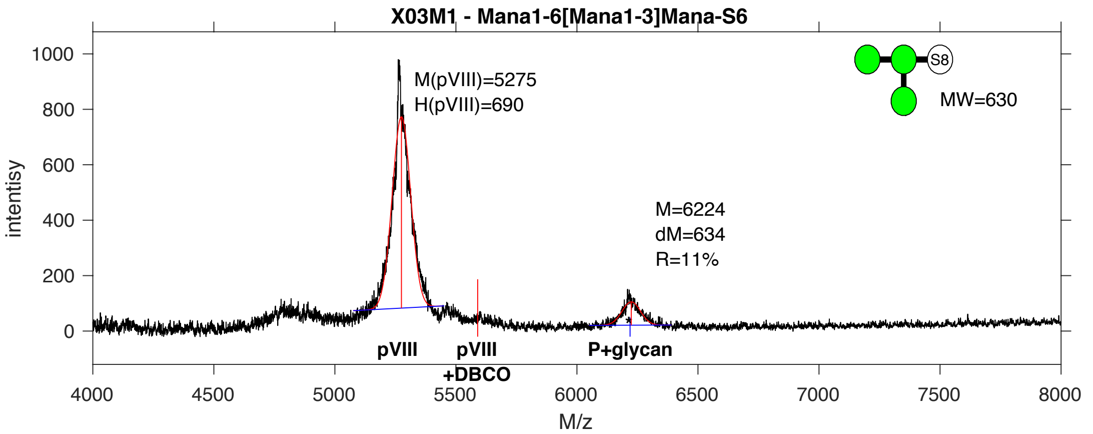


**SDB Number:** SDB15

**Sequencing File:** http://ligacloud.ca/searchLibInfo?f=0&b=0&d=20171128-87CLooOOBQBB-SS

**Barcode:**
CTGCTGTTCGCCATACCCCTTAGTGTGGAGAAGAATGATCAGAAGACTTATCATGCGGGTGGAGGT

**Axis Name:** (Man)3-[300]

**IUPAC:** Man(a1-6)[Man(a1-3)]Man(a1-S6

**CFG Name:** Mana1-6[Mana1-3]Mana-S6

**Common Name:** (Man)3

**Glytoucan ID:** G80926OA

**Compound Number:** X03M1

**Maldi File:** 81.txt

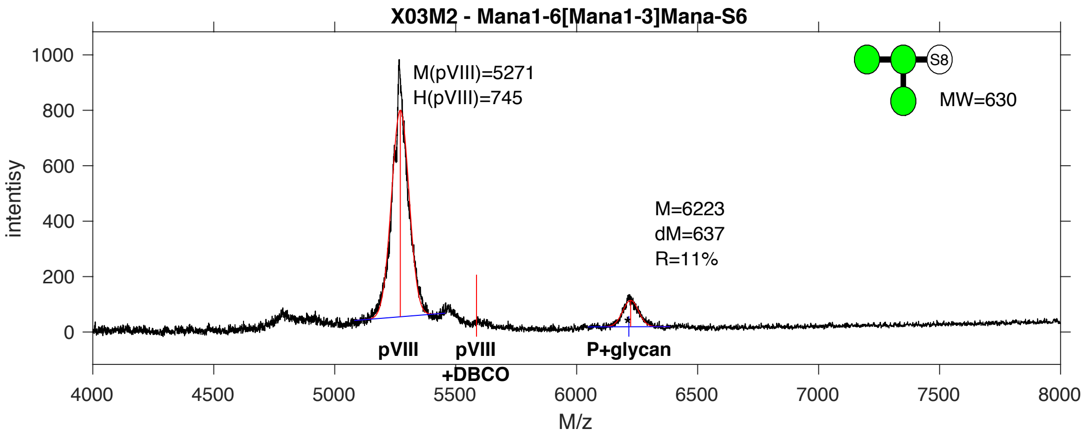


**SDB Number:** SDB16

**Sequencing File:** http://ligacloud.ca/searchLibInfo?f=0&b=0&d=20171128-87CLooOOIEEY-SS

**Barcode:**
CTGCTGTTCGCAATCCCGCTGAGTGTGGAGAAGAATGATCAGAAGACTTATCATGCGGGTGGAGGT

**Axis Name:** (Man)3-[300]

**IUPAC:** Man(a1-6)[Man(a1-3)]Man(a1-S6

**CFG Name:** Mana1-6[Mana1-3]Mana-S6

**Common Name:** (Man)3

**Glytoucan ID:** G80926OA

**Compound Number:** X03M2

**Maldi File:** 82.txt

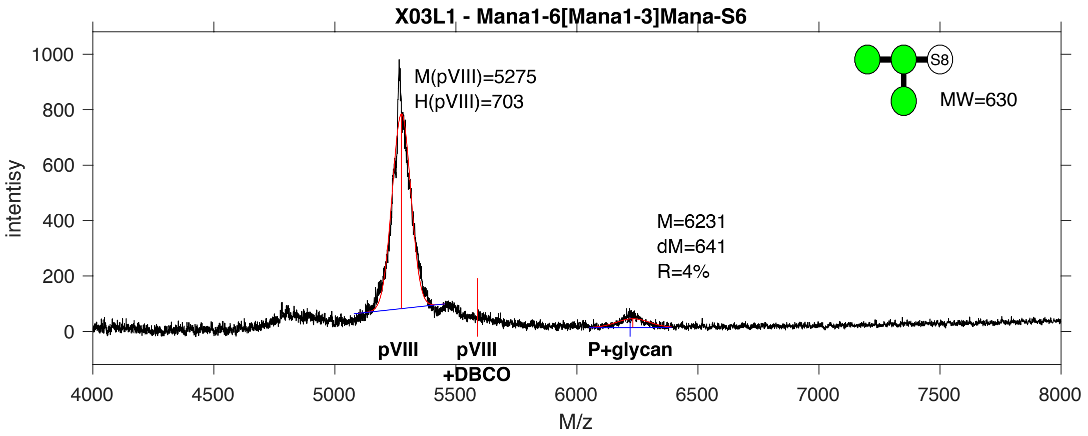


**SDB Number:** SDB17

**Sequencing File:** http://ligacloud.ca/searchLibInfo?f=0&b=0&d=20171128-87CLooOOKVTV-SS

**Barcode:**
CTACTCTTCGCGATTCCGCTTAGTGTGGAGAAGAATGATCAGAAGACTTATCATGCGGGTGGAGGT

**Axis Name:** (Man)3-[140]

**IUPAC:** Man(a1-6)[Man(a1-3)]Man(a1-S6

**CFG Name:** Mana1-6[Mana1-3]Mana-S6

**Common Name:** (Man)3

**Glytoucan ID:** G80926OA

**Compound Number:** X03L1

**Maldi File:** 83.txt

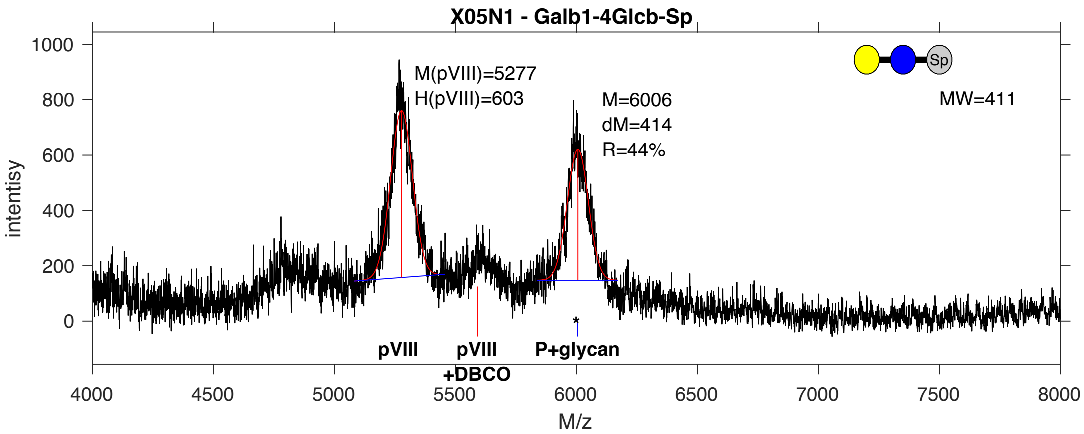


**SDB Number:** SDB20

**Sequencing File:** http://ligacloud.ca/searchLibInfo?f=0&b=0&d=20171128-87CLooOOXIFJ-SS

**Barcode:**
CTGCTCTTTGCCATCCCGCTTAGTGTGGAGAAGAATGATCAGAAGACTTATCATGCGGGTGGAGGT

**Axis Name:** Lac-[1190]

**IUPAC:** Gal(b1-4)Glc(b1-Sp

**CFG Name:** Galb1-4Glcb-Sp

**Common Name:** Lac

**Glytoucan ID:** G94144EF

**Compound Number:** X05N1

**Maldi File:** 84.txt

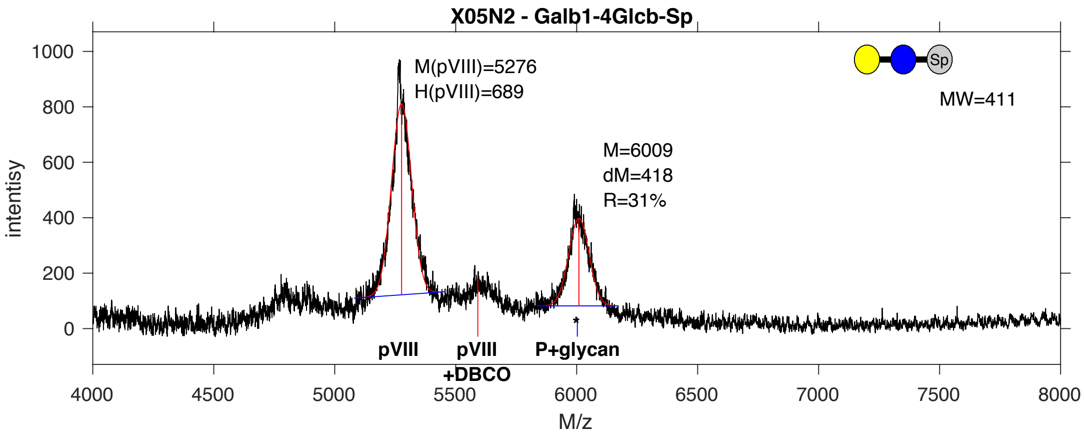


**SDB Number:** SDB21

**Sequencing File:** http://ligacloud.ca/searchLibInfo?f=0&b=0&d=20171128-87CLooOOEHIN-SS

**Barcode:**
CTACTCTTTGCAATTCCCCTTAGTGTGGAGAAGAATGATCAGAAGACTTATCATGCGGGTGGAGGT

**Axis Name:** Lac-[840]

**IUPAC:** Gal(b1-4)Glc(b1-Sp

**CFG Name:** Galb1-4Glcb-Sp

**Common Name:** Lac

**Glytoucan ID:** G94144EF

**Compound Number:** X05N2

**Maldi File:** 85.txt

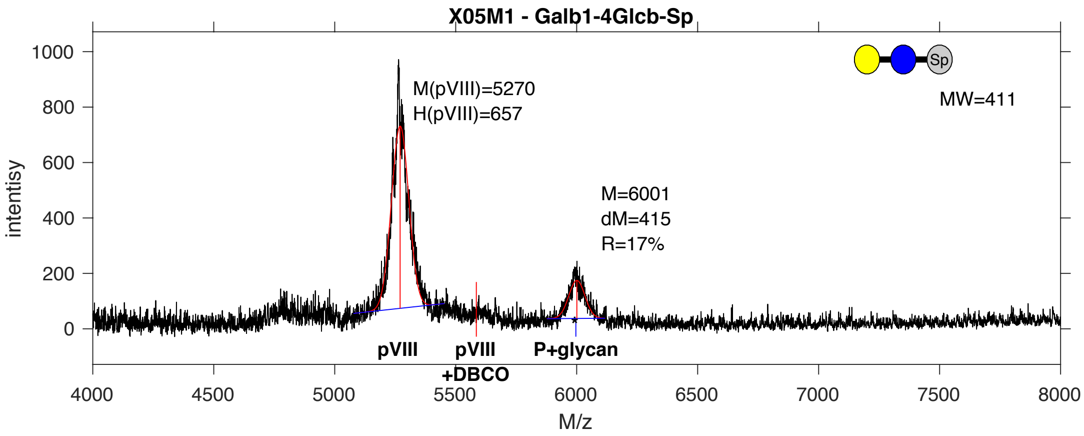


**SDB Number:** SDB22

**Sequencing File:** http://ligacloud.ca/searchLibInfo?f=0&b=0&d=20171128-87CLooOOTVNE-SS

**Barcode:**
CTACTGTTTGCTATCCCACTTAGTGTGGAGAAGAATGATCAGAAGACTTATCATGCGGGTGGAGGT

**Axis Name:** Lac-[490]

**IUPAC:** Gal(b1-4)Glc(b1-Sp

**CFG Name:** Galb1-4Glcb-Sp

**Common Name:** Lac

**Glytoucan ID:** G94144EF

**Compound Number:** X05M1

**Maldi File:** 86.txt

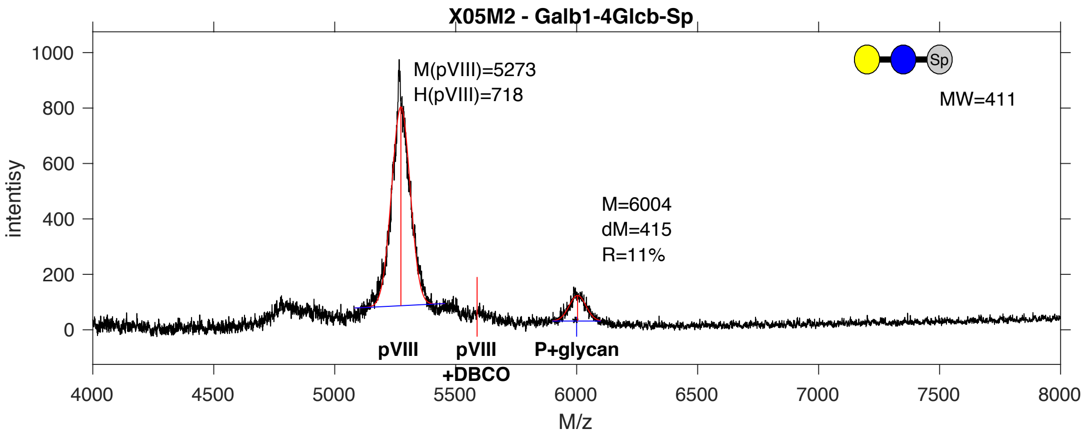


**SDB Number:** SDB23

**Sequencing File:** http://ligacloud.ca/searchLibInfo?f=0&b=0&d=20171128-87CLooOONHEY-SS

**Barcode:**
CTGCTCTTTGCAATACCTCTTAGTGTGGAGAAGAATGATCAGAAGACTTATCATGCGGGTGGAGGT

**Axis Name:** Lac-[320]

**IUPAC:** Gal(b1-4)Glc(b1-Sp

**CFG Name:** Galb1-4Glcb-Sp

**Common Name:** Lac

**Glytoucan ID:** G94144EF

**Compound Number:** X05M2

**Maldi File:** 87.txt

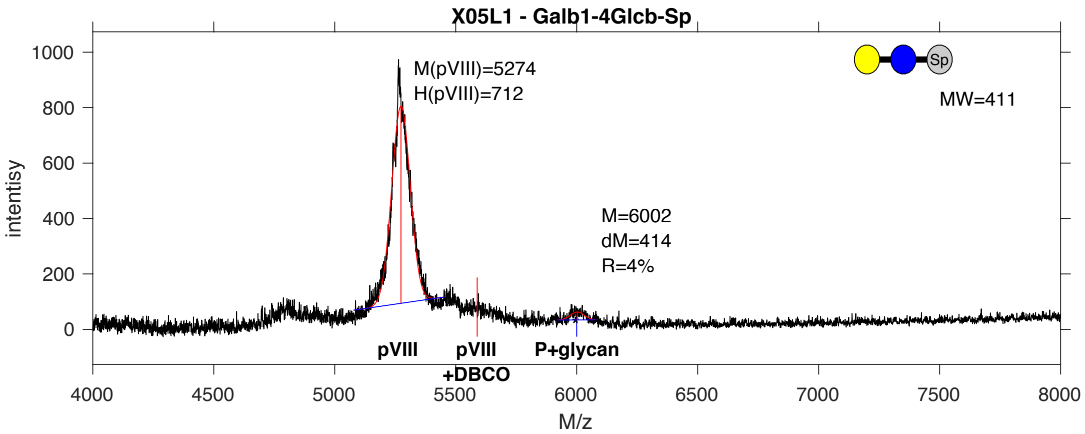


**SDB Number:** SDB24

**Sequencing File:** http://ligacloud.ca/searchLibInfo?f=0&b=0&d=20171128-87CLooOOJDTS-SS

**Barcode:**
CTACTATTCGCGATCCCGCTCAGTGTGGAGAAGAATGATCAGAAGACTTATCATGCGGGTGGAGGT

**Axis Name:** Lac-[140]

**IUPAC:** Gal(b1-4)Glc(b1-Sp

**CFG Name:** Galb1-4Glcb-Sp

**Common Name:** Lac

**Glytoucan ID:** G94144EF

**Compound Number:** X05L1

**Maldi File:** 88.txt

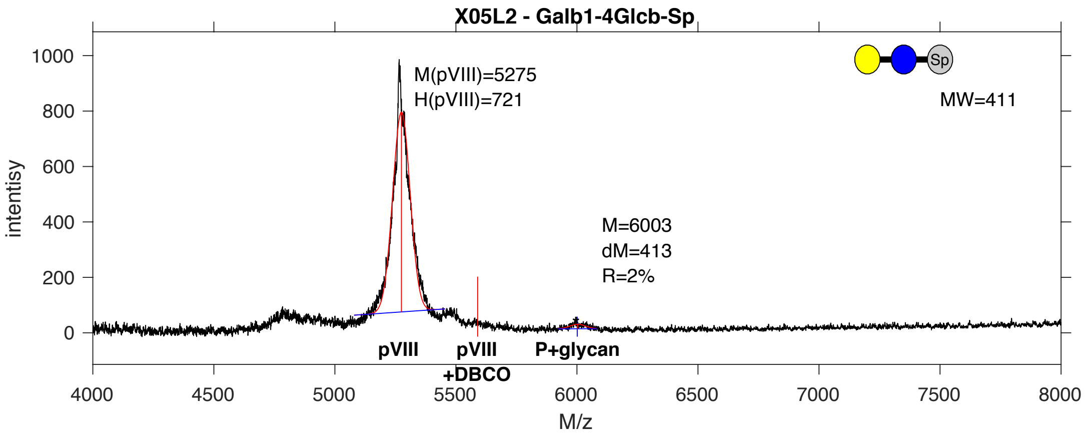


**SDB Number:** SDB25

**Sequencing File:** http://ligacloud.ca/searchLibInfo?f=0&b=0&d=20171128-87CLooOOQQEH-SS

**Barcode:**
CTGCTTTTCGCAATACCTCTAAGTGTGGAGAAGAATGATCAGAAGACTTATCATGCGGGTGGAGGT

**Axis Name:** Lac-[80]

**IUPAC:** Gal(b1-4)Glc(b1-Sp

**CFG Name:** Galb1-4Glcb-Sp

**Common Name:** Lac

**Glytoucan ID:** G94144EF

**Compound Number:** X05L2

**Maldi File:** 89.txt

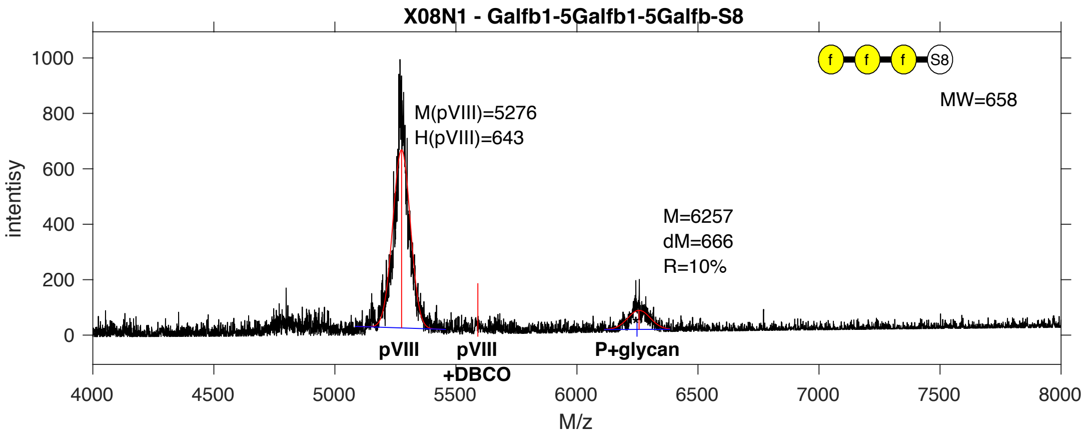


**SDB Number:** SDB26

**Sequencing File:** http://ligacloud.ca/searchLibInfo?f=0&b=0&d=NA

**Barcode:**
CTGCTATTCGCTATCCCACTCAGTGTGGAGAAGAATGATCAGAAGACTTATCATGCGGGTGGAGGT

**Axis Name:** (Galf)3-[270]

**IUPAC:** Galf(b1-5)Galf(b1-5)Galf(b1-S8

**CFG Name:** Galfb1-5Galfb1-5Galfb-S8

**Common Name:** (Galf)3

**Glytoucan ID:** G99590HR

**Compound Number:** X08N1

**Maldi File:** 90.txt

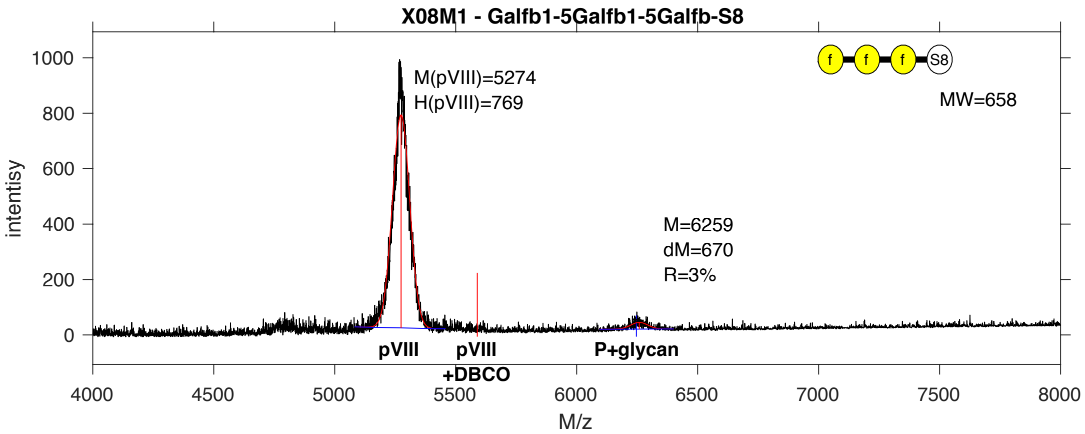


**SDB Number:** SDB28

**Sequencing File:** http://ligacloud.ca/searchLibInfo?f=0&b=0&d=20171128-87CLooOOIJYY-SS

**Barcode:**
CTACTCTTCGCCATTCCACTGAGTGTGGAGAAGAATGATCAGAAGACTTATCATGCGGGTGGAGGT

**Axis Name:** (Galf)3-[80]

**IUPAC:** Galf(b1-5)Galf(b1-5)Galf(b1-S8

**CFG Name:** Galfb1-5Galfb1-5Galfb-S8

**Common Name:** (Galf)3

**Glytoucan ID:** G99590HR

**Compound Number:** X08M1

**Maldi File:** 92.txt

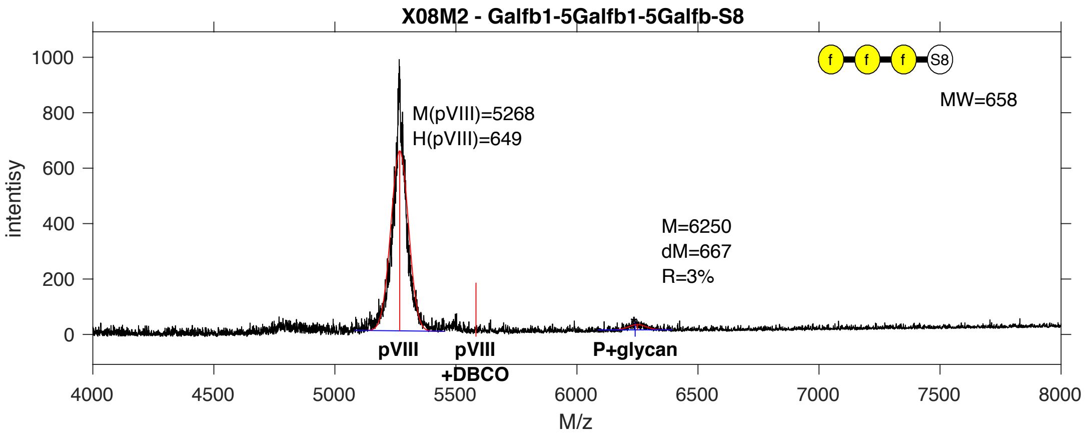


**SDB Number:** SDB29

**Sequencing File:** http://ligacloud.ca/searchLibInfo?f=0&b=0&d=20171128-87CLooOOSNHI-SS

**Barcode:**
CTGCTATTTGCGATCCCGCTGAGTGTGGAGAAGAATGATCAGAAGACTTATCATGCGGGTGGAGGT

**Axis Name:** (Galf)3-[80]

**IUPAC:** Galf(b1-5)Galf(b1-5)Galf(b1-S8

**CFG Name:** Galfb1-5Galfb1-5Galfb-S8

**Common Name:** (Galf)3

**Glytoucan ID:** G99590HR

**Compound Number:** X08M2

**Maldi File:** 93.txt

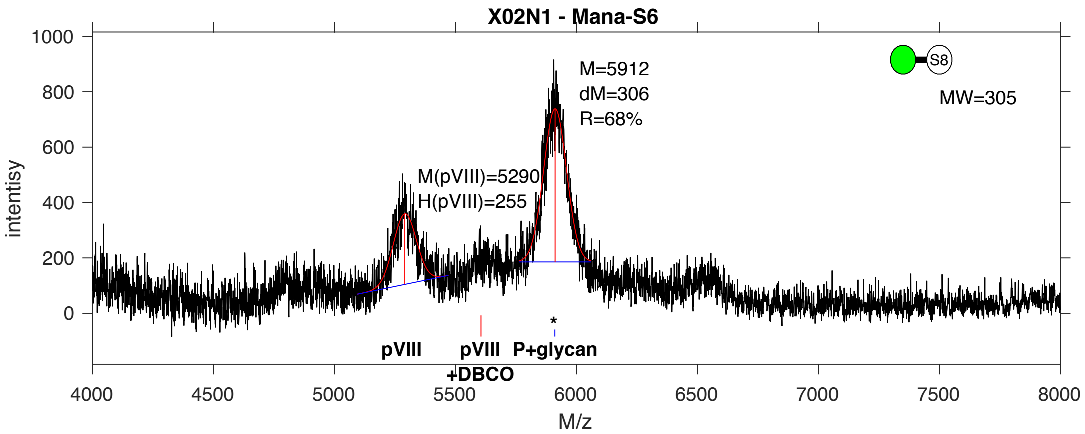


**SDB Number:** SDB48

**Sequencing File:** http://ligacloud.ca/searchLibInfo?f=0&b=0&d=20171128-87CLooOOONEV-SS

**Barcode:**
TTATTATTCGCAATTCCTTTAAGCGTAGAGAAAAACGACCAGAAGACCTATCACGCGGGAGGAGGT

**Axis Name:** aMan-[1860]

**IUPAC:** Man(a1-S6

**CFG Name:** Mana-S6

**Common Name:** aMan

**Glytoucan ID:** G19165TD

**Compound Number:** X02N1

**Maldi File:** 94.txt

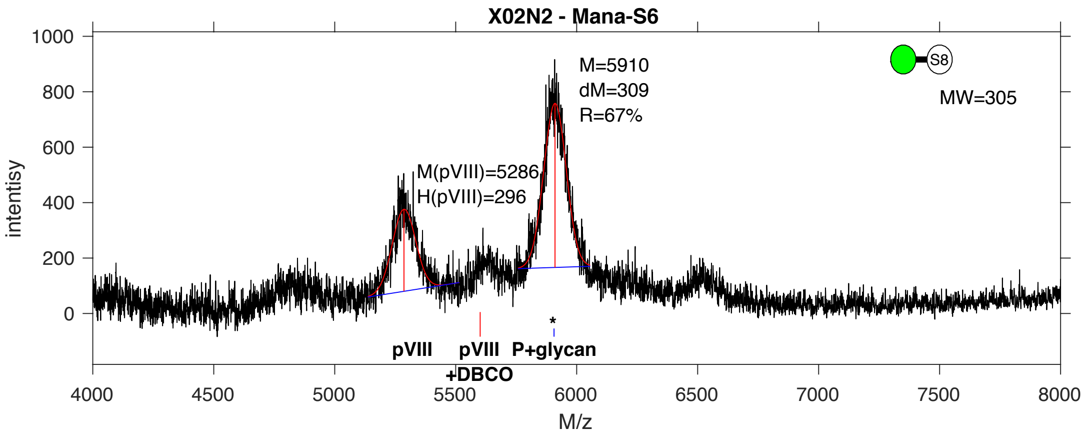


**SDB Number:** SDB49

**Sequencing File:** http://ligacloud.ca/searchLibInfo?f=0&b=0&d=20171128-87CLooOOIWYU-SS

**Barcode:**
CTACTGTTCGCTATCCCGCTGAGTGTGGAAAAGAATGATCAGAAGACTTACCACGCTGGTGGTGGG

**Axis Name:** aMan-[1810]

**IUPAC:** Man(a1-S6

**CFG Name:** Mana-S6

**Common Name:** aMan

**Glytoucan ID:** G19165TD

**Compound Number:** X02N2

**Maldi File:** 95.txt

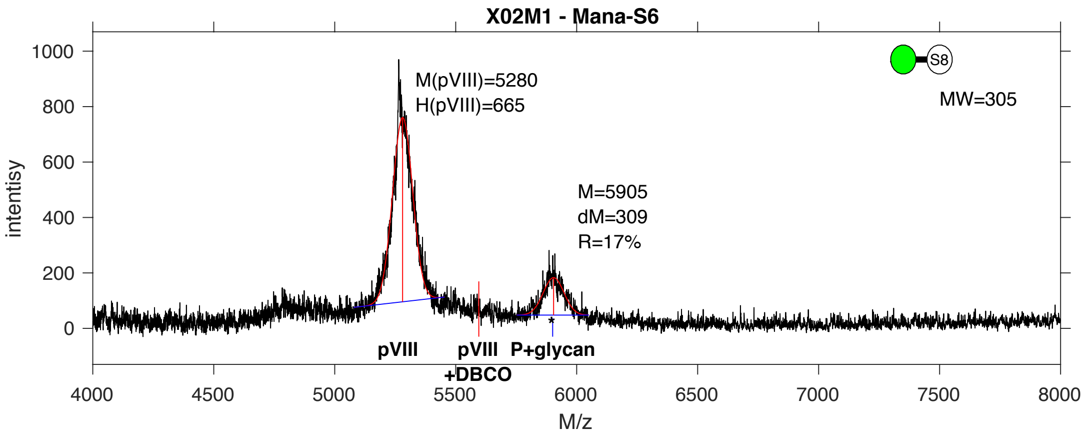


**SDB Number:** SDB52

**Sequencing File:** http://ligacloud.ca/searchLibInfo?f=0&b=0&d=20171128-87CLooOORRZY-SS

**Barcode:**
CTACTGTTTGCTATCCCGCTGAGTGTAGAGAAGAATGACCAAAAGACATATCATGCGGGAGGAGGT

**Axis Name:** aMan-[460]

**IUPAC:** Man(a1-S6

**CFG Name:** Mana-S6

**Common Name:** aMan

**Glytoucan ID:** G19165TD

**Compound Number:** X02M1

**Maldi File:** 100.txt

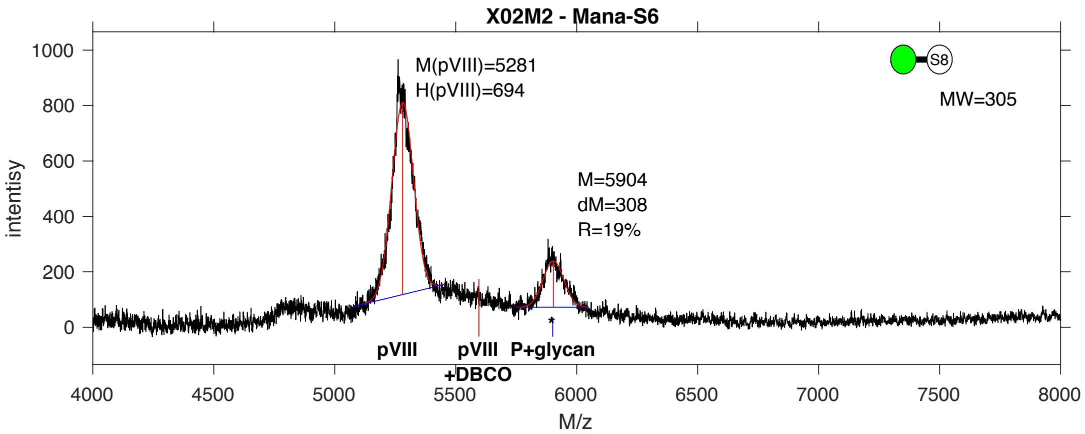


**SDB Number:** SDB53

**Sequencing File:** http://ligacloud.ca/searchLibInfo?f=0&b=0&d=20171128-87CLooOOWGUJ-SS

**Barcode:**
CTGCTGTTTGCGATCCCGCTCAGTGTGGAGAAAAATGACCAAAAGACCTATCATGCTGGAGGGGGT

**Axis Name:** aMan-[540]

**IUPAC:** Man(a1-S6

**CFG Name:** Mana-S6

**Common Name:** aMan

**Glytoucan ID:** G19165TD

**Compound Number:** X02M2

**Maldi File:** 101.txt

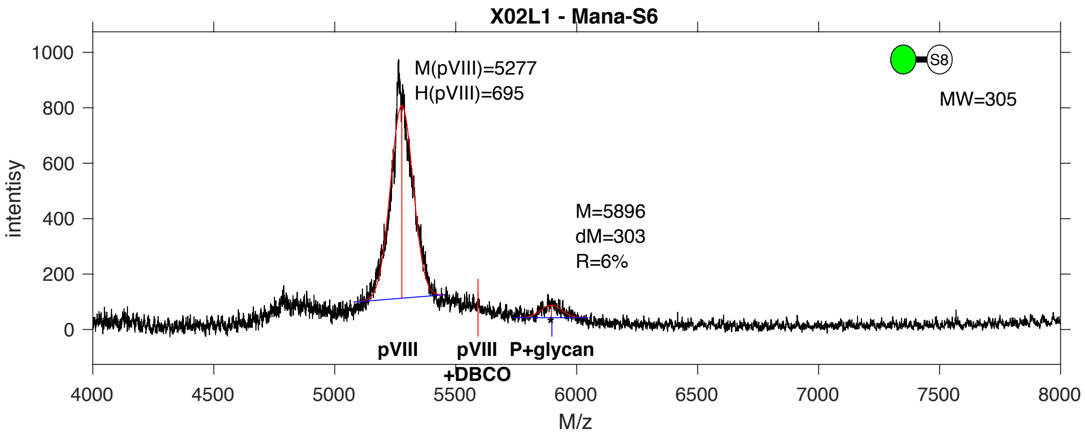


**SDB Number:** SDB54

**Sequencing File:** http://ligacloud.ca/searchLibInfo?f=0&b=0&d=20171128-87CLooOOBRPI-SS

**Barcode:**
TTATTATTCGCAATTCCTTTAAGTGTCGAAAAAAACGATCAGAAGACCTATCACGCAGGCGGGGGC

**Axis Name:** aMan-[160]

**IUPAC:** Man(a1-S6

**CFG Name:** Mana-S6

**Common Name:** aMan

**Glytoucan ID:** G19165TD

**Compound Number:** X02L1

**Maldi File:** 102.txt

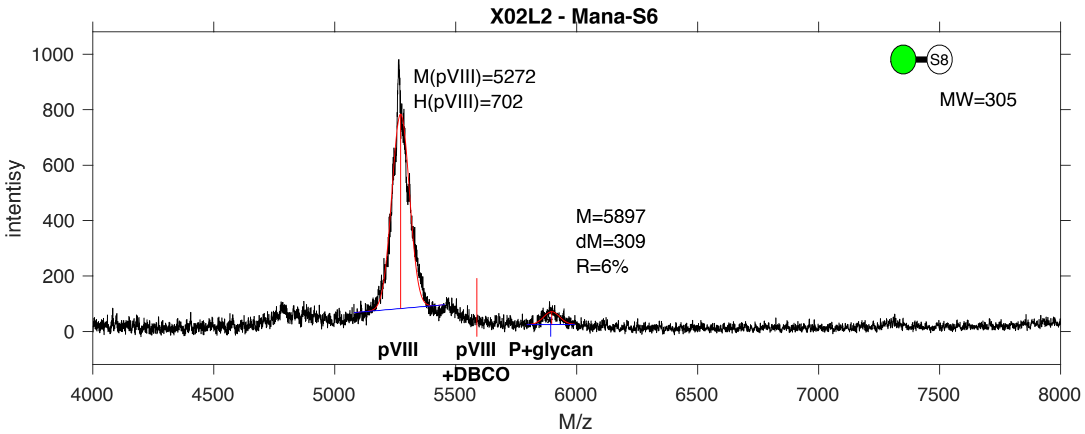


**SDB Number:** SDB57

**Sequencing File:** http://ligacloud.ca/searchLibInfo?f=0&b=0&d=20171128-87CLooOORPVC-SS

**Barcode:**
TTATTATTCGCAATTCCTTTAAGCGTCGAGAAGAACGACCAAAAAACCTATCACGCAGGTGGCGGG

**Axis Name:** aMan-[160]

**IUPAC:** Man(a1-S6

**CFG Name:** Mana-S6

**Common Name:** aMan

**Glytoucan ID:** G19165TD

**Compound Number:** X02L2

**Maldi File:** 103.txt

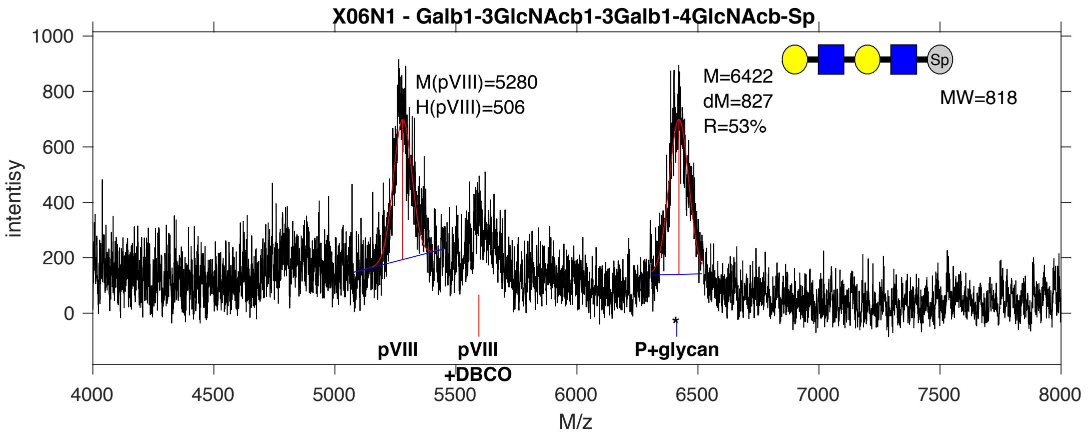


**SDB Number:** SDB58

**Sequencing File:** http://ligacloud.ca/searchLibInfo?f=0&b=0&d=20171128-87CLooOOQAMP-SS

**Barcode:**
CTACTCTTCGCTATACCCCTCAGCGTGGAGAAAAACGACCAAAAGACCTACCATGCAGGTGGTGGC

**Axis Name:** LNT-Nac-[1490]

**IUPAC:** Gal(b1-3)GlcNAc(b1-3)Gal(b1-4)GlcNAc(b1-Sp

**CFG Name:** Galb1-3GlcNAcb1-3Galb1-4GlcNAcb-Sp

**Common Name:** LNT-Nac

**Glytoucan ID:** G09681LD

**Compound Number:** X06N1

**Maldi File:** 104.txt

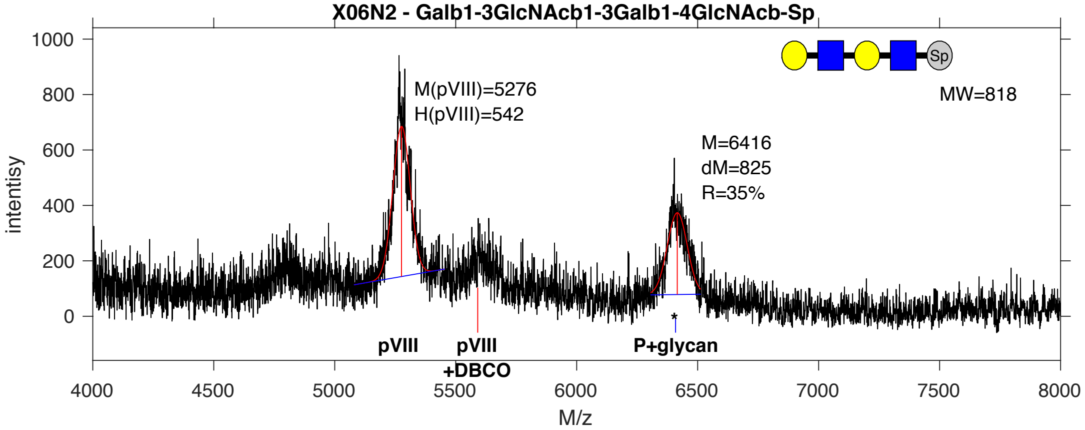


**SDB Number:** SDB67

**Sequencing File:** http://ligacloud.ca/searchLibInfo?f=0&b=0&d=20171128-87CLooOOSXNG-SS

**Barcode:**
CTTCTGTTCGCTATACCTCTCAGCGTGGAGAAGAATGACCAGAAAACTTATCACGCAGGAGGTGGA

**Axis Name:** LNT-Nac-[1030]

**IUPAC:** Gal(b1-3)GlcNAc(b1-3)Gal(b1-4)GlcNAc(b1-Sp

**CFG Name:** Galb1-3GlcNAcb1-3Galb1-4GlcNAcb-Sp

**Common Name:** LNT-Nac

**Glytoucan ID:** G09681LD

**Compound Number:** X06N2

**Maldi File:** 105.txt

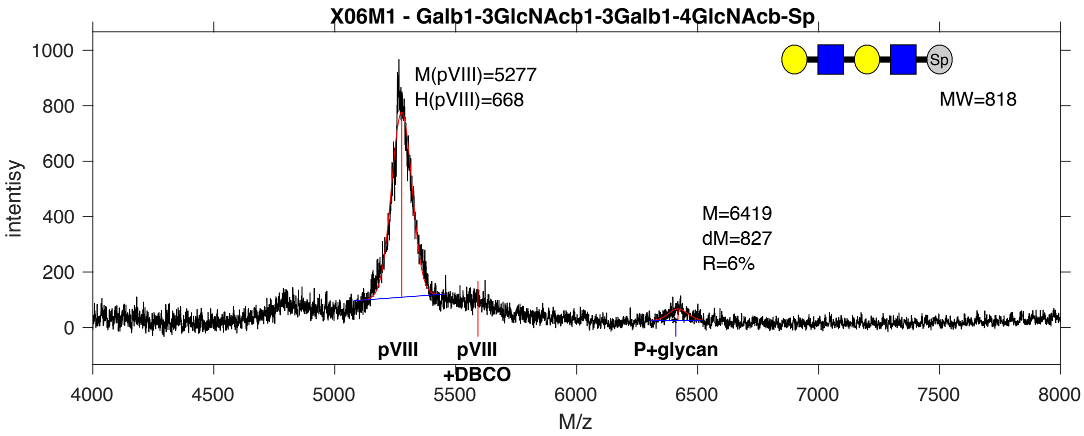


**SDB Number:** SDB68

**Sequencing File:** http://ligacloud.ca/searchLibInfo?f=0&b=0&d=20171128-67YYooOO1-1SS44

**Barcode:**
CTACTATTCGCGATCCCCCTCAGCGTGGAGAAGAACGACCAGAAGACGTATCACGCAGGGGGGGGG

**Axis Name:** LNT-Nac-[160]

**IUPAC:** Gal(b1-3)GlcNAc(b1-3)Gal(b1-4)GlcNAc(b1-Sp

**CFG Name:** Galb1-3GlcNAcb1-3Galb1-4GlcNAcb-Sp

**Common Name:** LNT-Nac

**Glytoucan ID:** G09681LD

**Compound Number:** X06M1

**Maldi File:** 106.txt

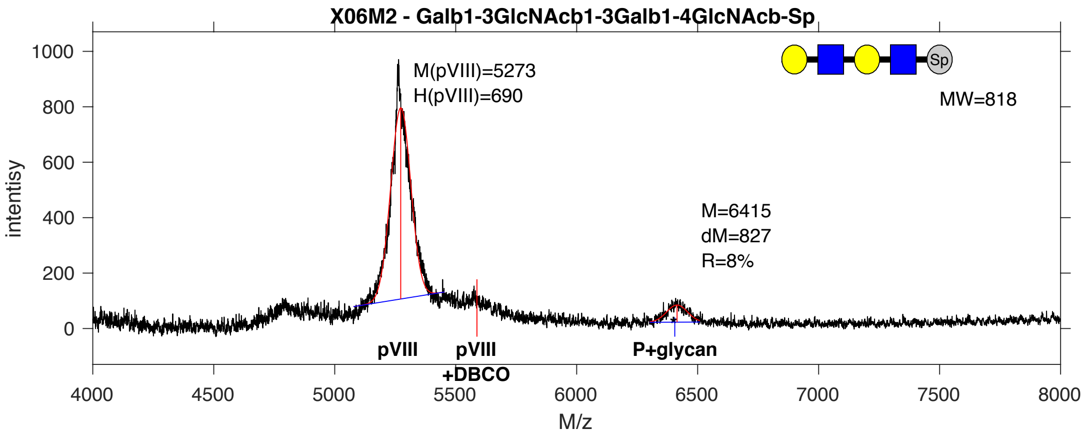


**SDB Number:** SDB69

**Sequencing File:** http://ligacloud.ca/searchLibInfo?f=0&b=0&d=20171128-87CLooOOKHLF-SS

**Barcode:**
CTTCTGTTTGCGATTCCACTGAGCGTGGAGAAGAACGACCAGAAAACATACCATGCTGGTGGTGGA

**Axis Name:** LNT-Nac-[220]

**IUPAC:** Gal(b1-3)GlcNAc(b1-3)Gal(b1-4)GlcNAc(b1-Sp

**CFG Name:** Galb1-3GlcNAcb1-3Galb1-4GlcNAcb-Sp

**Common Name:** LNT-Nac

**Glytoucan ID:** G09681LD

**Compound Number:** X06M2

**Maldi File:** 107.txt

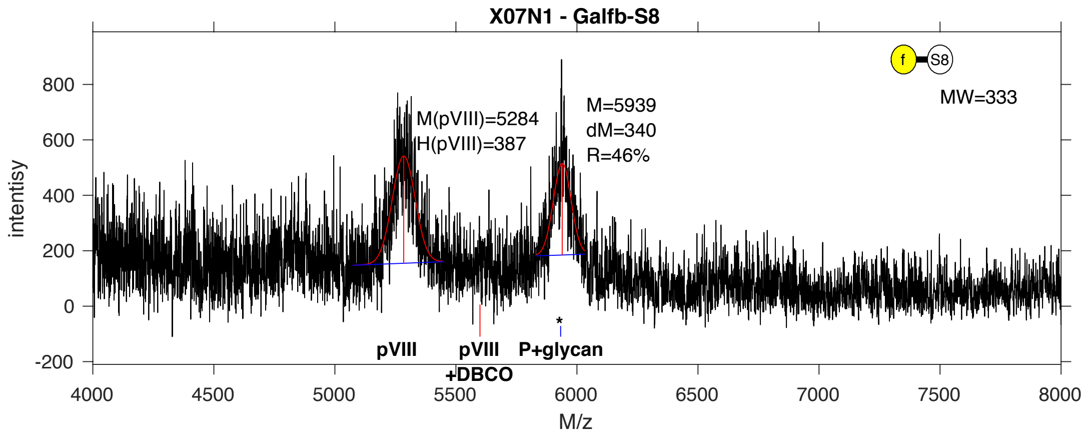


**SDB Number:** SDB80

**Sequencing File:** http://ligacloud.ca/searchLibInfo?f=0&b=0&d=20171128-87CLooOOINAX-SS

**Barcode:**
CTTCTGTTCGCGATCCCCCTAAGTGTAGAGAAGAATGATCAGAAAACGTATCATGCGGGAGGGGGT

**Axis Name:** Galf-[1300]

**IUPAC:** Galf(b1-S8

**CFG Name:** Galfb-S8

**Common Name:** Galf

**Glytoucan ID:** G38028NH

**Compound Number:** X07N1

**Maldi File:** 110.txt

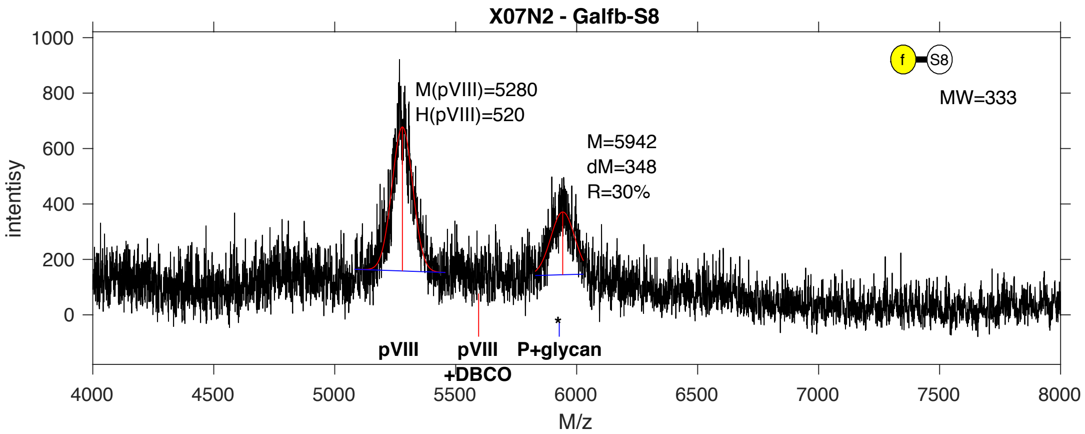


**SDB Number:** SDB81

**Sequencing File:** http://ligacloud.ca/searchLibInfo?f=0&b=0&d=20171128-87CLooOOIQKI-SS

**Barcode:**
CTACTCTTTGCTATTCCGCTTAGCGTGGAGAAGAATGATCAGAAAACGTACCACGCCGGTGGTGGC

**Axis Name:** Galf-[840]

**IUPAC:** Galf(b1-S8

**CFG Name:** Galfb-S8

**Common Name:** Galf

**Glytoucan ID:** G38028NH

**Compound Number:** X07N2

**Maldi File:** 111.txt

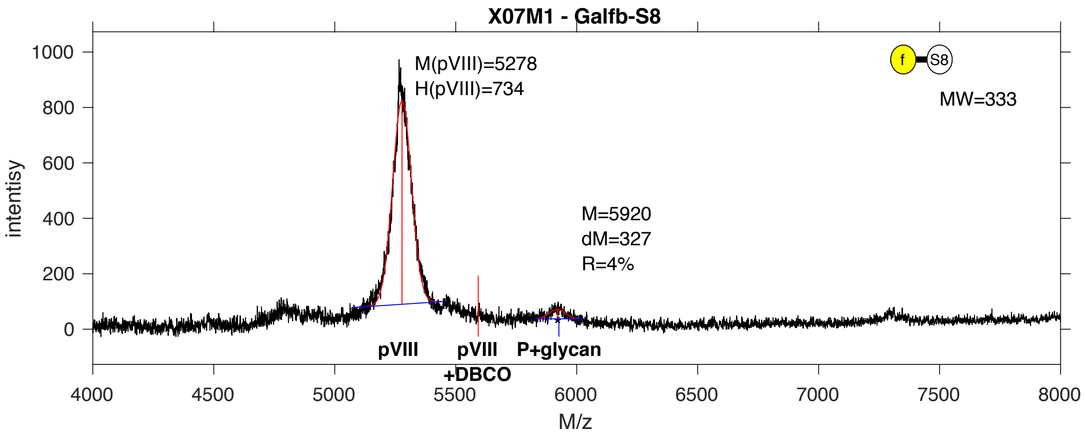


**SDB Number:** SDB82

**Sequencing File:** http://ligacloud.ca/searchLibInfo?f=0&b=0&d=20171128-87CLooOOEBYF-SS

**Barcode:**
TTATTATTCGCAATTCCTTTAAGCGTGGAGAAGAACGACCAAAAGACTTATCACGCGGCGGGGGG

**Axis Name:** Galf-[140]

**IUPAC:** Galf(b1-S8

**CFG Name:** Galfb-S8

**Common Name:** Galf

**Glytoucan ID:** G38028NH

**Compound Number:** X07M1

**Maldi File:** 112.txt

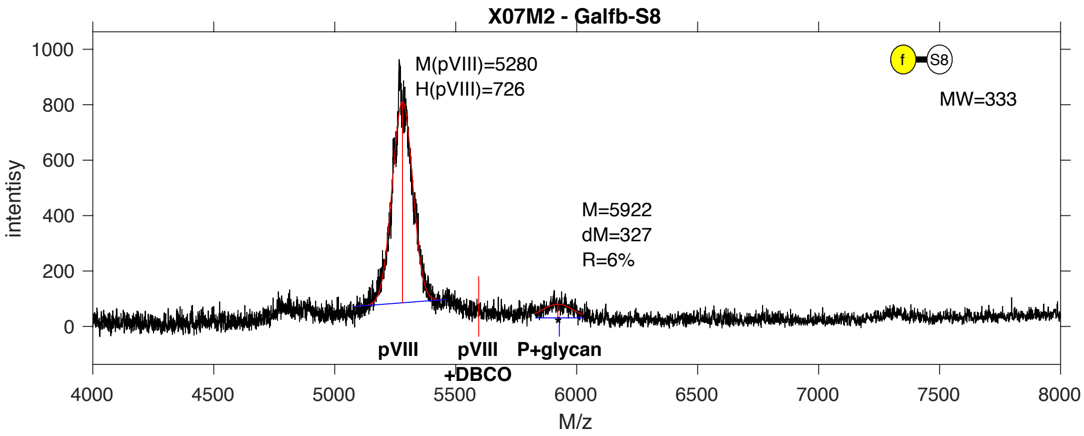


**SDB Number:** SDB83

**Sequencing File:** http://ligacloud.ca/searchLibInfo?f=0&b=0&d=20171128-87CLooOOUMWW-SS

**Barcode:**
CTGCTCTTCGCGATCCCTCTGAGTGTCGAAAAGAATGATCAAAAAACGTATCATGCGGGCGGTGGT

**Axis Name:** Galf-[160]

**IUPAC:** Galf(b1-S8

**CFG Name:** Galfb-S8

**Common Name:** Galf

**Glytoucan ID:** G38028NH

**Compound Number:** X07M2

**Maldi File:** 113.txt

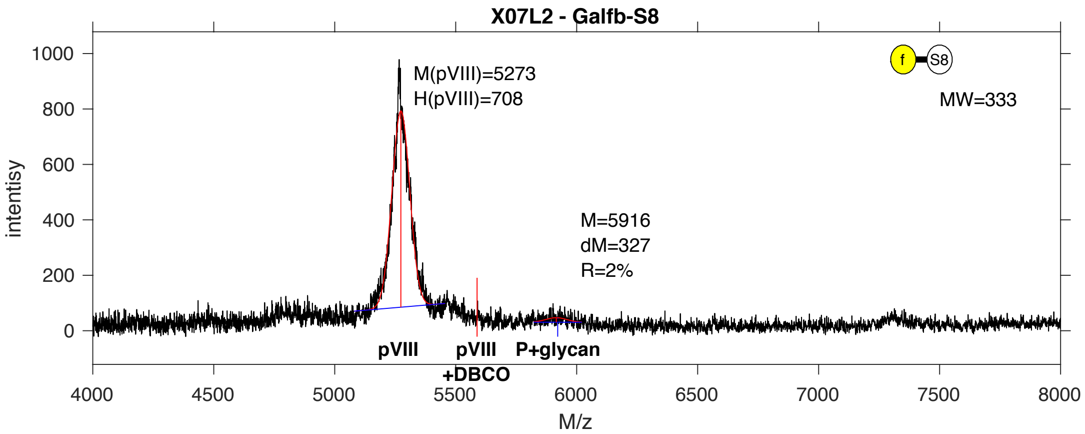


**SDB Number:** SDB86

**Sequencing File:** http://ligacloud.ca/searchLibInfo?f=0&b=0&d=20171128-87CLooOOKAWW-SS

**Barcode:**
CTTCTATTTGCTATTCCTCTAAGTGTGGAGAAGAATGATCAGAAGACATATCACGCTGGCGGGGGG

**Axis Name:** Galf-[110]

**IUPAC:** Galf(b1-S8

**CFG Name:** Galfb-S8

**Common Name:** Galf

**Glytoucan ID:** G38028NH

**Compound Number:** X07L2

**Maldi File:** 115.txt

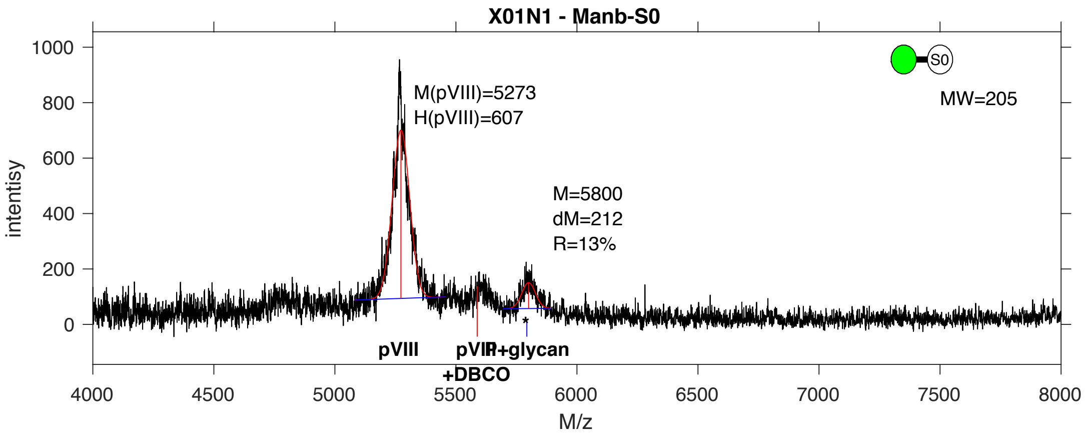


**SDB Number:** SDB87

**Sequencing File:** http://ligacloud.ca/searchLibInfo?f=0&b=0&d=20171128-87CLooOOPDDQ-SS

**Barcode:**
CTACTCTTCGCTATACCCCTCAGCGTAGAAAAAAATGACCAAAAGACCTACCATGCGGGAGGTGGG

**Axis Name:** bMan-[350]

**IUPAC:** Man(b1-S0

**CFG Name:** Manb-S0

**Common Name:** bMan

**Glytoucan ID:** G58490PL

**Compound Number:** X01N1

**Maldi File:** 116.txt

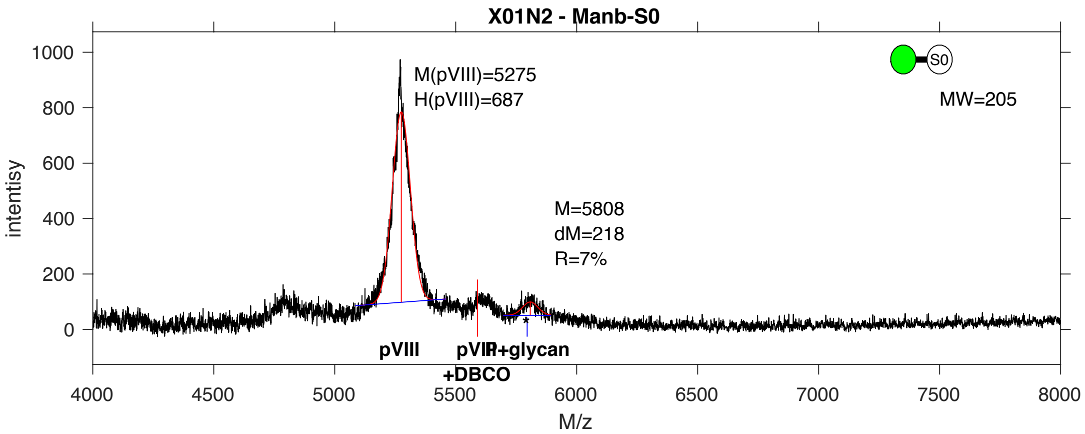


**SDB Number:** SDB88

**Sequencing File:** http://ligacloud.ca/searchLibInfo?f=0&b=0&d=20171128-87CLooOORWFR-SS

**Barcode:**
CTGCTGTTCGCGATTCCTCTAAGTGTAGAGAAGAATGACCAGAAGACCTACCACGCGGGCGGGGGG

**Axis Name:** bMan-[190]

**IUPAC:** Man(b1-S0

**CFG Name:** Manb-S0

**Common Name:** bMan

**Glytoucan ID:** G58490PL

**Compound Number:** X01N2

**Maldi File:** 117.txt

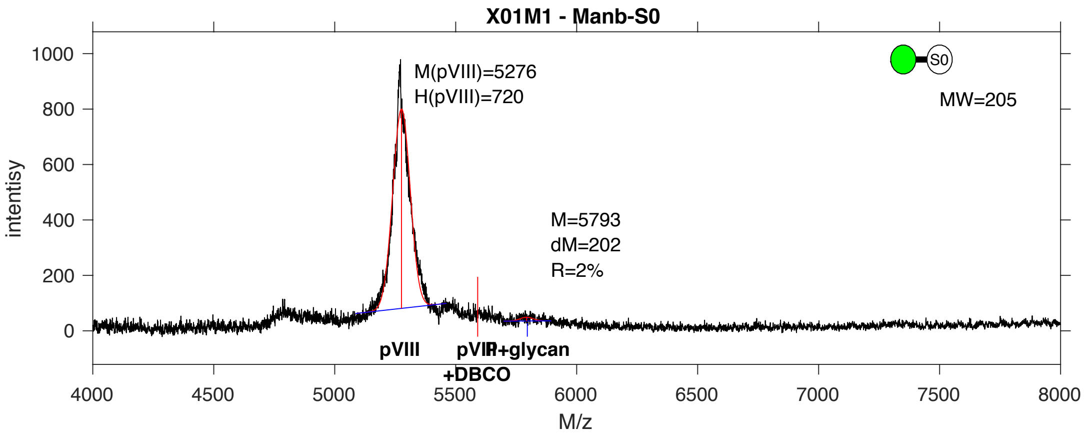


**SDB Number:** SDB89

**Sequencing File:** http://ligacloud.ca/searchLibInfo?f=0&b=0&d=20171128-87CLooOONRBH-SS

**Barcode:**
CTACTGTTCGCAATACCTCTCAGCGTGGAGAAAAATGATCAAAAAACATATCATGCAGGGGGTGGC

**Axis Name:** bMan-[80]

**IUPAC:** Man(b1-S0

**CFG Name:** Manb-S0

**Common Name:** bMan

**Glytoucan ID:** G58490PL

**Compound Number:** X01M1

**Maldi File:** 118.txt

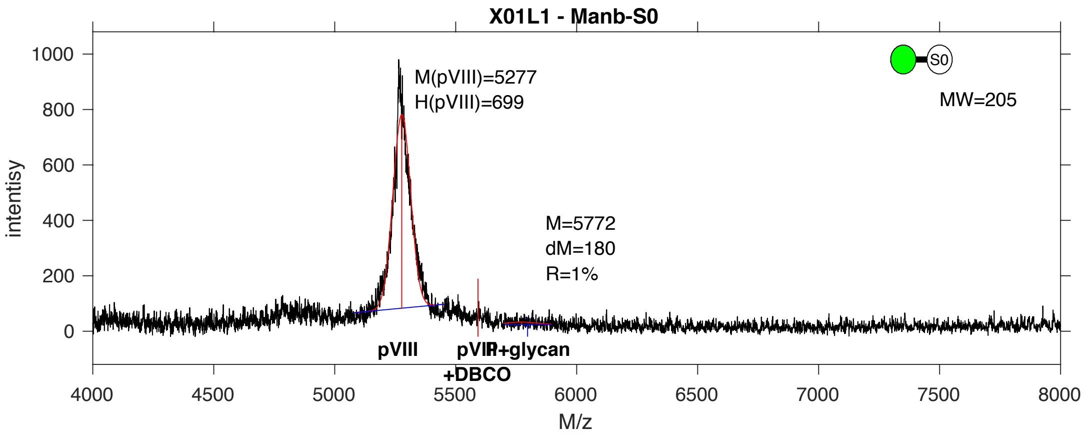


**SDB Number:** SDB99

**Sequencing File:** http://ligacloud.ca/searchLibInfo?f=0&b=0&d=20171128-87CLooOOZLUQ-SS

**Barcode:**
CTTCTCTTCGCGATACCGCTAAGTGTTGAGAAAAATGATCAAAAAACGTATCATGCGGGTGGTGGG

**Axis Name:** bMan-[<8]

**IUPAC:** Man(b1-S0

**CFG Name:** Manb-S0

**Common Name:** bMan

**Glytoucan ID:** G58490PL

**Compound Number:** X01L1

**Maldi File:** 119.txt

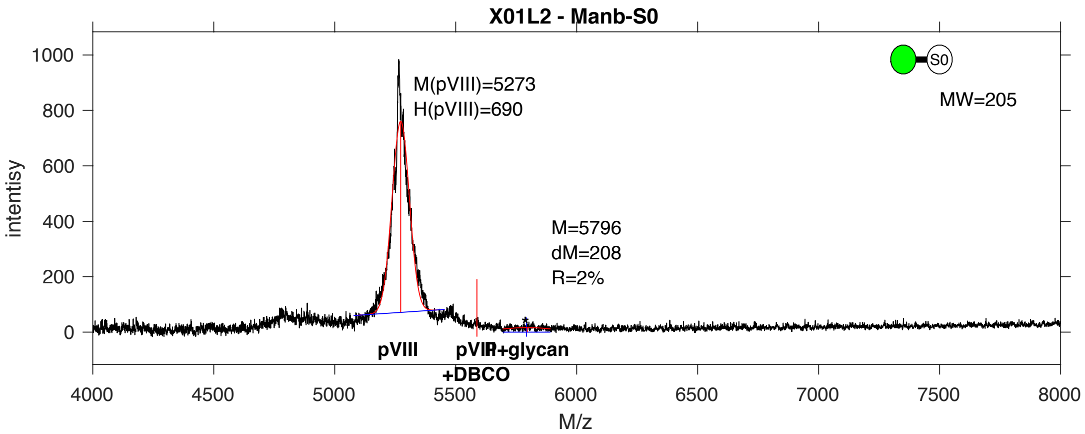


**SDB Number:** SDB126

**Sequencing File:** http://ligacloud.ca/searchLibInfo?f=0&b=0&d=20171128-87CLooOOXEGX-SS

**Barcode:**
CTGCTCTTTGCAATTCCGCTGAGTGTAGAAAAAAATGATCAGAAAACATACCATGCTGGTGGCGGA

**Axis Name:** bMan-[<8]

**IUPAC:** Man(b1-S0

**CFG Name:** Manb-S0

**Common Name:** bMan

**Glytoucan ID:** G58490PL

**Compound Number:** X01L2

**Maldi File:** 120.txt

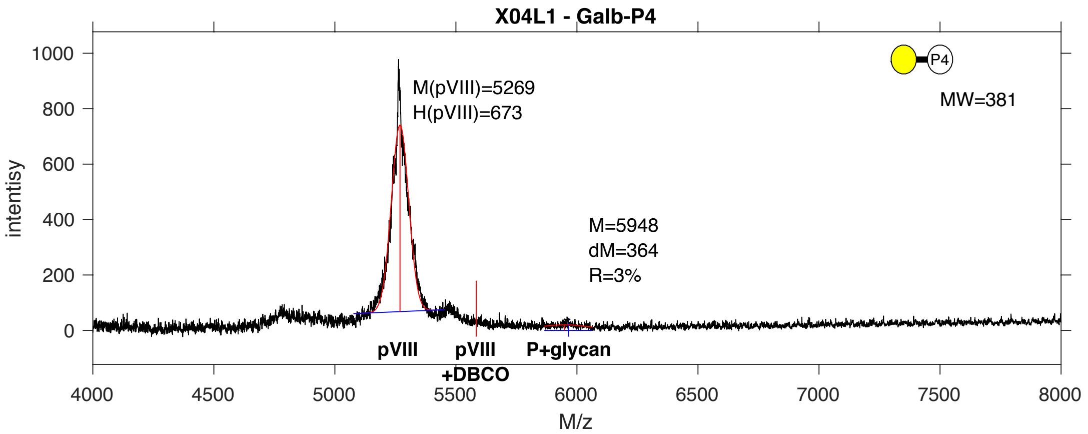


**SDB Number:** SDB132

**Sequencing File:** http://ligacloud.ca/searchLibInfo?f=0&b=0&d=20171128-87CLooOOTUHU-SS

**Barcode:**
CTACTGTTCGCAATACCTCTCAGCGTAGAAAAAAATGATCAAAAAACATATCACGCAGGTGGCGGA

**Axis Name:** Gal-[<8]

**IUPAC:** Gal(b1-P4

**CFG Name:** Galb-P4

**Common Name:** Gal

**Glytoucan ID:** G65889KE

**Compound Number:** X04L1

**Maldi File:** 122.txt

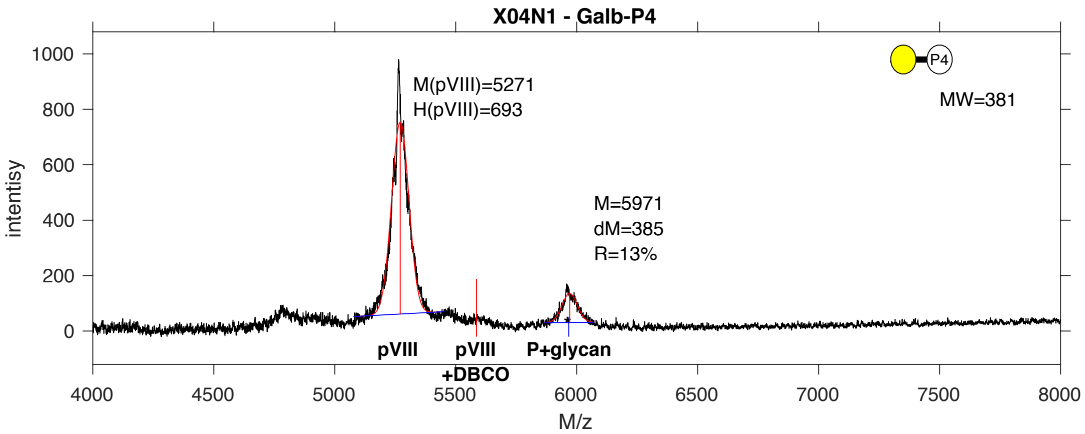


**SDB Number:** SDB127

**Sequencing File:** http://ligacloud.ca/searchLibInfo?f=0&b=0&d=20171128-87CLooOOZZNY-SS

**Barcode:**
CTGCTATTTGCCATCCCACTAAGTGTTGAGAAGAACGATCAGAAAACGTATCATGCAGGTGGTGGA

**Axis Name:** Gal-[380]

**IUPAC:** Gal(b1-P4

**CFG Name:** Galb-P4

**Common Name:** Gal

**Glytoucan ID:** G65889KE

**Compound Number:** X04N1

**Maldi File:** 123.txt

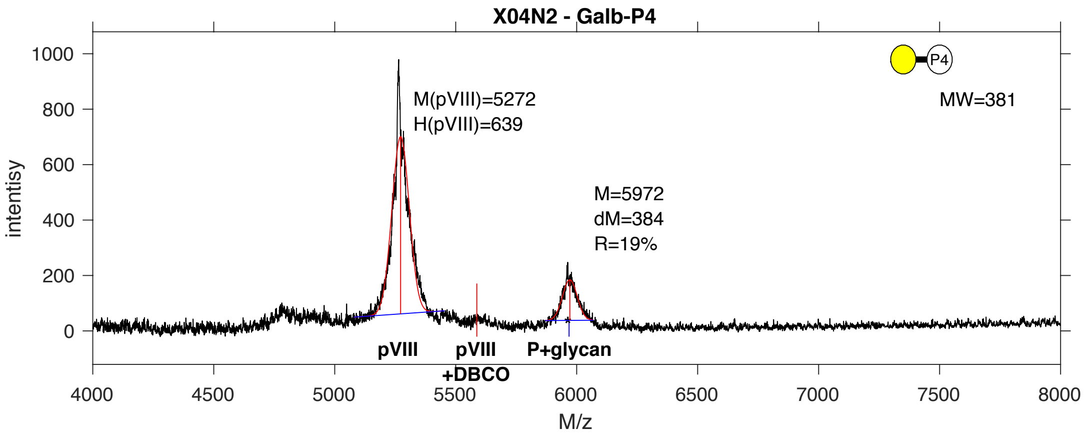


**SDB Number:** SDB128

**Sequencing File:** http://ligacloud.ca/searchLibInfo?f=0&b=0&d=20171128-87CLooOONWSB-SS

**Barcode:**
TTATTATTCGCAATTCCTTTAAGCGTTGAAAAGAATGACCAGAAAACATATCACGCTGGGGGAGGG

**Axis Name:** Gal-[510]

**IUPAC:** Gal(b1-P4

**CFG Name:** Galb-P4

**Common Name:** Gal

**Glytoucan ID:** G65889KE

**Compound Number:** X04N2

**Maldi File:** 124.txt

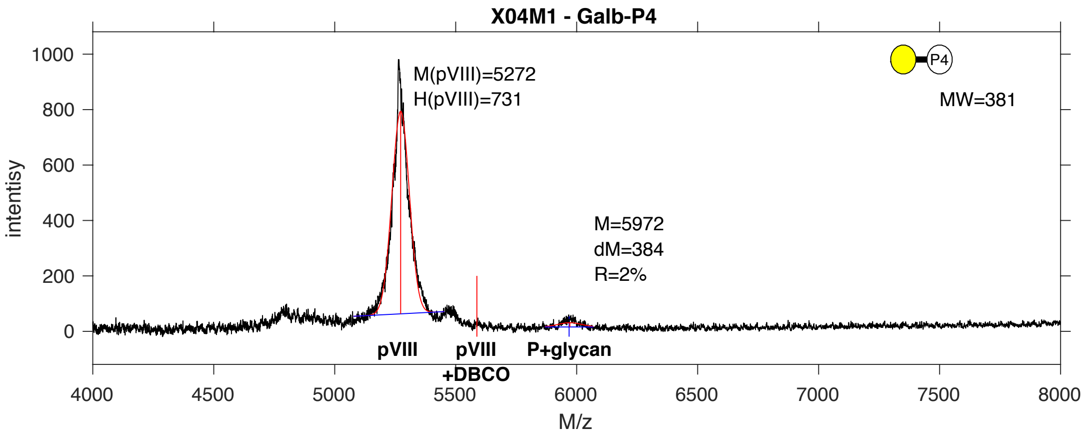


**SDB Number:** SDB129

**Sequencing File:** http://ligacloud.ca/searchLibInfo?f=0&b=0&d=20171128-87CLooOOOUNV-SS

**Barcode:**
CTTCTGTTCGCGATACCGCTGAGCGTCGAAAAGAATGATCAGAAAACTTATCATGCGGGGGGCGGG

**Axis Name:** Gal-[110]

**IUPAC:** Gal(b1-P4

**CFG Name:** Galb-P4

**Common Name:** Gal

**Glytoucan ID:** G65889KE

**Compound Number:** X04M1

**Maldi File:** 125.txt

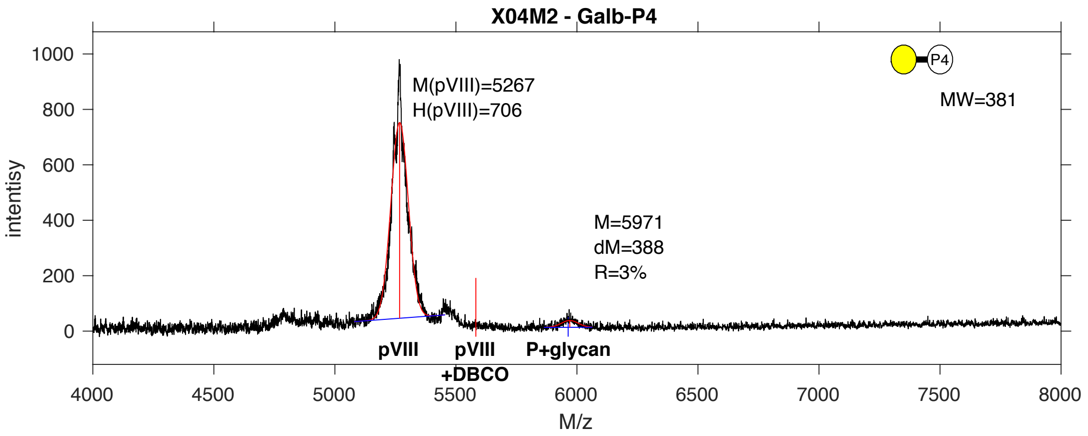


**SDB Number:** SDB130

**Sequencing File:** http://ligacloud.ca/searchLibInfo?f=0&b=0&d=20171128-87CLooOOTWPF-SS

**Barcode:**
CTTCTCTTTGCGATACCGCTAAGTGTGGAAAAGAATGATCAAAAAACGTACCATGCCGGTGGGGGT

**Axis Name:** Gal-[110]

**IUPAC:** Gal(b1-P4

**CFG Name:** Galb-P4

**Common Name:** Gal

**Glytoucan ID:** G65889KE

**Compound Number:** X04M2

**Maldi File:** 126.txt

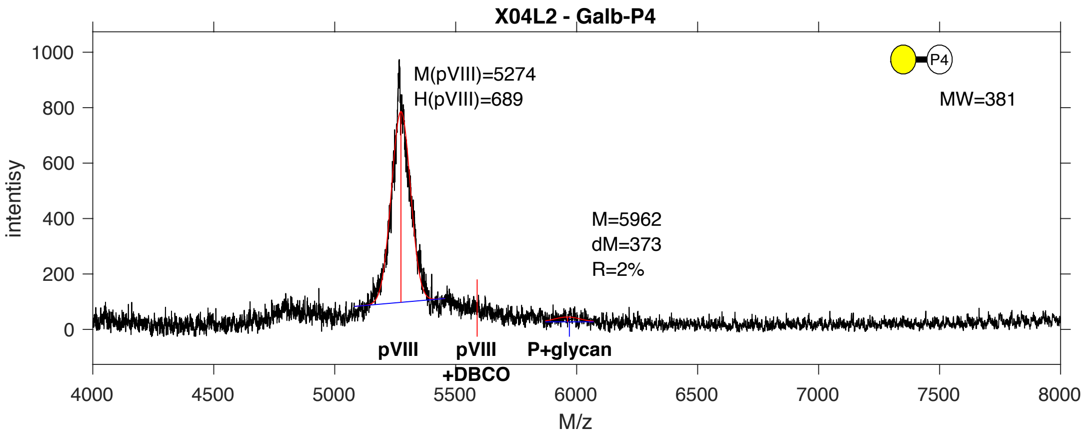


**SDB Number:** SDB107

**Sequencing File:** http://ligacloud.ca/searchLibInfo?f=0&b=0&d=20171128-87CLooOODAYD-SS

**Barcode:**
TTATTATTCGCAATTCCTTTAAGTGTGGAGAAGAACGATCAGAAGACCTATCACGCAGGAGGCGGA

**Axis Name:** Gal-[<8]

**IUPAC:** Gal(b1-P4

**CFG Name:** Galb-P4

**Common Name:** Gal

**Glytoucan ID:** G65889KE

**Compound Number:** X04L2

**Maldi File:** 127.txt

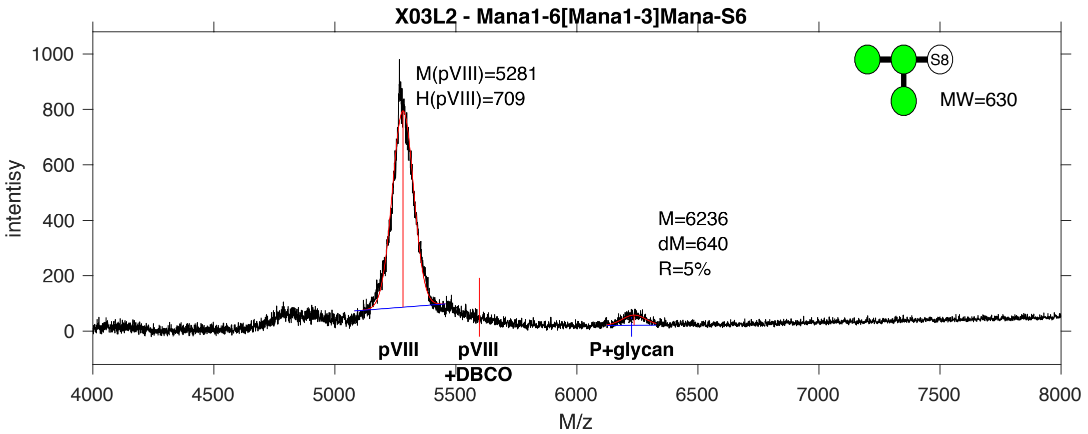


**SDB Number:** SDB18

**Sequencing File:** http://ligacloud.ca/searchLibInfo?f=0&b=0&d=20171128-87CLooOOAQTT-SS

**Barcode:**
CTGCTGTTTGCTATCCCTCTGAGTGTGGAGAAGAATGATCAGAAGACTTATCATGCGGGTGGAGGT

**Axis Name:** Tri-Man-[140]

**IUPAC:** Man(a1-6)[Man(a1-3)]Man(a1-S6

**CFG Name:** Mana1-6[Mana1-3]Mana-S6

**Common Name:** Tri-Man

**Glytoucan ID:** G80926OA

**Compound Number:** X03L2

**Maldi File:** 128.txt

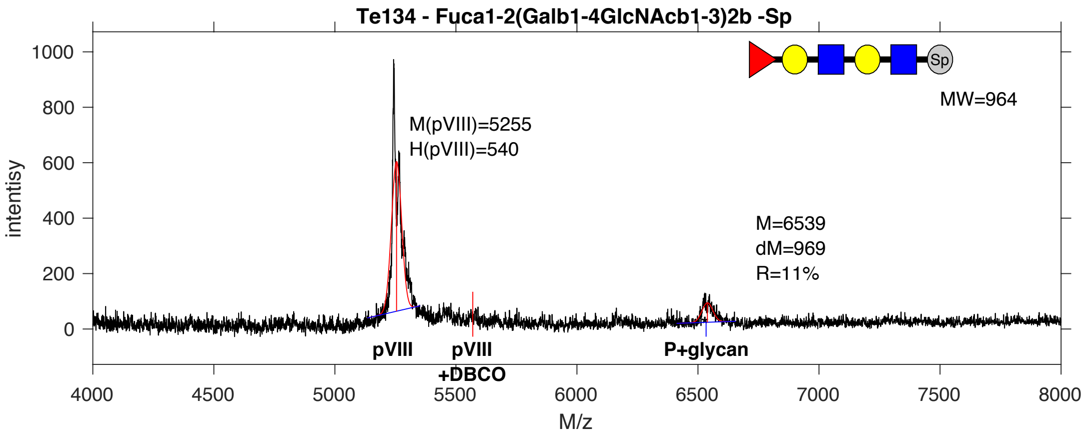


**SDB Number:** SDB76

**Sequencing File:** http://ligacloud.ca/searchLibInfo?f=0&b=0&d=20180522-87YJooOO1-1JM2

**Barcode:**
CTGCTTTTTGCCATACCGCTAAGTGTTGAGAAGAATGATCAGAAGACCTACCACGCAGGAGGTGGC

**Axis Name:** H2-[300]

**IUPAC:** Fuc(a1-2)Gal(b1-4)GlcNAc(b1-3)Gal(b1-4)GlcNAc(b1-Sp

**CFG Name:** Fuca1-2(Galb1-4GlcNAcb1-3)2b-Sp

**Common Name:** H2

**Glytoucan ID:** G99268MP

**Compound Number:** Te134

**Maldi File:** 134.txt

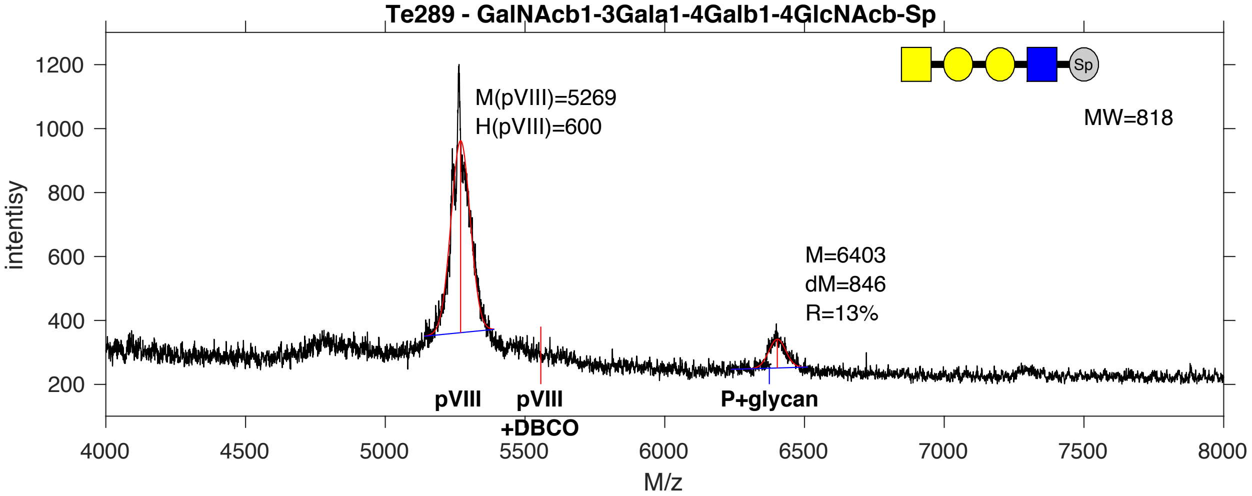


**SDB Number:** SDB5

**Sequencing File:** http://ligacloud.ca/searchLibInfo?f=0&b=0&d=NA

**Barcode:**
CTTCTGTTCGCCATTCCGCTGAGTGTGGAGAAGAATGATCAGAAGACTTATCATGCGGGTGGAGGT

**Axis Name:** P1 tetra-[350]

**IUPAC:** GalNAc(b1-3)Gal(a1-4)Gal(b1-4)GlcNAc(b1-Sp

**CFG Name:** GalNAcb1-3Gala1-4Galb1-4GlcNAcb-Sp

**Common Name:** P1 tetra

**Glytoucan ID:** G29548YX

**Compound Number:** Te289

**Maldi File:** 137.txt

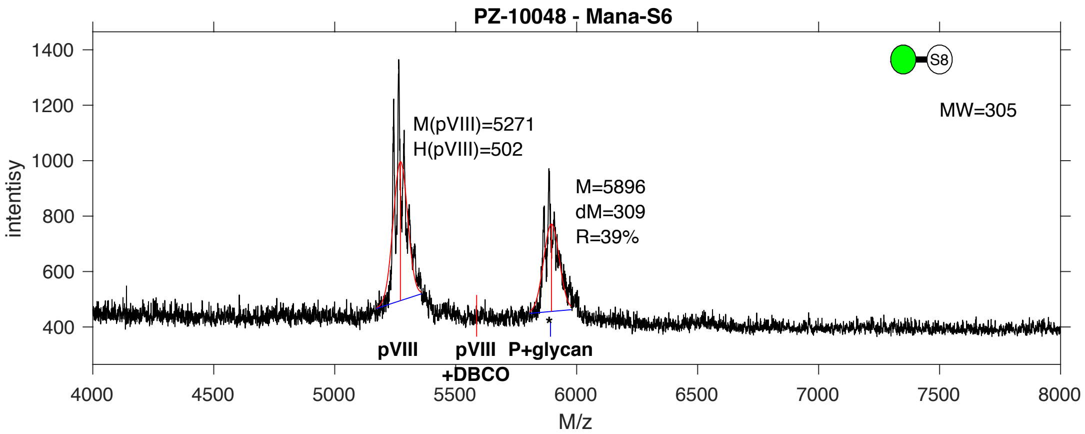


**SDB Number:** SDB28

**Sequencing File:** http://ligacloud.ca/searchLibInfo?f=0&b=0&d=20171128-87CLooOOIJYY-SS

**Barcode:**
CTACTCTTCGCCATTCCACTGAGTGTGGAGAAGAATGATCAGAAGACTTATCATGCGGGTGGAGGT

**Axis Name:** aMan-[1030]

**IUPAC:** Man(a1-S6

**CFG Name:** Mana-S6

**Common Name:** aMan

**Glytoucan ID:** G19165TD

**Compound Number:** PZ-10048

**Maldi File:** 142.txt

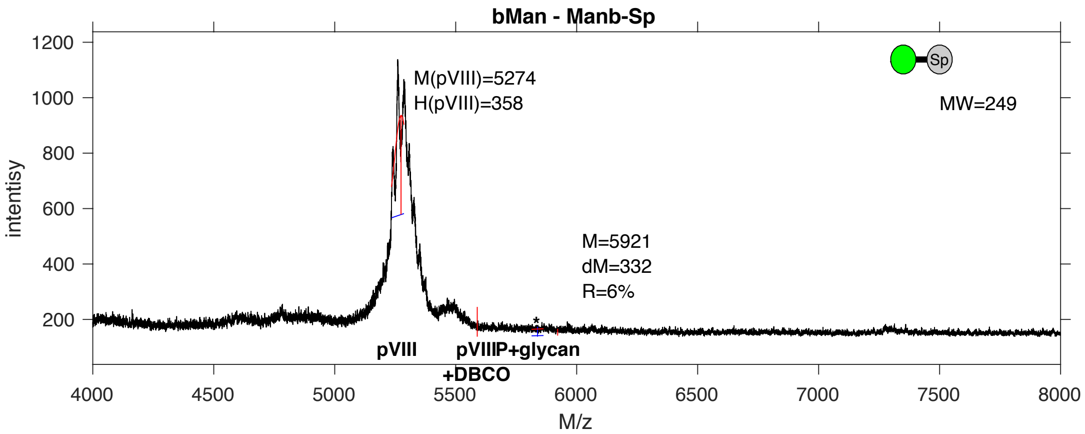


**SDB Number:** SDB3

**Sequencing File:** http://ligacloud.ca/searchLibInfo?f=0&b=0&d=NA

**Barcode:**
CTGCTTTTCGCAATTCCGCTTAGTGTGGAGAAGAATGATCAGAAGACTTATCATGCGGGTGGAGGT

**Axis Name:** bMan-[<8]

**IUPAC:** Man(b1-S0

**CFG Name:** Manb-S0

**Common Name:** bMan

**Glytoucan ID:** G58490PL

**Compound Number:** bMan-2

**Maldi File:** 150.txt

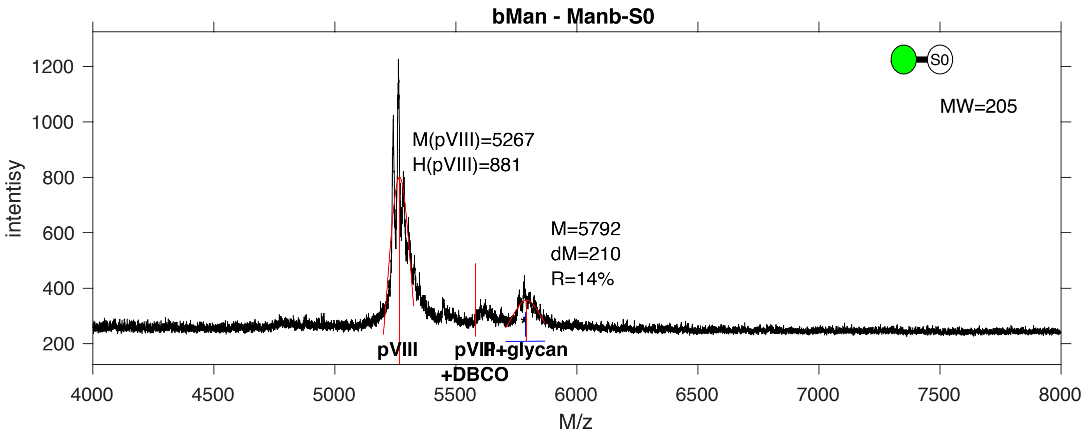


**SDB Number:** SDB8

**Sequencing File:** http://ligacloud.ca/searchLibInfo?f=0&b=0&d=20171128-87CLooOOOWCC-SS

**Barcode:**
CTGCTTTTTGCAATACCCCTCAGTGTGGAGAAGAATGATCAGAAGACTTATCATGCGGGTGGAGGT

**Axis Name:** bMan-[320]

**IUPAC:** Man(b1-S0

**CFG Name:** Manb-S0

**Common Name:** bMan

**Glytoucan ID:** G58490PL

**Compound Number:** bMan

**Maldi File:** 153.txt

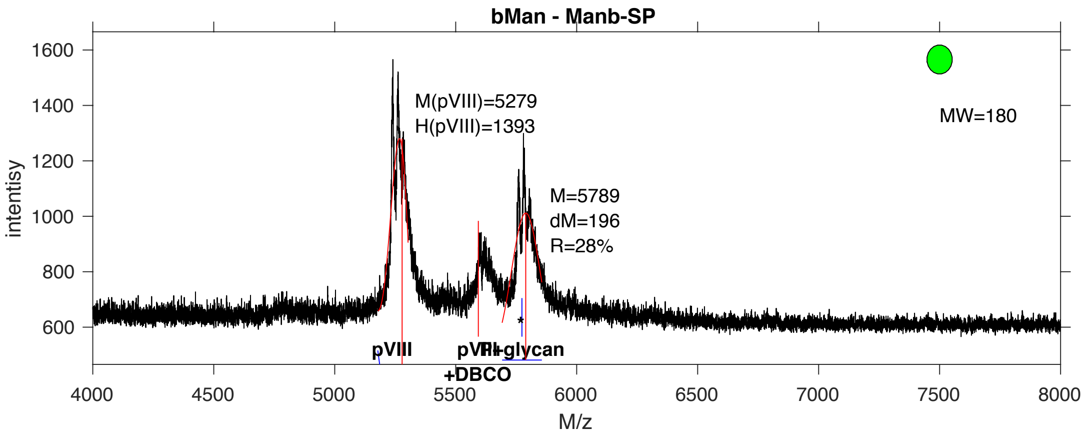


**SDB Number:** SDB11

**Sequencing File:** http://ligacloud.ca/searchLibInfo?f=0&b=0&d=20171128-87CLooOOZCHR-SS

**Barcode:**
CTACTATTTGCGATTCCCCTGAGTGTGGAGAAGAATGATCAGAAGACTTATCATGCGGGTGGAGGT

**Axis Name:** bMan-[1160]

**IUPAC:** Man(b1-S0

**CFG Name:** Manb-S0

**Common Name:** bMan

**Glytoucan ID:** G58490PL

**Compound Number:** bMan

**Maldi File:** 156.txt

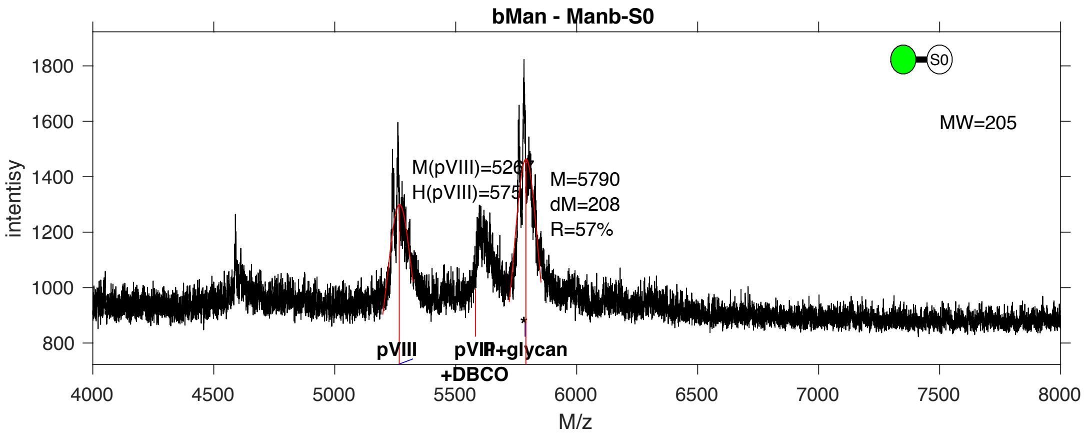


**SDB Number:** SDB12

**Sequencing File:** http://ligacloud.ca/searchLibInfo?f=0&b=0&d=20171128-87CLooOOHMWZ-SS

**Barcode:**
CTTCTGTTCGCGATACCTCTAAGTGTGGAGAAGAATGATCAGAAGACTTATCATGCGGGTGGAGGT

**Axis Name:** bMan-[1430]

**IUPAC:** Man(b1-S0

**CFG Name:** Manb-S0

**Common Name:** bMan

**Glytoucan ID:** G58490PL

**Compound Number:** bMan

**Maldi File:** 157.txt

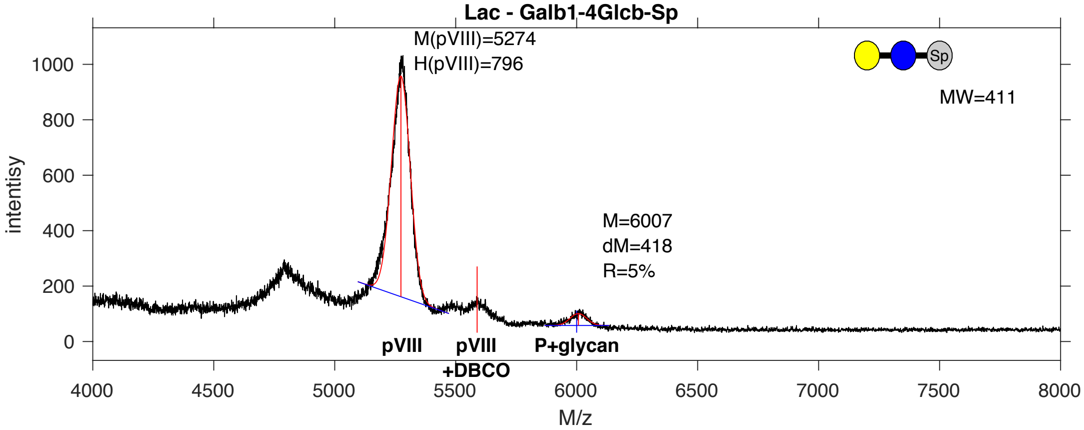


**SDB Number:** SDB25

**Sequencing File:** http://ligacloud.ca/searchLibInfo?f=0&b=0&d=20171128-87CLooOOQQEH-SS

**Barcode:**
CTGCTTTTCGCAATACCTCTAAGTGTGGAGAAGAATGATCAGAAGACTTATCATGCGGGTGGAGGT

**Axis Name:** Lac-[140]

**IUPAC:** Gal(b1-4)Glc(b1-Sp

**CFG Name:** Galb1-4Glcb-Sp

**Common Name:** Lac

**Glytoucan ID:** G94144EF

**Compound Number:** Lac

**Maldi File:** 158.txt

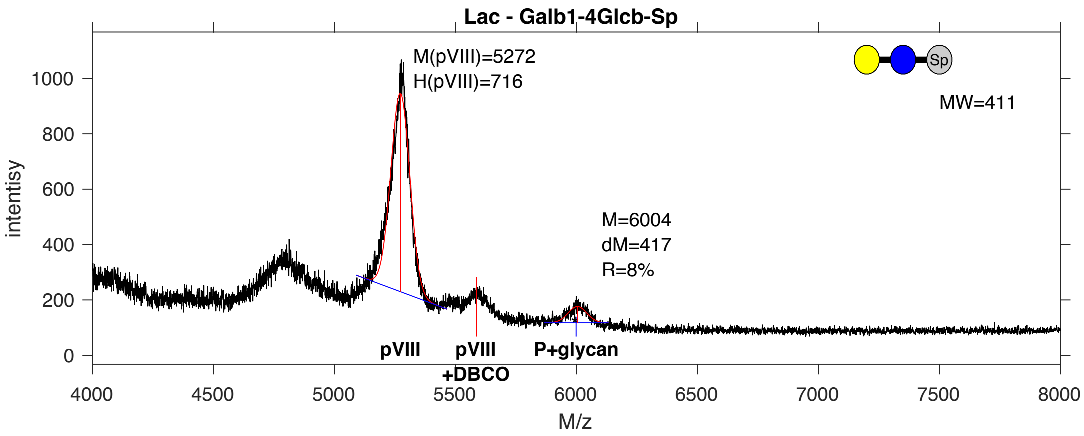


**SDB Number:** SDB27

**Sequencing File:** http://ligacloud.ca/searchLibInfo?f=0&b=0&d=NA

**Barcode:**
CTGCTTTTCGCTATTCCTCTCAGTGTGGAGAAGAATGATCAGAAGACTTATCATGCGGGTGGAGGT

**Axis Name:** Lac-[220]

**IUPAC:** Gal(b1-4)Glc(b1-Sp

**CFG Name:** Galb1-4Glcb-Sp

**Common Name:** Lac

**Glytoucan ID:** G94144EF

**Compound Number:** Lac

**Maldi File:** 159.txt

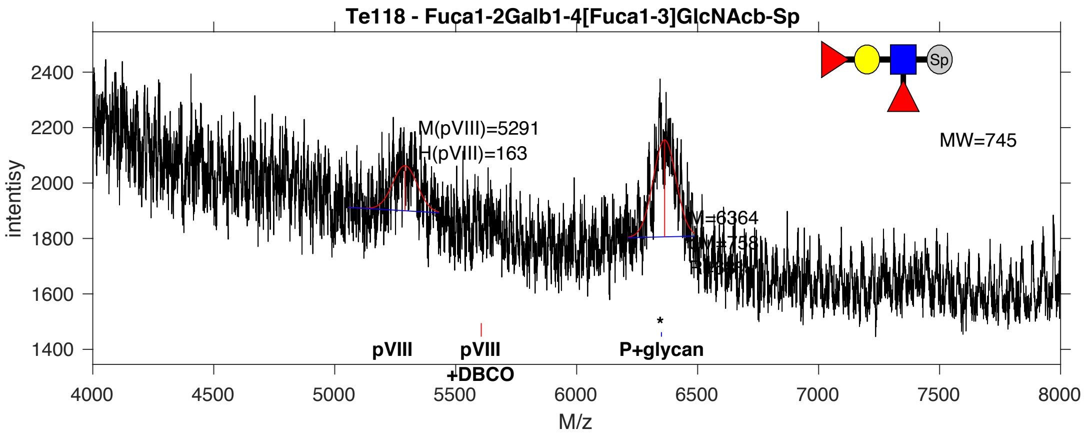


**SDB Number:** SDB50

**Sequencing File:** http://ligacloud.ca/searchLibInfo?f=0&b=0&d=20180809-87CLooPA1-1JM17

**Barcode:**
CTGCTCTTTGCGATTCCGCTGAGTGTCGAGAAGAACGACCAGAAAACATATCACGCGGGGGGGGGA

**Axis Name:** Ley-[1780]

**IUPAC:** Fuc(a1-2)Gal(b1-4)[Fuc(a1-3)]GlcNAc(b1-Sp

**CFG Name:** Fuca1-2Galb1-4[Fuca1-3]GlcNAcb-Sp

**Common Name:** Ley

**Glytoucan ID:** G35534DU

**Compound Number:** Te118

**Maldi File:** 170.txt

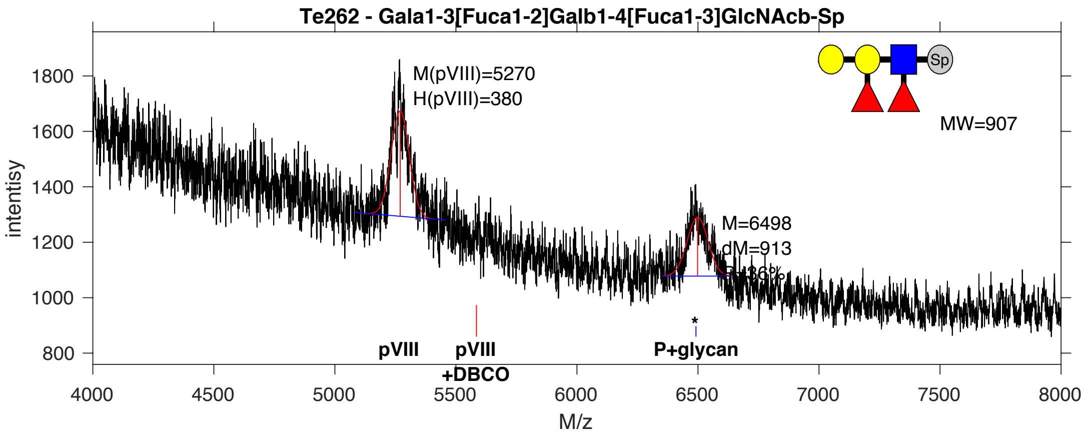


**SDB Number:** SDB39

**Sequencing File:** http://ligacloud.ca/searchLibInfo?f=0&b=0&d=NA

**Barcode:**
CTGCTGTTTGCGATCCCTCTAAGTGTCGAAAAGAACGACCAAAAAACTTACCTTGCAGGGGGGGGT

**Axis Name:** 2’F-B type 2-[970]

**IUPAC:** Gal(a1-3)[Fuc(a1-2)]Gal(b1-4)[Fuc(a1-3)]GlcNAc(b1-Sp

**CFG Name:** Gala1-3[Fuca1-2]Galb1-4[Fuca1-3]GlcNAcb-Sp

**Common Name:** 2’F-B type 2

**Glytoucan ID:** G04624DG

**Compound Number:** Te262

**Maldi File:** 171.txt

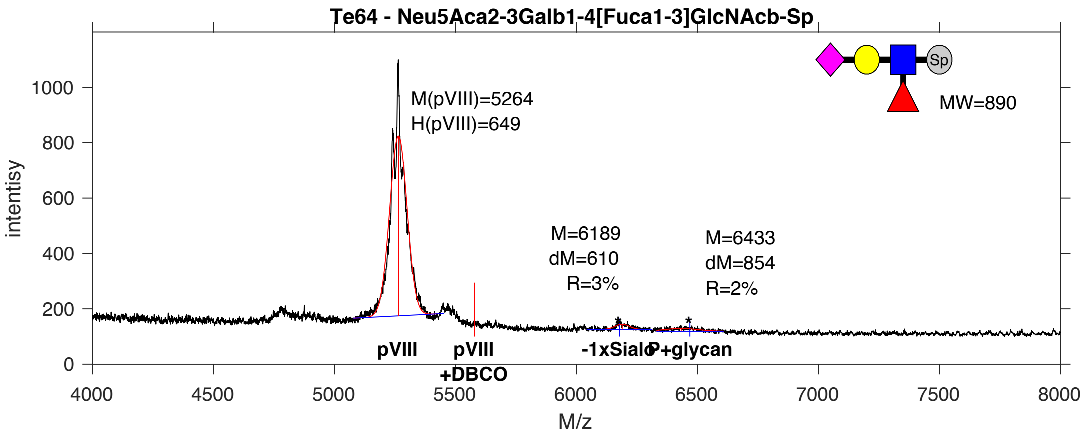


**SDB Number:** SDB65

**Sequencing File:** http://ligacloud.ca/searchLibInfo?f=0&b=0&d=NA

**Barcode:**
CTGCTTTTTGCCATCCCTCTTAGTGTTGAAAAGAACGATCAAAAAACTTATCACGCAGGGGGGGGG

**Axis Name:** 3’SLex-[140]

**IUPAC:** Neu5Ac(a2-3)Gal(b1-4)[Fuc(a1-3)]GlcNAc(b1-Sp

**CFG Name:** Neu5Aca2-3Galb1-4[Fuca1-3]GlcNAcb-Sp

**Common Name:** 3’SLex

**Glytoucan ID:** G55692SC

**Compound Number:** Te64

**Maldi File:** 173.txt

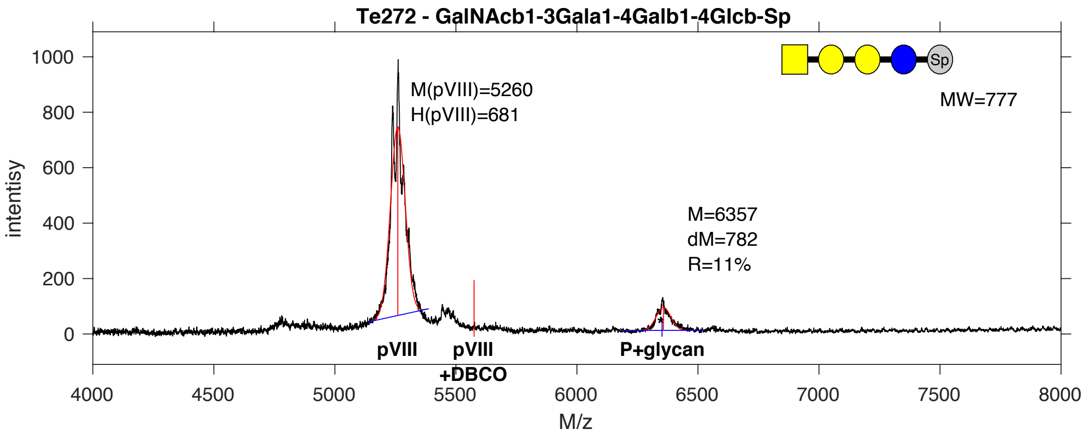


**SDB Number:** SDB44

**Sequencing File:** http://ligacloud.ca/searchLibInfo?f=0&b=0&d=20180809-87CLooPA1-1JM10

**Barcode:**
CTGCTATTTGCGATTCCCCTAAGTGTAGAGAAAAACGATCAAAAGACGTATCATGCCGGGGGCGGT

**Axis Name:** Globoside-P-[300]

**IUPAC:** GalNAc(b1-3)Gal(a1-4)Gal(b1-4)Glc(b1-Sp

**CFG Name:** GalNAcb1-3Gala1-4Galb1-4Glcb-Sp

**Common Name:** Globoside-P

**Glytoucan ID:** G24499XE

**Compound Number:** Te272

**Maldi File:** 176.txt

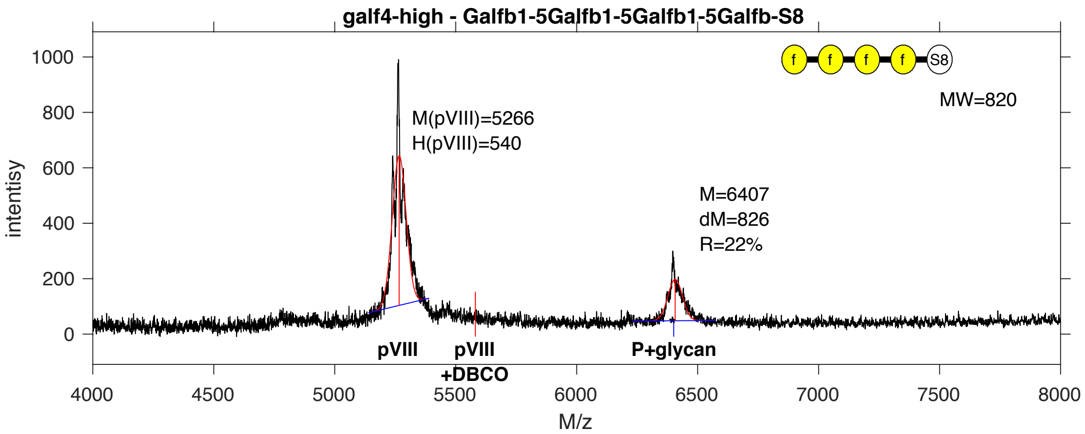


**SDB Number:** SDB20

**Sequencing File:** http://ligacloud.ca/searchLibInfo?f=0&b=0&d=20171128-87CLooOOXIFJ-SS

**Barcode:**
CTGCTCTTTGCCATCCCGCTTAGTGTGGAGAAGAATGATCAGAAGACTTATCATGCGGGTGGAGGT

**Axis Name:** (Galf)4-[590]

**IUPAC:** Galf(b1-5)Galf(b1-5)Galf(b1-5)Galf(b1-S8

**CFG Name:** Galfb1-5Galfb1-5Galfb1-5Galfb-S8

**Common Name:** (Galf)4

**Glytoucan ID:** G92890KE

**Compound Number:** galf4-high

**Maldi File:** 178.txt

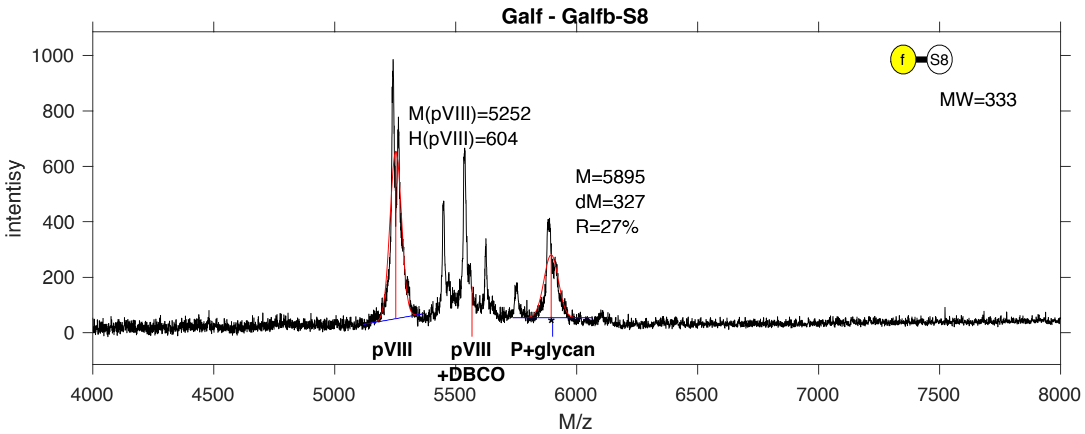


**SDB Number:** SDB130

**Sequencing File:** http://ligacloud.ca/searchLibInfo?f=0&b=0&d=20171128-87CLooOOTWPF-SS

**Barcode:**
CTTCTCTTTGCGATACCGCTAAGTGTGGAAAAGAATGATCAAAAAACGTACCATGCCGGTGGGGGT

**Axis Name:** Galf-[730]

**IUPAC:** Galf(b1-S8

**CFG Name:** Galfb-S8

**Common Name:** Galf

**Glytoucan ID:** G38028NH

**Compound Number:** Galf

**Maldi File:** 179.txt

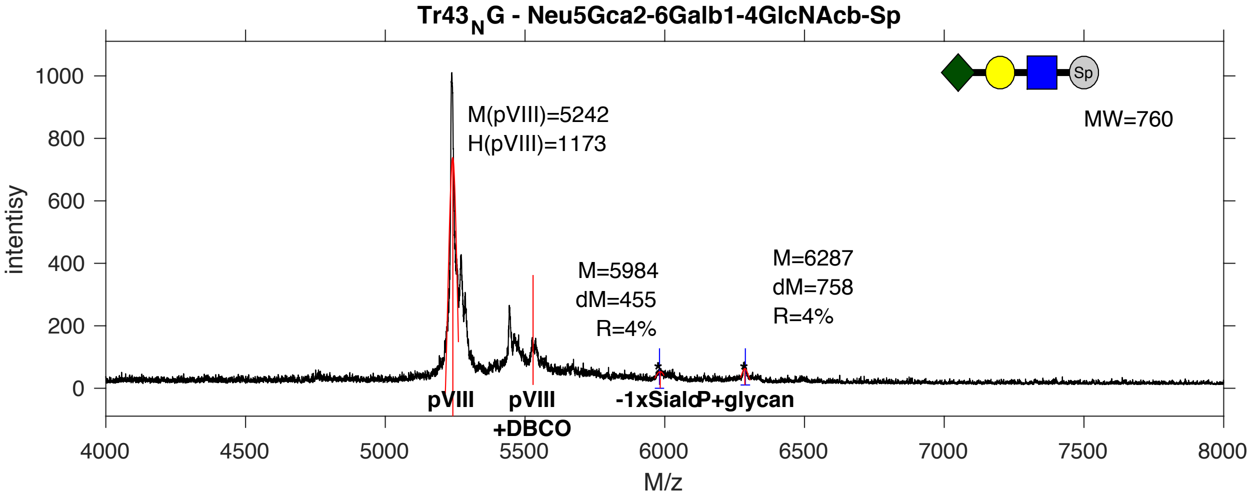


**SDB Number:** SDB179

**Sequencing File:** http://ligacloud.ca/searchLibInfo?f=0&b=0&d=NA

**Barcode:**
CTACTGTTCGCAATCCCGCTAAGTGTCGAAAAAAATGATCAAAAGACTTACCACGCTGGAGGGGGT

**Axis Name:** 6'SLN (Gc)-[220]

**IUPAC:** Neu5Gc(a2-6)Gal(b1-4)GlcNAc(b1-Sp

**CFG Name:** Neu5Gca2-6Galb1-4GlcNAcb-Sp

**Common Name:** 6'SLN (Gc)

**Glytoucan ID:** G24684AO

**Compound Number:** Tr43_NG

**Maldi File:** 380.txt

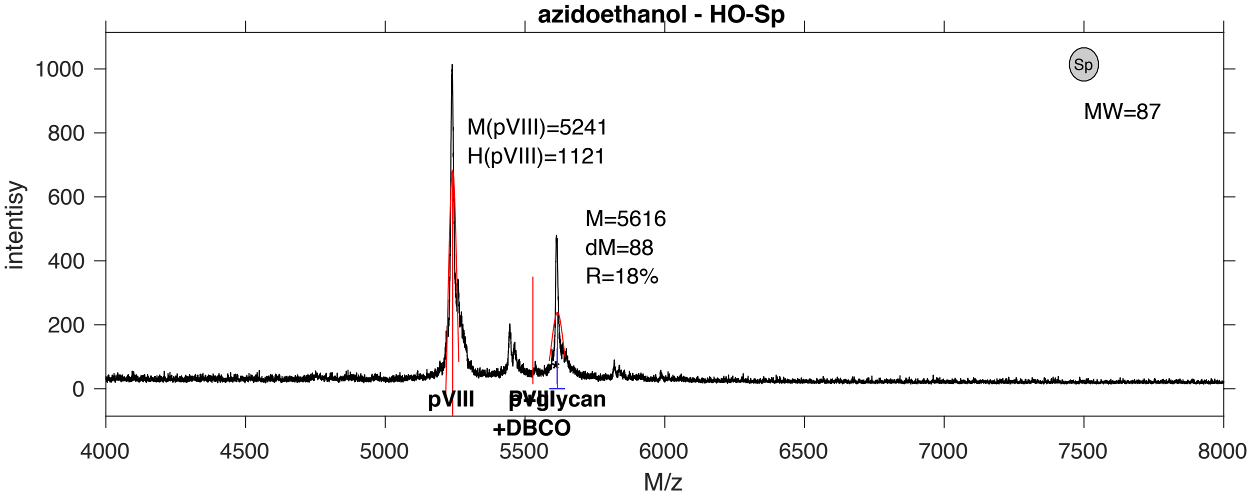


**SDB Number:** SDB183

**Sequencing File:** http://ligacloud.ca/searchLibInfo?f=0&b=0&d=NA

**Barcode:**
CTACTATTTGCGATCCCCCTAAGTGTAGAGAAAAATGATCAGAAAACCTACCACGCCGGAGGCGGT

**Axis Name:** AE-[490]

**IUPAC:** nonenone

**CFG Name:** HO-Sp

**Common Name:** AE

**Glytoucan ID:** none

**Compound Number:** azidoethanol

**Maldi File:** 382.txt

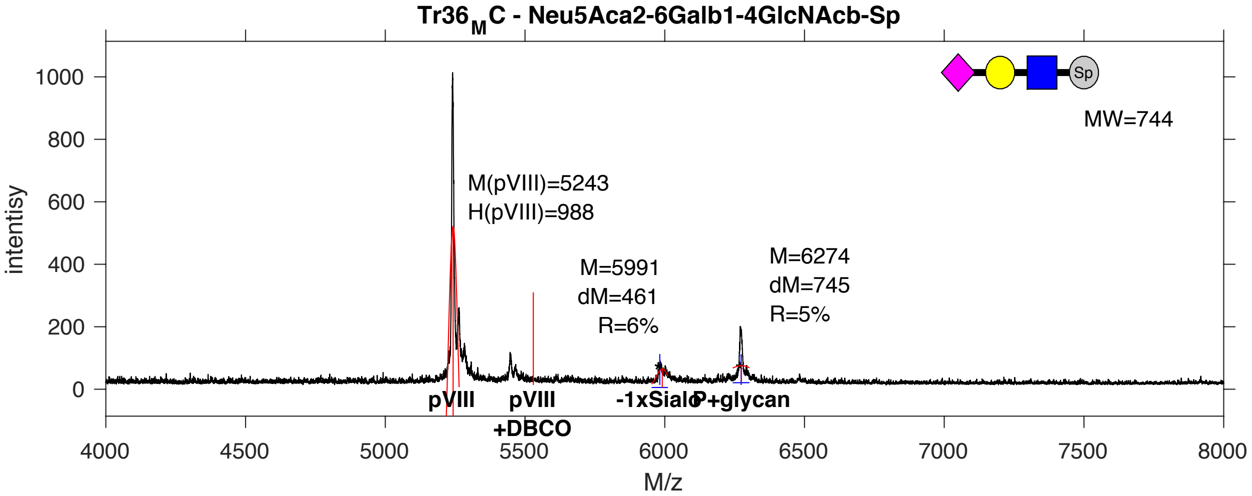


**SDB Number:** SDB177

**Sequencing File:** http://ligacloud.ca/searchLibInfo?f=0&b=0&d=20180809-87CLooPA-MR6

**Barcode:**
CTTCTATTTGCTATTCCTCTAAGCGTGGAGAAGAATGATCAAAAGACTTATCACGCTGGGGGGGGG

**Axis Name:** 6'SLN-[300]

**IUPAC:** Neu5Ac(a2-6)Gal(b1-4)GlcNAc(b1-Sp

**CFG Name:** Neu5Aca2-6Galb1-4GlcNAcb-Sp

**Common Name:** 6'SLN

**Glytoucan ID:** G73578JC

**Compound Number:** Tr36_MC

**Maldi File:** 383.txt

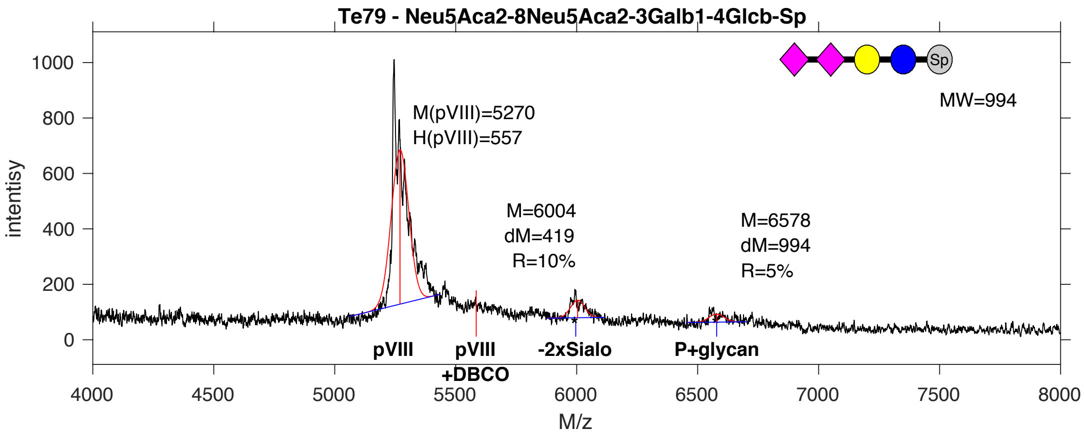


**SDB Number:** SDB45

**Sequencing File:** http://ligacloud.ca/searchLibInfo?f=0&b=0&d=20180809-87CLooPA1-1JM12

**Barcode:**
CTGCTGTTTGCGATTCCTCTGAGCGTGGAAAAAAATGACCAAAAAACCTACCATGCAGGGGGGGGA

**Axis Name:** GD3-[410]

**IUPAC:** Neu5Ac(a2-8)Neu5Ac(a2-3)Gal(b1-4)Glc(b1-Sp

**CFG Name:** Neu5Aca2-8Neu5Aca2-3Galb1-4Glcb-Sp

**Common Name:** GD3

**Glytoucan ID:** G07249ZL

**Compound Number:** Te79

**Maldi File:** 21-2.txt

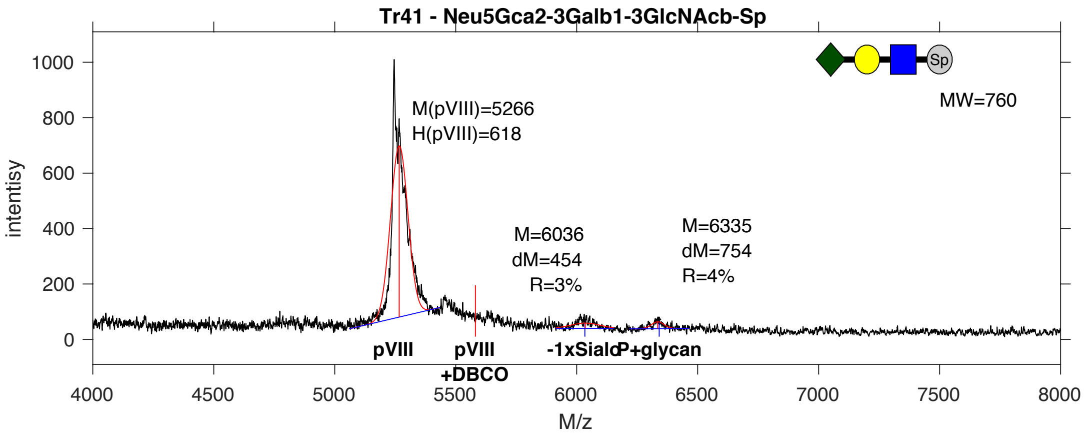


**SDB Number:** SDB74

**Sequencing File:** http://ligacloud.ca/searchLibInfo?f=0&b=0&d=20170807-87GYooOS1-2SS14

**Barcode:**
CTACTCTTTGCGATCCCTCTGAGTGTCGAAAAAAATGACCAAAAGACCTATCACGCGGGCGGGGGA

**Axis Name:** 3'SLec (Gc)-[190]

**IUPAC:** Neu5Gc(a2-3)Gal(b1-3)GlcNAc(b1-Sp

**CFG Name:** Neu5Gca2-3Galb1-3GlcNAcb-Sp

**Common Name:** 3'SLec (Gc)

**Glytoucan ID:** G01245TU

**Compound Number:** Tr41

**Maldi File:** 39-2.txt

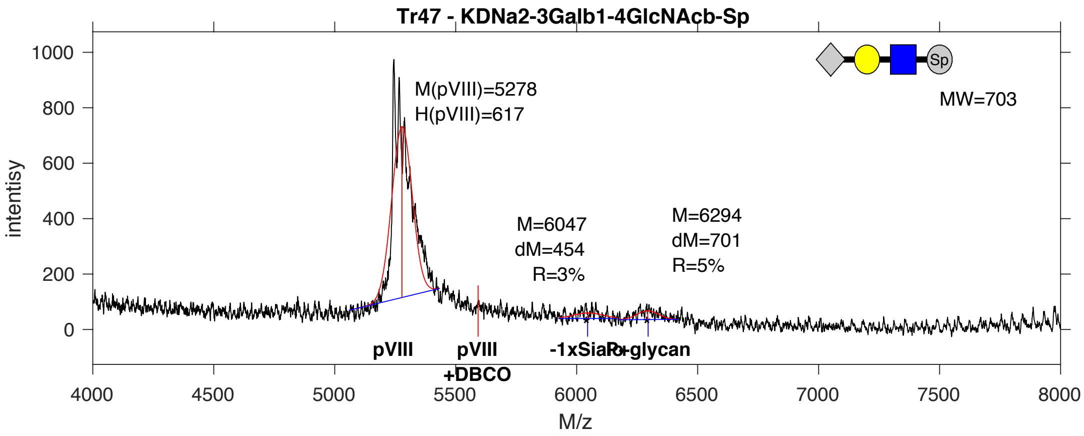


**SDB Number:** SDB87

**Sequencing File:** http://ligacloud.ca/searchLibInfo?f=0&b=0&d=20171128-87CLooOOPDDQ-SS

**Barcode:**
CTACTCTTCGCTATACCCCTCAGCGTAGAAAAAAATGACCAAAAGACCTACCATGCGGGAGGTGGG

**Axis Name:** 3'KDNLN-[220]

**IUPAC:** KDN(a2-3)Gal(b1-4)GlcNAc(b1-Sp

**CFG Name:** KDNa2-3Galb1-4GlcNAcb-Sp

**Common Name:** 3'KDNLN

**Glytoucan ID:** G61010HE

**Compound Number:** Tr47

**Maldi File:** 46-2.txt

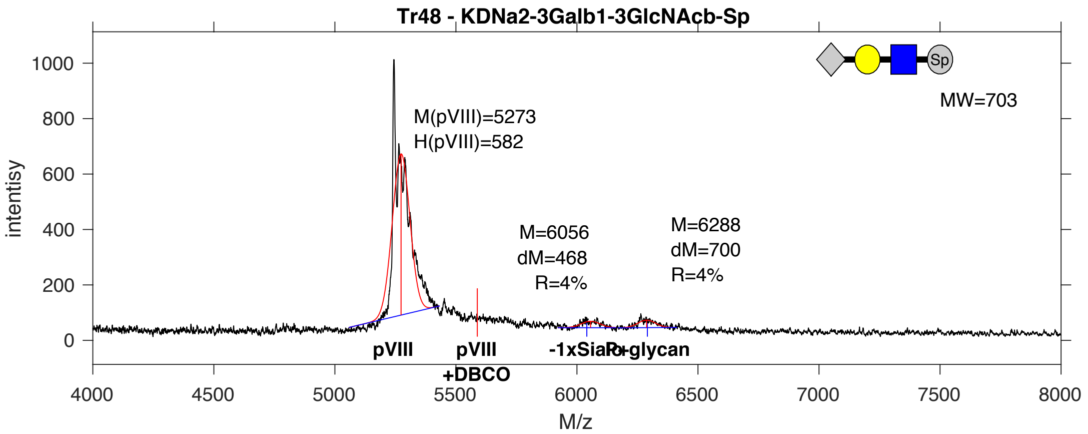


**SDB Number:** SDB111

**Sequencing File:** http://ligacloud.ca/searchLibInfo?f=0&b=0&d=NA

**Barcode:**
CTGCTGTTTGCTATCCCGCTAAGTGTAGAAAAAAATGACCAGAAAACGTACCATGCGGGCGGTGGG

**Axis Name:** 3’-KDNLec-[220]

**IUPAC:** KDN(a2-3)Gal(b1-3)GlcNAc(b1-Sp

**CFG Name:** KDNa2-3Galb1-3GlcNAcb-Sp

**Common Name:** 3’-KDNLec

**Glytoucan ID:** G13678CX

**Compound Number:** Tr48

**Maldi File:** 51-2.txt

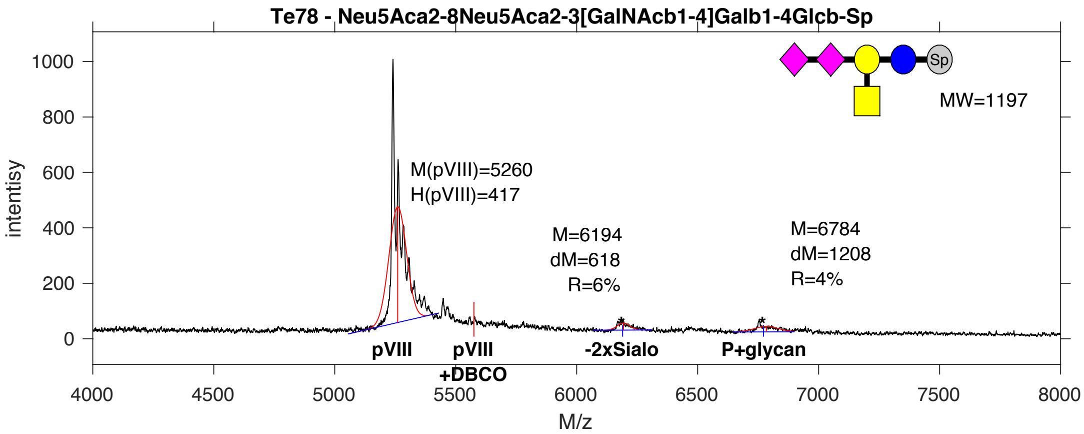


**SDB Number:** SDB122

**Sequencing File:** http://ligacloud.ca/searchLibInfo?f=0&b=0&d=20190703-87CLooOOUFCK-DF

**Barcode:**
CTGCTTTTTGCTATACCCCTGAGCGTGGAGAAGAACGATCAAAAGACCTATCACGCGGGGGGAGGC

**Axis Name:** GD2-[270]

**IUPAC:** Neu5Ac(a2-8)Neu5Ac(a2-3)[GalNAc(b1-4)]Gal(b1-4)Glc(b1-Sp

**CFG Name:** Neu5Aca2-8Neu5Aca2-3[GalNAcb1-4]Galb1-4Glcb-Sp

**Common Name:** GD2

**Glytoucan ID:** G79648LW

**Compound Number:** Te78

**Maldi File:** 57-2.txt

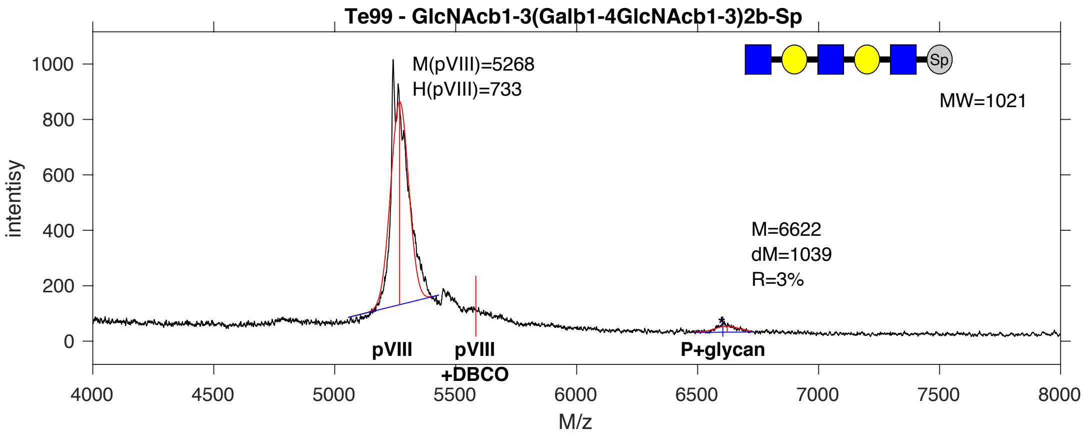


**SDB Number:** SDB150

**Sequencing File:** http://ligacloud.ca/searchLibInfo?f=0&b=0&d=NA

**Barcode:**
CTGCTCTTTGCTATCCCTCTAAGTGTTGAAAAAAACGACCAGAAGACTTATCACGCAGGTGGTGGG

**Axis Name:** 3’GN-Di-LN-[80]

**IUPAC:** GlcNAc(b1-3)Gal(b1-4)GlcNAc(b1-3)Gal(b1-4)GlcNAc(b1-Sp

**CFG Name:** GlcNAcb1-3(Galb1-4GlcNAcb1-3)2b-Sp

**Common Name:** 3’GN-Di-LN

**Glytoucan ID:** G73480MV

**Compound Number:** Te99

**Maldi File:** 60-2.txt

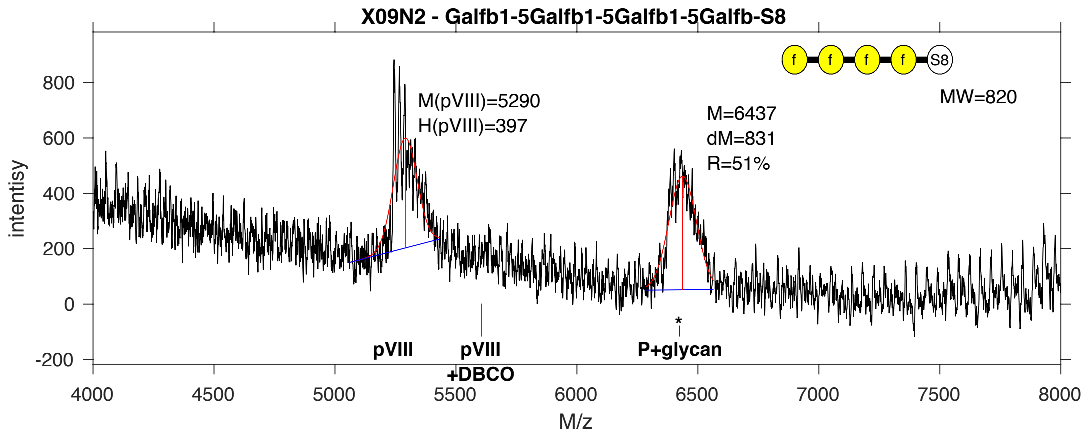


**SDB Number:** SDB6

**Sequencing File:** http://ligacloud.ca/searchLibInfo?f=0&b=0&d=20171128-87CLooOOPUXM-SS

**Barcode:**
CTGCTGTTTGCGATTCCACTGAGTGTGGAGAAGAATGATCAGAAGACTTATCATGCGGGTGGAGGT

**Axis Name:** (Galf)4-[1380]

**IUPAC:** Galf(b1-5)Galf(b1-5)Galf(b1-5)Galf(b1-S8

**CFG Name:** Galfb1-5Galfb1-5Galfb1-5Galfb-S8

**Common Name:** (Galf)4

**Glytoucan ID:** G92890KE

**Compound Number:** X09N2

**Maldi File:** 74-2.txt

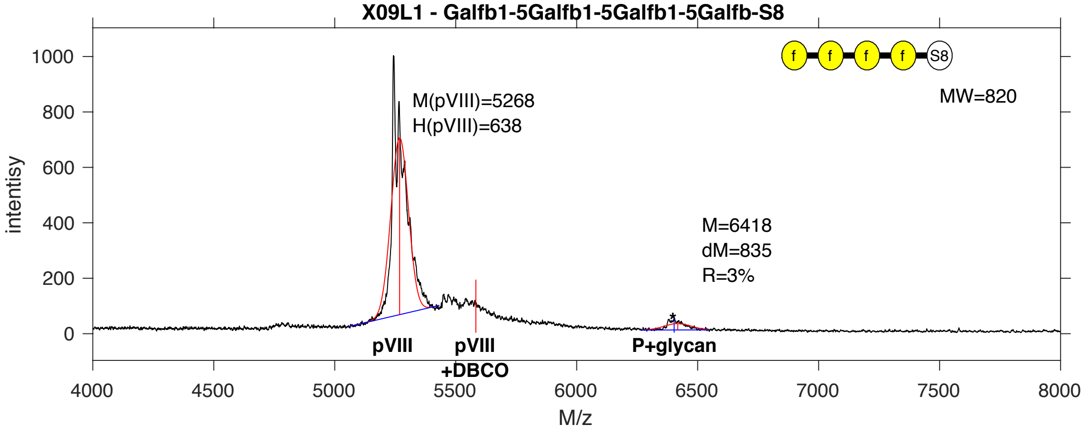


**SDB Number:** SDB10

**Sequencing File:** http://ligacloud.ca/searchLibInfo?f=0&b=0&d=20171128-87CLooOOPTIT-SS

**Barcode:**
CTACTGTTTGCTATACCGCTGAGTGTGGAGAAGAATGATCAGAAGACTTATCATGCGGGTGGAGGT

**Axis Name:** (Galf)4-[80]

**IUPAC:** Galf(b1-5)Galf(b1-5)Galf(b1-5)Galf(b1-S8

**CFG Name:** Galfb1-5Galfb1-5Galfb1-5Galfb-S8

**Common Name:** (Galf)4

**Glytoucan ID:** G92890KE

**Compound Number:** X09L1

**Maldi File:** 77-2.txt

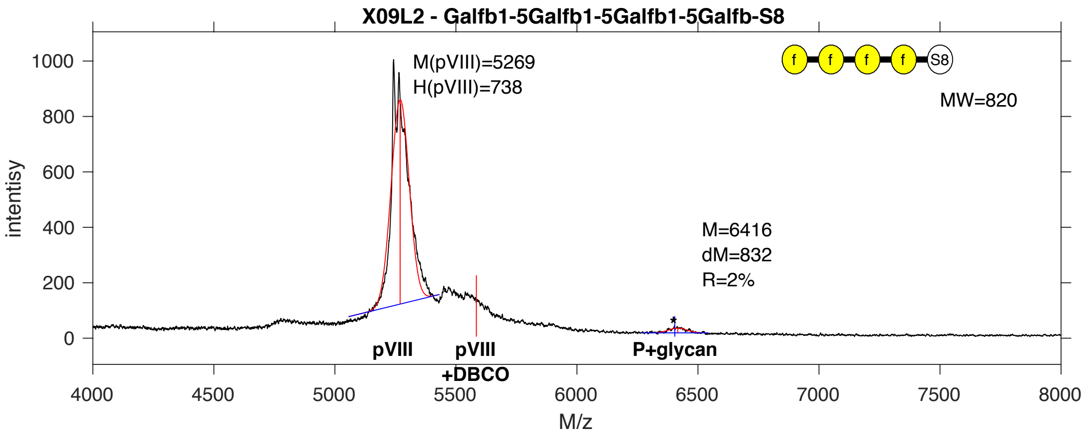


**SDB Number:** SDB11

**Sequencing File:** http://ligacloud.ca/searchLibInfo?f=0&b=0&d=20171128-87CLooOOZCHR-SS

**Barcode:**
CTACTATTTGCGATTCCCCTGAGTGTGGAGAAGAATGATCAGAAGACTTATCATGCGGGTGGAGGT

**Axis Name:** (Galf)4-[50]

**IUPAC:** Galf(b1-5)Galf(b1-5)Galf(b1-5)Galf(b1-S8

**CFG Name:** Galfb1-5Galfb1-5Galfb1-5Galfb-S8

**Common Name:** (Galf)4

**Glytoucan ID:** G92890KE

**Compound Number:** X09L2

**Maldi File:** 78-2.txt

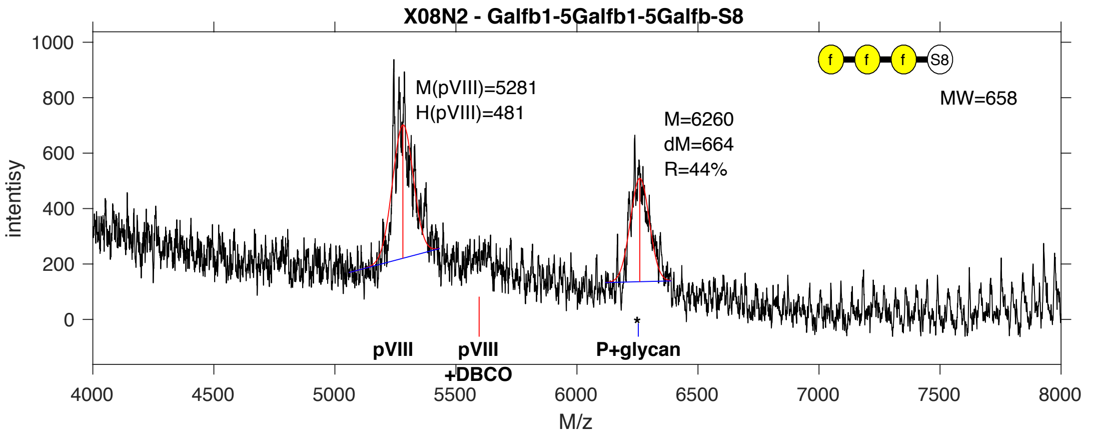


**SDB Number:** SDB27

**Sequencing File:** http://ligacloud.ca/searchLibInfo?f=0&b=0&d=NA

**Barcode:**
CTGCTTTTCGCTATTCCTCTCAGTGTGGAGAAGAATGATCAGAAGACTTATCATGCGGGTGGAGGT

**Axis Name:** (Galf)3-[1190]

**IUPAC:** Galf(b1-5)Galf(b1-5)Galf(b1-S8

**CFG Name:** Galfb1-5Galfb1-5Galfb-S8

**Common Name:** (Galf)3

**Glytoucan ID:** G99590HR

**Compound Number:** X08N2

**Maldi File:** 91-2.txt

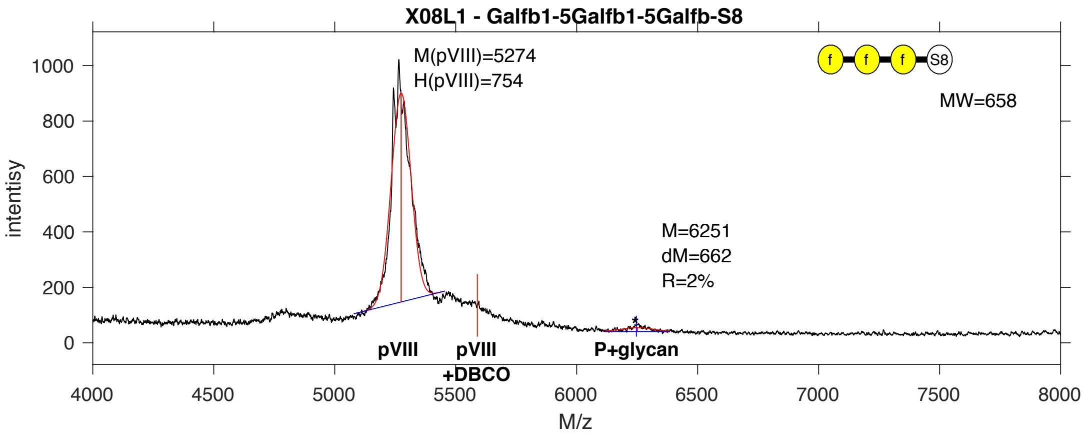


**SDB Number:** SDB34

**Sequencing File:** http://ligacloud.ca/searchLibInfo?f=0&b=0&d=20171128-87CLooOOTLAJ-SS

**Barcode:**
CTTCTTTTTGCGATTCCGCTGAGTGTGGAGAAGAATGATCAGAAGACTTATCATGCGGGTGGAGGT

**Axis Name:** (Galf)3-[50]

**IUPAC:** Galf(b1-5)Galf(b1-5)Galf(b1-S8

**CFG Name:** Galfb1-5Galfb1-5Galfb-S8

**Common Name:** (Galf)3

**Glytoucan ID:** G99590HR

**Compound Number:** X08L1

**Maldi File:** 98-2.txt

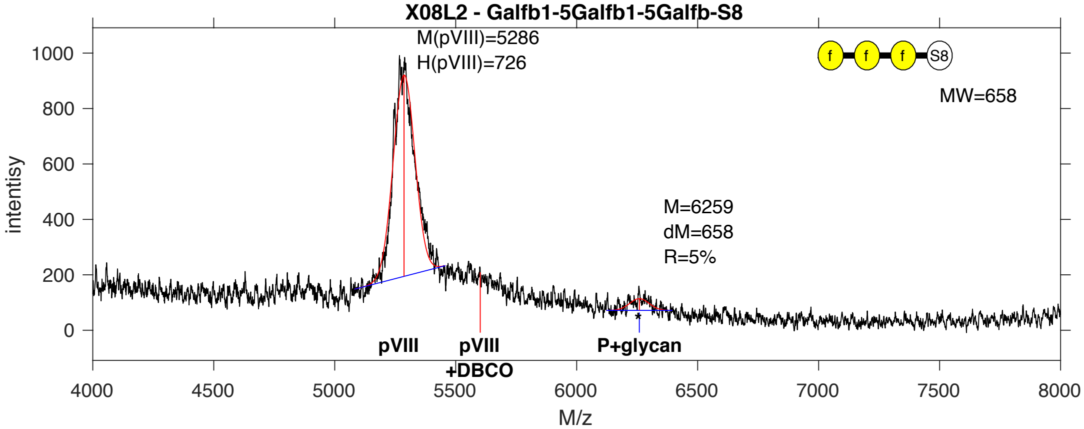


**SDB Number:** SDB35

**Sequencing File:** http://ligacloud.ca/searchLibInfo?f=0&b=0&d=20171128-87CLooOOBBJH-SS

**Barcode:**
CTGCTCTTCGCTATTCCACTTAGTGTGGAGAAGAATGATCAGAAGACTTATCATGCGGGTGGAGGT

**Axis Name:** (Galf)3-[140]

**IUPAC:** Galf(b1-5)Galf(b1-5)Galf(b1-S8

**CFG Name:** Galfb1-5Galfb1-5Galfb-S8

**Common Name:** (Galf)3

**Glytoucan ID:** G99590HR

**Compound Number:** X08L2

**Maldi File:** 99-2.txt

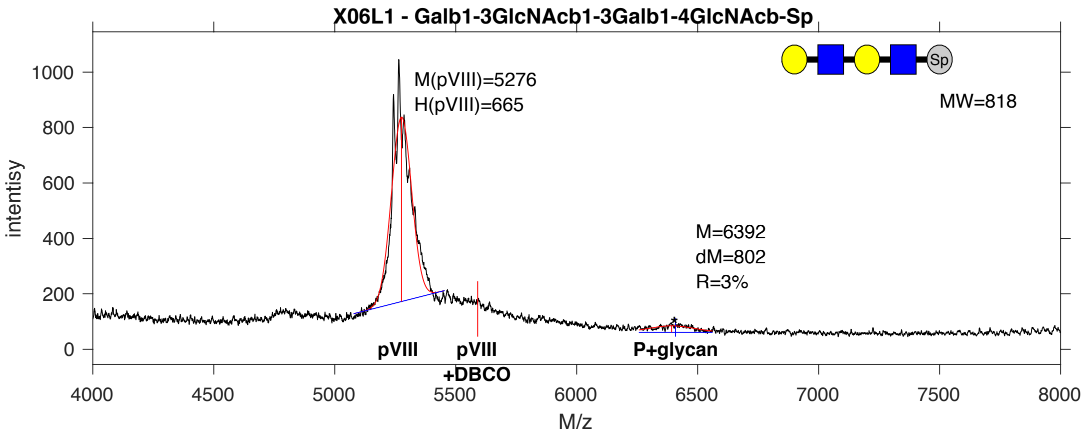


**SDB Number:** SDB70

**Sequencing File:** http://ligacloud.ca/searchLibInfo?f=0&b=0&d=20171128-87CLooOOJSQJ-SS

**Barcode:**
CTACTGTTCGCAATCCCGCTCAGTGTTGAAAAAAACGATCAAAAAACGTATCATGCTGGTGGAGGT

**Axis Name:** LNT-Nac-[80]

**IUPAC:** Gal(b1-3)GlcNAc(b1-3)Gal(b1-4)GlcNAc(b1-Sp

**CFG Name:** Galb1-3GlcNAcb1-3Galb1-4GlcNAcb-Sp

**Common Name:** LNT-Nac

**Glytoucan ID:** G09681LD

**Compound Number:** X06L1

**Maldi File:** 108-2.txt

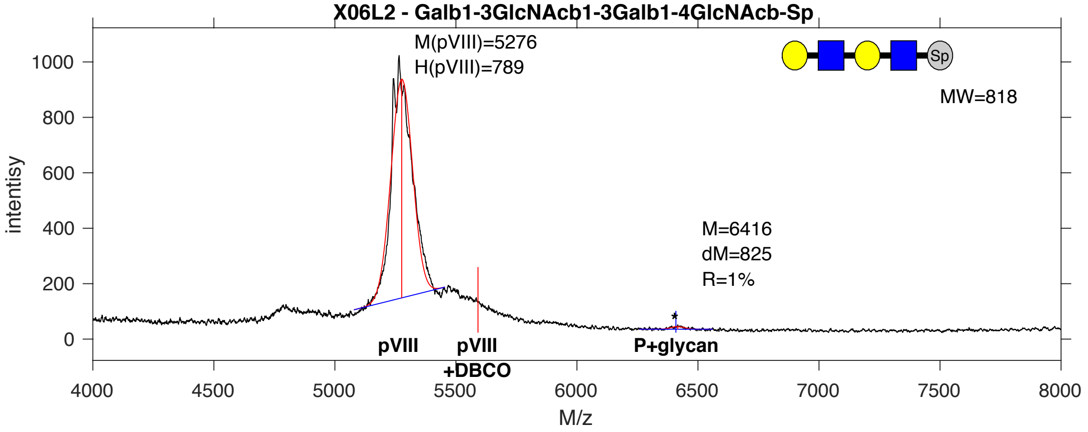


**SDB Number:** SDB71

**Sequencing File:** http://ligacloud.ca/searchLibInfo?f=0&b=0&d=20171128-87CLooOOBZUS-SS

**Barcode:**
CTGCTTTTTGCTATTCCTCTGAGTGTTGAAAAAAACGATCAGAAGACTTATCACGCGGGGGGCGGG

**Axis Name:** LNT-Nac-[30]

**IUPAC:** Gal(b1-3)GlcNAc(b1-3)Gal(b1-4)GlcNAc(b1-Sp

**CFG Name:** Galb1-3GlcNAcb1-3Galb1-4GlcNAcb-Sp

**Common Name:** LNT-Nac

**Glytoucan ID:** G09681LD

**Compound Number:** X06L2

**Maldi File:** 109-2.txt

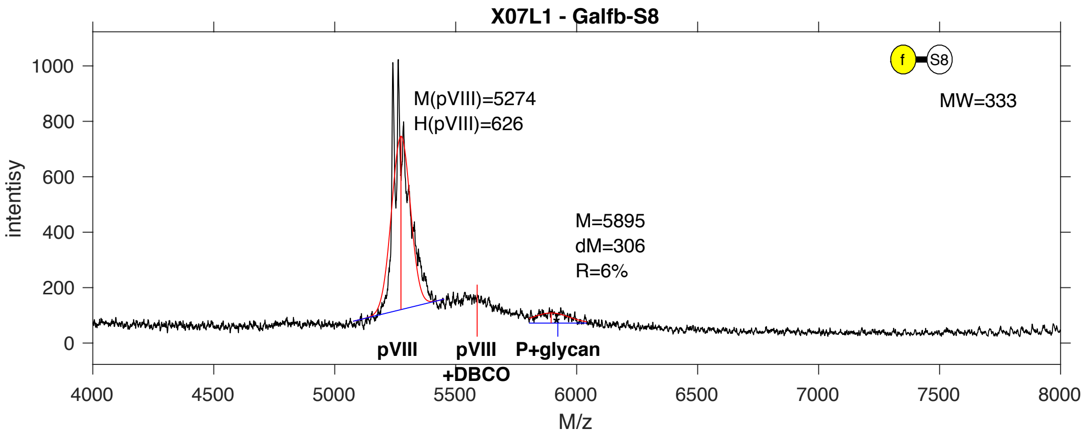


**SDB Number:** SDB84

**Sequencing File:** http://ligacloud.ca/searchLibInfo?f=0&b=0&d=20171128-87CLooOOPFPA-SS

**Barcode:**
CTACTCTTCGCGATCCCACTGAGCGTCGAGAAGAACGATCAGAAGACTTATCACGCTGGCGGCGGC

**Axis Name:** Galf-[160]

**IUPAC:** Galf(b1-S8

**CFG Name:** Galfb-S8

**Common Name:** Galf

**Glytoucan ID:** G38028NH

**Compound Number:** X07L1

**Maldi File:** 114-2.txt

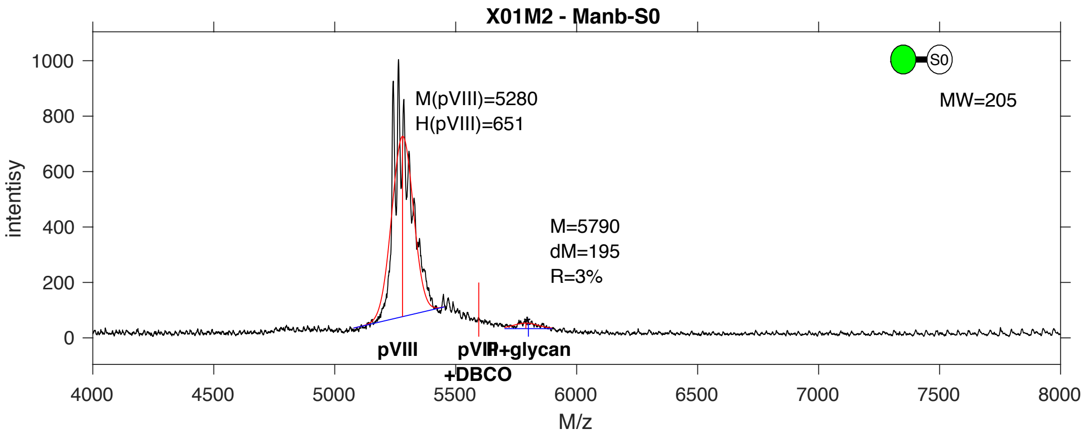


**SDB Number:** SDB125

**Sequencing File:** http://ligacloud.ca/searchLibInfo?f=0&b=0&d=20171128-87CLooOOEYOY-SS

**Barcode:**
CTGCTATTCGCTATCCCGCTGAGTGTAGAAAAAAACGACCAGAAAACCTATCATGCCGGTGGCGGG

**Axis Name:** bMan-[80]

**IUPAC:** Man(b1-S0

**CFG Name:** Manb-S0

**Common Name:** bMan

**Glytoucan ID:** G58490PL

**Compound Number:** X01M2

**Maldi File:** 121-2.txt
